# Quantifying and controlling the proteolytic degradation of cell adhesion peptides

**DOI:** 10.1101/2024.04.19.590329

**Authors:** Samuel J. Rozans, Abolfazl Salehi Moghaddam, Yingjie Wu, Kayleigh Atanasoff, Liliana Nino, Katelyn Dunne, E. Thomas Pashuck

## Abstract

Peptides are widely used within biomaterials to improve cell adhesion, incorporate bioactive ligands, and enable cell-mediated degradation of the matrix. While many of the peptides incorporated into biomaterials are intended to be present throughout the life of the material, their stability is not typically quantified during culture. In this work we designed a series of peptide libraries containing four different N-terminal peptide functionalizations and three C-terminal functionalization to better understand how simple modifications can be used to reduce non-specific degradation of peptides. We tested these libraries with three cell types commonly used in biomaterials research, including mesenchymal stem/stromal cells (hMSCs), endothelial cells, and macrophages, and quantified how these cell types non-specifically degraded peptide as a function of terminal amino acid and chemistry. We found that peptides in solution which contained N-terminal amines were almost entirely degraded by 48 hours, irrespective of the terminal amino acid, and that degradation occurred even at high peptide concentrations. Peptides with C-terminal carboxylic acids also had significant degradation when cultured with cells. We found that simple modifications to the termini could significantly reduce or completely abolish non-specific degradation when soluble peptides were added to cells cultured on tissue culture plastic or within hydrogel matrices, and that functionalizations which mimicked peptide conjugations to hydrogel matrices significantly slowed non-specific degradation. We also found that there were minimal differences across cell donors, and that sequences mimicking different peptides commonly-used to functionalized biomaterials all had significant non-specific degradation. Finally, we saw that there was a positive trend between RGD stability and hMSC spreading within hydrogels, indicating that improving the stability of peptides within biomaterial matrices may improve the performance of engineered matrices.

## Introduction

Hydrogels show great promise for mimicking the extracellular matrix (ECM) that surrounds cells^1^ and are frequently used to improve our understanding of basic physiological processes and the efficacy of regenerative therapies.^2^ Cells have a dynamic relationship with their microenvironment and cell behavior within the ECM is influenced by the presence of adhesion domains present in the local matrix.^3^ Cells actively modulate the local microenvironment by secreting biomolecules, including enzymes, that modify their local niche.^4,5^ This includes proteases, a class of over 600 enzymes that cleave proteins and peptide bonds.^6^ Proteases are expressed by all human cell types^7^ and play important roles in basic physiological processes,^6^ including cell spreading and migration.^8,9^ Individual proteases can be highly specific, cleaving only certain peptide sequences, or broadly degrade the termini of proteins or peptides.^10^

It is well-understood that non-specific degradation of therapeutic peptide drugs significantly limits their clinical potential.^11^ However, bioactive peptides are frequently incorporated into biomaterials to better mimic native ECMs,^3,12,13^ but typically without any characterization of their stability during culture. This includes the canonical RGD cell adhesion peptide^14^ and also signaling peptides which mimic growth factors, among other sequences.^15,16^ These peptides are generally intended to be present throughout the lifetime of the material and any degradation is undesirable and could reduce the efficacy of the biomaterial system. Cell-secreted proteases, such as MMPs^17^ or cathepsins,^18^ are often used as stimuli to modify biomaterials.^19^ However, MMPs and cathepsins are only a fraction of the total number of proteases expressed by cells.^7^ The enzymes harnessed within bioengineered systems, including MMPs and cathepsincs, are almost exclusively endopeptidases,^19,20^ which cleave interior of peptides and protein sequences. Exopeptidases, another class of proteases which cleave amino acids at the termini of peptides and proteins (**Fig. 1A**), are ubiquitously expressed in human tissues,^7^ and have been used to tailor the adhesion environment within biomaterials.^21^ Notably, exopeptidases are believed to be largely responsible for the rapid degradation of peptide drugs,^22,23^ and modifications to the N-terminus of peptides can slow non-specific degradation.^22,24^ Modification of the N-terminus of proteins is widespread, and more than 80% of human proteins have acetylated N-termini.^25^ The exopeptidase family of proteases includes both aminopeptidases that are active against the N-terminus of peptides and carboxypeptidases that are active against the C-terminus, and the extent to which these classes of proteases degrade peptides is known to be heavily dependent upon the chemistry of the termini.^26^

**Fig. 1.**
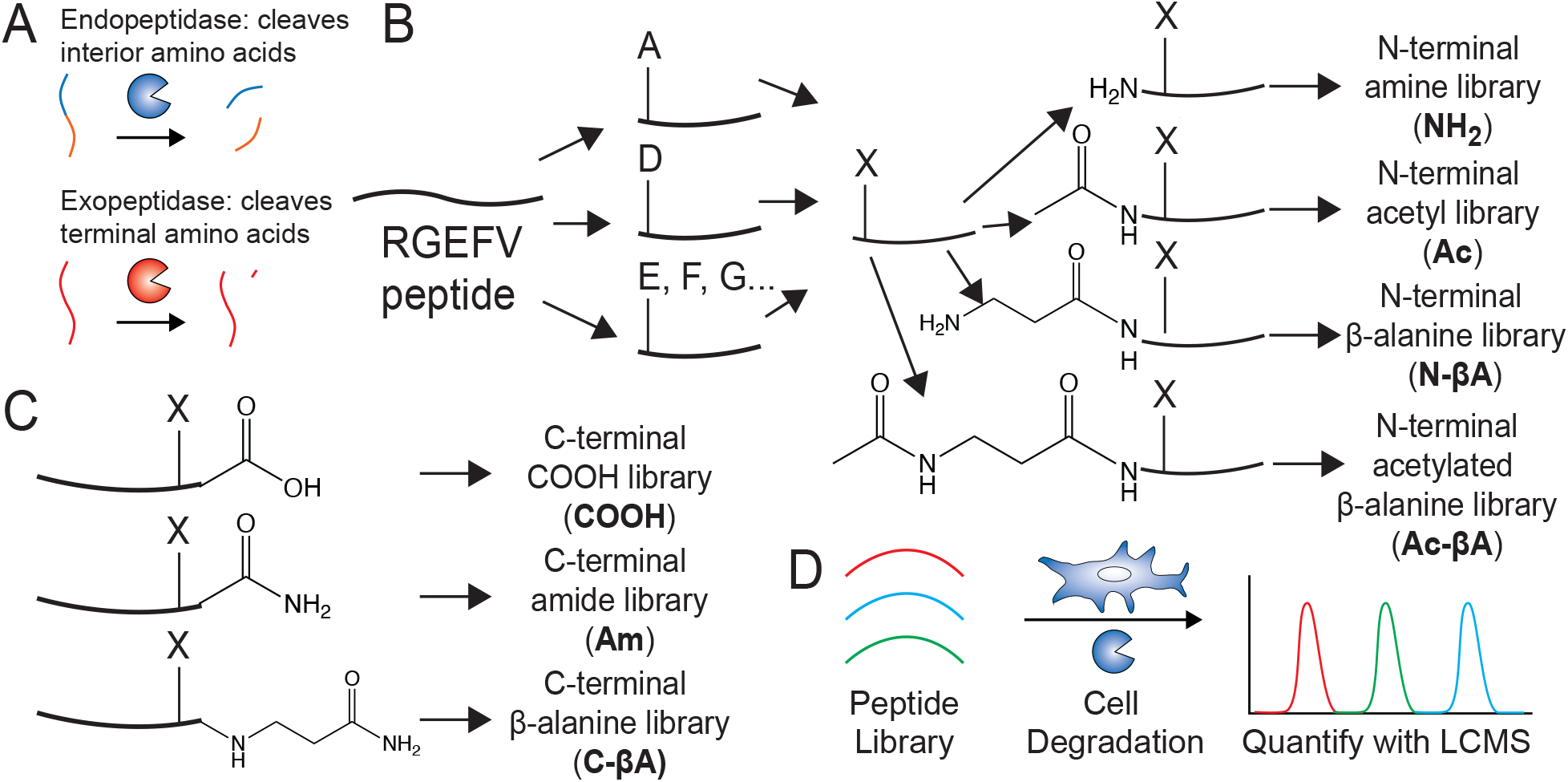
Synthetic scheme to develop design rules to understand non-specific degradation of peptides. **a**. Proteases can be classified as either endopeptidases, which cleave on the interior of proteins and peptides, or exopeptidases, which cleave terminal amino acids. **b**. A model cell adhesion peptide was split 19 ways during synthesis and every canonical amino acid except cysteine was added to the N-terminus. The reactions were then pooled together, and the N-terminus was then split again to generate four libraries with different N-terminal modifications. **c**. This process was also performed on the C-terminus of the model peptide. **d**. These peptide libraries were incubated with three different cell types, mesenchymal stem cells, endothelial cells, and macrophages, and the extent of peptide degradation was quantified using LCMS.

In this work we synthesized a series of peptide libraries to better understand how cell-secreted proteases degrade bioactive peptides. Peptide libraries were synthesized with a range of N- and C-terminal modifications to all canonical amino acids except cysteine. Peptides with N-terminal amines were rapidly degraded by human mesenchymal stem/stromal cells (hMSCs), human umbilical vein endothelial cells (hUVECs), and pro-inflammatory macrophages across almost all terminal amino acids. However, simple modifications could significantly reduce this non-specific degradation across most cell types. These results offer generalized design rules to improve the design and bioactivity of peptide-functionalized hydrogels.

## Methods and Materials

### Materials

All peptide synthesis reagents were purchased from Chemscene or Ambeed. N,N-Dimethylformamide (DMF) and dichloromethane (DCM) (both from VWR BDH Chemicals), piperidine (Millipore Sigma), trifluoroacetic acid (Millipore Sigma), diethyl ether (Fisher Scientific), N, N-Diisopropylethylamine (DIPEA) (VWR), were used as purchased. 20 kDa 8-arm poly(ethylene glycol) dibenzocyclooctyne (PEG-DBCO) was purchased from Biopharm PEG. Batches which dissolved in water in under 30 seconds were used as received. Otherwise they were quickly purified by dissolving them in isopropanol and removing impurities using a 10 kDa Amicon centrifugal filter. Approximately 200 mg of PEG was dissolved in 5 mLs of isopropanol and it was centrifuged at 4,500 RPM until there was less than one milliliter of PEG-DBCO/ isopropanol left in the filter. This was repeated twice, and then the filtrate was lyophilized and used.

### Methods

#### Peptide Synthesis Procedure

Peptides were synthesized using standard solid phase peptide synthesis (SPPS) protocols using either manual synthesis or an automated peptide synthesizer (CEM Liberty Blue) using standard Fmoc-protected amino acids (Chemscene) on a Rink amide resin (Supra Sciences) unless otherwise noted. All amide couplings were done using O-(6-chlorobenzotriazol-1-yl)-N,N,N′,N′-tetramethyluronium hexafluorophosphate (HCTU) in DMF unless otherwise noted. For each coupling the amino acid, HCTU, and DIPEA were added in a 4:4:6 molar ratio to the peptide. During peptide synthesis a ninhydrin test was performed after every addition to test for the presence of free amines. Upon a positive test, the coupling was replicated until the test was negative. A capping step was then performed with acetic anhydride (Sigma-Aldrich) in a 10:5:100 acetic anhydride:DIPEA:DMF solution twice for 5 min, and then a ninhydrin test was performed to check for complete capping of the free amines. After successful coupling, the Fmoc group was removed, washing the resin with 20% piperidine in DMF twice for 5 min. A ninhydrin test was performed to check for a positive result.

For split-and-pool steps the resin was washed 3× with DMF and then the entire amount of resin was weighed on a scale. This number was divided by approximately 22 and this was split 19 ways into 19 separate tubes, with some excess to account for resin loss during transport and weighing. The reactions were performed in 15 mL tubes and upon successful coupling of all 19 amino acids all 19 fractions of resin were re-combined. Peptide libraries attached to the C-terminal end were completed in one vessel. Peptide libraries attached to the N-terminal end were washed with DMF 3X, weighed and split four ways followed by adding the desired chain-end chemistry. Once the N-terminal libraries were split four ways, they were not recombined again. Peptides with C-terminal carboxylic acids were synthesized using 2-chlorotrityl chloride resin. For the COOH library 2 mmol of resin was weight out and 52.6 µmol of each amino acid (1 mmol total synthesis scale) was added into a single 50 mL tube. 25 mL of DCM was added and then DMF was added dropwise until all of the amino acids went fully into solution. The 2-chlorotrityl chloride resin was washed with DCM for five minutes to swell the resin, at which time the DCM was drained and the amino acid solution was added with 5 equivalents of DIPEA. After 30 minutes another 1.5 equivalents of DIPEA was added. After another 30 minutes 5 mL of methanol was added to cap the resin, at which time the Fmoc groups were deprotected and the peptides were synthesized using standard Fmoc-peptide synthesis procedures.

Peptides libraries and all peptides containing tryptophan were cleaved using 92.5% trifluoroacetic acid (TFA), 2.5% H_2_O, 2.5% triisopropylsilane (TIPS), and 2.5% dithiothreitol (DTT). Peptides which lacked a tryptophan were cleaved without DTT. Peptides were typically cleaved for 2-3 hours at room temperature using approximately 25 mL of cleavage solution per mM of peptide. However, peptides containing azides were cleaved for 30 minutes to prevent degradation of the azide group. Peptide mass was checked using electrospray ionization and if protecting groups remained the peptide was re-cleaved for 30 minutes. The masses of the peptide libraries were checked using liquid chromatography mass spectrometry (LCMS) and if protecting groups remained the peptides were re-cleaved for 30 minutes. At the end of the cleavage the peptides were precipitated in diethyl ether. These were then centrifuged for 5 minutes at 4,000 rpm, and the supernatant was discarded. The peptide pellet was washed with diethyl ether, and centrifuged, and this was repeated twice. The peptide pellet was allowed to dry, and then dissolved in water and neutralized with ammonium hydroxide prior to purification.

The cyclic RGD peptide was synthesized on a 2-chlorotrityl chloride resin. The first amino acid (0.3 mmol of amino acid per gram of resin) was dissolved in dichloromethane (DCM) and added to the resin in a shaker vessel. Five equivalents of DIPEA was then added, and after 5 min of shaking another 1.5 equivalents of DIPEA was added. After 1 h the unreacted 2-chlorotrityl chloride resin was capped with an excess of methanol for 30 min with another five equivalents of DIPEA. The rest of the amino acids were then coupled using standard solid phase Fmoc-synthesis protocols. After the last Fmoc group was deprotected the resin was washed 3× in DMF and 3× in DCM. The peptide was then cleaved under mild acidic conditions consisting of 5% trifluouroacetic acid and 2.5% triisopropylsilane in DCM. The mild cleavage solution was added to the resin for 5 minutes and then collected into a round bottom flask and this was repeated until the resin turned dark red or black. The collected liquid was then precipitated in ether. This cyclization was then dissolved in 50:50 acetonitrile:water, neutralized with 1M NH_4_OH, and lyophilized. The protected linear NH_2_-GRGDSK(N_3_)-OH peptide was then cyclized. This was done by dissolving the peptide into DMF at 1 mg/mL and adding 3 equivalents of (1-[bis(dimethylamino)methylene]-1H-1,2,3-triazolo[4,5-b]-pyridinium 3-oxid hexafluorophosphate) (HATU), and three equivalents of DIPEA. After six hours the DMF was removed using rotary evaporation at 65°C.

The PanMMP crosslinking peptide was functionalized with 2-azido acetic acid on the N-terminus. 2-azido acetic acid was synthesized by mixing bromoacetic acid (70.168 g, 505 mmol) and sodium azide (32.504 g, 500 mmol) and water (250ml), the solution was stirred overnight at RT under ambient conditions. The next day the solution was acidified to pH ∼ 1 using hydrochloric acid and extracted using ethyl acetate (5 × 100 mL). The organic layers were combined and dried in vacuo to afford the 2-azidoacetic acid as a colorless liquid. 2-azidoace+c acid was stored in a −20°C freezer until needed for synthesis.

All peptides were purified using HPLC using a Phenomenex Gemini 5 µm NX-C18 110 Å LC Column 150 x 21.2 mm. Gradients were run from 95% Mobile Phase A (water with 0.1% TFA) and 5% Mobile Phase B (acetonitrile with 0.1% TFA) to 100% Mobile Phase B. A typical HPLC run featured a two-minute equilibration step, followed by a 10 minute ramp from 95% Mobile Phase A to 100% Mobile Phase B, and then two minutes of equilibration at 100% Mobile Phase B, before ramping back down to the starting conditions. Notably, the split-and-pool libraries were ramped up to 100% Mobile Phase B over two minutes, since these libraries consisted of approximately 19 different peptides which weren’t intended to be separated from each other. The protected cyclic RGD peptides was purified using a gradient that ramped from 30% Mobile Phase B to 100% Mobile Phase B. After purification all peptides were lyophilized and were ready to use, except for the protected cyclic RGD peptide which was deprotected using 95% TFA, 2.5% TIPS, and 2.5% H_2_O for one hour.

PEGylated RGEFV peptides were prepared using standard SPPS as described above. Immediately adjacent to the chain-end chemistry of interest, on the N or C terminus of the peptide, an Fmoc-Azidolysine-OH (AR001RXM, Aaron Chemicals) was added. Once seven variations of the azidolysine functionalized peptides were synthesized, HPLC purified, dissolved in water at 10mM, and verified via LCMS, and 20 mg of peptide was transferred to new Eppendorf tubes. 100 mg of m-dPEG_12_-DBCO (QBD-10596, Vector Laboratories) was dissolved in water and evenly distributed 7 ways, amongst all azide-functionalized RGEFV peptides. The strained alkyne was allowed to click to the azide-functionalized peptides overnight. The next day, the pegylated peptides were HPLC purified, validated via LCMS, and lyophilized until needed.

#### Peptide Degradation Studies

##### hUVECs on Tissue culture plastic (TCP)

Passage 2 of human umbilical vein endothelial cells (hUVECs) (Life Technology) were seeded into 24 well plates at a seeding density of 150,000 cells per well in 1ml of expansion media (Lifeline Cell Technology, LM-0002) containing ascorbic acid, hydrocortisone, FBS, L-Glutamine, rh-EGF, heparin, and EnGS-US (All supplements LifeFactors, LS-1122). After 24 hour the media was changed. Peptide libraries, each containing 19 peptides and the non-proteolytically degradable βFβAβAβAβAβA-NH_2_ internal standard, were added to the cell media for a final concentration of 37 µM per peptide. Each library was tested in triplicate per per cell type per study. 40 µl samples were collected from the media at hours 0, 1, 4, 8, 24, and 48. In-between timepoints samples were frozen −80°C. After 48 hours, samples were kept frozen and thawed just prior to LCMS. The following donors were used for these studies: Lot 08119 from a Caucasian male, Lot 08478 from a Caucasian/African American female, and Lot 04608 from an African American male. Initial degradation studies with cells on tissue culture plastic were done using all three donors, and subsequent studies were done using Lot 04608.

##### hMSCs on TCP

Passage 3 of human mesenchymal stem cells (hMSCs) (Rooster Bio) were seeded into 24 well plates at a seeding density of 75,000 cells per well in 1ml of RoosterBasal™-MSC-CC (RoosterBio, SU-022) containing RoosterBooster™-MSC (RoosterBio SU-003). After 24 hour the media was changed and peptide libraries were added to the cell media for a final concentration of 37 µM per peptide. Each library was tested simultaneously, in triplicate per cell lot for a total of 63 wells. 40 µl samples were collected from the media at hours 0, 1, 4, 8, 24, and 48. In-between timepoints samples were frozen −80°C. After 48 hours, samples were kept frozen and thawed just prior to LCMS. The following donors were used for these studies: Lot 310264 a 20 year old African-american male, Lot 310268 a 19 year old Eritrean/east african female, and Lot 210280 Asian 26 year old male. Initial degradation studies with cells on tissue culture plastic were done using all three donors, and subsequent studies were done using Lot 310268.

##### PBMC Derived Macrophages on TCP

Peripheral blood mononuclear cells (PBMCs) were purchased from Allcells placed in a 24 well plate with a seeding density of one million cells per well using RPMI (Cytiva, SJ30027.1) containing 10% FBS (Foundation Fetal Bovine Serum, Gemini Bioproducts) and 1% antibiotic-antimycotic (gibco, 15240-062). Human M-CSF (PEPROTECH, 300-25-50UG) was immediately added at a concentration of 20ng/ml for five days to induce differentiation to M0 macrophages. After day 5 the media was changed to RPMI containing interferon gamma (IFN-γ) (PEPROTECH, 300-02-20UG), and 100 ng/ml Lipopolysaccharide (LPS) (Sigma L4391-1MG) and allowed to polarize for three days (Day 8 of culture). On Day 9, the media was changed to macrophage serum free media with L-Glutamine (Gibco 12065-074). Peptide libraries were added 24 hours later to the cell media for a final concentration of 37 µM per peptide. Each library was tested simultaneously, in triplicate per cell lot for a total of 63 wells. 40 µl samples were collected from the media at hours 0, 1, 4, 8, 24, and 48. In-between timepoints samples were frozen −80°C. After 48 hours, samples were kept frozen and thawed just prior to LCMS. Three donors and one cell line was used for these studies. The donors came from Lot 3087423 a 26-year-old Asian male, Lot 3088202 a 52 year old White male, and Lot 3091412 a 21 year old African-american Female. The primary PBMC-derived macrophages were used for initial studies on tissue culture plastic.

##### THP1 derived macrophages on TCP

THP1 cells were placed into a 24 well plate with a seeding density of one million cells per well using RPMI (Cytiva, SJ30027.1) containing 10% FBS and 1% anti-anti, with PMA at a concentration of 100ng/ml for two days. After day 2 the media was changed to RPMI containing IFN-γ, 20 ng/ml MCSF, and 100 ng/ml LPS and allowed to polarize for three days (Day 5 of culture). On day six, the media was changed to macrophage serum free media with L-Glutamine (Gibco 12065-074) Peptide libraries were added 24 hours later to the cell media for a final concentration of 37 µM per peptide. Each library was tested simultaneously, in triplicate for a total of 21 wells. 40 µl samples were collected from the media at hours 0, 1, 4, 8, 24, and 48. In-between timepoints samples were frozen −80°C. After 48 hours, samples were kept frozen and thawed just prior to LCMS. The THP-1 cell line was used for initial degradation experiments on tissue culture plastic and all other cell studies.

##### hUVECs in PEG

Passage 2 hUVECs were seeded into T75 flasks in basal media (LIFELINE CELL TECHNOLOGY, LM-0002) containing ascorbic acid, hydrocortisone, FBS, L-Glutamine, rh EGF, heparin, and EnGS-US (All supplements - LifeFactors, LS-1122) until they were 80-90% confluent. Cells were then washed with PBS and trypsinized with 2 mL of 0.25% Trypsin in HBSS with EDTA (Cytiva, SH30042.01) and incubated for 5 minutes. Once cells have detached they’re centrifuged at 0.2 RCF for 5 minutes and counted using a hemocytometer. At this time they were seeded into 28 µl of 3.5% (W/V) 8-arm-PEG-DBCO (MW 20,000 KDa) hydrogels at a density of 150,000 cells per gel with 35% of PEG arms crosslinked with a PanMMP peptide N_3_KGPQGIWGQKK(N_3_), and 10% of the arms are tethered with cyclic GRGDSK(N_3_) using copper free click chemistry of the strained alkyne DBCO with the azides. After 24 hours peptide libraries were added to the cell media for a final concentration of 37 µM per peptide. Each library was tested simultaneously, in triplicate for a total of 21 wells. 40 µl samples were collected from the media at hours 0, 1, 4, 8, 24, and 48. In-between timepoints samples were frozen −80°C. After 48 hours, samples were kept frozen and thawed just prior to LCMS.

##### hMSC in PEG

Passage 3 of hMSCs were seeded into T-75 flasks until they 80-90% confluent. Cells were then washed with PBS and trypsinized in 2 mL of trypsin and incubated for 5 minutes. Once cells have detached they were centrifuged at 0.2 RCF for 5 minutes and counted using a hemocytometer. At this time they werere seeded into 28 µl of 3.5% (W/V) 8-arm-peg-DBCO (MW 20,000 KDa) hydrogels at a density of 75,000 cells per gel with 35% of PEG arms crosslinked with a PanMMP peptide N_3_KGPQGIWGQKK(N_3_), and 10% of the arms are tethered with cyclic GRGDSK(N_3_) using copper free click chemistry of the strained alkyne DBCO with the azides. After 24 hours peptide libraries were added to the cell media for a final concentration of 37 µM per peptide. Each library was tested simultaneously, in triplicate for a total of 21 wells per study. 40 µl samples were collected from the media at hours 0, 1, 4, 8, 24, and 48. In-between timepoints samples were frozen −80°C. After 48 hours, samples were kept frozen and thawed just prior to LCMS.

##### THP1 derived macrophages in PEG

THP1 cells were placed into untreated six well plates with RPMI (Cytiva, SJ30027.1) containing 10% FBS and 1% anti-anti, with PMA at a concentration of 100ng/ml for two days. After day 2 the media was changed to RPMI containing IFN-γ, MCSF, and LPS and allowed to polarize for three days (Day 5 of culture). On day six, the media was changed to fresh RPMI and detached using a cell scraper. They were then centrifuged at 0.2 RCF for 5 minutes, counted using a hemocytometer and seeded into 40uL gels at a density of 1million cells per gel containing 28 uL of 3.5% (W/V) 8-arm-PEG-DBCO (MW 20,000 KDa) hydrogels at a density of 75,000 cells per gel with 35% of PEG arms crosslinked with a PAN-MMP sensitive peptide N_3_KGPQGIWGQKK(N_3_), and 10% of the arms are tethered with cyclic GRGDSK(N_3_) using copper free click chemistry of the strained alkyne DBCO with the azides. After 24 hours peptide libraries were added to the cell media for a final concentration of 37 µM per peptide. Each library was tested simultaneously, in triplicate for a total of 21 wells. 40 samples were collected from the media at hours 0, 1, 4, 8, 24, and 48. In-between timepoints samples were frozen −80°C. After 48 hours, samples were kept frozen and thawed just prior to LCMS.

### LCMS Data Acquisition and Analysis

4 µL of neat acetic acid were added to each well immediately after the plates were removed from the −80°C freezer to acidify the media and prevent further peptide degradation by proteases. From each sample, 10 µL of crude solution was introduced by the LC-MS through an Thermo Scientific™ Vanquish™ LC System (Thermo Fisher Scientific) which outputted to a Thermo Scientific™ LTQ XL™ Linear Ion Trap Mass Spectrometer (Thermo Fisher Scientific). The sampled mixture was trapped on a column (ProntoSIL C18 AQ, 120 Å, 3 µm, 2.0 x 50 mm HPLC Column, PN 0502F184PS030, MAC-MOD Analytical Inc.). The samples were loaded onto the column with a solvent containing acetonitrile/water, 5:95 (v/v) containing 1% Acetic Acid at a flow rate of 300 µL/min and held for one minute. The sample was then eluted from the column with a linear gradient of 5-40% Solvent B (1% Acetic acid in Acetonitrile) at the same flow rate for five minutes. This was followed by a 1 min ramp up to 100% Solvent B, where it was re-equilibrated with Solvent A (1% Acetic Acid) to 5% solvent B over the course of 1 min and held there for 2 min. The column temperature was a constant 29 °C. The Mass Spectrometer was operated in positive ion mode. Using a Heated ESI, the Source Voltage was set to 4.1 kV, and the capillary temperature was 350 °C.

Data Analysis was performed on Xcalibur Software (Thermo Scientific). Peptides were identified automatically using the Thermo Xcalibur Processing Setup window where the mass (m/z) ±0.5 AMU and expected retention times for each quantified peptide were inputted. This was later used to isolate individual peptide species from the total ion count (TIC) trace using the Thermo Xcalibur Quan Setup window, where the area under the curve was calculated and visually inspected for accuracy. All peptides were normalized to a non-changing internal standard, NH_2_-βFβAβAβAβAβA-NH_2_, where βA is a β-alanine and β-homophenylalanine. The integrated peak area of the peptide of interest was divided by the βFβAβAβAβAβA-NH_2_ internal stadard to create an area ratio. Relative amounts of a peptide of peptide were then calculated by normalized all values to their corresponding time zero area ratio. Calculated data was visualized using RStudio.

### Viability Assay

Cells were cultured in gels as described. A working solution was made by diluting the alamarBlue dye (Y00-100, Thermo Scientific) (1:10) with the same media cells were cultured in. The media the cells were in was replaced with one ml of working solution. One ml of working solution was also placed into 3 blank wells. The cells were then placed in the incubator 36°C, 5% CO_2_ for 3 hours. After 3 hours, 100 µL of the solution was sampled and placed into a 96 well plate, and fluorescence intensity was measured using a Tecan plate reader using 560_Ex_/590_Em_ nm filter settings, reading from the bottom of the plate. Immediately after measurement, the alamar blue working solution was washed three times with appropriate media for each cell type being measured and placed back in the incubator. The same cells were used for each measurement during days 0, 3, and 7.

### DNA Quantification Assay

Cells were cultured in gels as described above. However, gels were placed on SilGuard™ coated 24 well plates to prevent cell adhesion to the tissue culture plastic, ensuring the origin of the quantified DNA are exclusively taken from cells located within the gels. To coat plates in Sylgard, the Sylgard 184 elastomer was mixed with its curing agent in a 1:10 ratio. Subsequently, the mixture was poured into each well of a 24-well plate. The elastomer was allowed to cure at room temperature overnight for all experiments. Following the curing process, an alcohol-based reagent was applied through spraying, and the samples were subjected to UV sterilization for 24 hours.

A two-step procedure was used to quantify the double-stranded DNA (dsDNA): 1) sample homogenization, and 2) dsDNA quantification. Sample homogenization was performed via enzymatic digestion of the gel and cellular components using Papain (P4762-50mg, Sigma-Aldrich). Papain was reconstituted to a concentration of 10mg/mL using PBS, 300µL was aliquoted into Eppendorf tubes, and the papain stock solution was stored in −20°C until needed. L-cysteine (25-7210-00, Chemical Dynamics Co.) was prepared at a concentration of 24.2 mg/ mL in DI water, placed into 0.5 ml aliquots and stored at −20°C until use. EDTA was prepared into a stock of 0.333M in DI water and stored at 4°C in the fridge until needed (shelf life 3 months). Papain solution was prepared fresh just prior to hominization by combining and diluting papain, L-cysteine, and EDTA to a final concentration of 125 µg/ml, 0.242 mg/ml, and 0.333 M respectively with PBS. 300 µL of papain digestion solution was used per 28 µL gel. Each gel was then incubated at overnight at 6°C. The next day, homogenization was verified by pipetting samples up and down, noting absence of crosslinked gel material.

Double Stranded DNA was quantified using the Quantiflour® dsDNA System (E2670, Promega). 1X TE buffer was prepared by diluting the stock TE buffer 20 fold with microbial cell culture grade water (BP2820-100, Fisher BioReagents). The Quantifluor dsDNA dye was then diluted 400 fold with 1X TE buffer to create the working dye solution. 200 µL. A standard curve was prepared in a 96 well plate by placing 2µL of DNA standard into 200 µL of the working dye solution for a total amount of 200 ng of dsDNA standard. Then a 1:4 dilution series was performed down to 0.05 ng of dsDNA standard using the working dye solution as the diluent, and a blank containing just 2 µL of 1X TE buffer in the working dye solution. This was performed in triplicate. 15 µL of the homogenized samples were placed into an empty well of the plate, and 185 µL of the working dye solution was added ontop of the unknown samples. Samples were protected from light and incubated for 5-10 minutes, and the fluorescence (504nm_Ex_/531nm_Em_) was measured using a plate reader (SpectraMax® iD3, Molecular Devices). The three standard curves were averaged, and then used to calculate the total dsDNA present within each sample.

### Actin Staining

Cells in gels were cultured as described above, washed two times with 1X PBS and stained in 10% NBF for 15 min and washed three times with 1X PBS. Cells were then permeabilized with 100% Methanol at 4 °C for 30 min, washed two times with 1X PBS and permeabilized with 0.5% Trition-X-100 for 15 min at RT. Gels were washed two times with 1X PBS and blocked using blocking solution (1% BSA, 0.3 M glycine, and 0.01% Triton-X-100) for one hour. Cells were then incubated with anti-pan actin mouse monoclonal antibody (AANO2, Cytoskeleton inc.) (1:500)in blocking solution and agitated overnight at 4 °C. Gels were washed with 0.1% Triton-X in 1X PBS, and then three times with 1X PBS. Cells were then incubated with Goat anti-mouse IgG2b-AF555 (1091-32, SouthernBiotech) (1:400) in blocking solution for 60 min at RT in the dark. Gels were washed three times with 1X PBS and then incubated in DAPI (Anaspec AS-83210) (1:10,000) in 1X PBS for 20min at RT. Cells were mounted on coverslips just prior to imaging. Cells were imaged using a confocal microscopy (Nikon Eclipse Ti). Cell area was quantified by first importing a z-stack into the Fiji distribution of ImageJ2. The Z-stack was then projected into two dimensional space using the maximum projection function. After splitting the color channels, the threshold was adjusted such that the only the DAPI or the phalloidin staining was applied. Last, the “analyze particles” function was applied to gain two pieces of information: 1) the number of particles taken from the DAPI channel, and the total particle area taken from the phalloidin area. Should two nuclei be very close and be counted as one particle, the particle count was adjusted manually for accuracy. The total particle area was then normalized to the number of particles to get an average area per cell. Each group tested was done using three replicates, each done in triplicate.

### Statistical Analysis

Statistical analysis was done using multi-way analysis of variance (ANOVA) with a Tukey post-hoc test. All comparisons in this manuscript are statistically significant (p <0.05) unless otherwise noted. Statistical data can be found in **Fig. S24**.

## Results and Discussion

### Design and synthesis of peptide libraries to quantify non-specific peptide degradation

There are dozens of human exopeptidases^27,28^ and the activity of each protease is typically dependent on the amino acid at the termini of the peptide.^29^ Significant effort has gone into preventing degradation of therapeutic proteins, and it modification of the N-terminal amine or C-terminal carboxylic acid or the inclusion of β-amino acids frequently reduces non-specific peptide degradation.^24,30,31^ While most work assessing proteolytic degradation of peptides has focused on the substrate preferences for individual proteases,^32^ or degradation of specific peptides,^33^ we set out to better understand peptide degradation due to the total protease expression of entire cell types across a range of terminal chemistries. In order to characterize the effects of non-specific proteolytic degradation as a function of end group, we designed a series of RGEFV peptide libraries based upon the widely-used RGD peptide. Seven peptide libraries were synthesized, each having a different terminal chemistries and containing 19 of the 20 canonical amino acids (excluding cysteine) to quantify peptide degradation by cell type (**Fig. 1**). The aspartic acid (D) on the RGD was mutated to glutamic acid (E) to prevent the peptides from binding cell-surface integrins while retaining their physiochemical properties,^34^ and also included a hydrophobic Phe-Val utilized in the RGDFV adhesion peptide^35^ to improve chromatographic retention during analysis.

We utilized a split-and-pool peptide synthesis technique to build the libraries.^36^ This was done by splitting the resin equally into 19 different vials at the desired synthesis step, followed by the addition of a different amino acid into each of the 19 vials (**Fig. 1B**). Upon completion of the coupling steps the 19 resins were then recombined into a single batch for further synthesis. For the N-terminal libraries the split-pool synthesis was performed on the N-terminal side of the RGEFV peptide. The N-terminal library resin was split again make four different 19-peptide libraries with different N-terminal modifications. These N-terminal chemistries are an amine (**NH**_**2**_), an acetyl (**Ac**), an N-terminal β-alanine (**N-βA**), and an N-terminal acetylated β-alanine (**Ac-βA**) (**Fig. 1B**). For the C-terminal libraries the split pool was performed on the C-terminal side of the RGEFV peptide with one of three C-terminal chemistries: a C-terminal carboxylic acid (**COOH**), a C-terminal amide (**Am**), or a C-terminal amidated β-alanine (**C-βA**) (**Fig. 1C**). Peptide libraries were added to cell culture media and analyzed by liquid chromatography mass spectrometry (LCMS) (**Fig. 1D**). All N-terminal libraries had an amidated C-terminal β-alanine, and all C-terminal libraries had an acetylated β-alanine on the N-terminus.

### Quantification of peptide degradation by cells

Each of the seven peptide libraries was added to the cell culture media of three cell types commonly used in biomedical research: human mesenchymal stem/stromal cells (hMSCs), human umbilical vein endothelial cells (hUVECs), and classically polarized macrophages (Macrophages) cultured on tissue culture plastic. Samples of each peptide library were taken in cell culture media at 0, 1, 4, 8, 24h, and 48h, acidified with acetic acid to prevent further enzymatic degradation, and quantified using LCMS. The extent of degradation was determined by calculating the ratio of peptide found at later time points to the zero hour time point. While peptides with 19 different terminal amino acids were present in every library, isoleucine and leucine have identical masses and were combined for analysis. It should be noted that any chemical modification to the peptide that changes the mass of the peptide will reduce the intensity of the peptide peak in mass spectrometry (**Fig. 2**). While our data suggest that a significant fraction of degradation is due to exopeptidase activity, it is possible and even likely that endopeptidases or other enzymes are also degrading these peptides.

**Fig. 2.**
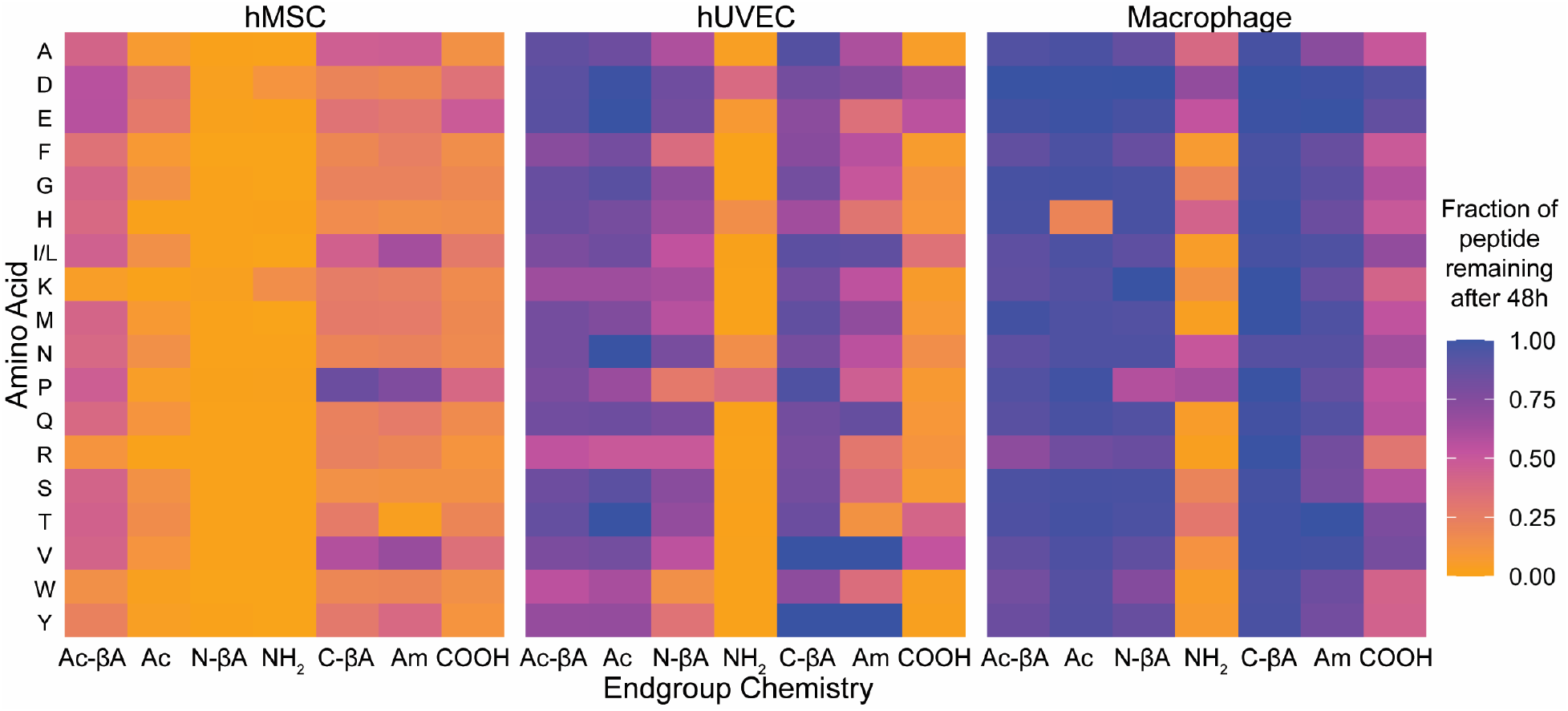
Non-specific degradation of peptides depends upon cell type and the chemistry of peptide termini. Peptide libraries were incubated with three different cell types for 48 hours and the fraction of peptides remaining was quantified. Notably, peptides with N-terminal amines (NH_2_) were almost completely degraded irrespective of the N-terminal amino acid, while adding a β-amino acid to the C-terminus of the peptide (C-βA), or an acetylated β-amino acid to the N-terminus (Ac-βA) significantly reduced degradation.

Our results show that non-specific degradation is primarily controlled by the chemistry of the peptide termini, and that simple modifications can drastically reduce peptide degradation (**Fig. 3 and S1**). The amino acids present at the termini influenced how quickly a peptide was degraded, but the difference between amino acids was much smaller than terminal chemistries. Peptides with N-terminal amines were rapidly degraded for almost every amino acid by all three cell types. In hMSCs after eight hours in culture 15 of the 18 peptides with N-terminal amines (**NH**_**2**_) had less than 50% of the peptide remaining, 13 peptides had less than 25% remaining, and 9 peptides had less than 5% of the original peptide remaining. hUVECs demonstrated less degradation of NH_2_ peptides, but 13 peptides had less than 50% remaining after 8 hours, and 9 peptides had less than 25% remaining. Macrophages had the least degradation, but nine of the 18 peptides had less than 50% remaining after 8 hours in culture. For all cell types, modifications of the N-terminus reduced peptide degradation, although they different levels of efficacy in preventing degradation. While peptides with N-terminal β-alanines (**N-βA**) were completely degraded by 48 hours when cultured with hMSCs, 42% of N-βA peptides remained after eight hours versus 17% for peptides with N-terminal amines, suggesting that the N-βA does slow down degradation in hMSCs. The N-βA modification was more effective at inhibiting peptide degradation in hUVECs, which increased the fraction of peptides remaining after 48 hours from 7% in NH_2_ to 58% in N-βA, and macrophages, which went from 25% remaining for NH_2_ to over 90% for N-βA.

**Fig. 3.**
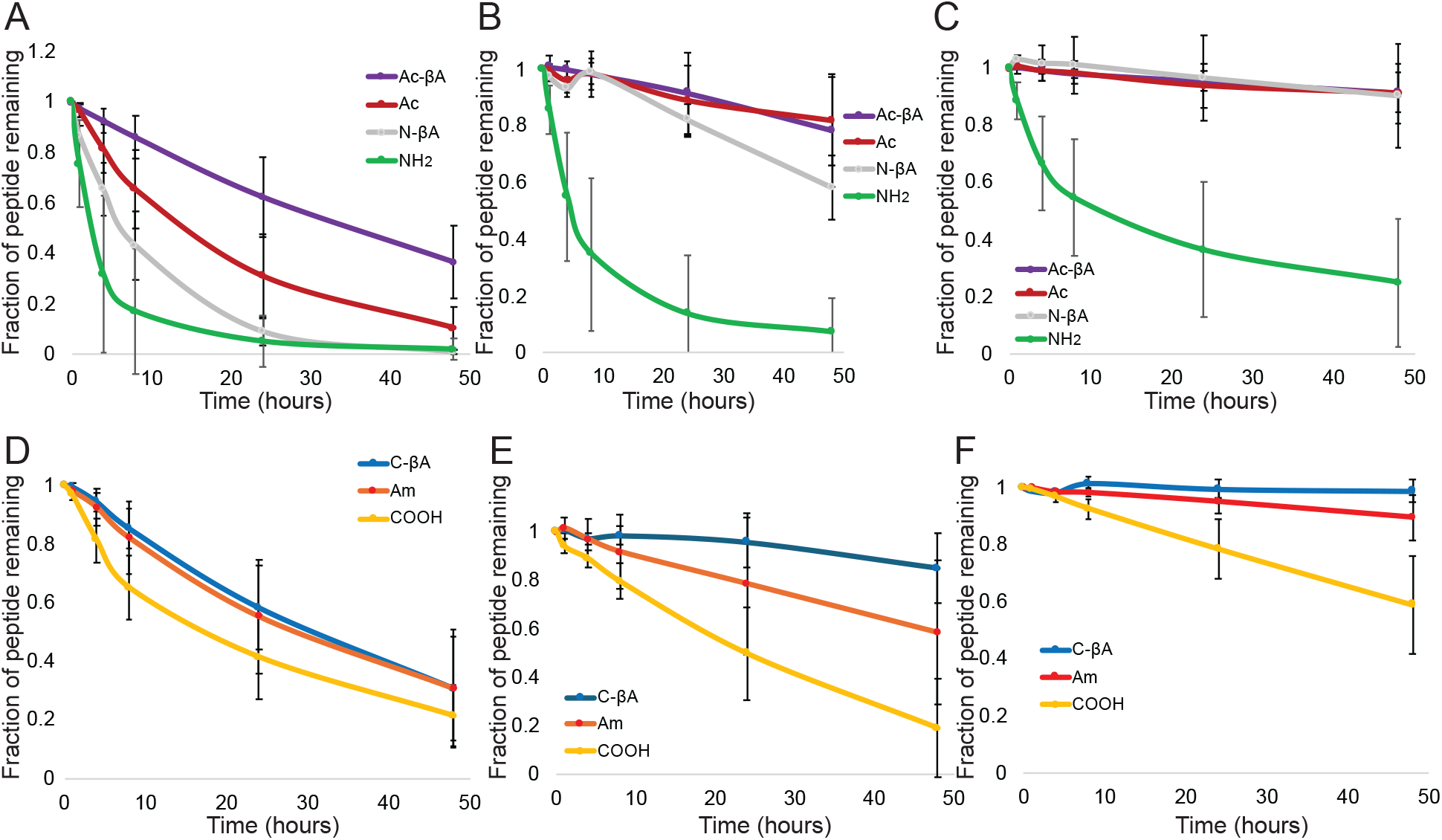
Non-specific degradation of peptides with N-terminal amines across three different cell types. The amount of degradation at different time points was quantified. Averaging over all amino acids, quantified the effects of N-terminal modifications for (A) hMSCs, (B) hUVECs, and (C) macrophages, and C-terminal modifications for (D) hMSCs, (E) hUVECs, and (F) macrophages. Error bars represent the standard deviation across all terminal amino acids.

The positive charge of the N-terminal amine can play a role in aminopeptidase recognition of the peptide substrate.^37^ Both the NH_2_ and N-βA termini are positively charged, and the uncharged acetylation (**Ac**) and acetylated β-alanines (**Ac-βA**) were both more effective at preventing degradation. While there was essentially complete degradation of all peptides with N-terminal amines or N-terminal β-alanines when cultured with hMSCs on tissue culture plastic, 10% of acetylated peptides remained, and 37% of Ac-βA peptides. For hUVECs, 82% of acetylated peptides remained, and 78% of peptides with acetylated β-alanines, and for macrophages approximately 91% of peptides remained for both Ac and Ac-βA.

The chemistry of the C-terminus also had a significant effect on peptide degradation, although to a lesser extent than the chemistry of the N-terminus. Peptides with C-terminal carboxylic acids typically had the most degradation, with hUVECs having the most degradation, with 19% remaining, followed by hMSCs (21%) and macrophages (59%). Modification of the C-terminus reduced degradation for all cell types. C-terminal amidation and an amidated C-terminal β-alanine both had 31% of peptide remaining after 48 hours for hMSCs. However, the C-β A modification was superior to C-terminal amidation in preventing peptide degradation hUVECs (85% remaining in C-βA versus 59% in Am) and macrophages (99% remaining in C-βA versus 90% for Am) than peptides which had C-terminal amides, and those with C-terminal β-alanines had the least. Current strategies to prevent exopeptidase degradation of peptides are largely focused on non-natural amino acids or acetylation of the N-terminus. Peptides with N-terminal acetylated β-alanines and C-terminal amidated β-alanines would largely have reduced degradation over existing strategies for all cell types. Acetylated N-terminal β-alanines result in reduced degradation compared to standard acetylation for hMSCs, and amidated C-terminal β-alanines have significantly reduced degradation for hUVECs and macrophages. Globally peptide degradation varied by cell type, and across all peptides hMSCs had more degradation than hUVECs, which had more degradation the macrophages for every end group at 24 and 48 hours, with the exception of the C-terminal carboxylic acid, which was more degraded at 48 hours by hUVECs than hMSCs.

We tested the peptide libraries with three different donors for each cell type, including both sexes and multiple ethnicities, to ensure that the results were robust across biological variation. We found that the trends for degradation across different chemistries and amino acids was similar across donors for all three cell types, and typically there was less than a 10% difference in degradation rate across all amino acids between donors. Averaging across all end groups and amino acids, after 48 hours there was between 54% and 59% of peptide remaining across the three hUVEC donors, 13% and 22% across the three hMSC donors, and 76% to 84% for the three macrophage donors (**Fig. S2-4**). Additionally, the THP-1 monocyte cell line had 70% of peptides remaining across all conditions, and the pattern of degradation across end groups and amino acids closely matched the human primary cells.

The split-and-pool libraries contain 19 different peptides each at a 37 µM concentration. Most biomaterials are functionalized by a single peptide at a higher concentration. We performed degradation studies using an individual peptide with a glycine at the terminus at concentrations ranging from 19.5 µM to 5,000 µM. We broadly saw that peptides at lower concentrations were more rapidly degraded than peptides at higher concentrations (**Fig. 4 and Fig. S5**). The effects of terminal chemistry on peptide degradation was broadly conserved across concentration ranges, with peptides having N-terminal amines being rapidly degraded even at 5 millimolar concentrations. After 48 hours only 5% of amine-terminated RGEFV peptides remained after starting with an initial concentration was 5,000 µM. C-terminal degradation could also be rapid, and only 26% of the initial peptide remained of an initial 5,000 µM COOH peptide when cultured with hUVECs (**Fig. S5**). Overall these results indicate that increasing peptide concentration leads to higher amounts of peptide at later time points, but that modifying the chemistry at the terminus is the most effective method for increasing the concentration of peptide after 48 hours.

**Fig. 4.**
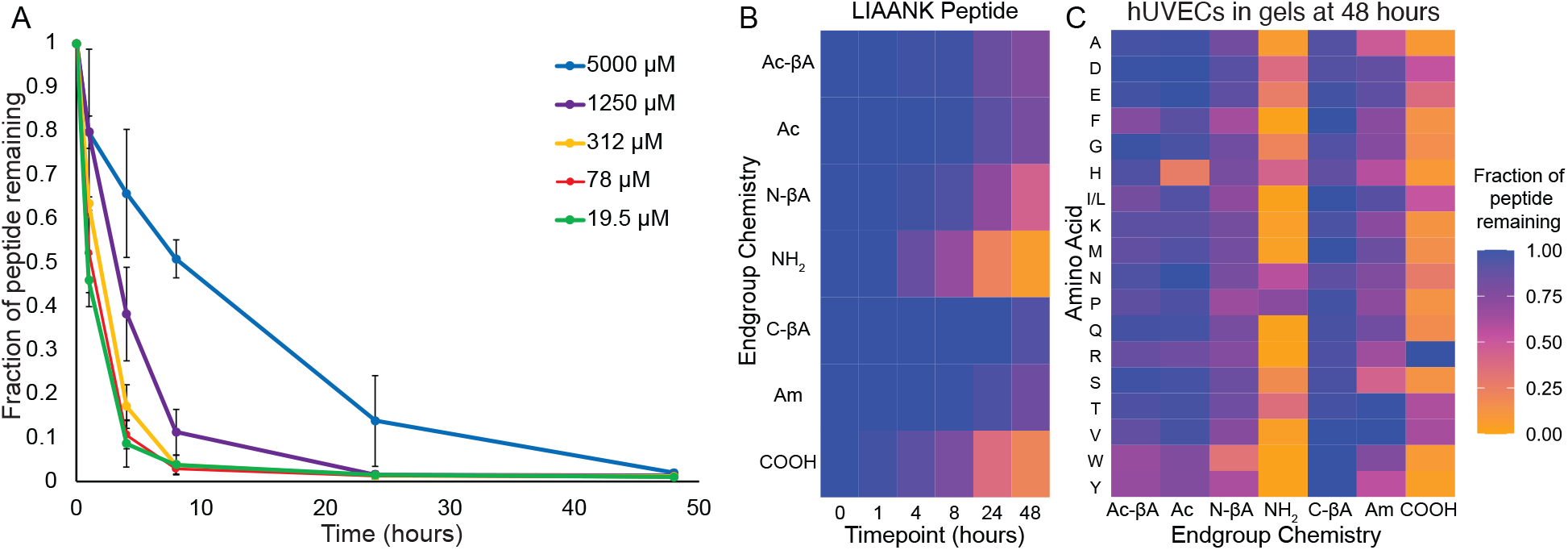
Non-specific degradation of peptides with N-terminal amines is rapid across peptide sequences and for cells in hydrogels. (**A**) Degradation of peptides with N-terminal amines can be rapid even at 5 millimolar peptide concentrations. (**B**). A scrambled version of the TGF-β1 mimicking LIANAK peptide was rapidly degraded by hUVECs when the peptide feature an N-terminal amine or C-terminal carboxylic acid. (**C**) hUVECs cultured in PEG hydrogels rapidly degrade RGEFV peptides with free N-terminal amines or C-terminal carboxylic acids.

Our initial libraries were derivatives of an RGEFV peptide based upon the RGD cell adhesion peptide. To ensure that the results are not specific to this sequence we synthesized two other peptides, an IVKVA peptide based upon the IKVAV laminin-mimetic peptide,^38^ and an LIAANK based upon the LIANAK TGF-β mimetic peptide. We made peptides with each of the terminal chemistries having glycine at the terminus, since it both does not have any special chemical groups and had degradation in the RGEFV peptides near the average (**Figs. 4B and S6**). We found that the same broad trends held, with most peptides having more C-terminal degradation in the COOH condition, and acetylating the N-terminus, either Ac or Ac-βA reduced N-terminal degradation.

### Peptide degradation by cells in hydrogels

The overall goal of this work is to understand how cells in hydrogel scaffolds degrade peptides. Since cell spreading and migration often involve proteases *in vivo*, we cultured hMSCs, hUVECs, and macrophages within protease-substrate peptide crosslinked 8-arm poly(ethylene glycol) (PEG) hydrogels. We found that hUVECs peptide degradation with hUVECs was similar between tissue culture plastic (**Fig. 2**) and within hydrogels (**Fig. 4B**). Macrophages also had similar peptide degradation kinetics when soluble peptides were added to cells on tissue culture plastic or cells in gels. Interestingly, hMSCs had significantly less non-specific degradation of peptides when cultured in gels (**Fig. S6**) compared to cells cultured on tissue culture plastic (**Fig. 2**). While acetylation of the N-terminus was largely effective at reducing non-specific degradation, peptides with an N-terminal histidine were an exception, and peptides with an acetylated histidine were cleaved for all three cell types when cultured in gels (**Fig. 4, Fig. S7**). Overall the same broad peptide degradations trends held for cells in gels, with N-terminal amines undergoing significant degradation for all three cell types versus other N-terminal functionalizations, and C-terminal carboxylic acids being significantly degraded by hMSCs and hUVECs compared to other terminal chemistries (**Table 1**, **Fig**. **S 7**). Macrophages cultured on tissue culture plastic had significantly reduced degradation of C-terminal libraries compared to the two other cell types (**Fig. 2-3**), but were still only had 59% of peptides remaining after 24 hours across amino acids. However, macrophages encapsulate in hydrogels had greater than 96% remaining of all C-terminal peptide chemistries after 48 hours in culture (**Table 1**).

**Table 1.** Quantifying the degradation of peptide libraries after 48 hours in culture with cells encapsulate in PEG hydrogels. Each value is the average across 19 amino acids.

| Terminal Chemistry | Cell Type | Fraction Remaining | Standard Deviation |
| --- | --- | --- | --- |
| Ac-βA | hMSC | 0.81 | 0.12 |
|  | hUVEC | 0.90 | 0.10 |
|  | Macrophage | 0.99 | 0.03 |
| Ac | hMSC | 0.82 | 0.15 |
|  | hUVEC | 0.90 | 0.18 |
|  | Macrophage | 0.87 | 0.20 |
| N-βA | hMSC | 0.62 | 0.17 |
|  | hUVEC | 0.76 | 0.14 |
|  | Macrophage | 0.95 | 0.06 |
| NH<sub>2</sub> | hMSC | 0.20 | 0.23 |
|  | hUVEC | 0.19 | 0.22 |
|  | Macrophage | 0.13 | 0.25 |
| C-βA | hMSC | 0.83 | 0.07 |
|  | hUVEC | 0.99 | 0.08 |
|  | Macrophage | 0.98 | 0.03 |
| Am | hMSC | 0.73 | 0.11 |
|  | hUVEC | 0.75 | 0.17 |
|  | Macrophage | 0.98 | 0.03 |
| COOH | hMSC | 0.36 | 0.15 |
|  | hUVEC | 0.25 | 0.20 |
|  | Macrophage | 0.96 | 0.04 |

### Effects of peptide conjugation on degradation

In addition to their roles as adhesion ligands or growth factor mimetic peptides, peptides are also widely used to crosslink hydrogel matrices.^17,39^ This is often done by utilizing chemistries that can react with canonical amino acids, such as thiol-maleimide reactions,^40^ or incorporating non-natural chemistries into the peptide side chain, such as azides, which can then undergo click reactions with reactive groups present on polymers.^41^ To better understand how these peptide modifications influence degradation kinetics, we synthesized a series peptides and put an azide-lysine on either the N-terminus or C-terminus and then functionalized the termini with each of the seven different terminal chemistries (**Fig. 5**). A portion of these peptides were then functionalized with a (PEG)_12_ chain modified with a dibenzocyclooctyne (DBCO) group that is commonly used in biomaterial synthesis. We added both the azide-containing peptides and PEG-modified peptides to cell culture media and quantified degradation by hMSCs, hUVECs, and macrophages (**Fig. 5** and **S8**). We found that PEG modification slow down peptide degradation for every N-terminal and C-terminal chemistry across hMSCs, hUVECs, and macrophages. These results indicate that non-specific peptide degradation depends upon both the end group chemistry of the peptide and also the presence of bulky groups which can prevent the degradation of peptides conjugated to matrices, such as crosslinking peptides.

**Fig. 5.**
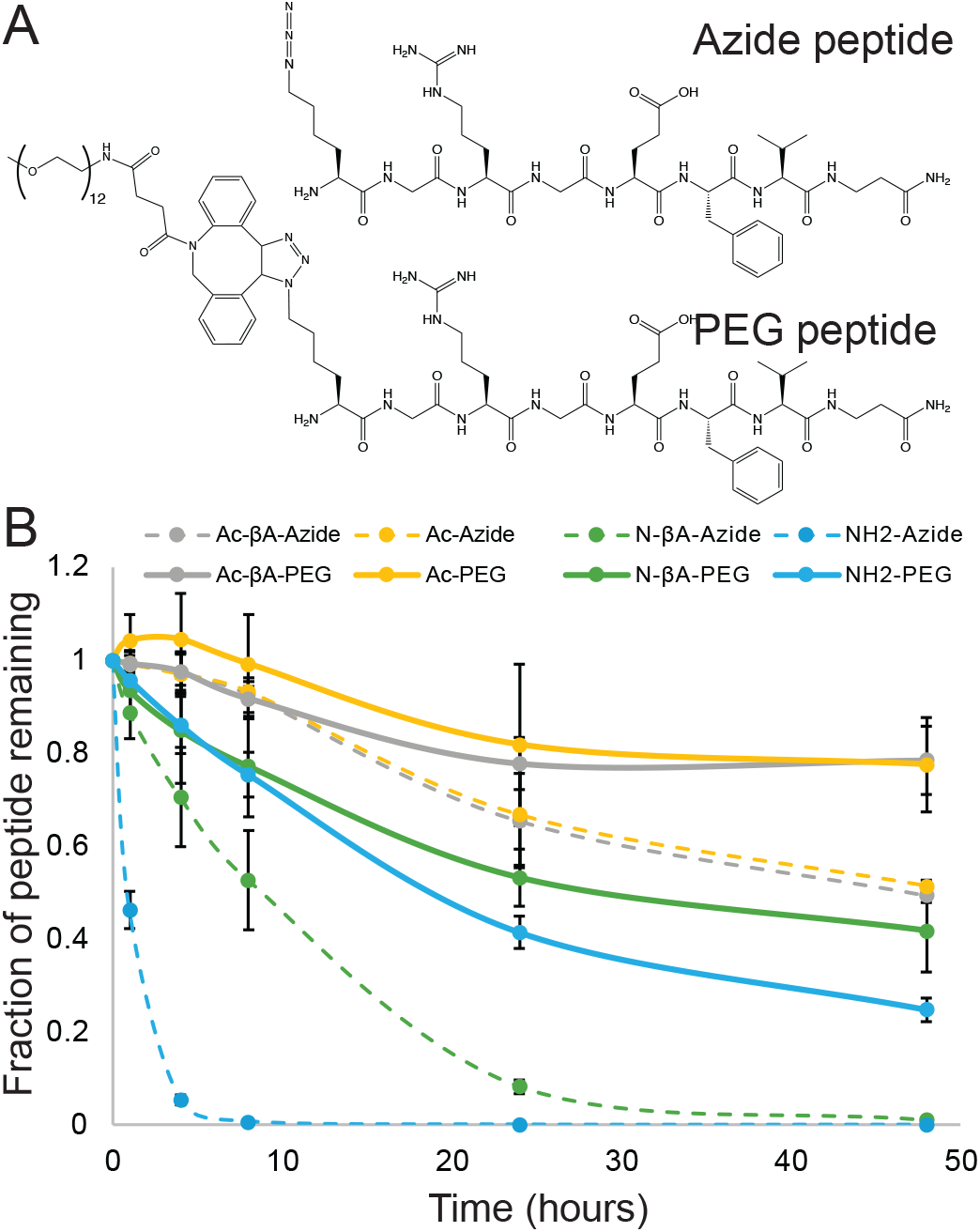
PEG conjugation to peptides reduces degradation rates. (**A**) Peptides with different terminal chemistries were synthesized with azides on either their N- or C-terminus a portion were modified with (PEG)_12_-DBCO. (**B**). Both the azide and PEG modified peptides were incubated with hMSCs for 48 hours, and peptides featuring PEG conjugations showed less degradation across all N-terminal chemistries.

**Fig. 6.**
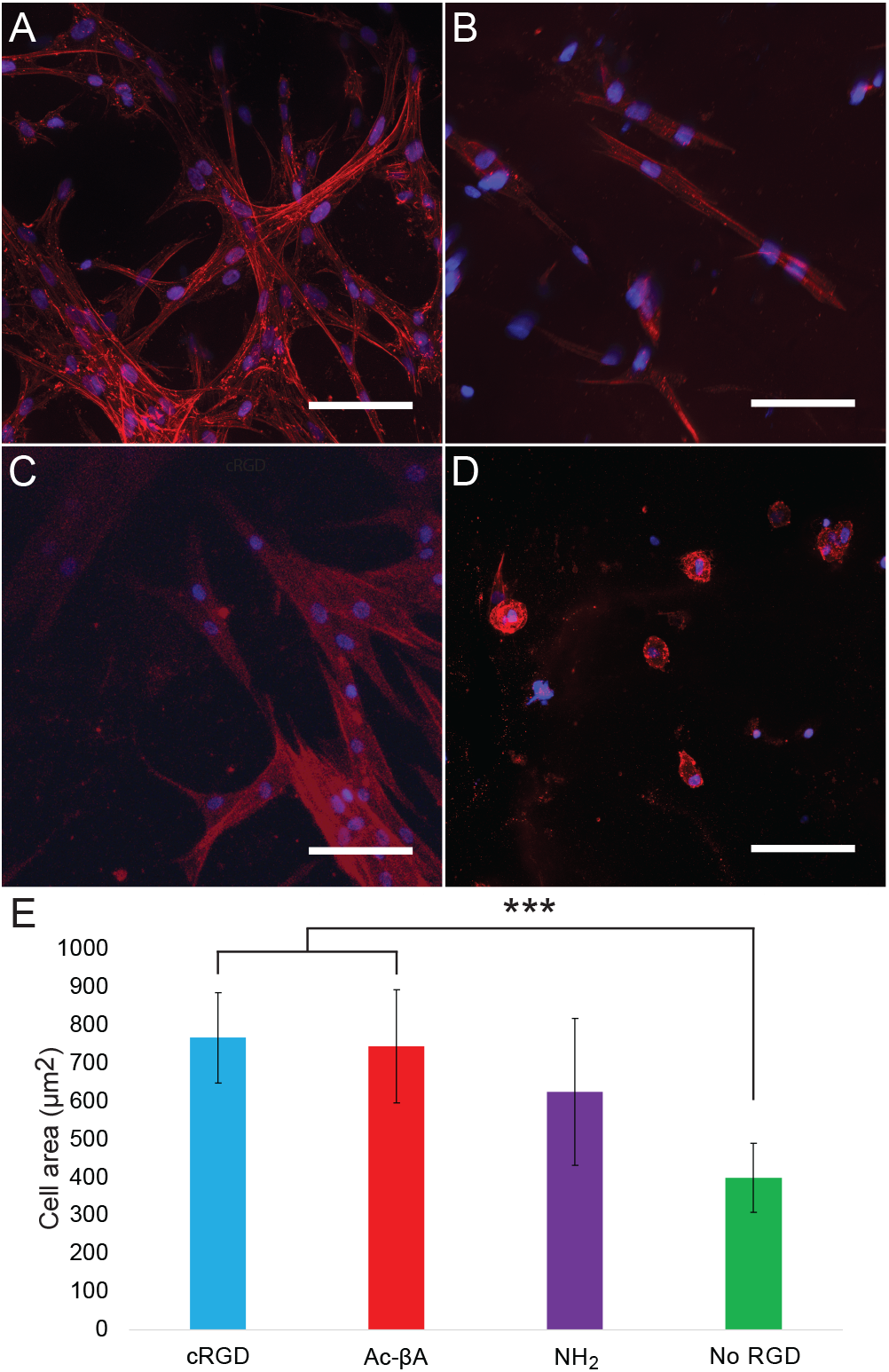
RGD is required for cell spreading and viability within hydrogels. (**A**) Ac-βA-GRGDS, (**B**) NH_2_-GRGDS, (**C**) cyclic RGDS (cRGD), and (**D**) no added RGDS. (**E**) hMSC spreading was increased in hydrogels which were functionalized with RGD peptides. Red is actin and blue is the nuclei. Scale bar is 100 µm and * indicates p < 0.05, *** indicates p < 0.001 by Tukey’s post hoc test.

### Effects of RGD chemistry on cell behavior in hydrogels

We next sought to better understand how the presence of fast and slow degrading RGD sequences influenced cell behavior and and peptide degradation when covalently coupled to a hydrogel. To do this we made hydrogels with both “Fast” degrading NH_2_-GRGDS peptides and slow degrading Ac-βA-GRGDS peptides. As a positive control we used a cyclic RGD peptide (cRGD), which does not have a terminus and is not susceptible to degradation by exopeptidases. We also included a negative control which consisted of gels without any RGD peptides. To better understand the effects of RGD stability on cell morphology, we quantified hMSC spreading at Day 7 in hydrogels with cyclic RGD, slow-degrading Ac-βA-GRGDS, fast degrading NH_2_-GRGDS and gels with no RGD peptides (**Fig. 5E** and **S9**). We found that hMSCs in the cRGD gels had the most spreading (769 ± 119 µm^2^), followed by the Ac-βA-GRGDS (746 ± 149 µm^2^), NH_2_-GRGDS (626 ± 193 µm^2^), and gels without RGD (400 ± 91 µm^2^).

Hydrogels with cRGD and Ac-βA-GRGDS had more spreading than gels with either NH_2_-GRGDS or no RGD, although the differences with the NH_2_-GRGDS were not statistically significant (p > 0.05). We also found that viability within gels was primarily dependent upon the presence of RGD, however this varied by cell type (**Fig. S10**). Gels containing the cyclic RGD peptide had the greatest viability across all cell types, which could be due to both the inability of cyclic peptides to undergo exopeptidase degradation, or the beneficial effect that the cyclization of RGD sequences can have on adhesion.

## Conclusion

In conclusion, we synthesized a series of peptide libraries to better understand how the proteases secreted by cells degrade peptides in culture. Soluble peptides with canonical termini, such as an N-terminal amine, were rapidly degraded by two cell types found in most tissues, endothelial cells and macrophages, and one commonly used in biomedical applications, hMSCs, irrespective of the N-terminal amino acid. Peptide degradation with N-terminal amines and C-terminal carboxylic acids was also found for cells cultured in hydrogels, and multiple peptide sequences. We found that simple modifications to the protein termini could greatly slow down or abolish the non-specific degradation of soluble peptides by cells on tissue culture plastic or cells within a hydrogel. Finally, we found that the RGD was important for cell spreading and viability within hydrogel matrices.

## Acknowledgements

We would like to acknowledge our funding sources, the NIH (1R21GM143593-01) and NSF (Award 2138723). We are grateful to the lab of Lesley Chow for the use of their preparative high performance liquid chromatography.

