## Supplemental Information for "Quantifying and controlling the proteolytic degradation of cell adhesion peptides"

Supporting Information for the manuscript:
Quantifying and controlling the proteolytic degradation of cell adhesion peptides

Samuel J. Rozans, Abolfazl Salehi Moghaddam, Yingjie Wu, Kayleigh Atanasoff, Liliana Nino, Katelyn Dunne, E. Thomas Pashuck

**Table of Contents:**

**Figure S1** - Degradation of soluble peptides cultured with cells on TCP. Pages 1-5.
**Figure S2** - Comparison of peptide degradation by different hMSC donors. Pages 6-9.
**Figure S3** - Comparison of peptide degradation by different hUVEC donors. Pages 10-13.
**Figure S4**. Comparison of peptide degradation by different PBMC donors. Pages 14-17.
**Figure S5**. Degradation of peptides at different concentrations. Pages 18-20.
**Figure S6**. Effect of different peptide sequences on non-specific degradation. Pages 21-23.
**Figure S7**. Degradation of soluble peptide libraries by cells in PEG hydrogels.Pages 24-29.
**Figure S8**. PEG conjugation to peptides slows degradation. Page 30.
**Figure S9**. Microscopy of cells growing in gels with different RGD presentations. Page 31.
**Figure S10**. Quantification of viability and proliferation in hydrogels containing different RGD sequences. Page 32.
**Figure S11**. Standard curves validating the use of LCMS to measure the concentration of peptides. Page 33.
**Figure S12**. LCMS spectra of the Ac-X-RGEFV-βA-$NH_2$ libraries. Pages 34-42.
**Figure S13**. LCMS spectra of the Ac-βA-X-RGEFV-βA-$NH_2$ libraries. Pages 43-51.
**Figure S14**. LCMS spectra of the Ac-βA-RGEFV-X-$NH_2$ libraries. Pages 52-60.
**Figure S15**. LCMS spectra of the Ac-βA-RGEFV-X-βA-$NH_2$ libraries. Pages 61-69.
**Figure S16**. LCMS spectra of the Ac-βA-RGEFV-X-COOH libraries. Pages 70-78.
**Figure S17**. LCMS spectra of the $NH_2$-βA-X-RGEFV-βA-$NH_2$ libraries. Pages 79-87.
**Figure S18**. LCMS spectra of the $NH_2$-X-RGEFV-βA-$NH_2$ libraries. Pages 88-96.
**Figure S19**. LCMS spectra of the peptides for the concentration studies. Pages 97-100.
**Figure S20**. LCMS spectra of the LIAANK peptides. Pages 100-103.
**Figure S21**. LCMS spectra of the IVKVA peptides. Pages 104-107.
**Figure S22**. LCMS spectra of the Azide and PEG modified peptides. Pages 107-114.
**Figure S23**. LCMS spectra of the peptides used for cell culture. Pages 115-117.
**Figure S24**. Statistical analyses. Pages 118-135.

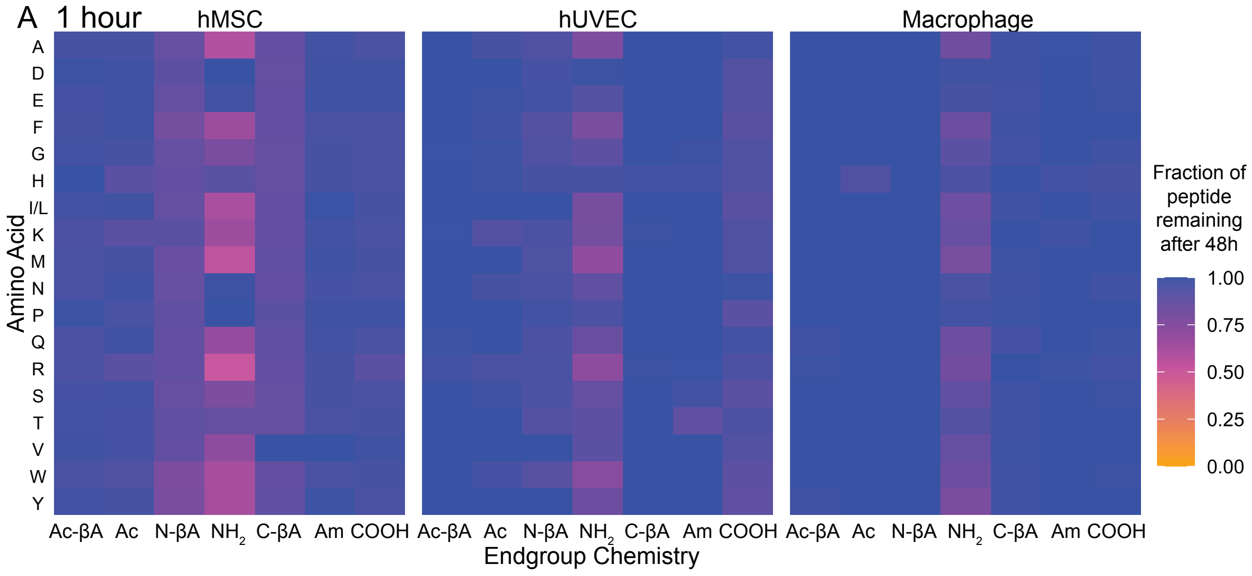

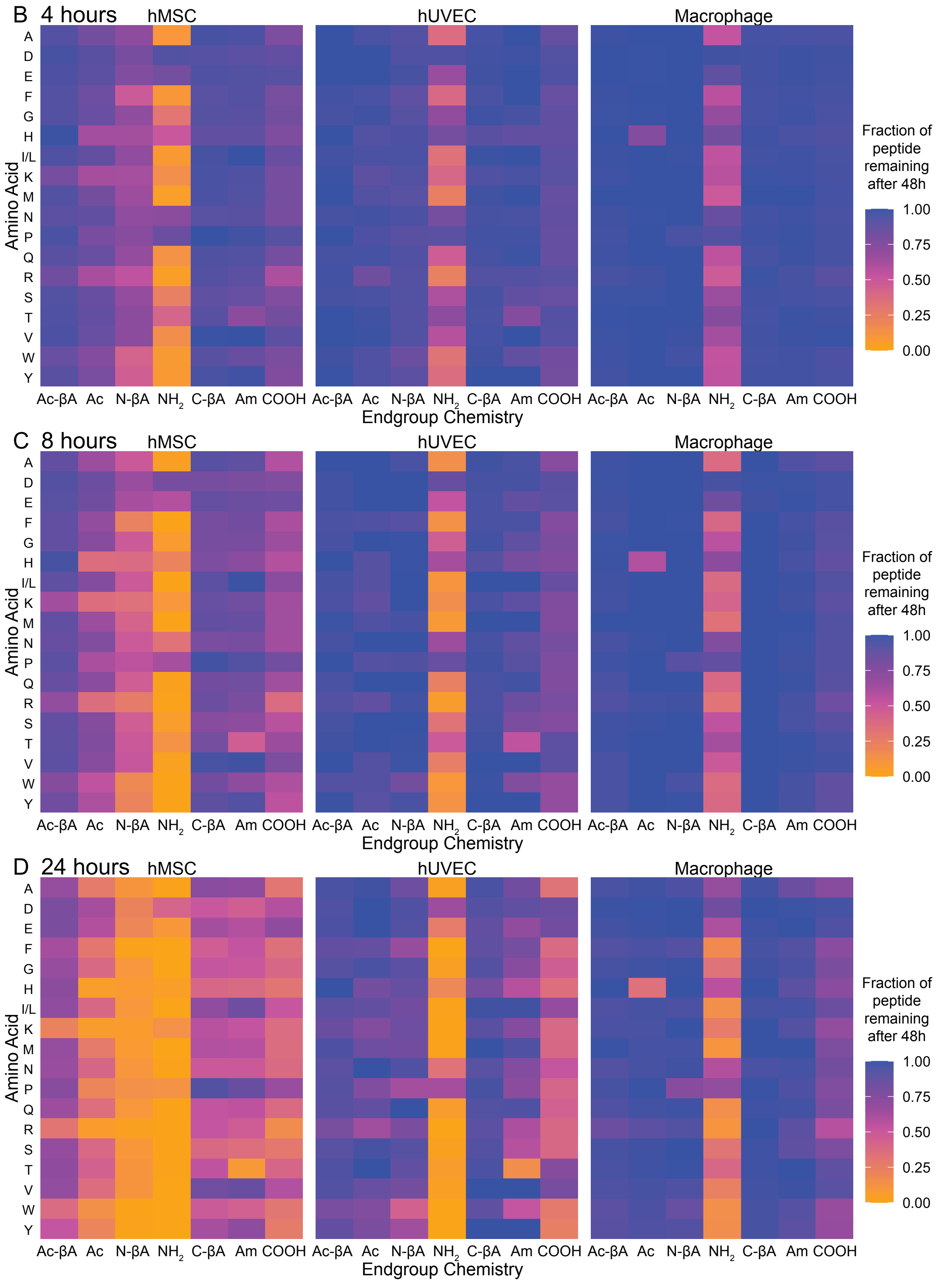
B 4 hours
hMSC
hUVEC
Macrophage
C 8 hours
hMSC
hUVEC
Macrophage
D 24 hours
hMSC
hUVEC
Macrophage
Amino Acid
A
D
E
F
G
H
I/L
K
M
N
P
Q
R
S
T
V
W
Y
Ac-βA Ac N-βA NH2 C-βA Am COOH
Endgroup Chemistry
Fraction of peptide remaining after 48h
1.00
0.75
0.50
0.25
0.00

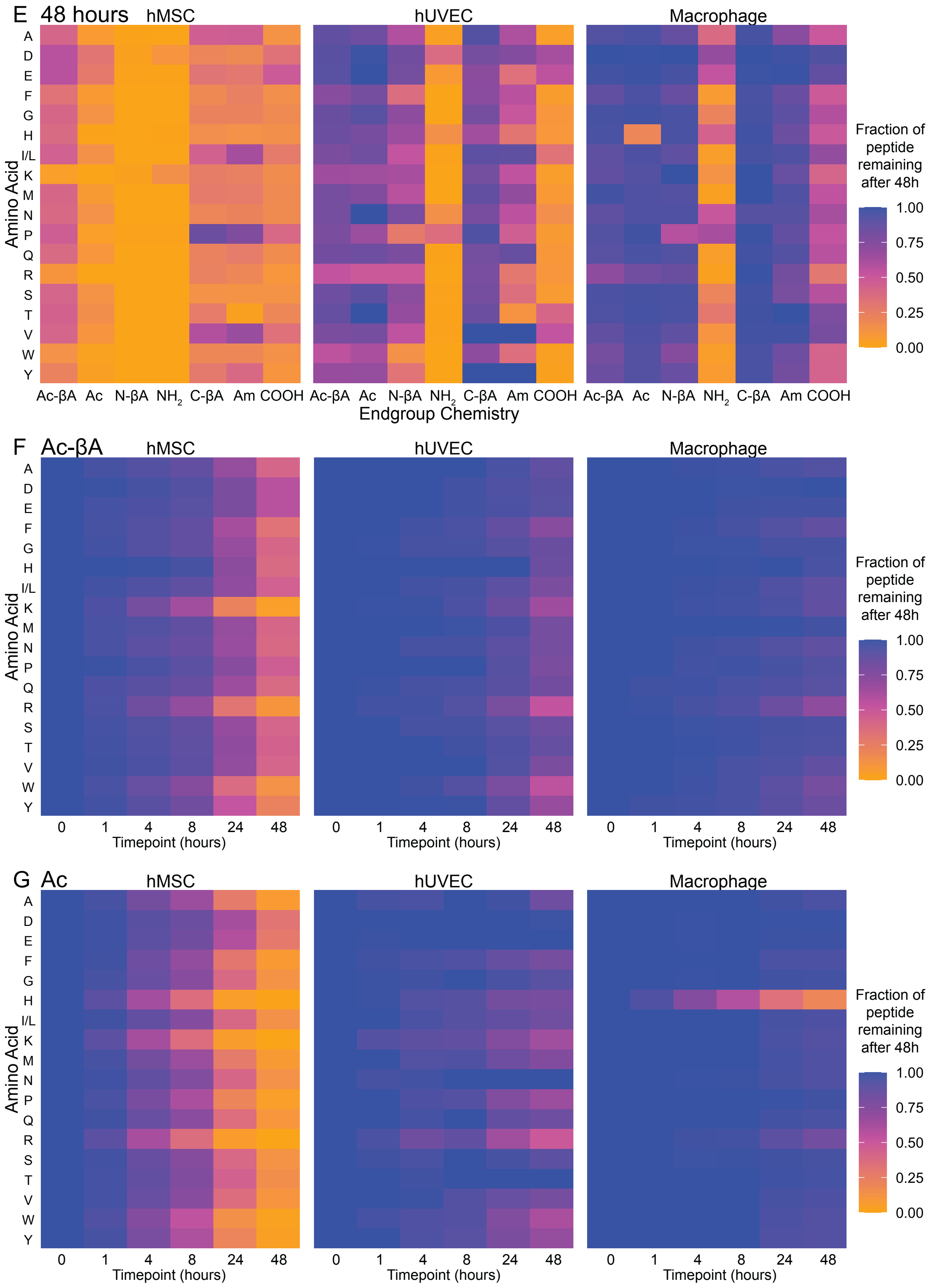
E 48 hours
hMSC
hUVEC
Macrophage
Amino Acid
A
D
E
F
G
H
I/L
K
M
N
P
Q
R
S
T
V
W
Y
Ac-βA Ac N-βA $NH_2$ C-βA Am COOH
Endgroup Chemistry
Fraction of peptide remaining after 48h
1.00
0.75
0.50
0.25
0.00
F Ac-βA
G Ac
0 1 4 8 24 48
Timepoint (hours)

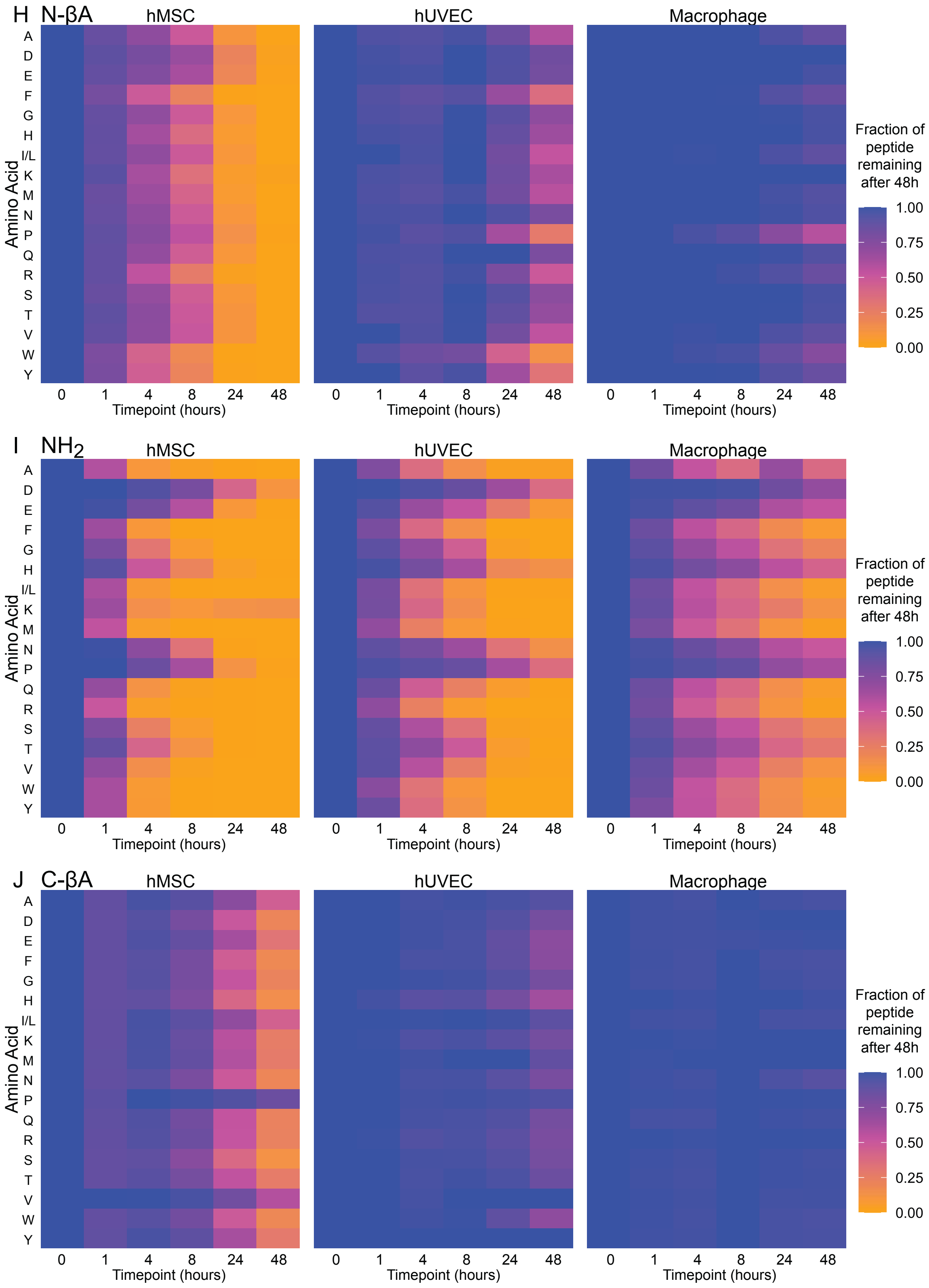

H N-βA
I NH2
J C-βA
hMSC
hUVEC
Macrophage
Amino Acid
A
D
E
F
G
H
I/L
K
M
N
P
Q
R
S
T
V
W
Y
0
1
4
8
24
48
Timepoint (hours)
Fraction of peptide remaining after 48h
1.00
0.75
0.50
0.25
0.00

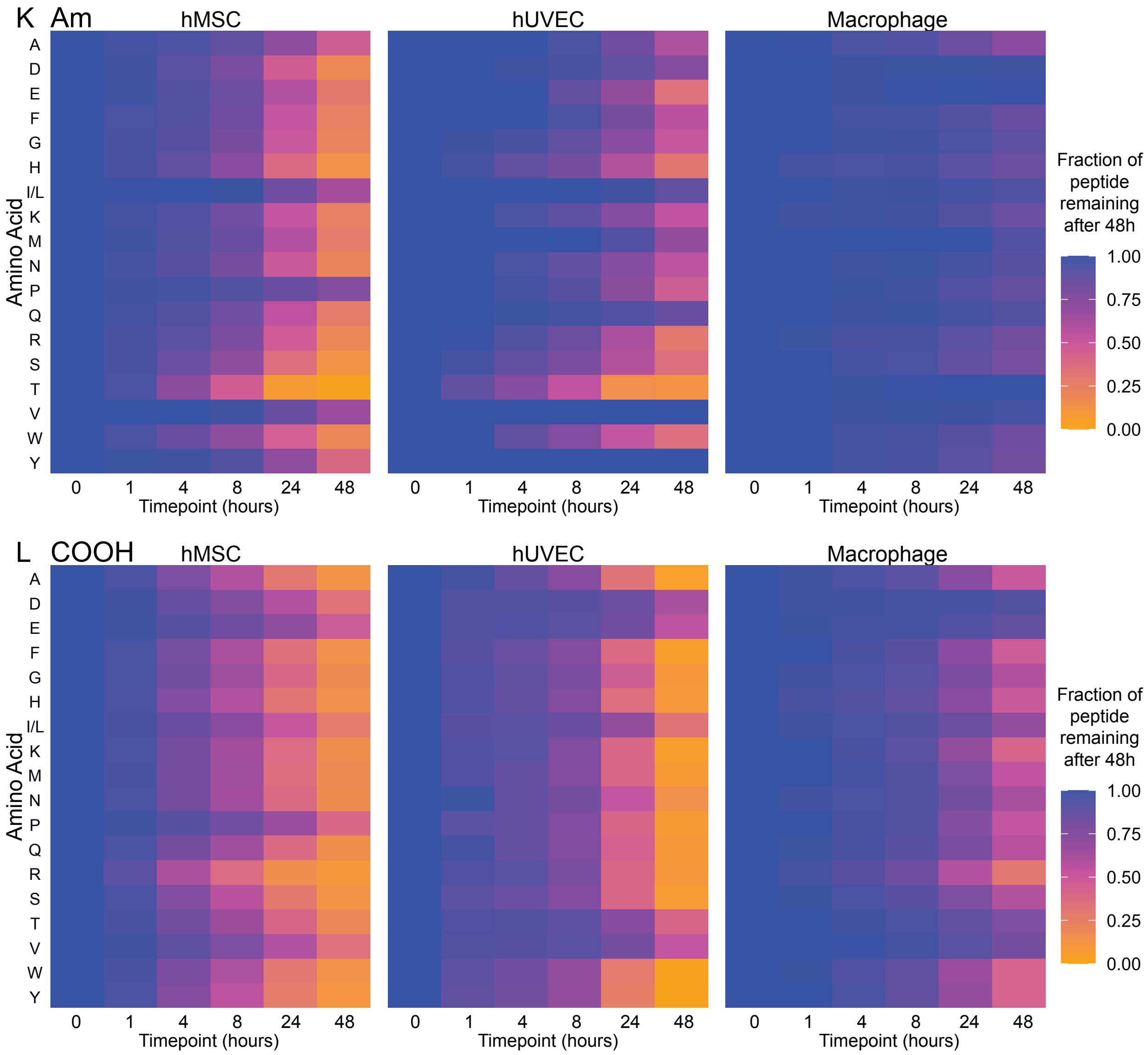


**Figure S1**. Degradation of soluble peptides cultured with cells on tissue culture plastic. Degradation was quantified at (**A**) 1 hour, (**B**) 4 hours, (**C**) 8 hours, (**D**) 24 hours and (**E**) 48 hours. Degradation was also quantified by the chemistry of the peptide termini, including (**F**) Ac-βA, (**G**) Ac, (**H**) N-βA, (**I**) $NH_2$, (**J**) C-βA, (**K**) Am, (**L**) COOH.

A hMSC: Ac-βA

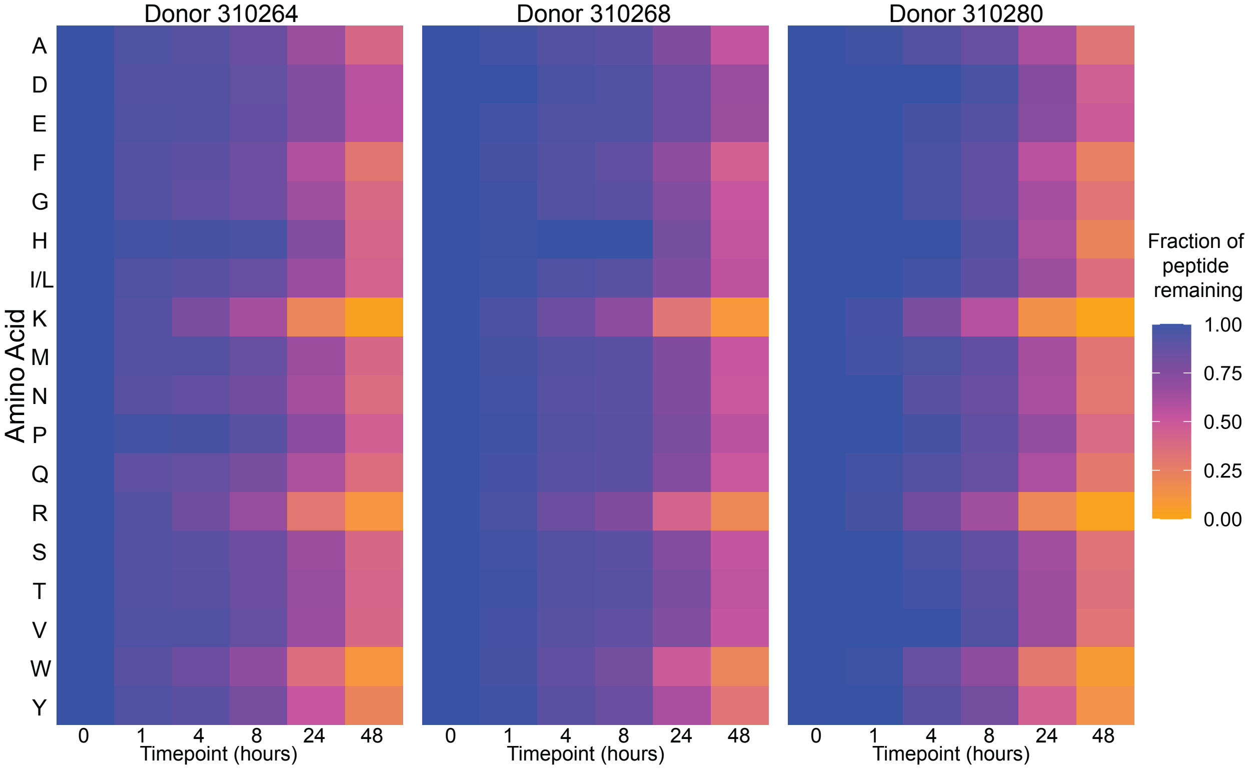


B hMSC: Ac

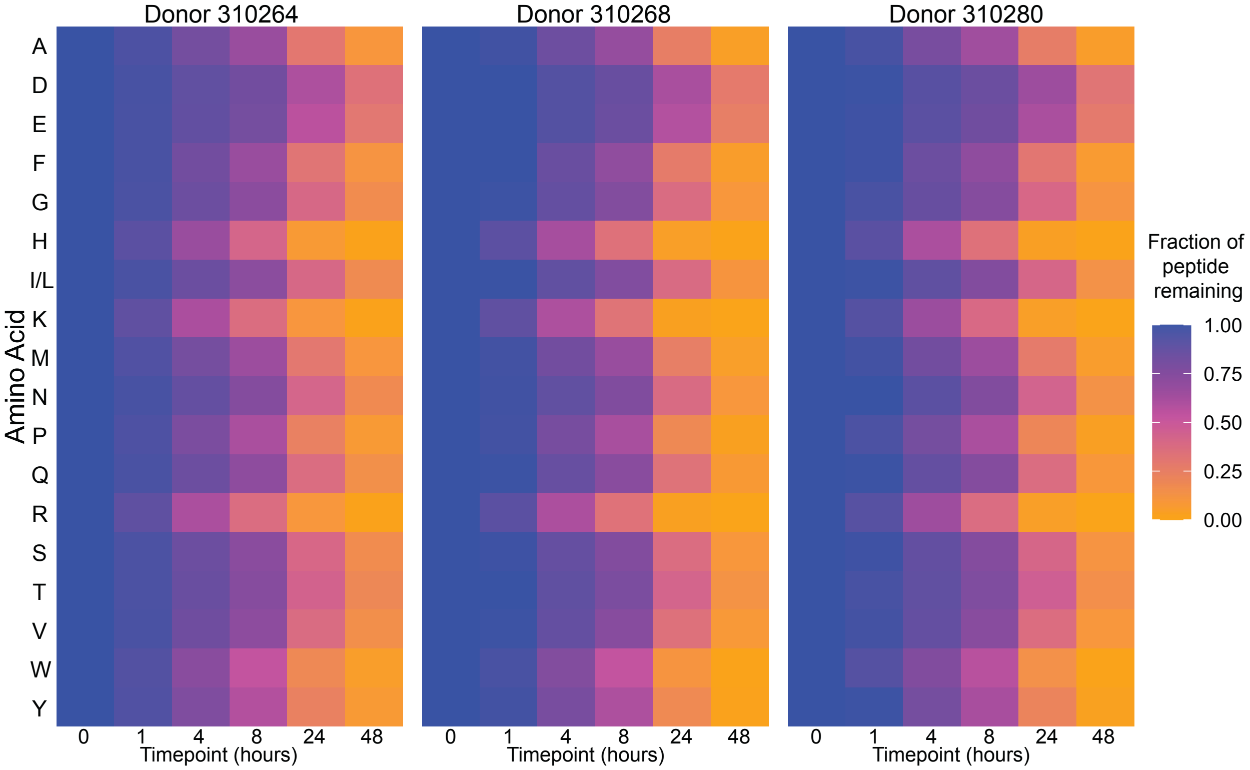

C hMSC: N-βA

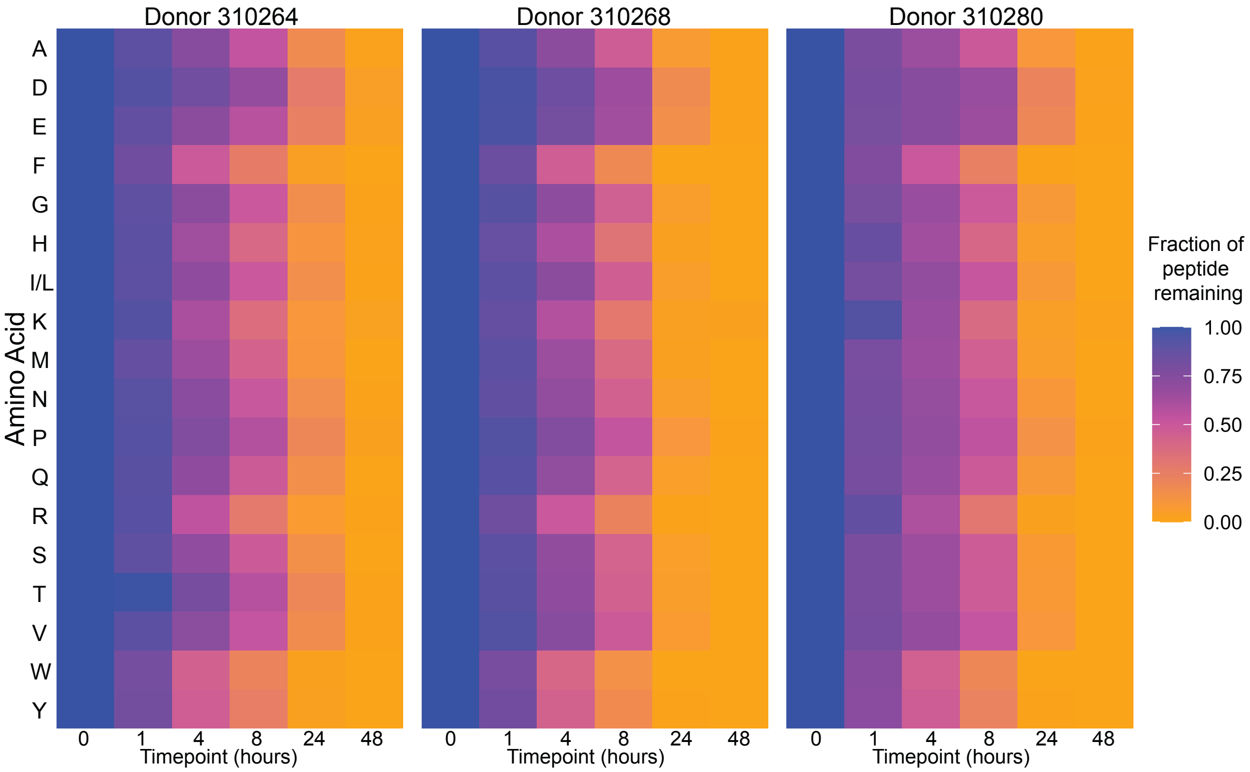


D hMSC: $NH_2$

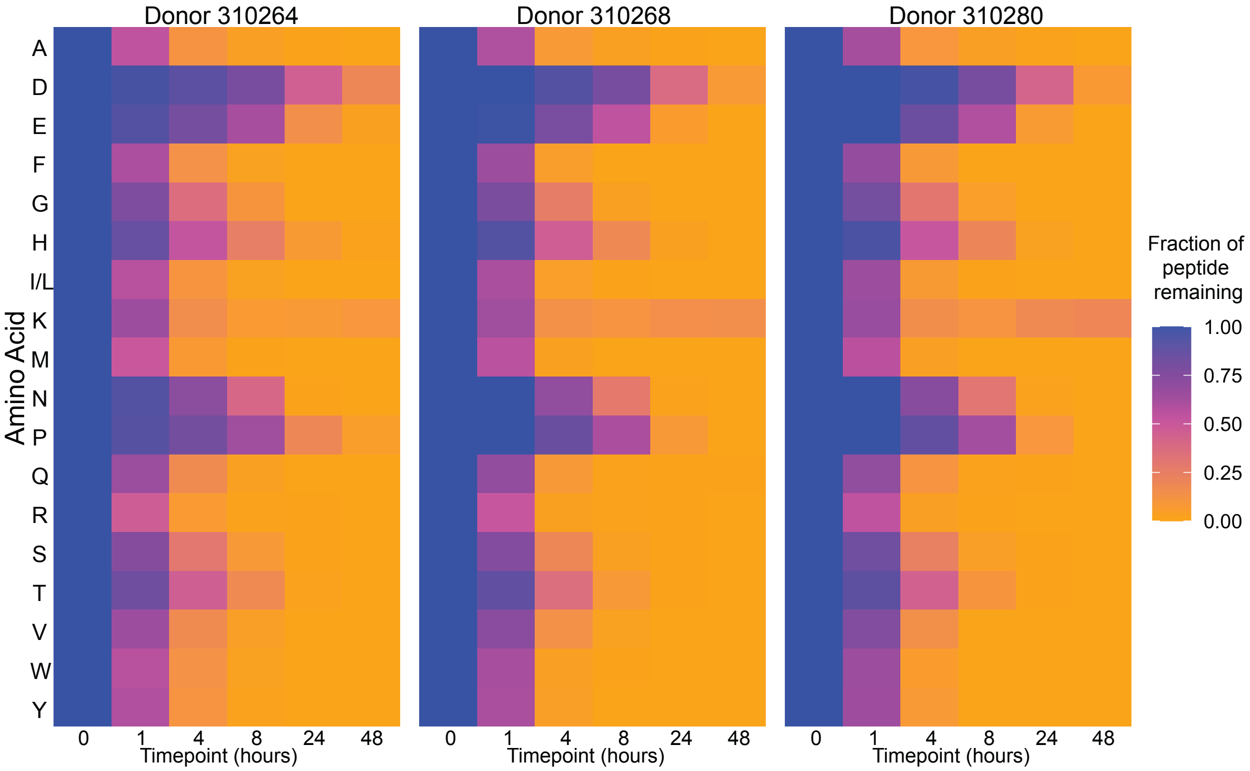

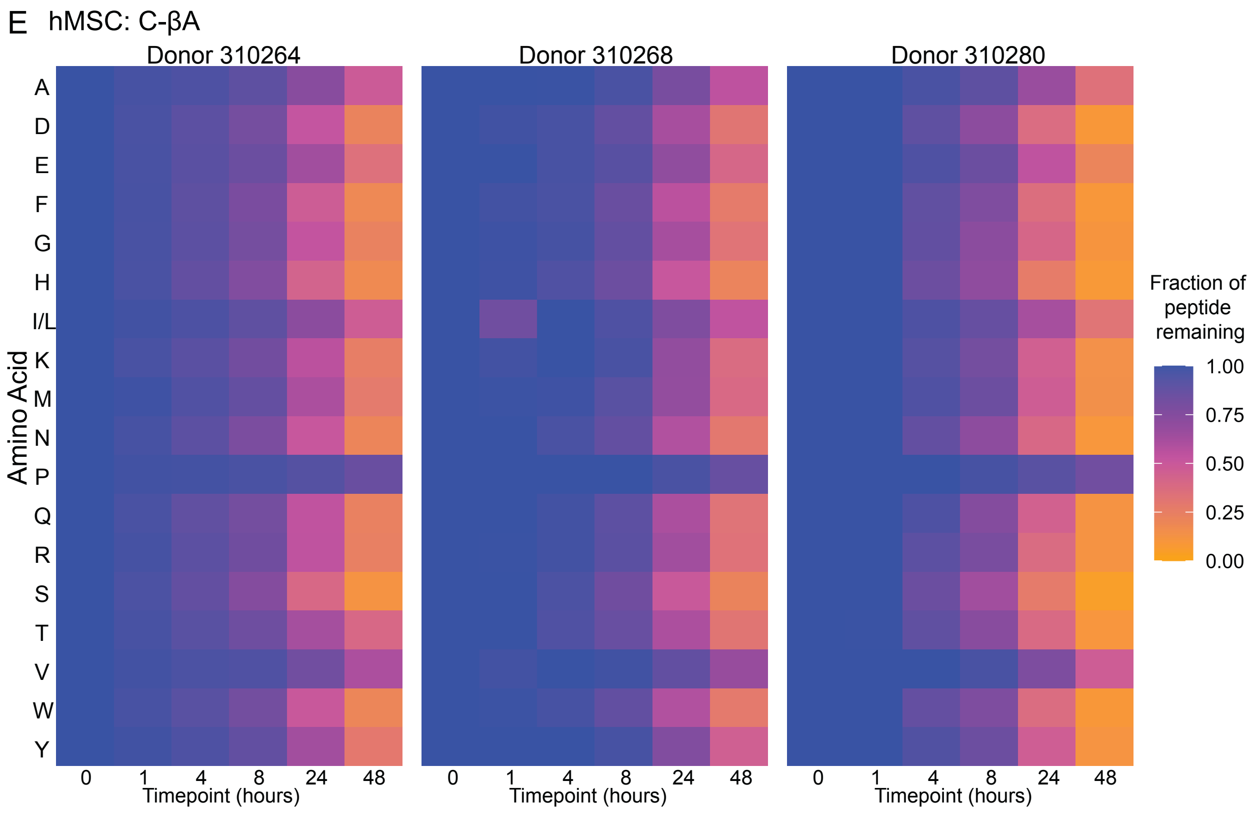
E hMSC: C-βA
Donor 310264
Donor 310268
Donor 310280
Amino Acid
A
D
E
F
G
H
I/L
K
M
N
P
Q
R
S
T
V
W
Y
0 1 4 8 24 48
Timepoint (hours)
Fraction of peptide remaining
1.00
0.75
0.50
0.25
0.00

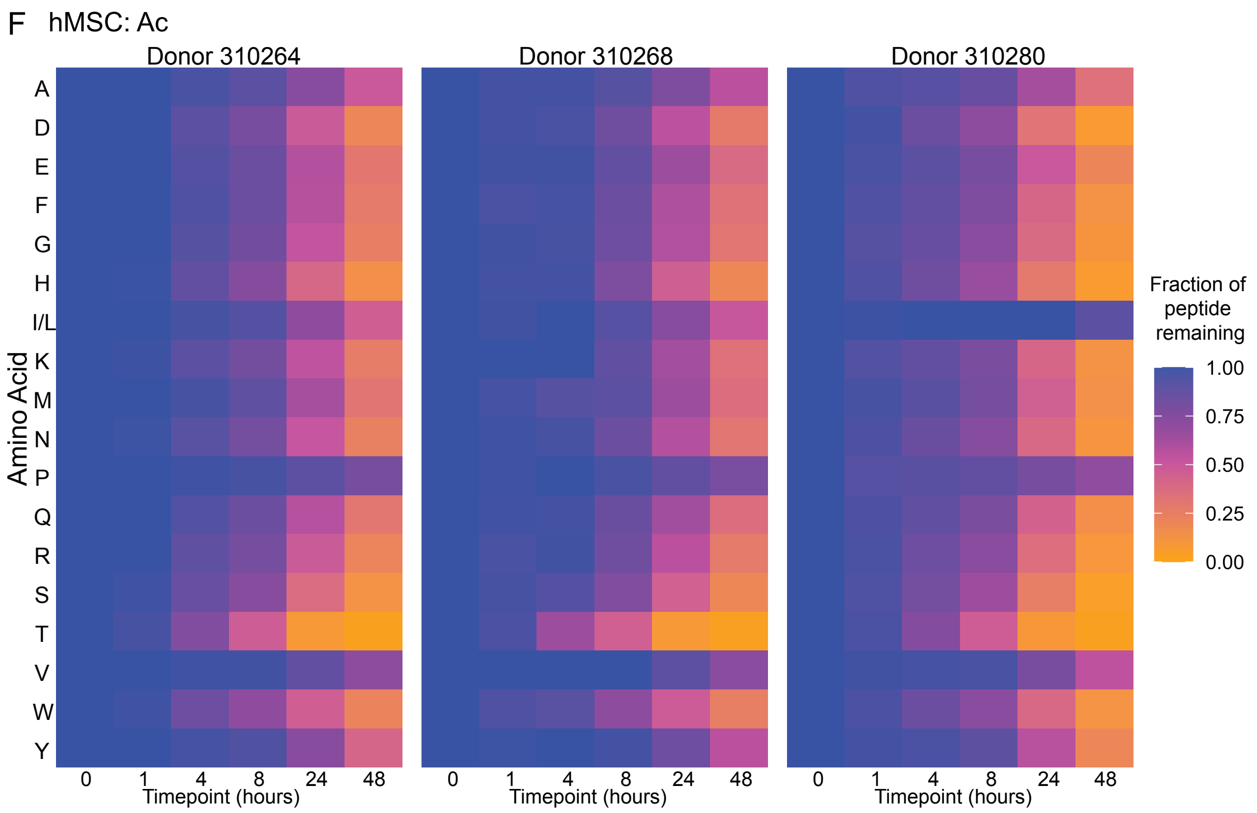
F hMSC: Ac
Donor 310264
Donor 310268
Donor 310280
Amino Acid
A
D
E
F
G
H
I/L
K
M
N
P
Q
R
S
T
V
W
Y
0 1 4 8 24 48
Timepoint (hours)
Fraction of peptide remaining
1.00
0.75
0.50
0.25
0.00

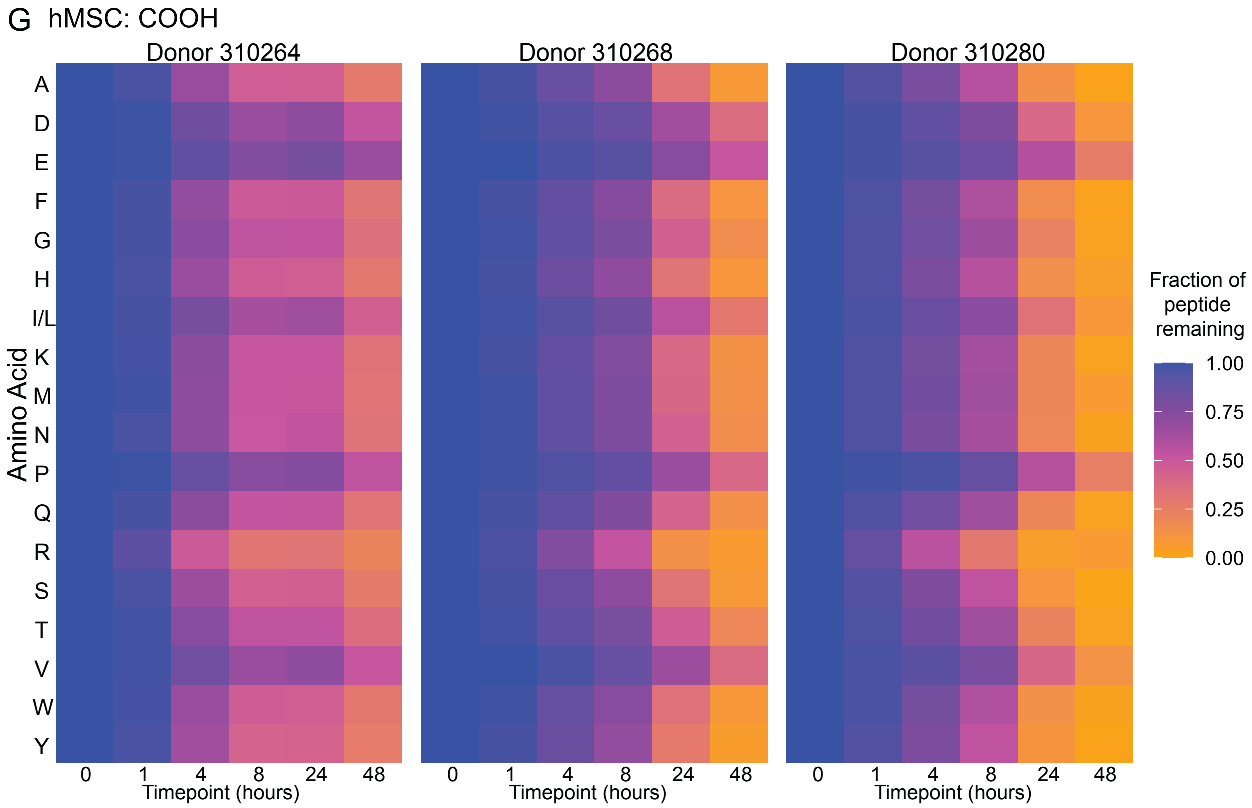


**Figure S2**. Comparison of peptide degradation by different hMSC donors. hMSCs were cultured with (**A**) Ac-βA, (**B**) Ac, (**C**) N-βA, (**D**) $NH_2$, (**E**) C-βA, (**F**) Am, (**G**) COOH.

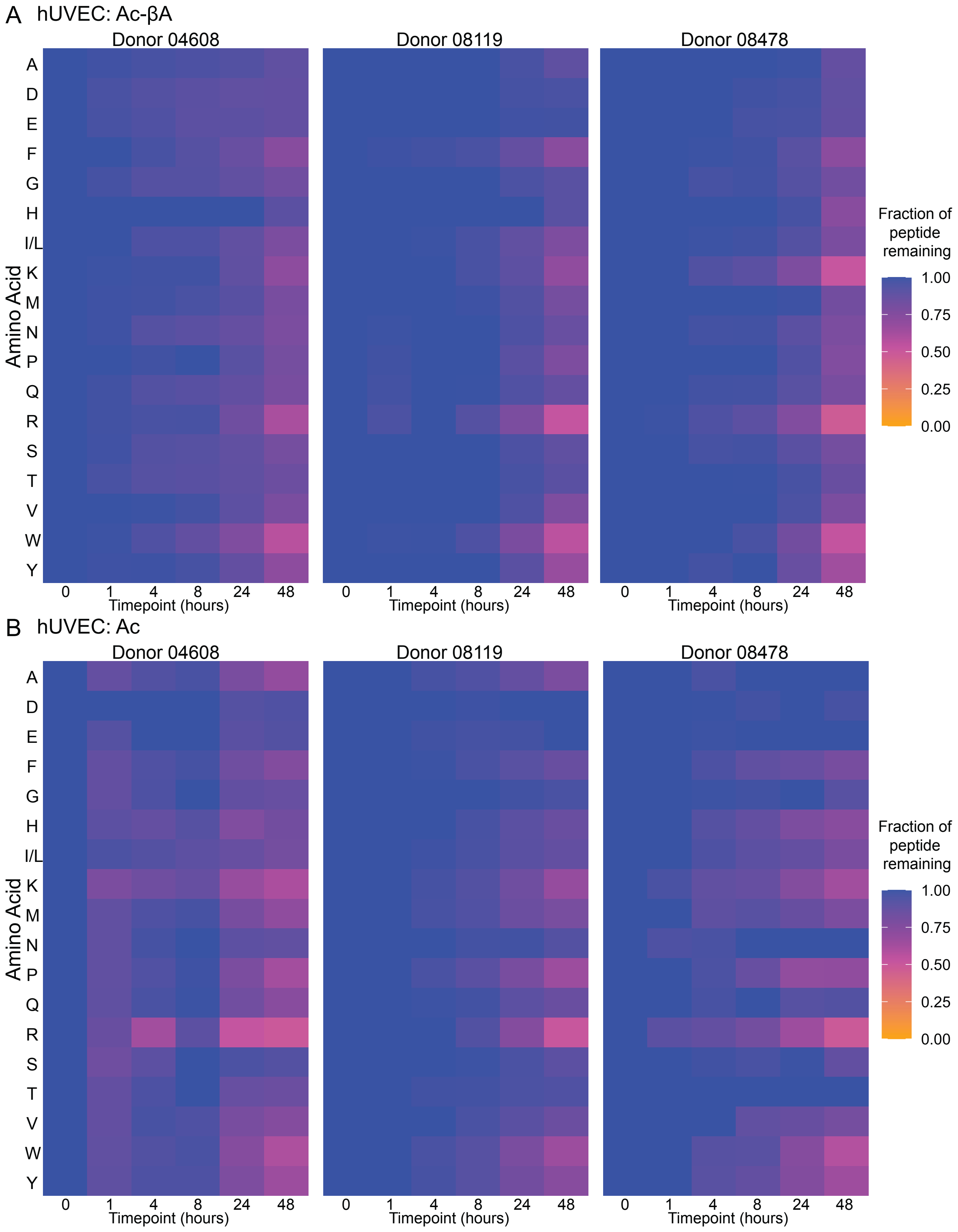
A hUVEC: Ac-βA
Donor 04608
Donor 08119
Donor 08478
Amino Acid
A
D
E
F
G
H
I/L
K
M
N
P
Q
R
S
T
V
W
Y
0 1 4 8 24 48
Timepoint (hours)
Fraction of peptide remaining
1.00
0.75
0.50
0.25
0.00
B hUVEC: Ac
Donor 04608
Donor 08119
Donor 08478
Amino Acid
Timepoint (hours)
Fraction of peptide remaining

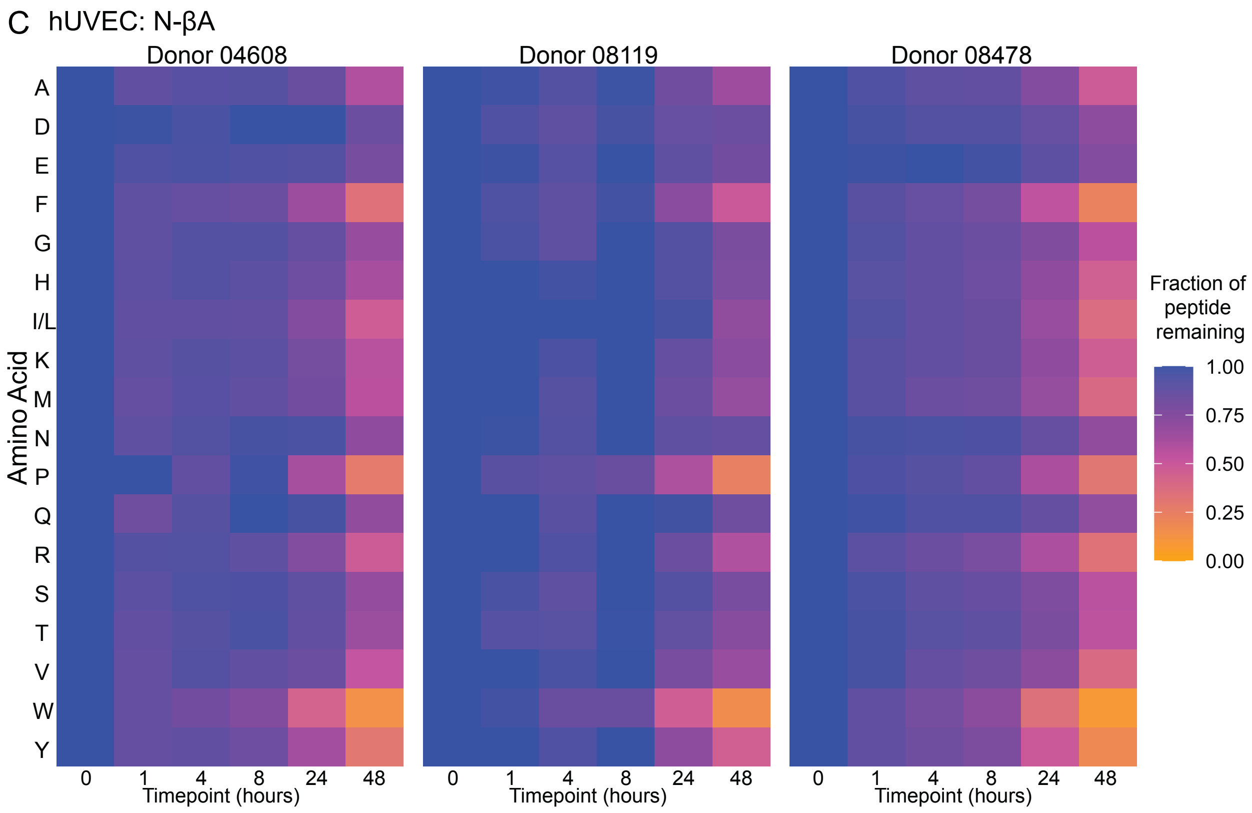
C hUVEC: N-βA
Donor 04608
Donor 08119
Donor 08478
Amino Acid
A
D
E
F
G
H
I/L
K
M
N
P
Q
R
S
T
V
W
Y
0 1 4 8 24 48
Timepoint (hours)
Fraction of peptide remaining
1.00
0.75
0.50
0.25
0.00

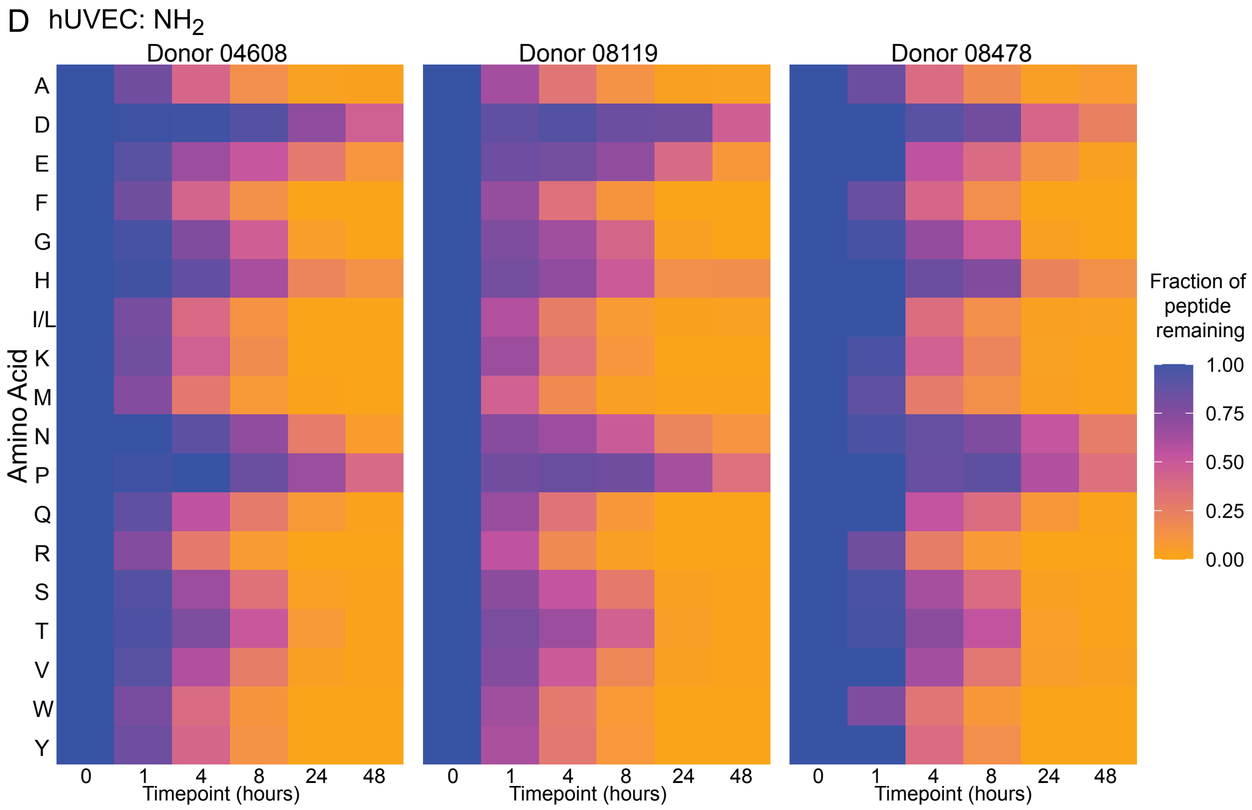
D hUVEC: NH2
Donor 04608
Donor 08119
Donor 08478
Amino Acid
A
D
E
F
G
H
I/L
K
M
N
P
Q
R
S
T
V
W
Y
0 1 4 8 24 48
Timepoint (hours)
Fraction of peptide remaining
1.00
0.75
0.50
0.25
0.00

E hUVEC: C-βA

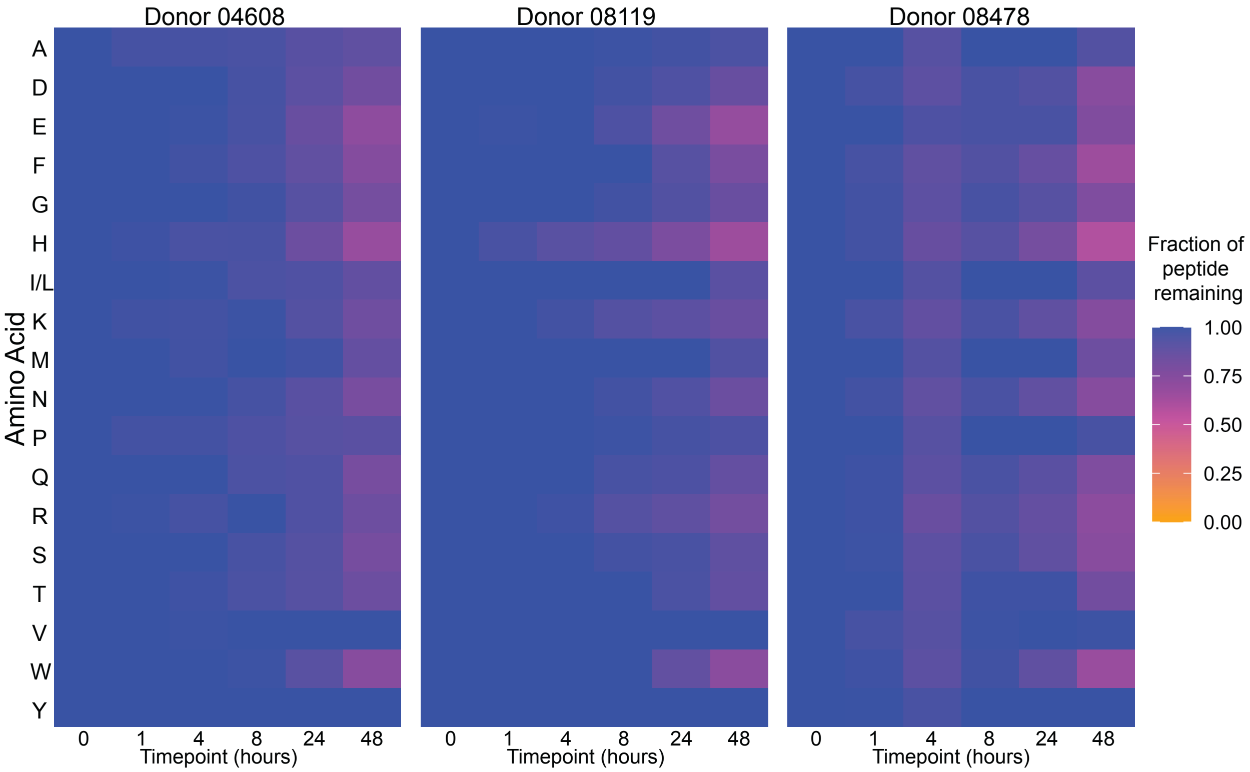


F hUVEC: Am

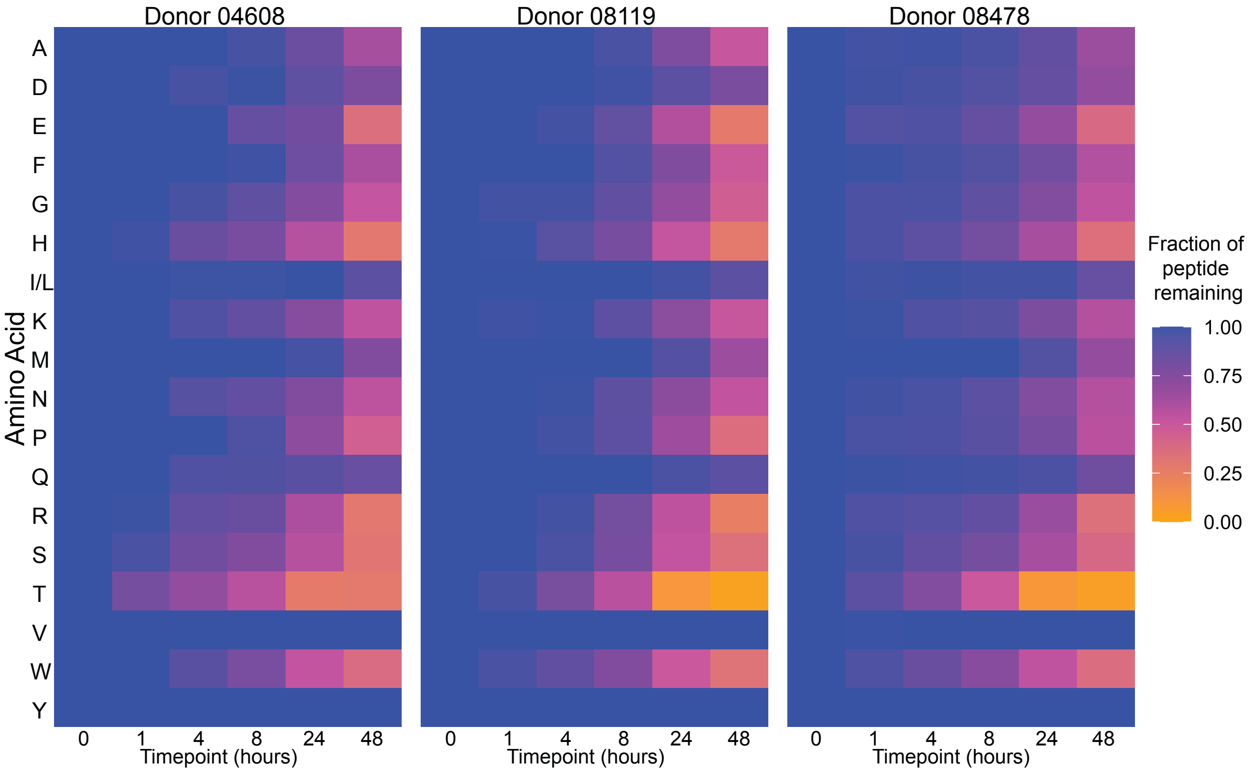

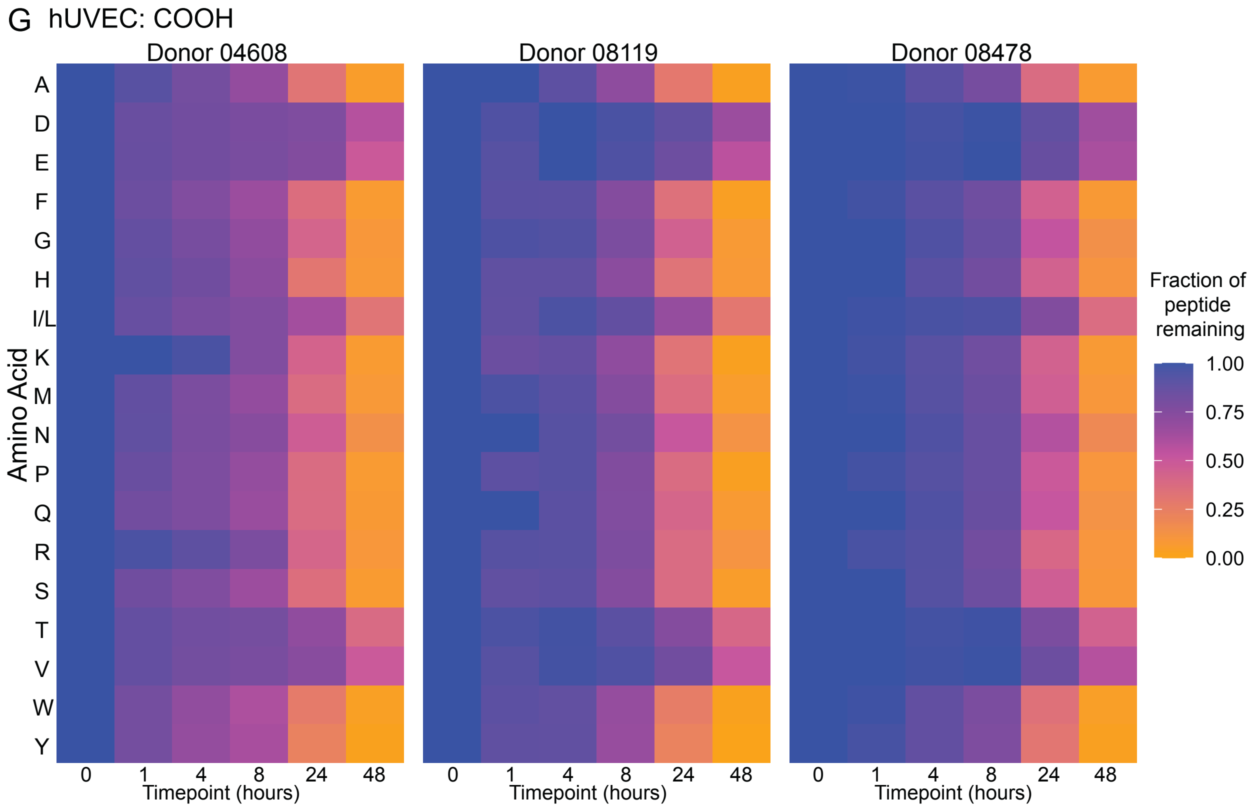


**Figure S3**. Comparison of peptide degradation by different hUVEC donors. hUVECs were cultured with (**A**) Ac-βA, (**B**) Ac, (**C**) N-βA, (**D**) $NH_2$, (**E**) C-βA, (**F**) Am, (**G**) COOH.

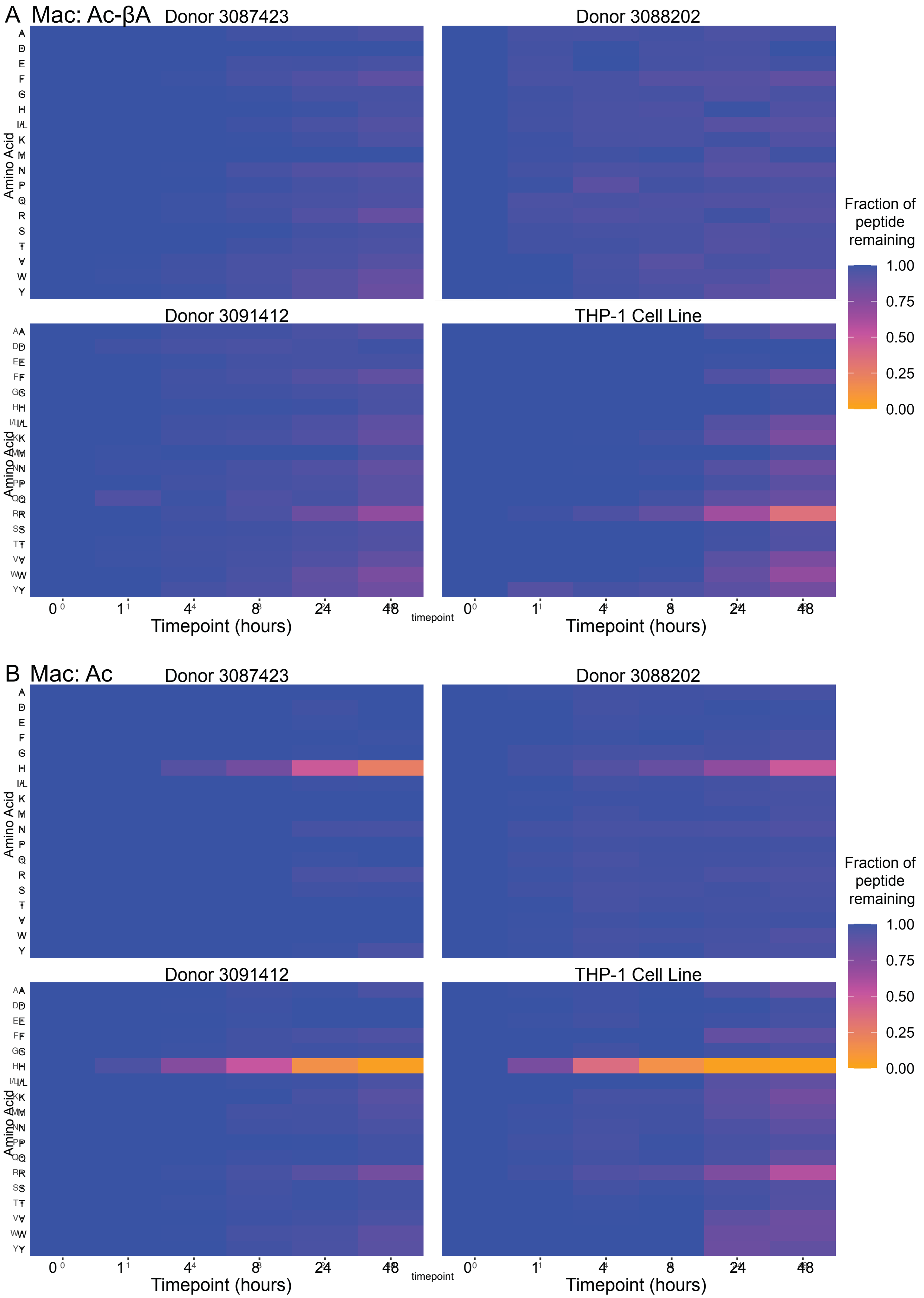
A Mac: Ac-βA
Donor 3087423
Donor 3088202
Donor 3091412
THP-1 Cell Line
Amino Acid
A
D
E
F
G
H
I/L
K
M
N
P
Q
R
S
T
V
W
Y
Fraction of peptide remaining
1.00
0.75
0.50
0.25
0.00
0
1
4
8
24
48
timepoint
Timepoint (hours)
B Mac: Ac
Donor 3087423
Donor 3088202
Donor 3091412
THP-1 Cell Line
Fraction of peptide remaining
Timepoint (hours)

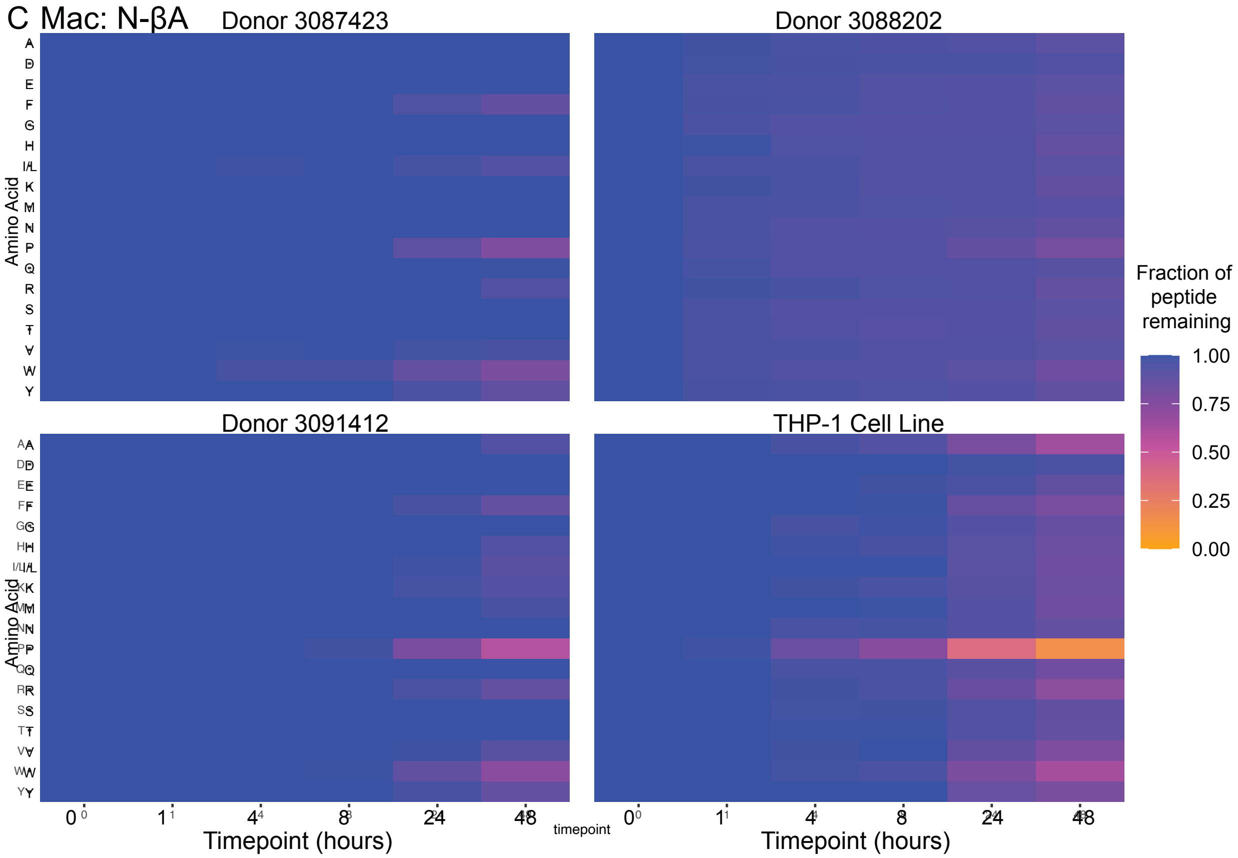
C Mac: N-βA
Donor 3087423
Donor 3088202
Donor 3091412
THP-1 Cell Line
Amino Acid
A
D
E
F
G
H
I/L
K
M
N
P
Q
R
S
T
V
W
Y
Fraction of peptide remaining
1.00
0.75
0.50
0.25
0.00
0
1
4
8
24
48
timepoint
Timepoint (hours)

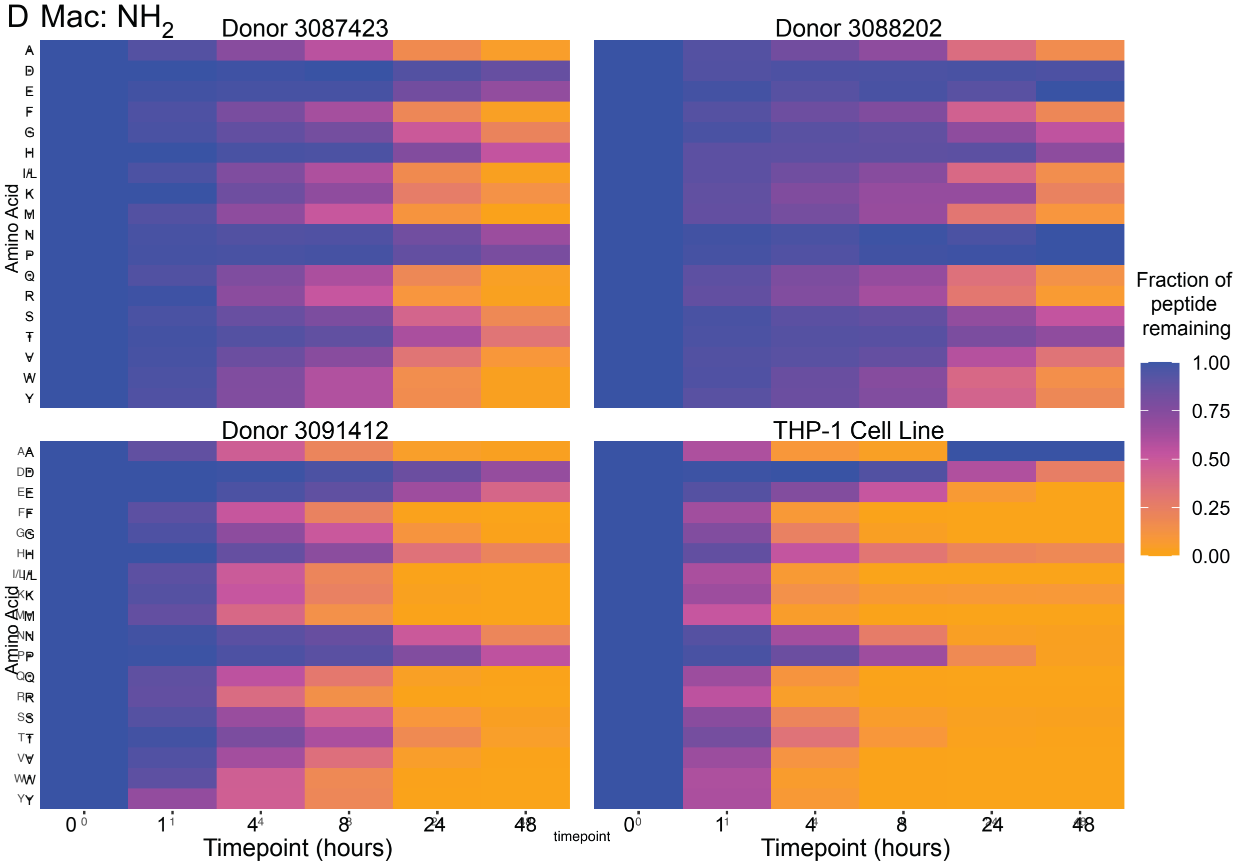
D Mac: NH2
Donor 3087423
Donor 3088202
Donor 3091412
THP-1 Cell Line
Amino Acid
A
D
E
F
G
H
I/L
K
M
N
P
Q
R
S
T
V
W
Y
Fraction of peptide remaining
1.00
0.75
0.50
0.25
0.00
0
1
4
8
24
48
timepoint
Timepoint (hours)

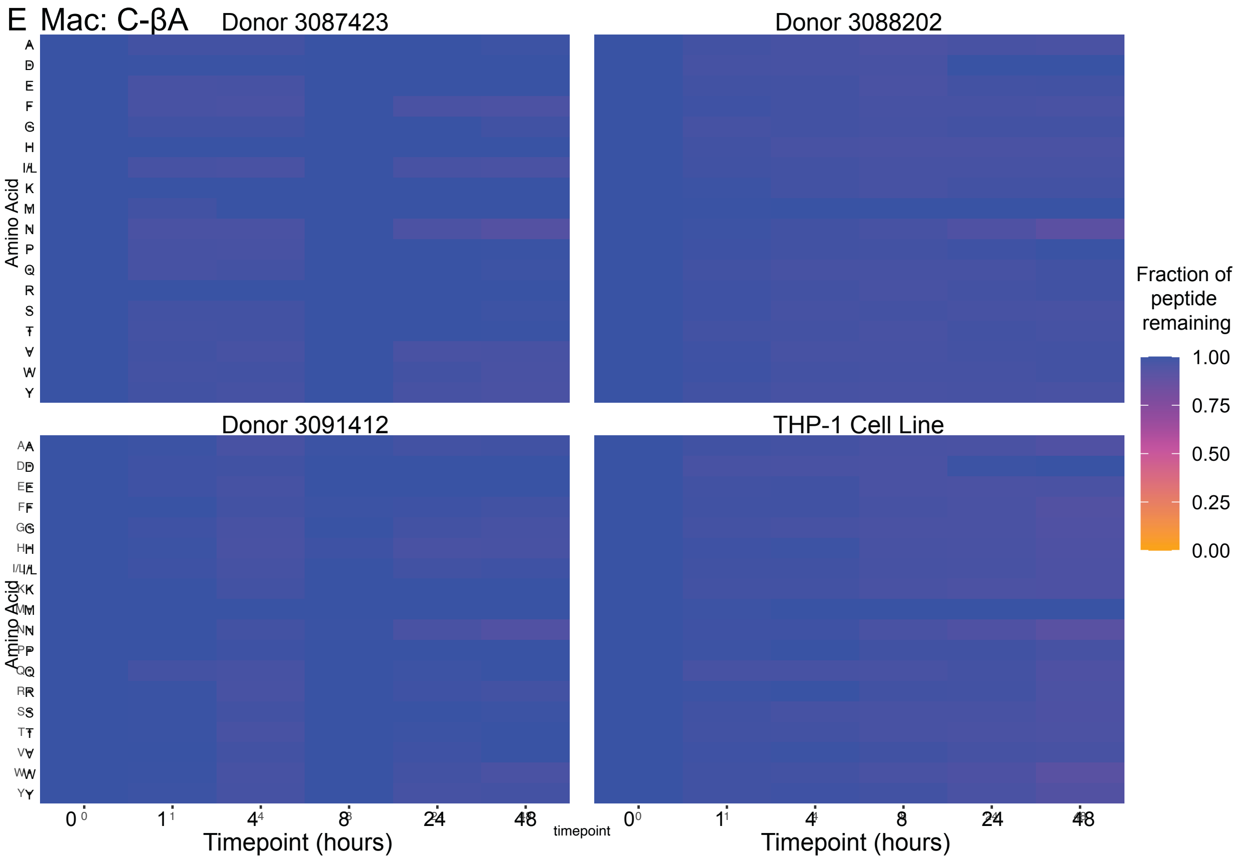
E Mac: C-βA
Donor 3087423
Donor 3088202
Donor 3091412
THP-1 Cell Line
Amino Acid
A
D
E
F
G
H
I/L
K
M
N
P
Q
R
S
T
V
W
Y
Fraction of peptide remaining
1.00
0.75
0.50
0.25
0.00
0
1
4
8
24
48
timepoint
Timepoint (hours)

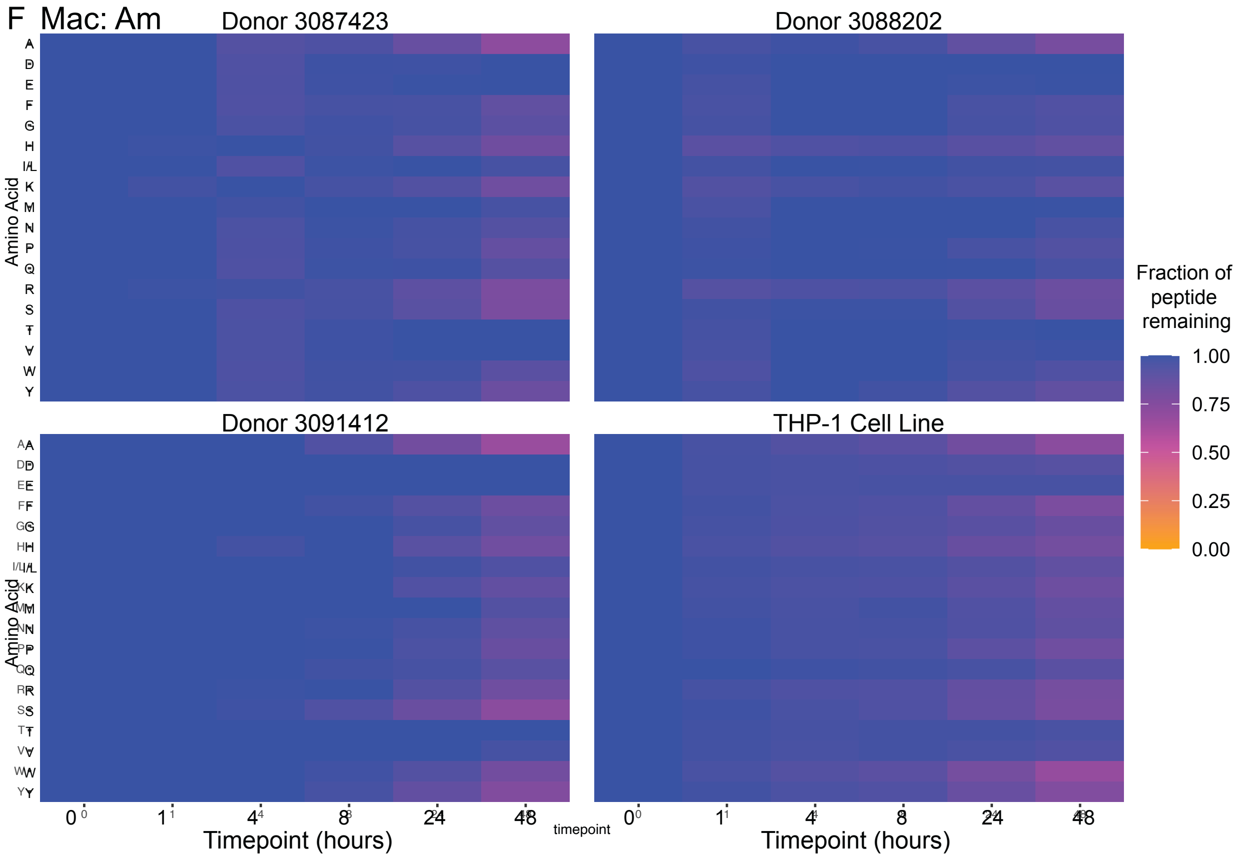
F Mac: Am
Donor 3087423
Donor 3088202
Donor 3091412
THP-1 Cell Line
Amino Acid
A
D
E
F
G
H
I/L
K
M
N
P
Q
R
S
T
V
W
Y
Fraction of peptide remaining
1.00
0.75
0.50
0.25
0.00
0
1
4
8
24
48
timepoint
Timepoint (hours)

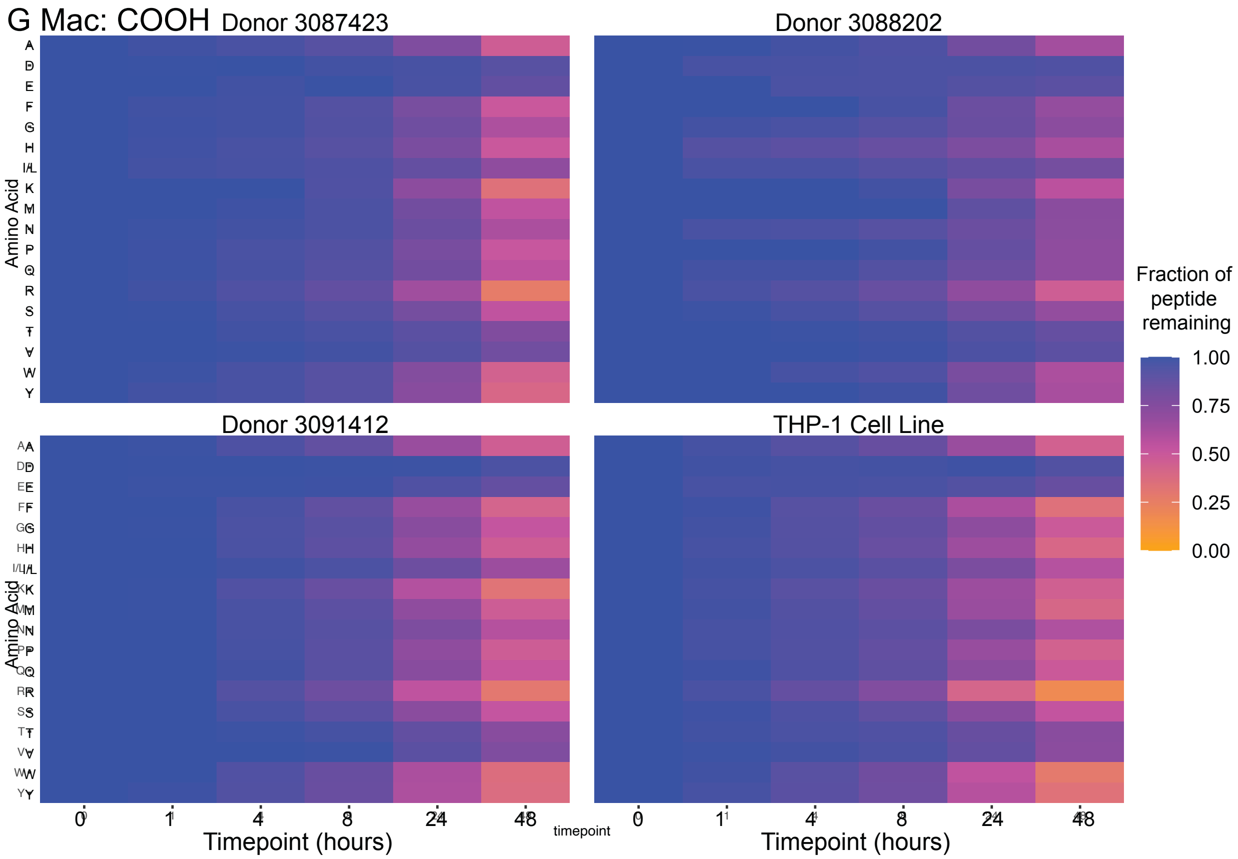


**Figure S4**. Comparison of peptide degradation by different donors. Macrophages were cultured with (**A**) Ac-βA, (**B**) Ac, (**C**) N-βA, (**D**) $NH_2$, (**E**) C-βA, (**F**) Am, (**G**) COOH.

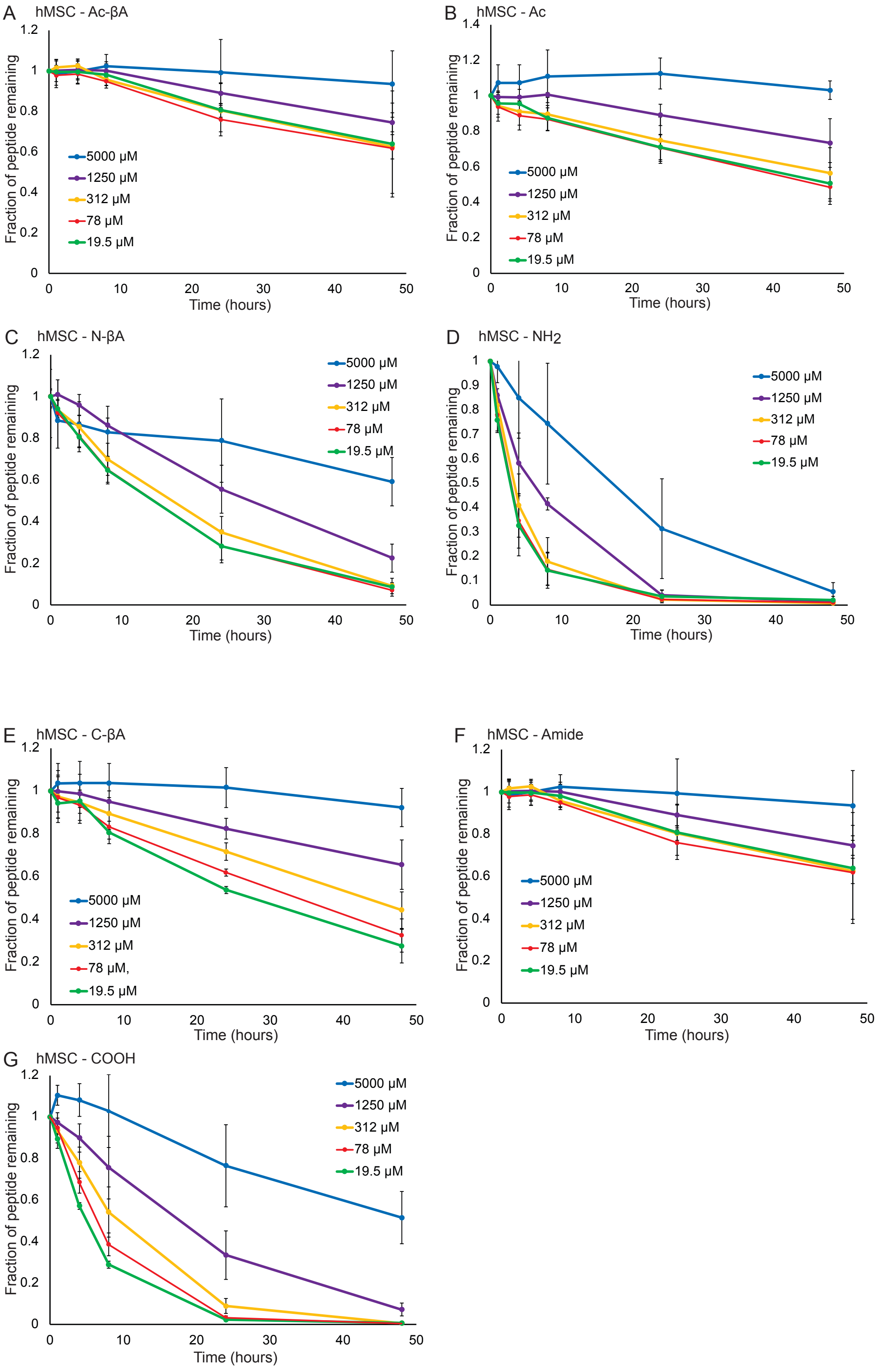
A
hMSC - Ac-βA
Fraction of peptide remaining
Time (hours)
5000 μM
1250 μM
312 μM
78 μM
19.5 μM
B
hMSC - Ac
Fraction of peptide remaining
Time (hours)
5000 μM
1250 μM
312 μM
78 μM
19.5 μM
C
hMSC - N-βA
Fraction of peptide remaining
Time (hours)
5000 μM
1250 μM
312 μM
78 μM
19.5 μM
D
hMSC - NH2
Fraction of peptide remaining
Time (hours)
5000 μM
1250 μM
312 μM
78 μM
19.5 μM
E
hMSC - C-βA
Fraction of peptide remaining
Time (hours)
5000 μM
1250 μM
312 μM
78 μM,
19.5 μM
F
hMSC - Amide
Fraction of peptide remaining
Time (hours)
5000 μM
1250 μM
312 μM
78 μM
19.5 μM
G
hMSC - COOH
Fraction of peptide remaining
Time (hours)
5000 μM
1250 μM
312 μM
78 μM
19.5 μM

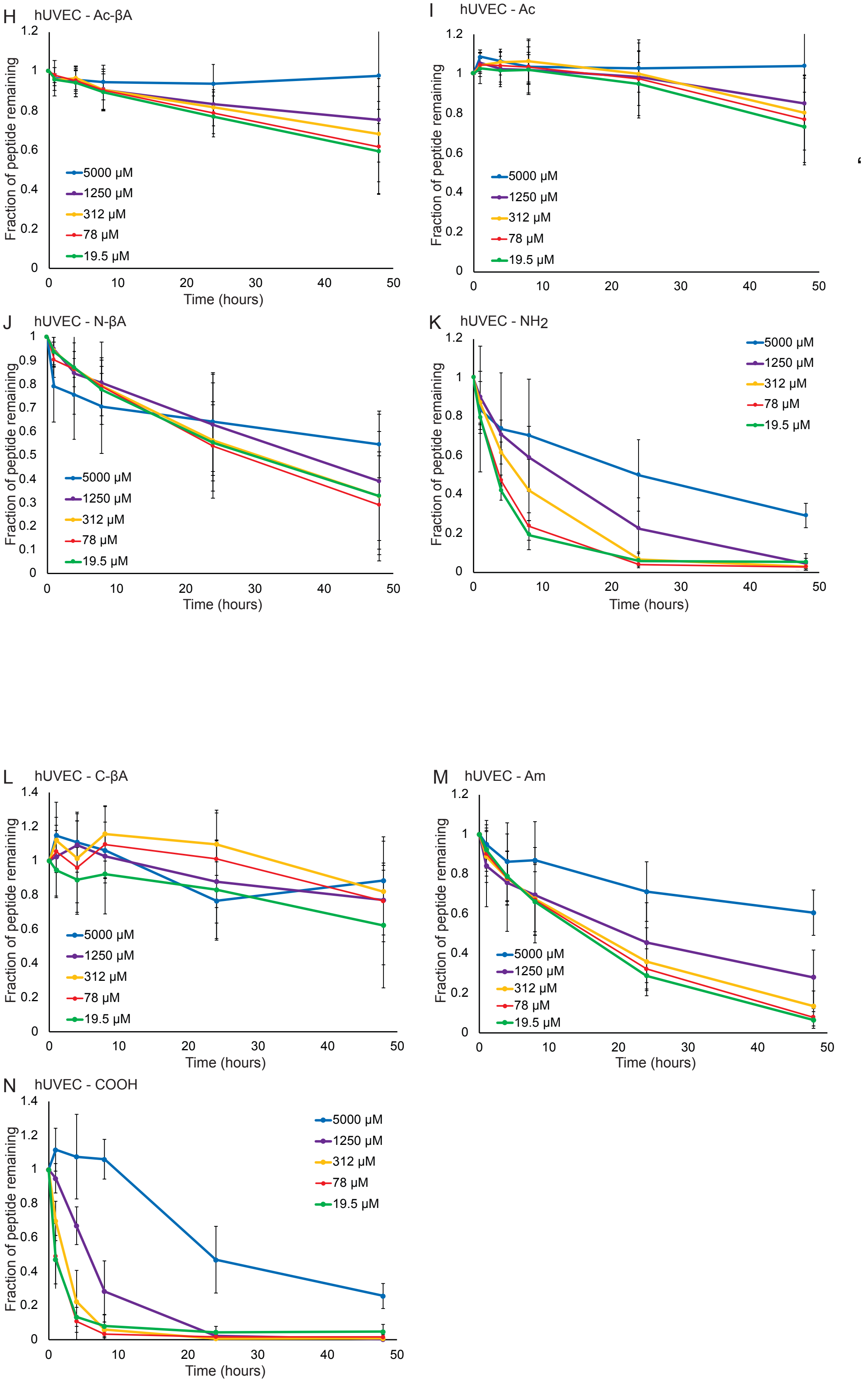
H
hUVEC - Ac-βA
Fraction of peptide remaining
Time (hours)
5000 µM
1250 µM
312 µM
78 µM
19.5 µM
I
hUVEC - Ac
Fraction of peptide remaining
5000 µM
1250 µM
312 µM
78 µM
19.5 µM
J
hUVEC - N-βA
Fraction of peptide remaining
Time (hours)
5000 µM
1250 µM
312 µM
78 µM
19.5 µM
K
hUVEC - NH2
Fraction of peptide remaining
Time (hours)
5000 µM
1250 µM
312 µM
78 µM
19.5 µM
L
hUVEC - C-βA
Fraction of peptide remaining
Time (hours)
5000 µM
1250 µM
312 µM
78 µM
19.5 µM
M
hUVEC - Am
Fraction of peptide remaining
Time (hours)
5000 µM
1250 µM
312 µM
78 µM
19.5 µM
N
hUVEC - COOH
Fraction of peptide remaining
Time (hours)
5000 µM
1250 µM
312 µM
78 µM
19.5 µM

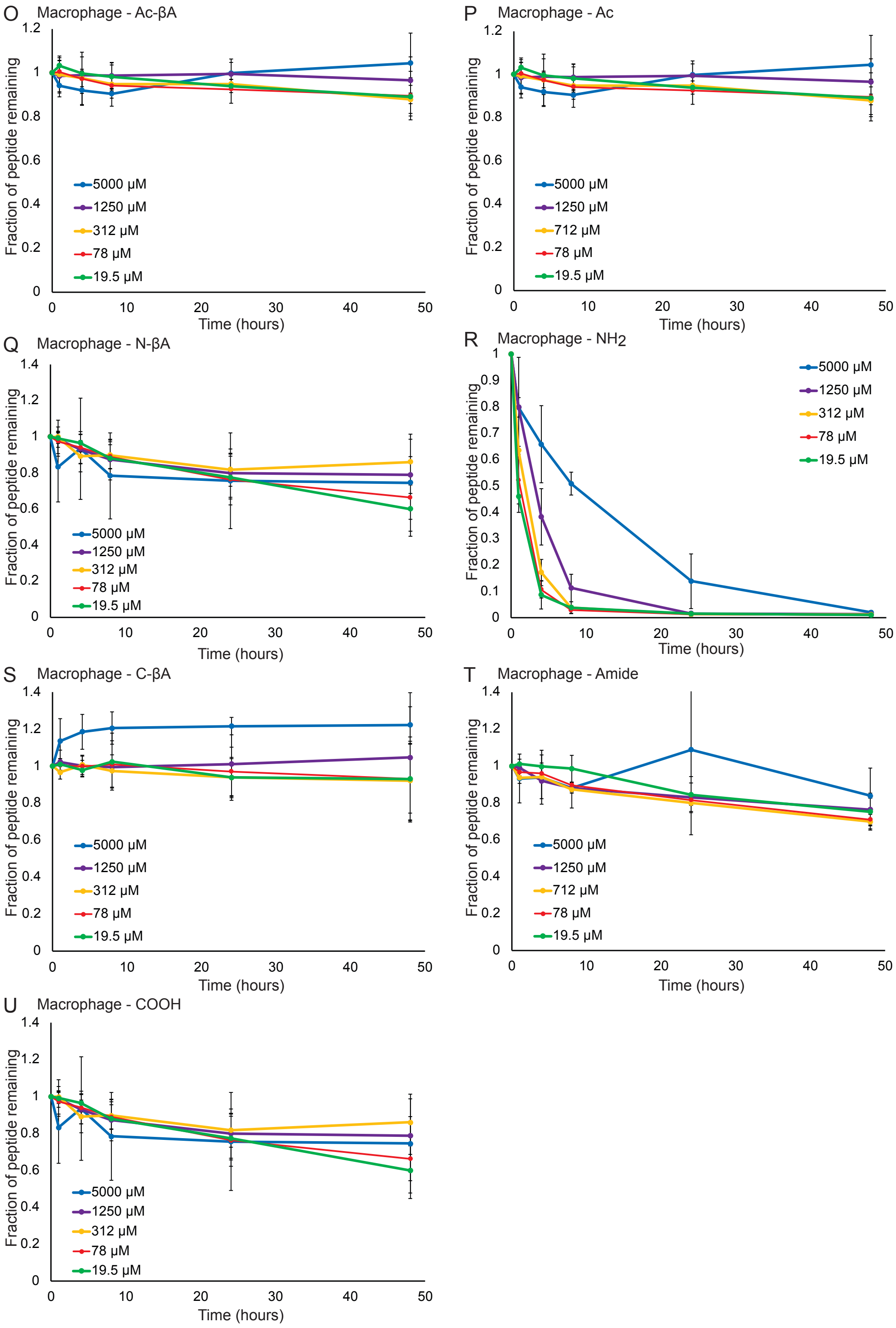


**Figure S5**. Degradation of peptides at different concentrations. Data was quantified for hMSCs (**A**) Ac-βA, (**B**) Ac, (**C**) N-βA, (**D**) $NH_2$, (**E**) C-βA, (**F**) Am, (**G**) COOH, hUVECs (**H**) Ac-βA, (**I**) Ac, (**J**) N-βA, (**K**) $NH_2$, (**L**) C-βA, (**M**) Am, (**N**) COOH, and macrophages (**O**) Ac-βA, (**P**) Ac, (**Q**) N-βA, (**R**) $NH_2$, (**S**) C-βA, (**T**) Am, (**U**) COOH.

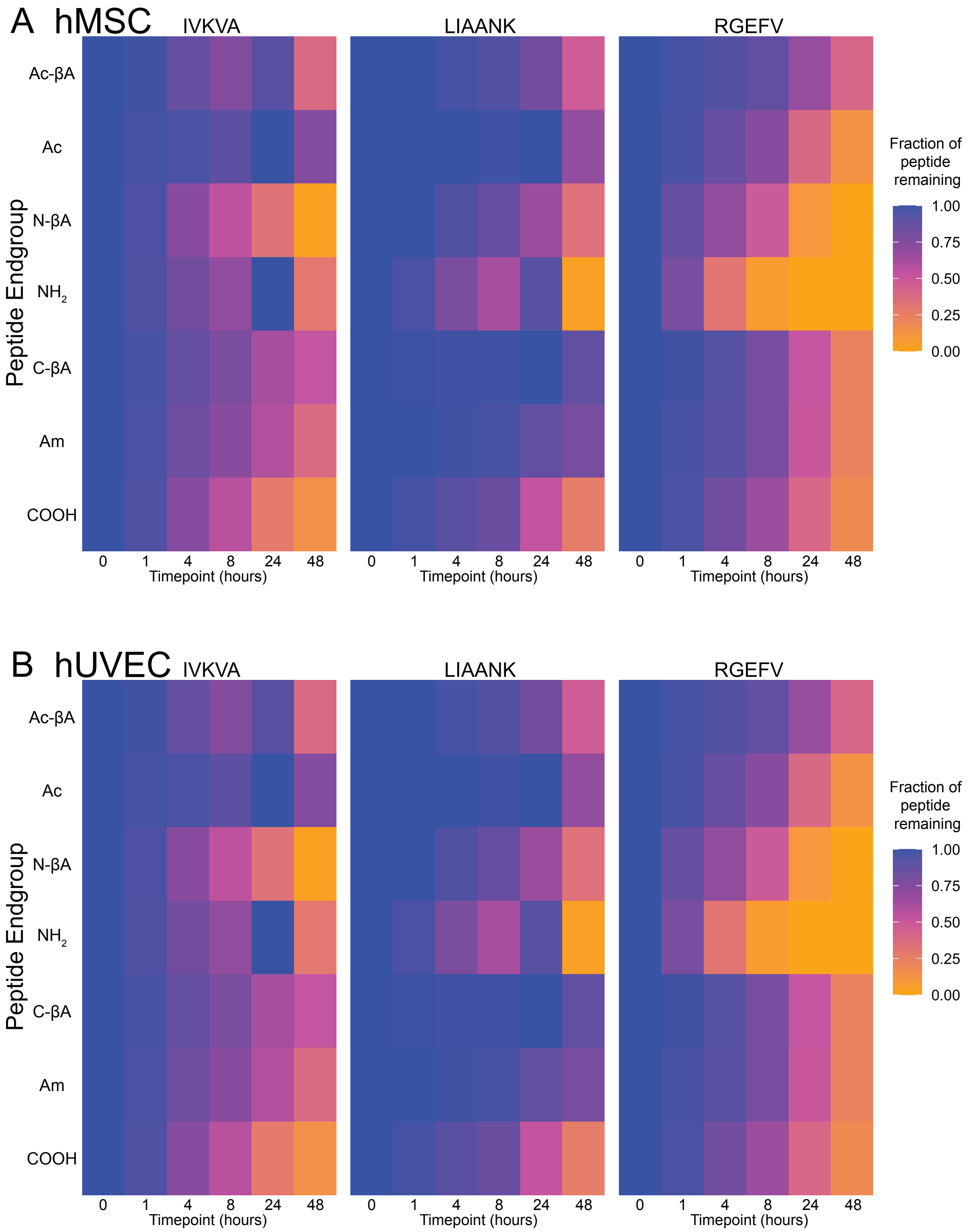

A hMSC
IVKVA
LIAANK
RGEFV
Peptide Endgroup
Ac-βA
Ac
N-βA
NH2
C-βA
Am
COOH
0
1
4
8
24
48
Timepoint (hours)
Fraction of peptide remaining
1.00
0.75
0.50
0.25
0.00
B hUVEC

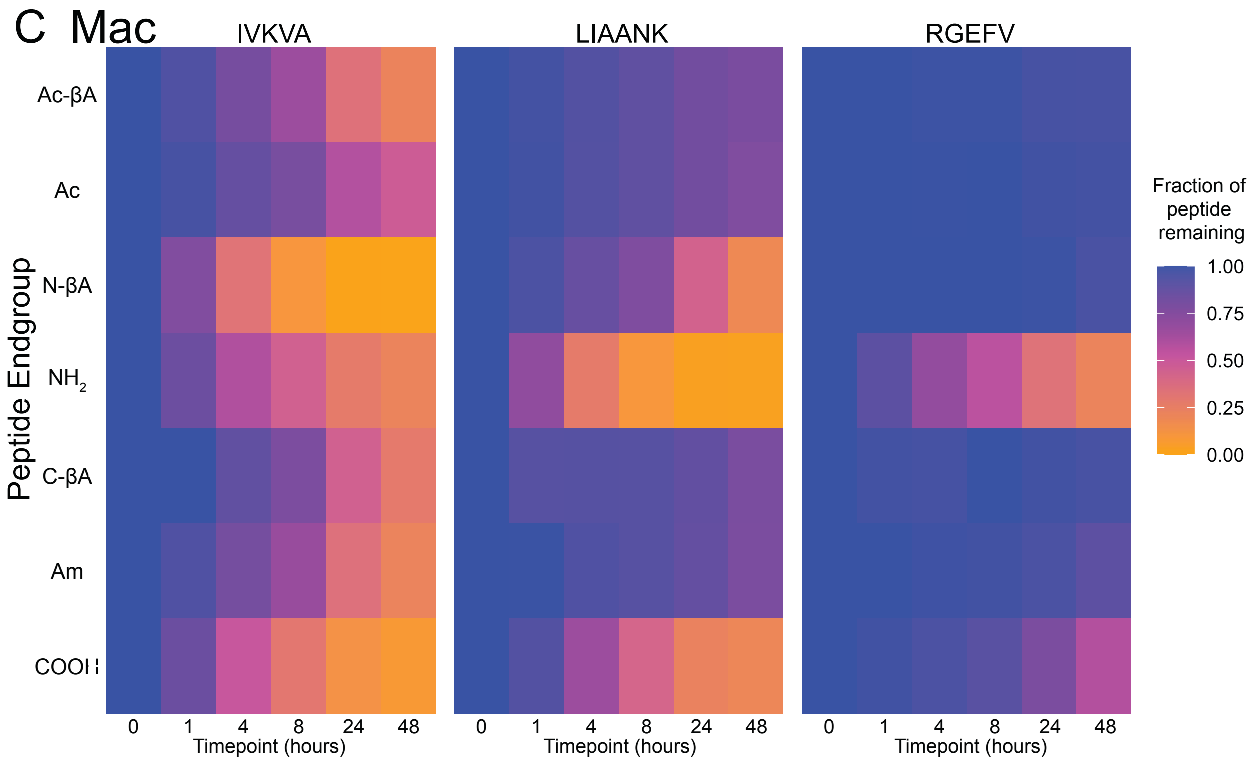
C Mac
IVKVA
LIAANK
RGEFV
Peptide Endgroup
Ac-βA
Ac
N-βA
NH2
C-βA
Am
COOH
0
1
4
8
24
48
Timepoint (hours)
Fraction of peptide remaining
1.00
0.75
0.50
0.25
0.00

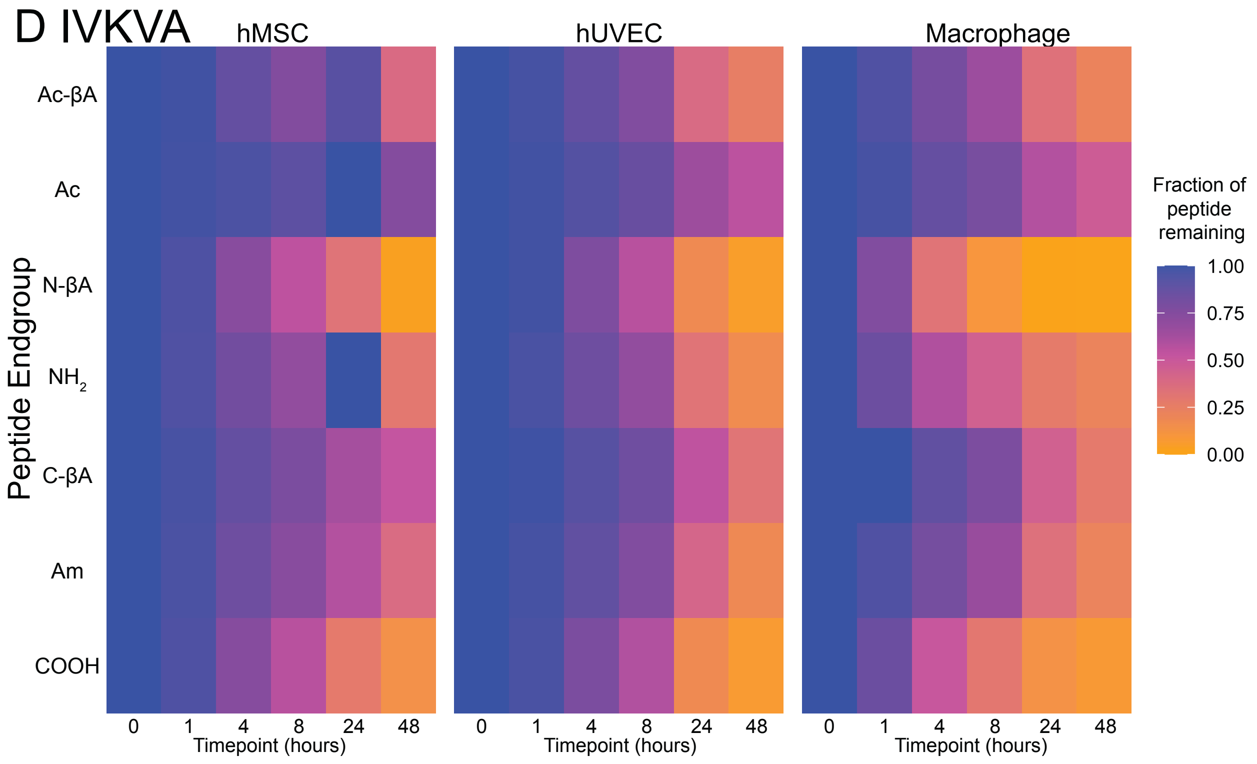
D IVKVA
hMSC
hUVEC
Macrophage
Peptide Endgroup
Ac-βA
Ac
N-βA
NH2
C-βA
Am
COOH
0
1
4
8
24
48
Timepoint (hours)
Fraction of peptide remaining
1.00
0.75
0.50
0.25
0.00

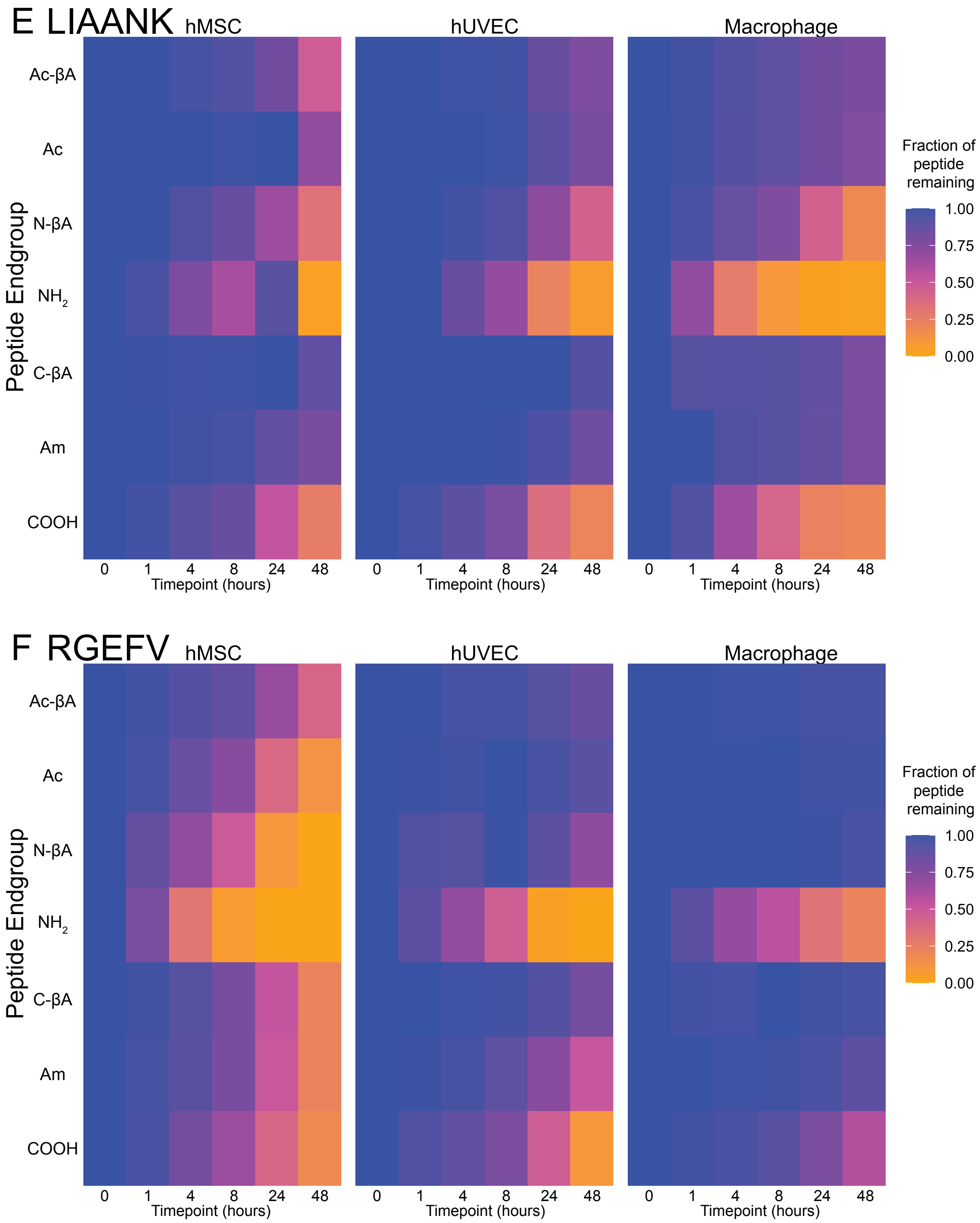


**Figure S6**. Effect of different peptide sequences on non-specific degradation. Degradation was quantified for the cell types (**A**) hMSCs, (**B**) hUVECs, (**C**), macrophages, and peptides (**D**) IVKVA, (**E**) LIAANK, and (**F**) RGEFV.

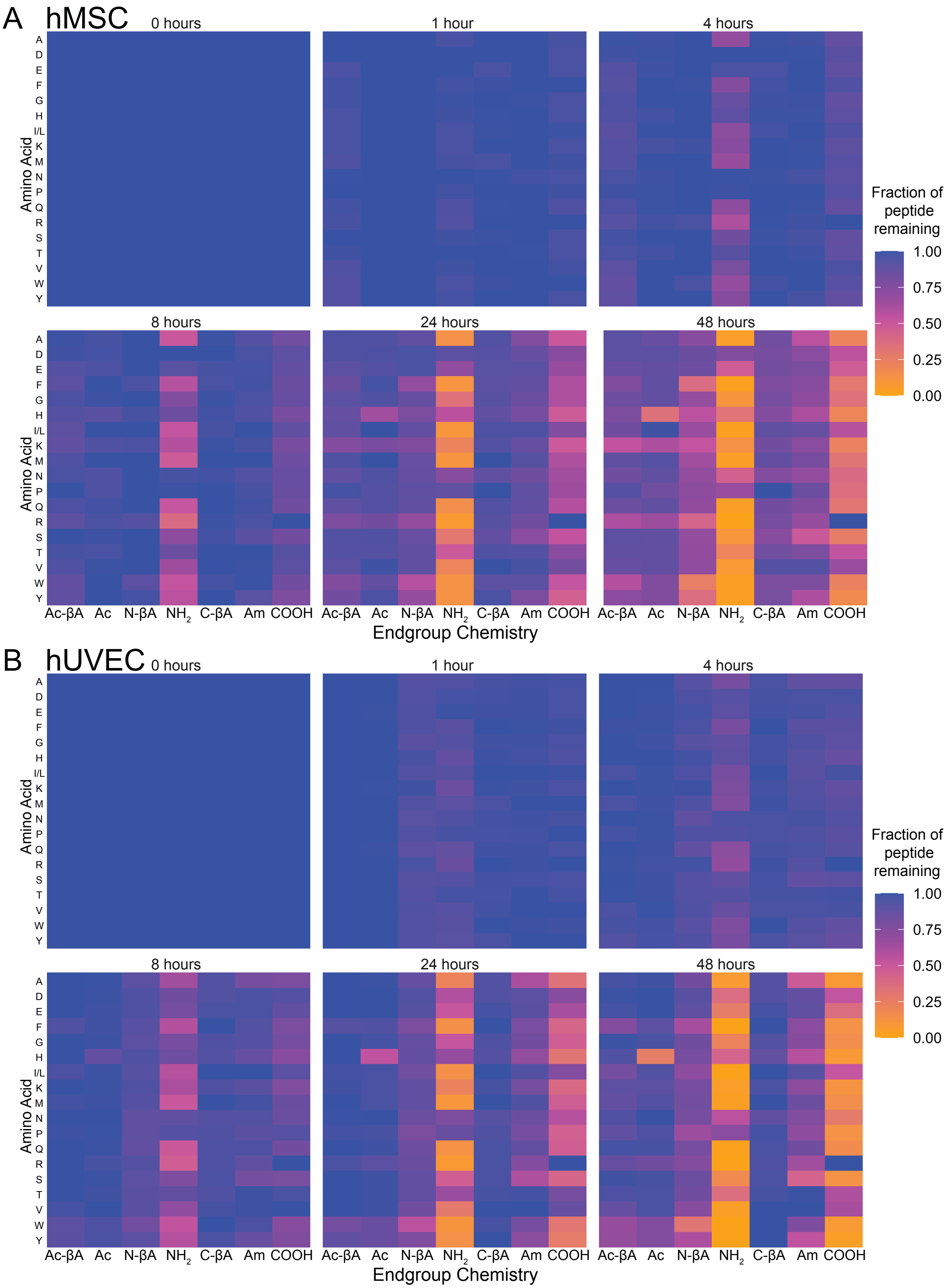
A
hMSC
0 hours
1 hour
4 hours
8 hours
24 hours
48 hours
Amino Acid
A
D
E
F
G
H
I/L
K
M
N
P
Q
R
S
T
V
W
Y
Ac-βA
Ac
N-βA
NH2
C-βA
Am
COOH
Endgroup Chemistry
Fraction of peptide remaining
1.00
0.75
0.50
0.25
0.00
B
hUVEC
0 hours
1 hour
4 hours
8 hours
24 hours
48 hours
Amino Acid
Endgroup Chemistry
Fraction of peptide remaining
1.00
0.75
0.50
0.25
0.00

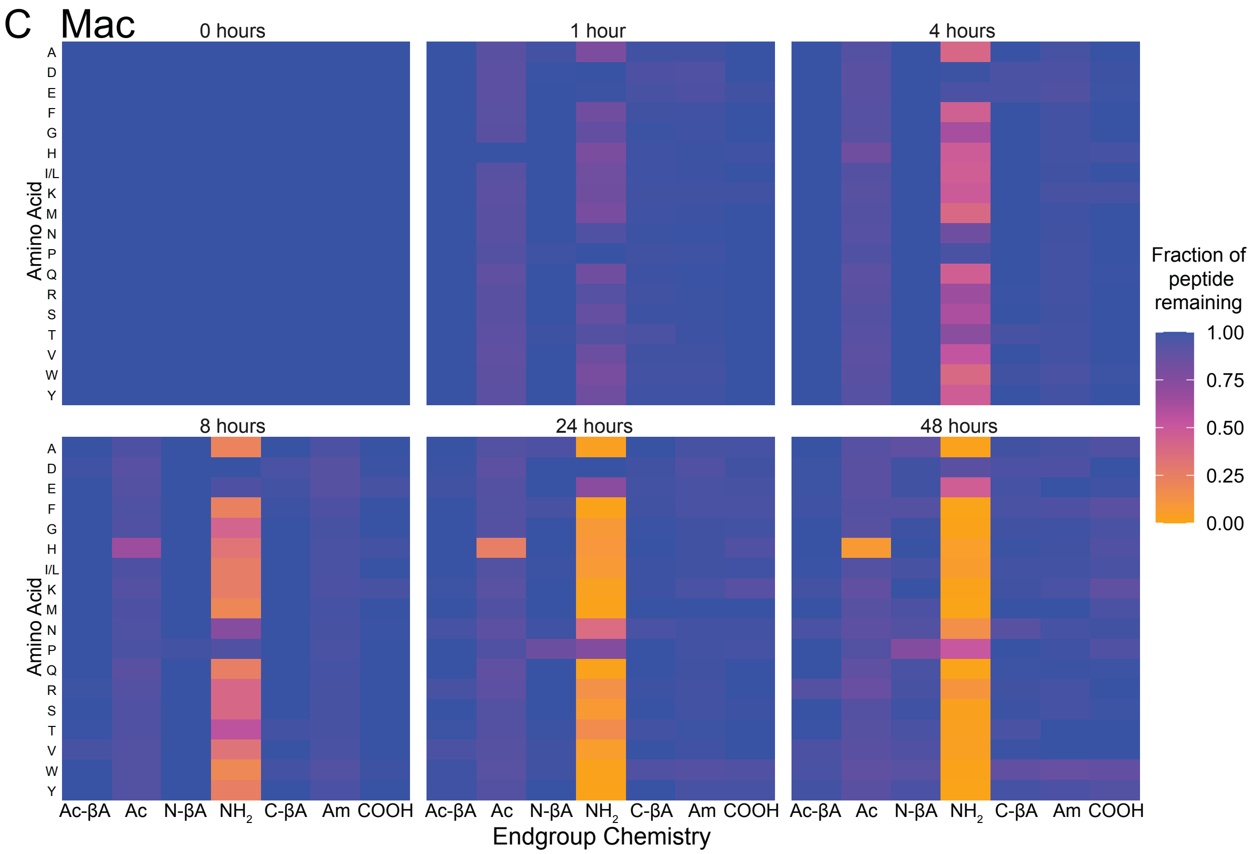
C Mac
0 hours
1 hour
4 hours
8 hours
24 hours
48 hours
Amino Acid
A
D
E
F
G
H
I/L
K
M
N
P
Q
R
S
T
V
W
Y
Fraction of peptide remaining
1.00
0.75
0.50
0.25
0.00
Ac-βA Ac N-βA NH2 C-βA Am COOH
Endgroup Chemistry

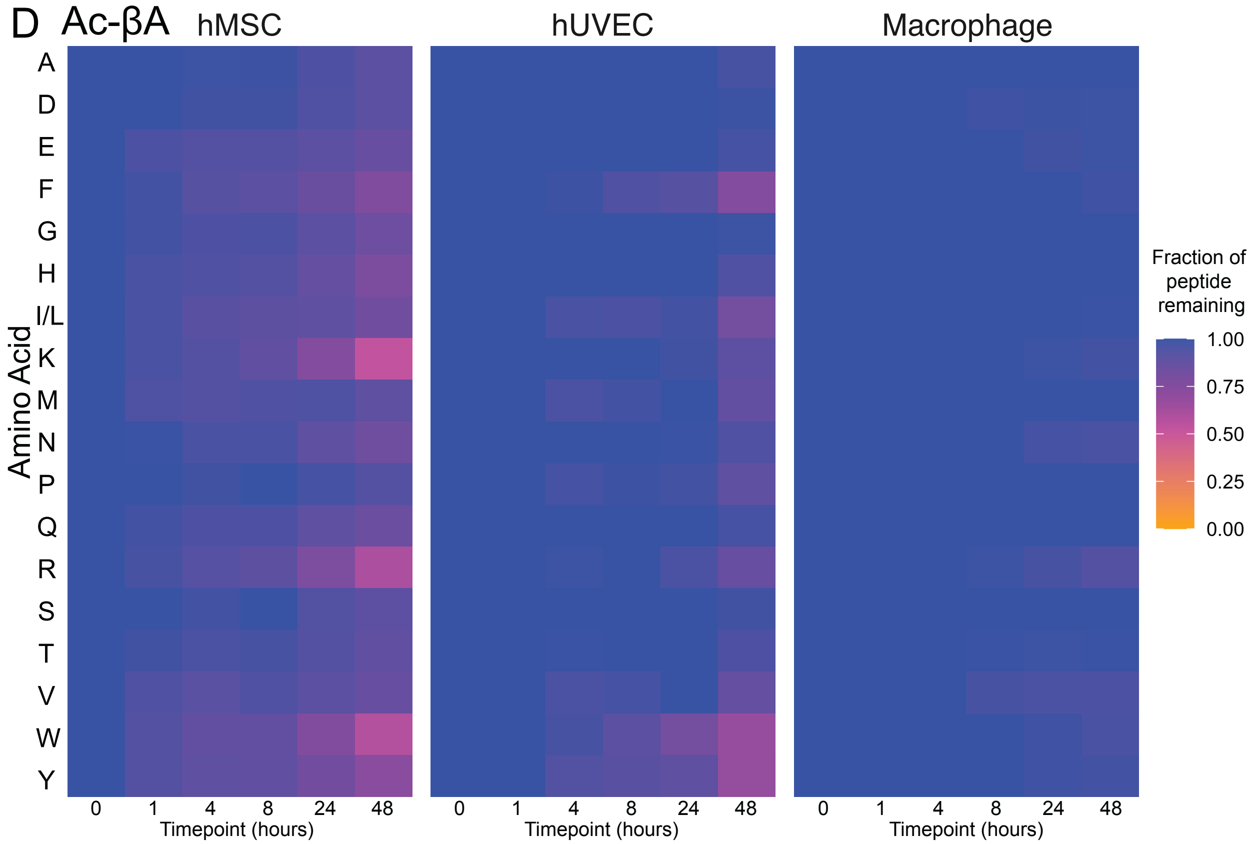
D Ac-βA
hMSC
hUVEC
Macrophage
Amino Acid
A
D
E
F
G
H
I/L
K
M
N
P
Q
R
S
T
V
W
Y
Fraction of peptide remaining
1.00
0.75
0.50
0.25
0.00
0 1 4 8 24 48
Timepoint (hours)

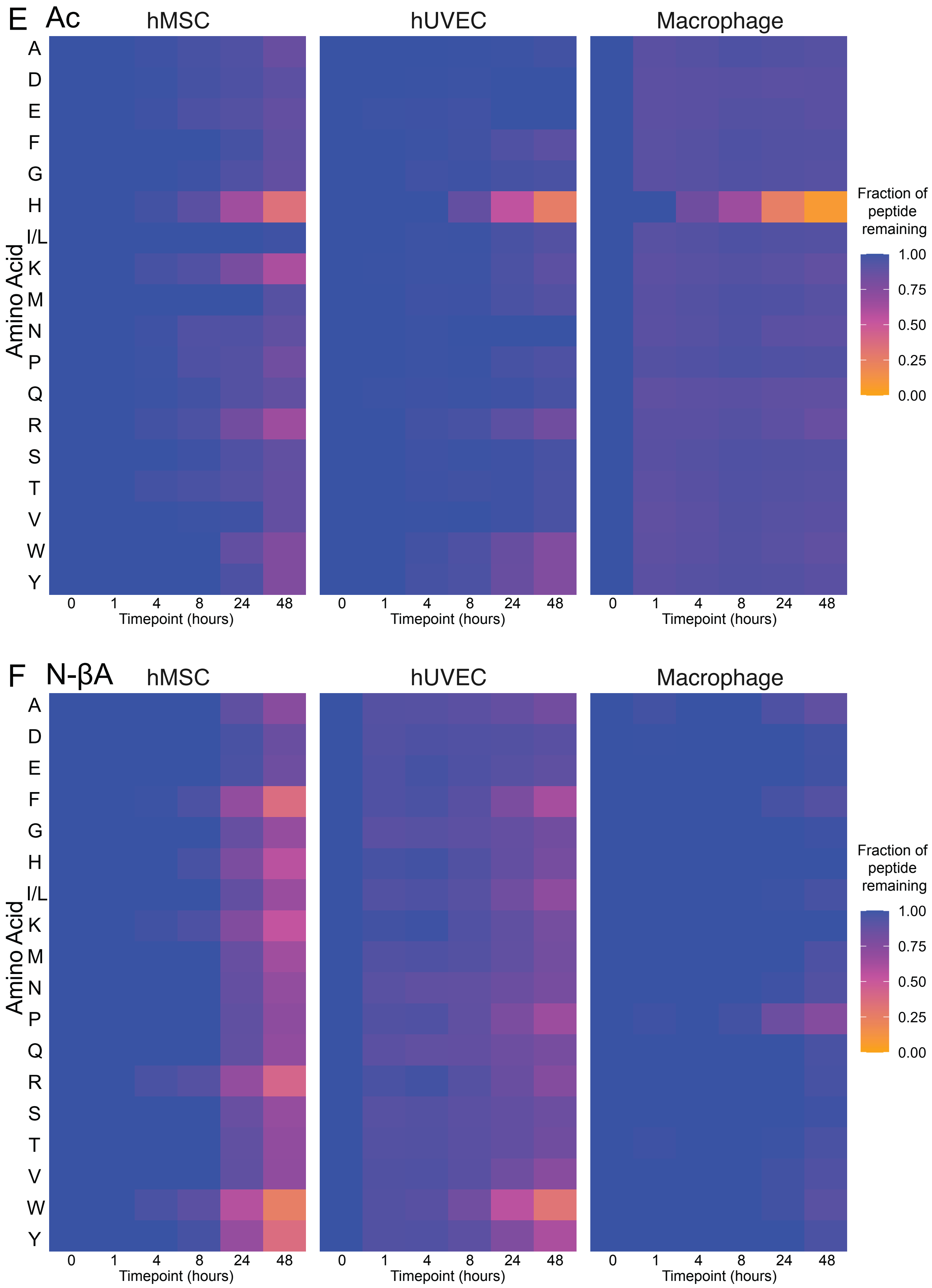
E
Ac
hMSC
hUVEC
Macrophage
Amino Acid
A
D
E
F
G
H
I/L
K
M
N
P
Q
R
S
T
V
W
Y
0
1
4
8
24
48
Timepoint (hours)
Fraction of peptide remaining
1.00
0.75
0.50
0.25
0.00
F
N-βA
hMSC
hUVEC
Macrophage
Amino Acid
Timepoint (hours)
Fraction of peptide remaining

G

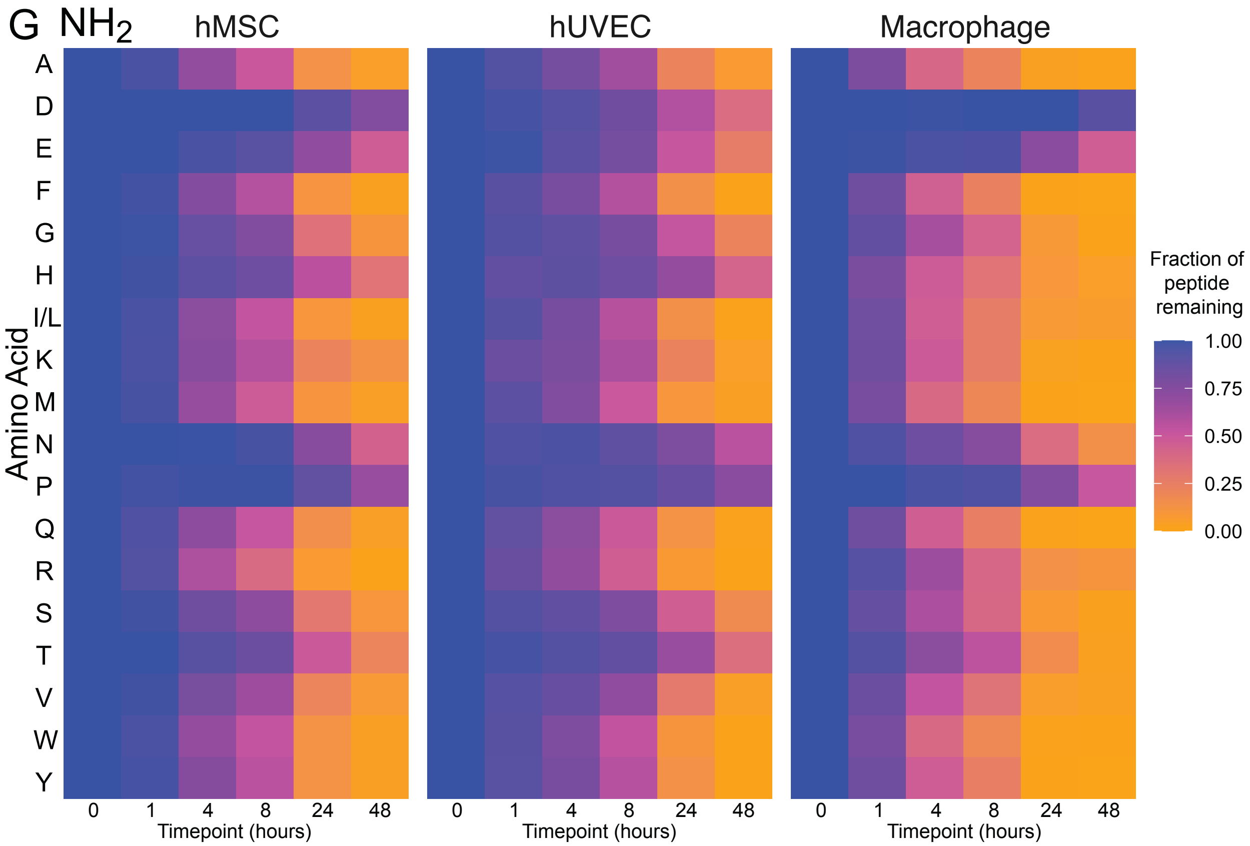
NH2
hMSC
hUVEC
Macrophage
Amino Acid
A
D
E
F
G
H
I/L
K
M
N
P
Q
R
S
T
V
W
Y
0 1 4 8 24 48
Timepoint (hours)
Fraction of peptide remaining
1.00
0.75
0.50
0.25
0.00

H

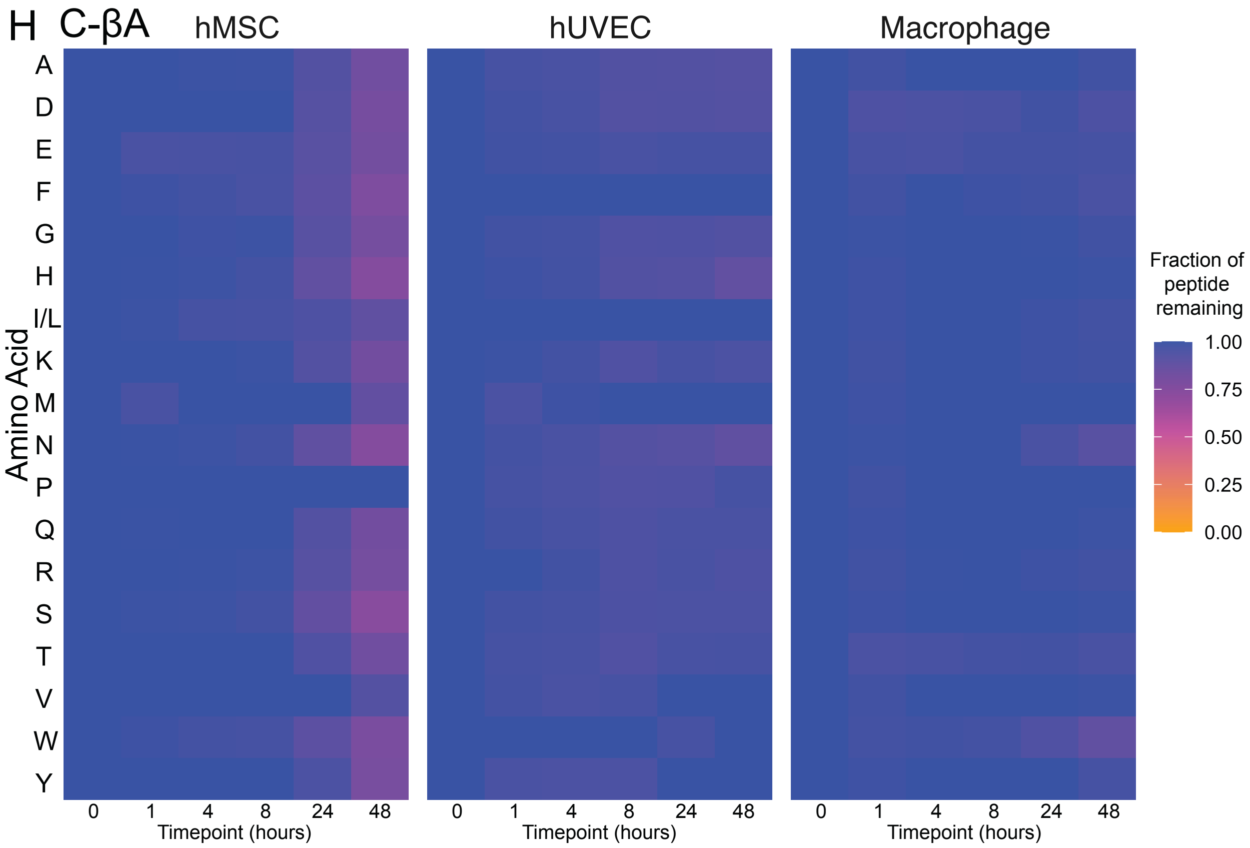
C-βA
hMSC
hUVEC
Macrophage
Amino Acid
A
D
E
F
G
H
I/L
K
M
N
P
Q
R
S
T
V
W
Y
0 1 4 8 24 48
Timepoint (hours)
Fraction of peptide remaining
1.00
0.75
0.50
0.25
0.00

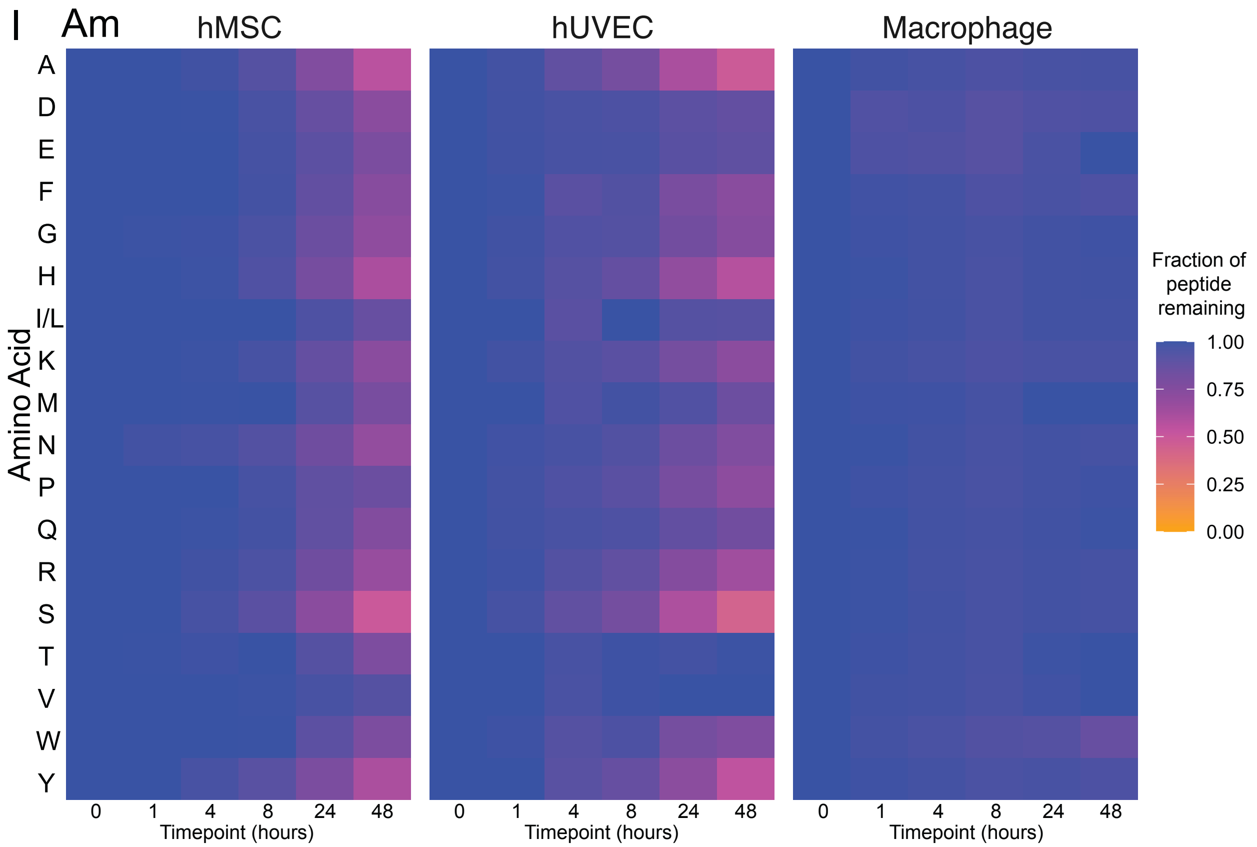
I
Am
hMSC
hUVEC
Macrophage
A
D
E
F
G
H
I/L
K
M
N
P
Q
R
S
T
V
W
Y
Amino Acid
0
1
4
8
24
48
Timepoint (hours)
Fraction of peptide remaining
1.00
0.75
0.50
0.25
0.00

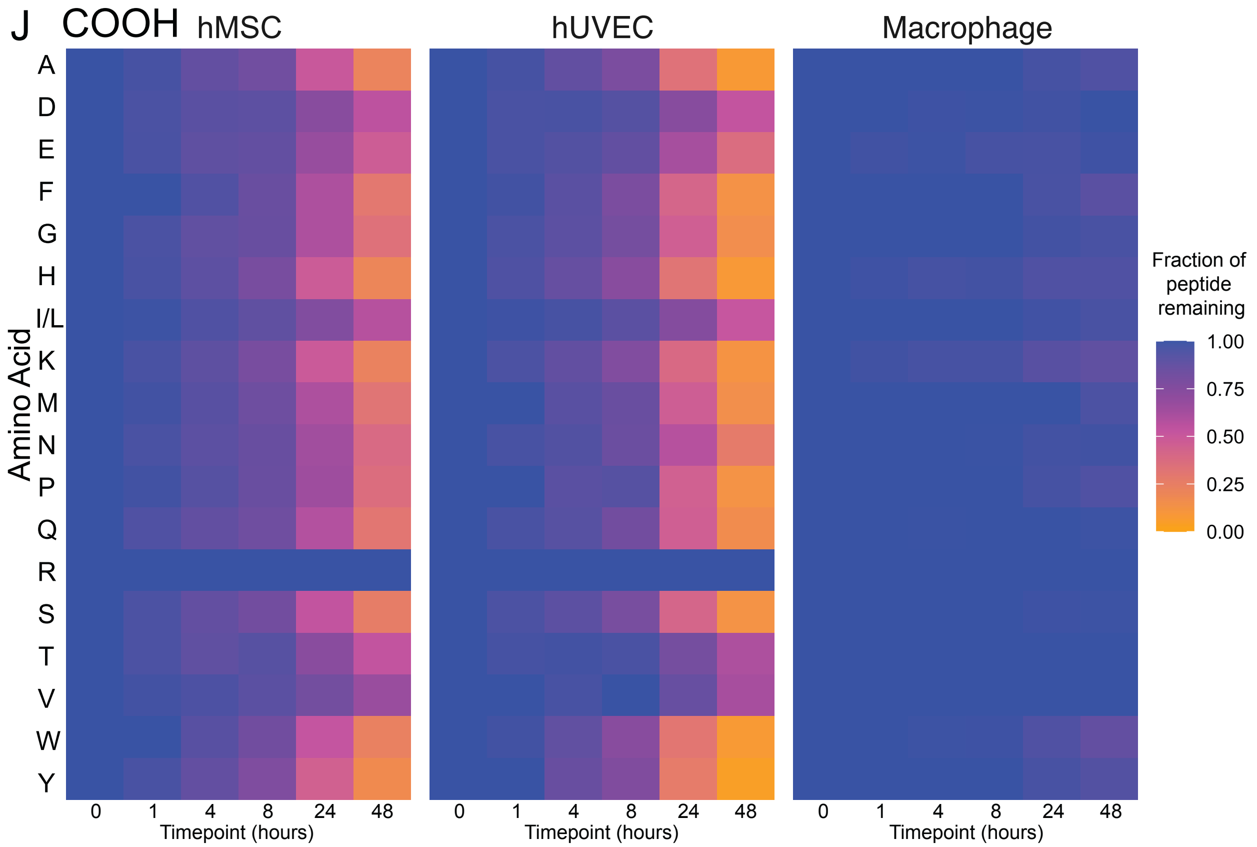
J
COOH
hMSC
hUVEC
Macrophage
A
D
E
F
G
H
I/L
K
M
N
P
Q
R
S
T
V
W
Y
Amino Acid
0
1
4
8
24
48
Timepoint (hours)
Fraction of peptide remaining
1.00
0.75
0.50
0.25
0.00

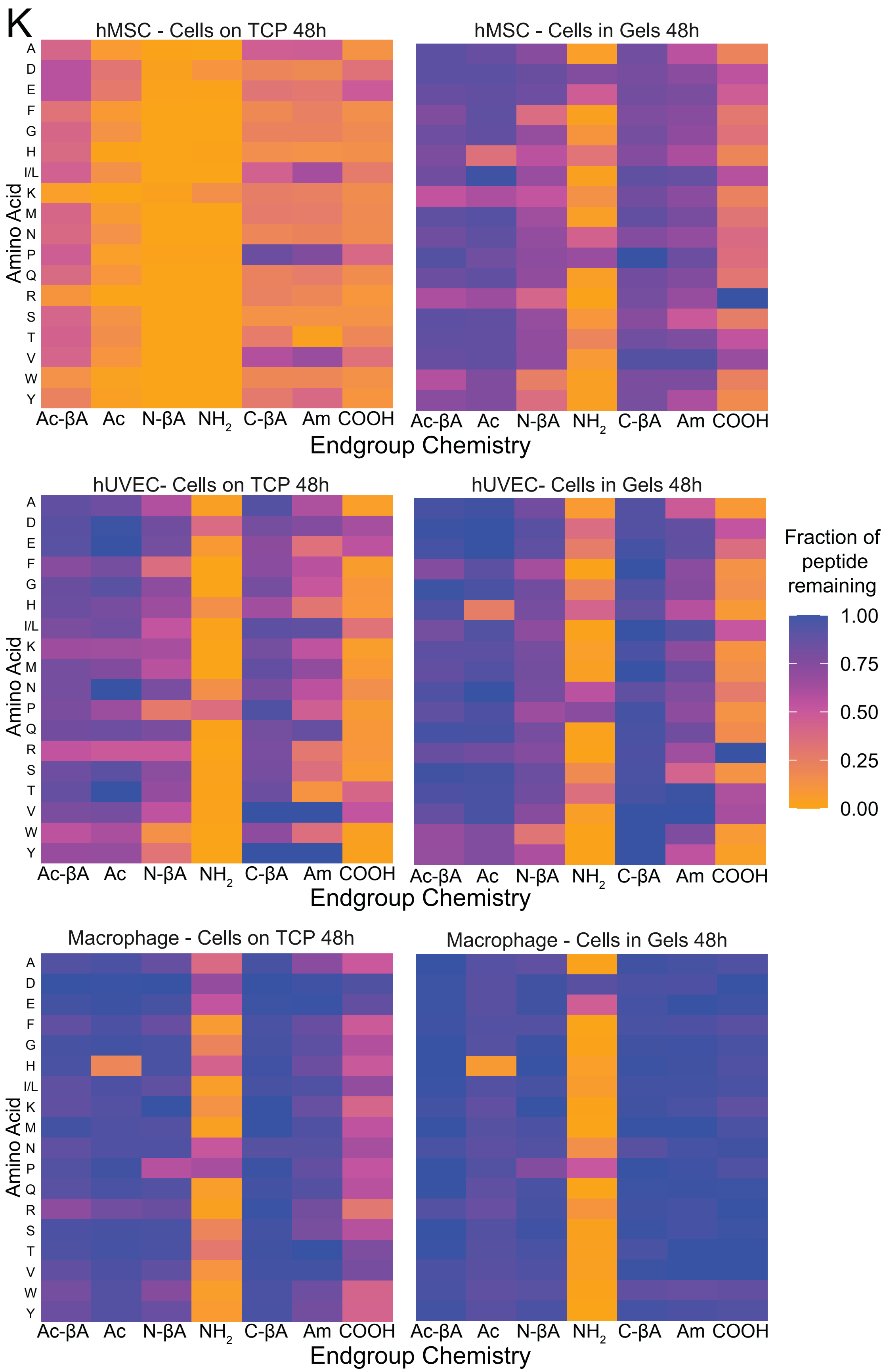
K
hMSC - Cells on TCP 48h
hMSC - Cells in Gels 48h
hUVEC- Cells on TCP 48h
hUVEC- Cells in Gels 48h
Macrophage - Cells on TCP 48h
Macrophage - Cells in Gels 48h
Amino Acid
A D E F G H I/L K M N P Q R S T V W Y
Ac-βA Ac N-βA NH2 C-βA Am COOH
Endgroup Chemistry
Fraction of peptide remaining
1.00
0.75
0.50
0.25
0.00

**Figure S7**. Degradation of soluble peptide libraries by cells in PEG hydrogels. (**A**) hMSCs, (**B**) hUVECs, (**C**), macrophages, and by peptide endgroup (**D**) Ac-βA, (**E**) Ac, (**F**) N-βA, (**G**) $NH_2$, (**H**) C-βA, I) Am, (**J**) COOH. (**K**) Direct comparison of peptide degradation by cells on tissue culture plastic (TCP) and cells in gels.

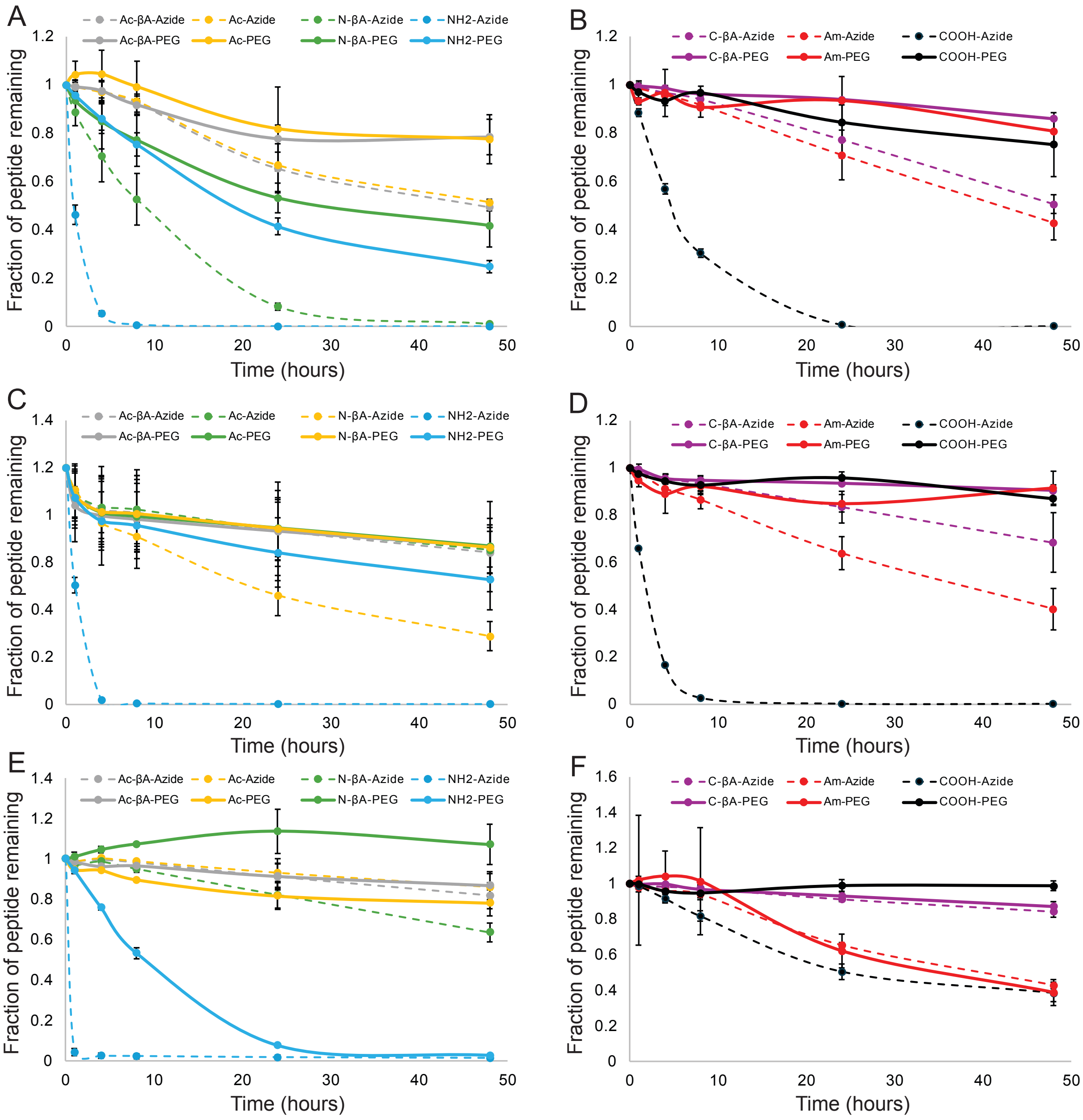


**Figure S8**. PEG conjugation to peptides slows degradation across endgroups and cell types. (**A**) hMSC N-terminal chemistries, (**B**) hMSC C-terminal chemistries, (**C**) hUVEC N-terminal chemistries, (**D**) hUVEC C-terminal chemistries, (**E**) macrophage N-terminal chemistries, (**F**) macrophage C-terminal chemistries.

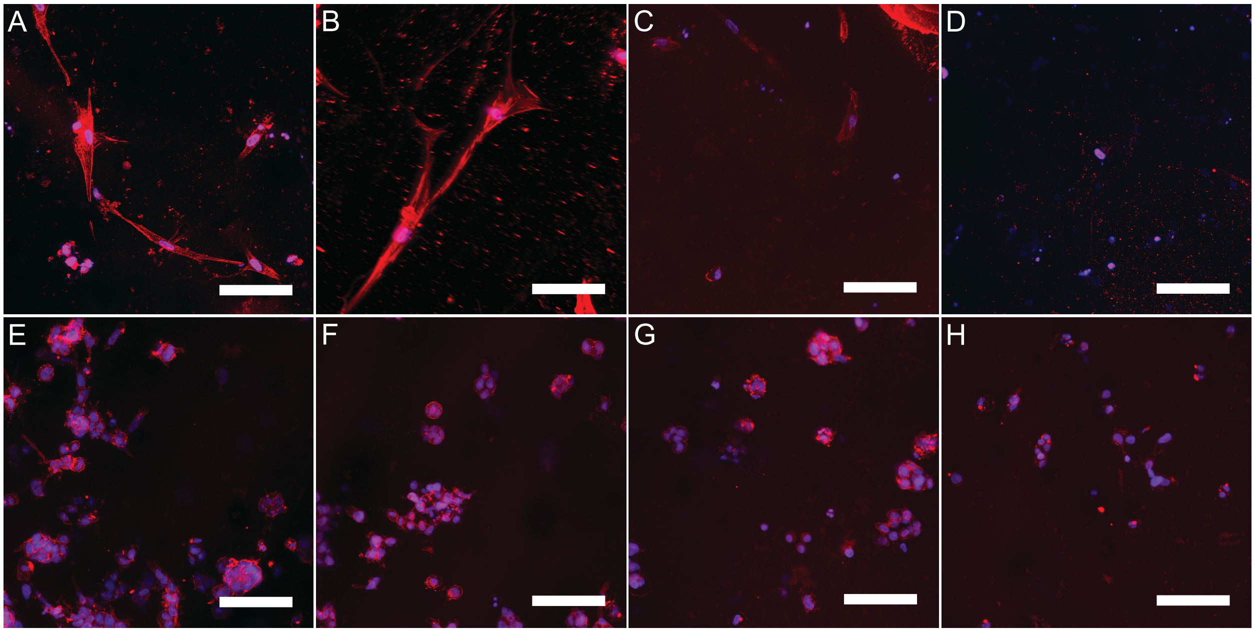


**Figure S9**. Cells growing in gels with different RGD presentations. hUVECs growing in gels with (**A**) cyclic RGDS, (**B**) Ac-βA-GRGDS, (**C**) $NH_2$-GRGDS, and (**D**) no added RGDS. Macrophages growing in gels with (**E**) cyclic RGDS, (**F**) Ac-βA-GRGDS, (**G**) $NH_2$-GRGDS, and (**H**) no added RGDS. It should be noted that the viability assay (Fig. S9) indicates negligible of metabolic activity within hUVEC gels lacking RGD sequences. Scale bar is 100 μm, and red is actin and blue is the cell nuclei.

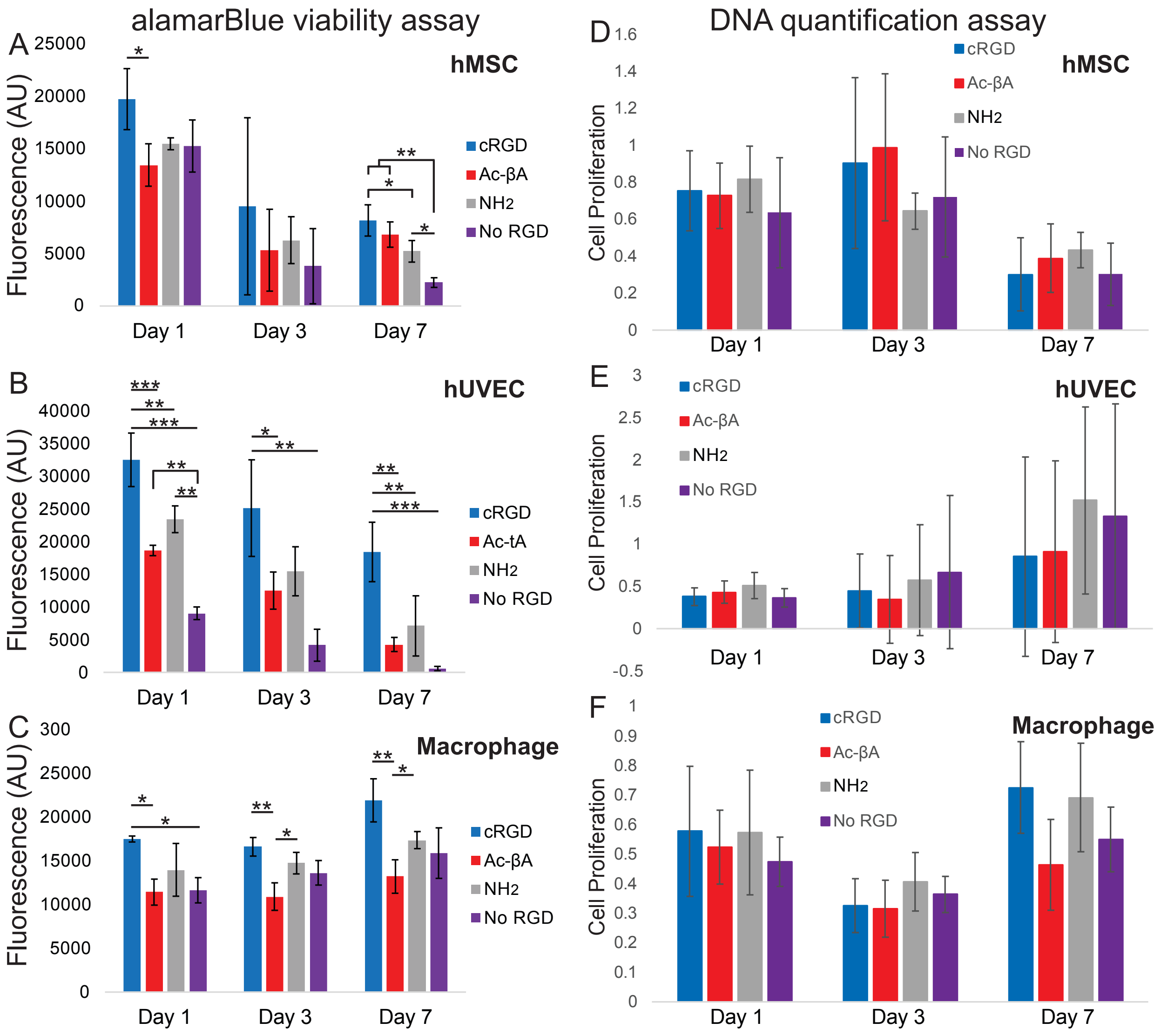


**Figure S10**. Quantification of viability and proliferation in hydrogels containing different RGD sequences. An alamarBlue metabolic activity assays was performed on (**A**) hMSCs, (**B**) hUVECs, and (**C**) macrophages and a DNA quantification assay was performed on (**D**) hMSCs, (**E**) hUVECs, and (**F**) macrophages. * indicates $p < 0.05$, ** indicates $p < 0.01$, *** indicates $p < 0.0001$ by Tukey's post hoc test.

A

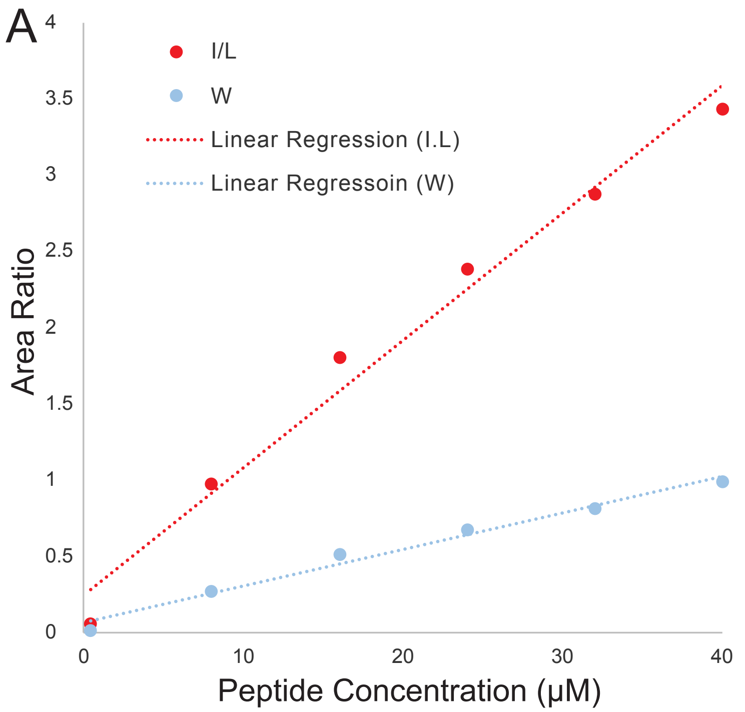


B

| | K | H | R | N | S | Q | G | A | T | E |
|---|---|---|---|---|---|---|---|---|---|---|
| Ac-βA | 0.977 | 0.965 | 0.978 | 0.979 | 0.978 | 0.986 | 0.978 | 0.977 | 0.977 | 0.983 |
| Ac | 0.936 | 0.931 | 0.938 | 0.791 | 0.977 | 0.936 | 0.986 | 0.976 | 0.979 | 0.979 |
| N-βA | 0.962 | 0.943 | 0.976 | 0.995 | 0.998 | 0.992 | 0.997 | 0.998 | 0.997 | 0.996 |
| NH2 | 0.984 | 0.983 | 0.969 | 0.981 | 0.981 | 0.980 | 0.984 | 0.981 | 0.982 | 0.993 |
| C-βA | 0.994 | 0.989 | 0.982 | 0.992 | 0.992 | 0.991 | 0.989 | 0.994 | 0.993 | 0.995 |
| Am | 0.998 | 0.995 | 0.996 | 0.999 | 0.994 | 0.996 | 0.999 | 0.990 | 0.998 | 0.997 |
| COOH | 0.871 | 0.903 | | 0.934 | 0.900 | 0.914 | 0.939 | 0.896 | 0.974 | 0.979 |

| | D | P | V | Y | M | I/L | F | W | Average | St Dev |
|---|---|---|---|---|---|---|---|---|---|---|
| Ac-βA | 0.983 | 0.979 | 0.977 | 0.973 | 0.979 | 0.981 | 0.979 | 0.984 | 0.978 | 0.004 |
| Ac | 0.738 | 0.898 | 0.870 | 0.808 | 0.858 | 0.891 | 0.841 | 0.855 | 0.899 | 0.073 |
| N-βA | 0.998 | 0.997 | 0.996 | 0.989 | 0.992 | 0.995 | 0.992 | 0.982 | 0.989 | 0.015 |
| NH2 | 0.983 | 0.993 | 0.985 | 0.978 | 0.979 | 0.985 | 0.984 | 0.983 | 0.983 | 0.005 |
| C-βA | 0.992 | 0.992 | 0.983 | 0.972 | 0.991 | 0.987 | 0.991 | 0.988 | 0.989 | 0.006 |
| Am | 0.999 | 0.999 | 0.999 | 0.997 | 0.997 | 0.998 | 0.998 | 0.998 | 0.997 | 0.002 |
| COOH | 0.988 | 0.928 | 0.990 | 0.918 | 0.935 | 0.982 | 0.944 | 0.953 | 0.938 | 0.036 |

**Figure S11**. Standard curves validating the use of LCMS to measure the concentration of peptides. Peptide libraries were run on the LCMS at different concentrations ranging from 0.4-40 µM and the amount of peptide was quantified. (**A**) Two representative conditions, I/L and W peptides in Ac-βA and the linear regression line fitting the data points. (**B**) A table of the R-squared values for every endgroup/amino acid combinations.

**A**

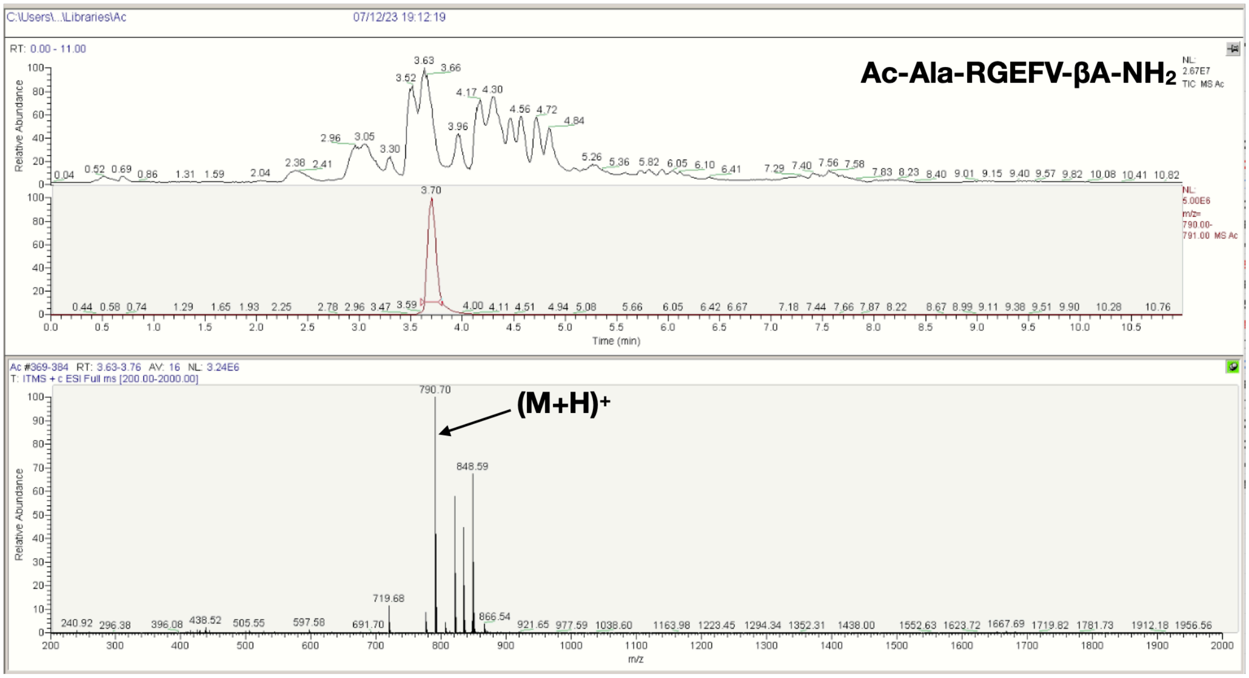
C:\Users\...\Libraries\Ac
07/12/23 19:12:19
Ac-Ala-RGEFV-βA-NH2
RT: 0.00 - 11.00
Relative Abundance
Time (min)
Ac #369-384 RT: 3.63-3.76 AV: 16 NL: 3.24E6
T: ITMS + c ESI Full ms [200.00-2000.00]
(M+H)+
790.70
848.59
m/z

**B**

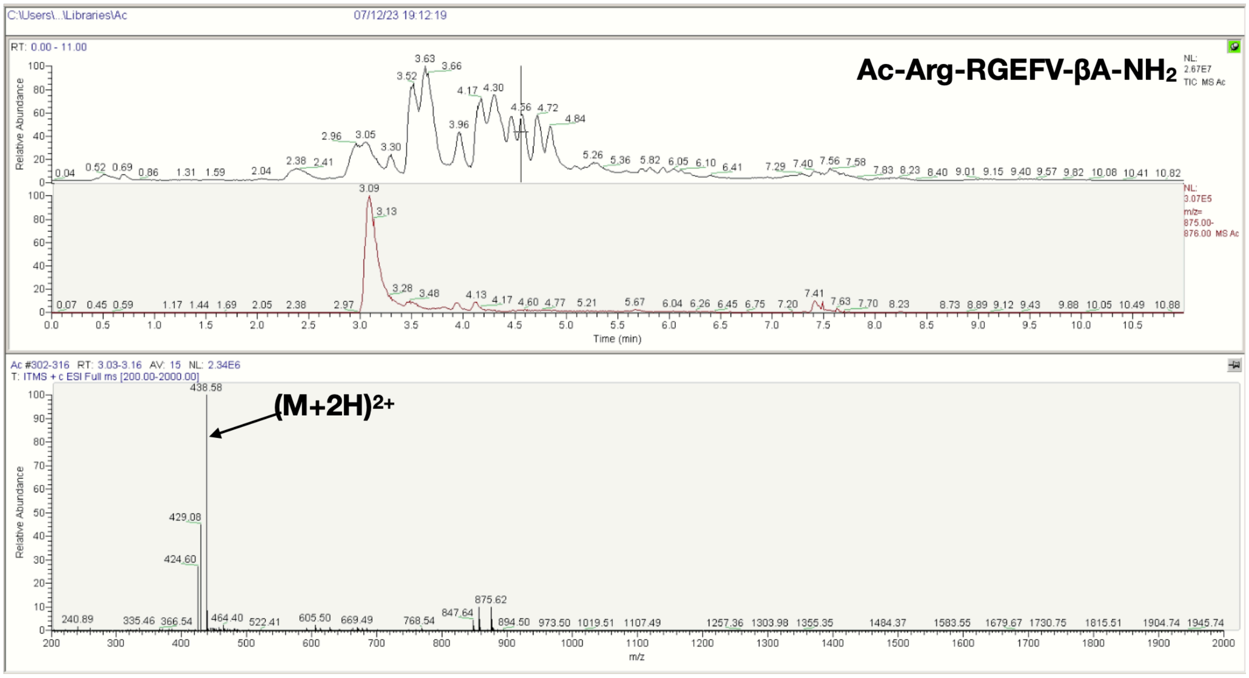
C:\Users\...\Libraries\Ac
07/12/23 19:12:19
Ac-Arg-RGEFV-βA-NH2
RT: 0.00 - 11.00
Relative Abundance
Time (min)
Ac #302-316 RT: 3.03-3.16 AV: 15 NL: 2.34E6
T: ITMS + c ESI Full ms [200.00-2000.00]
(M+2H)2+
438.58
429.08
424.60
875.62
m/z

C

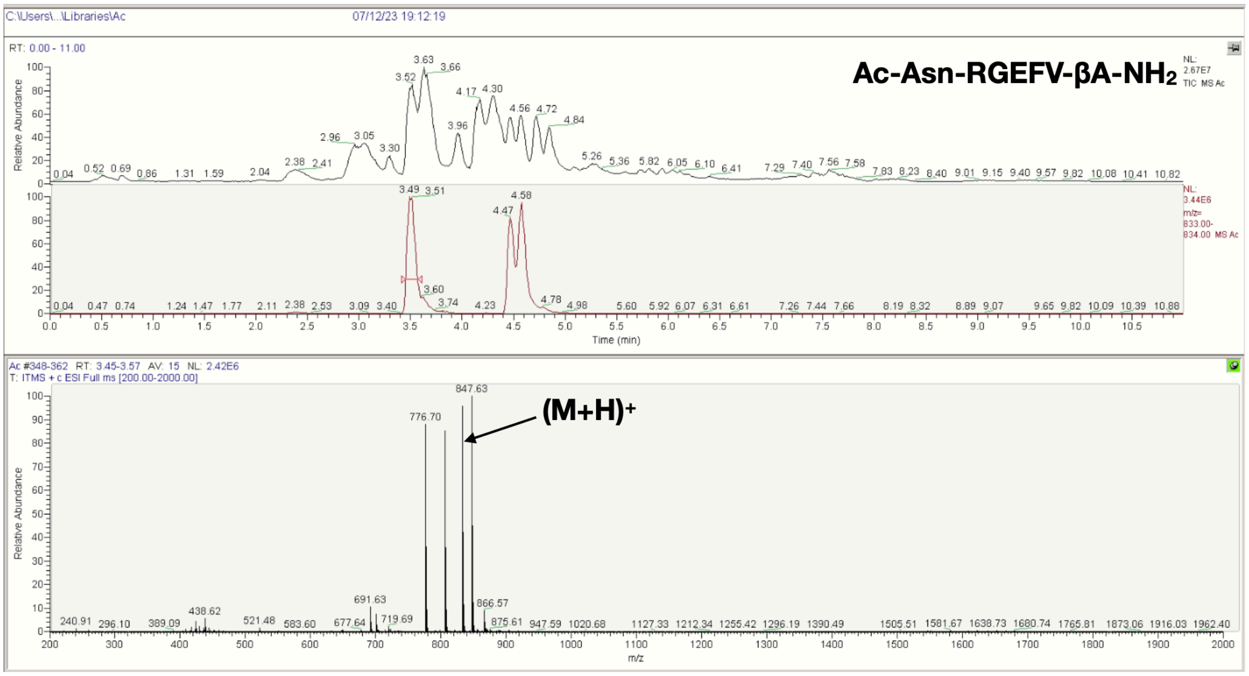
C:\Users\...\Libraries\Ac
07/12/23 19:12:19
Ac-Asn-RGEFV-βA-NH2
(M+H)+
Relative Abundance
Time (min)
m/z
Ac #348-362 RT: 3.45-3.57 AV: 15 NL: 2.42E6
T: ITMS + c ESI Full ms [200.00-2000.00]
847.63
776.70

D

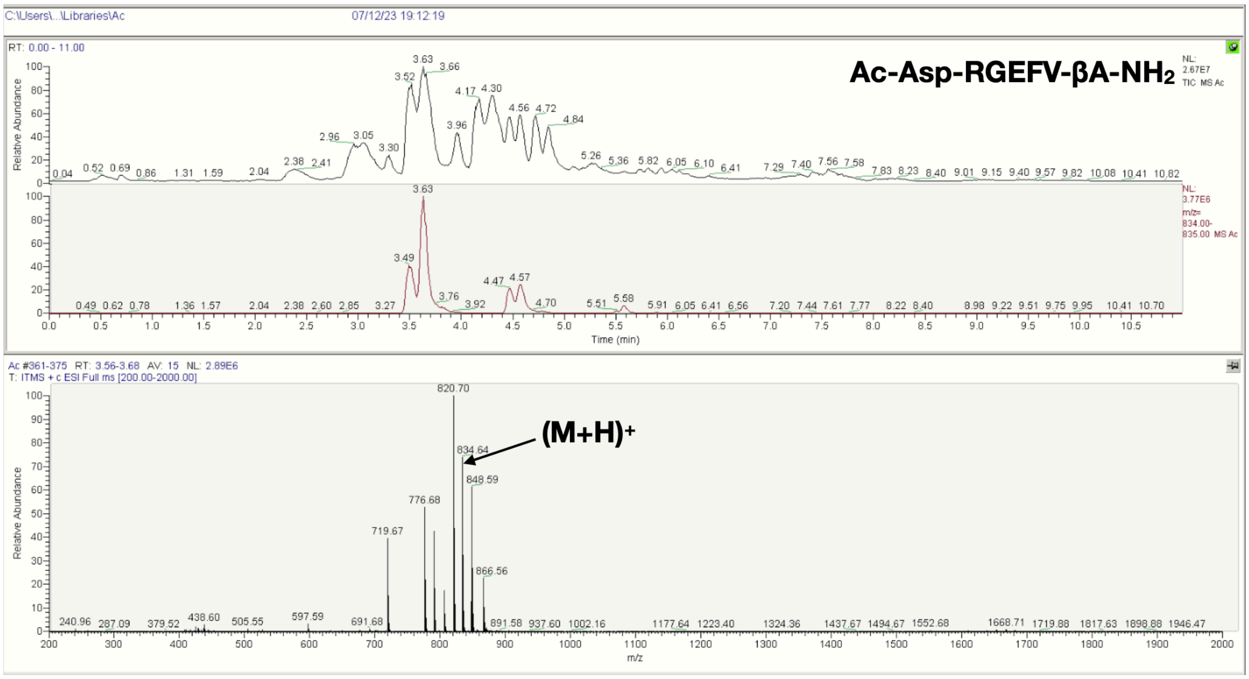
C:\Users\...\Libraries\Ac
07/12/23 19:12:19
Ac-Asp-RGEFV-βA-NH2
(M+H)+
Relative Abundance
Time (min)
m/z
Ac #361-375 RT: 3.56-3.68 AV: 15 NL: 2.89E6
T: ITMS + c ESI Full ms [200.00-2000.00]
820.70
834.64
848.59
776.68
719.67

E

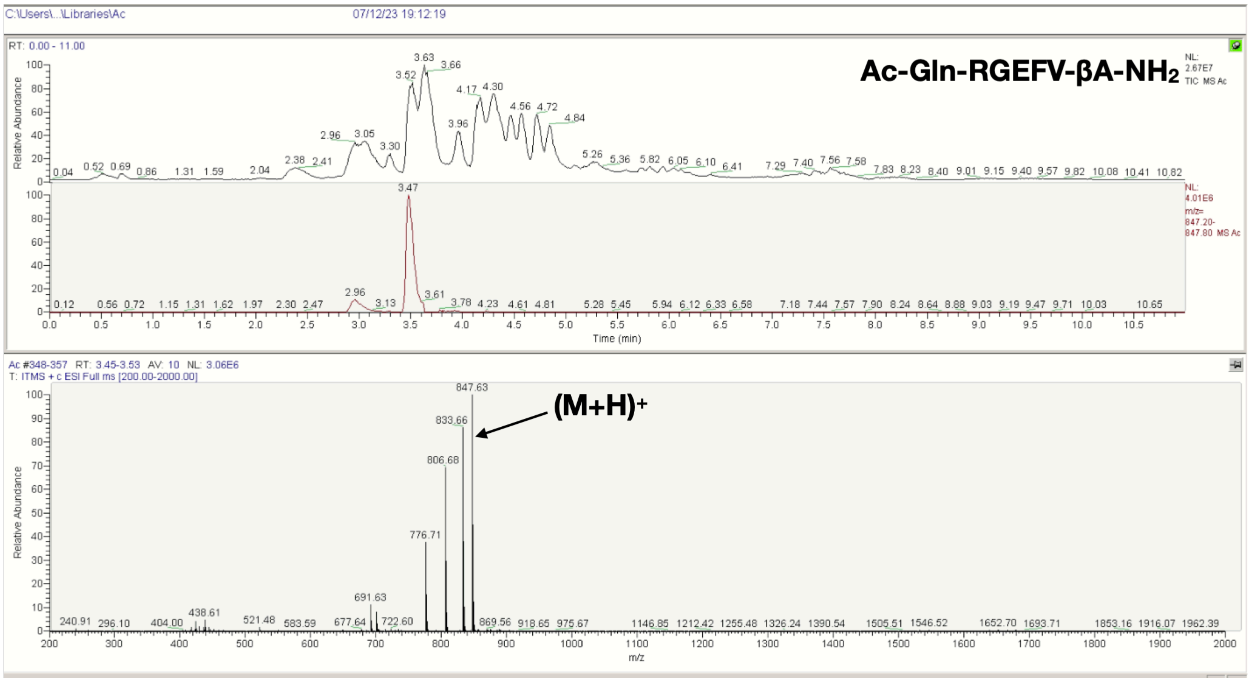


F

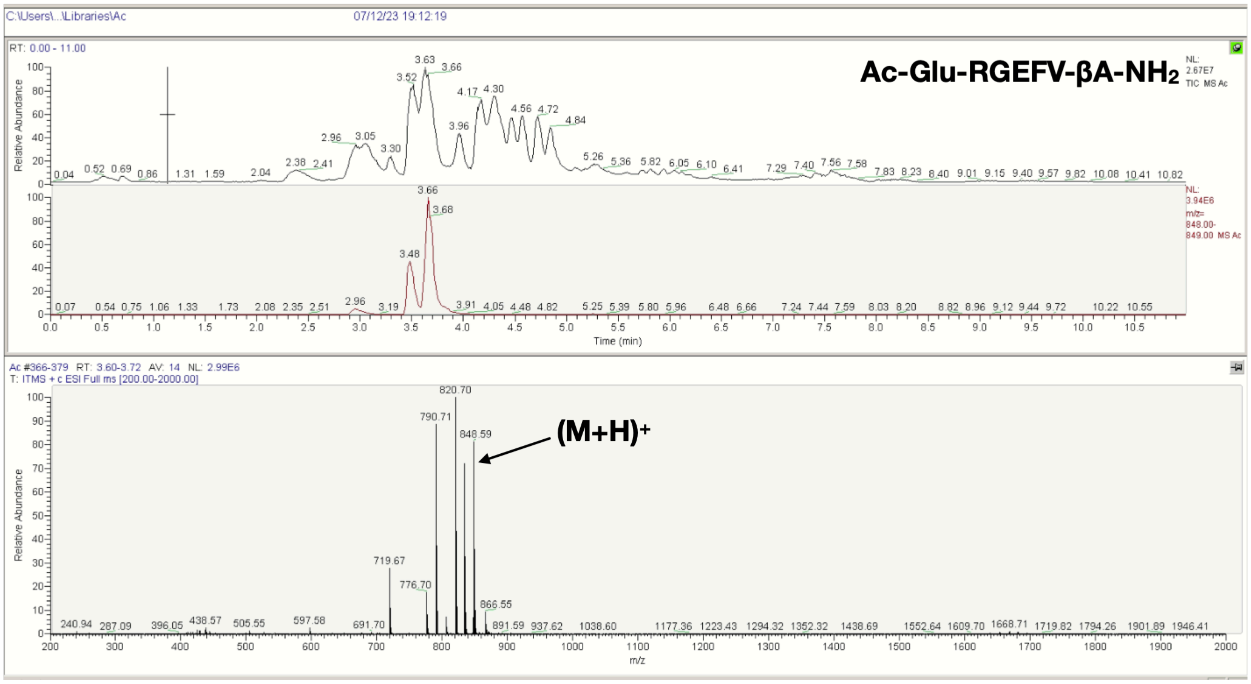

G

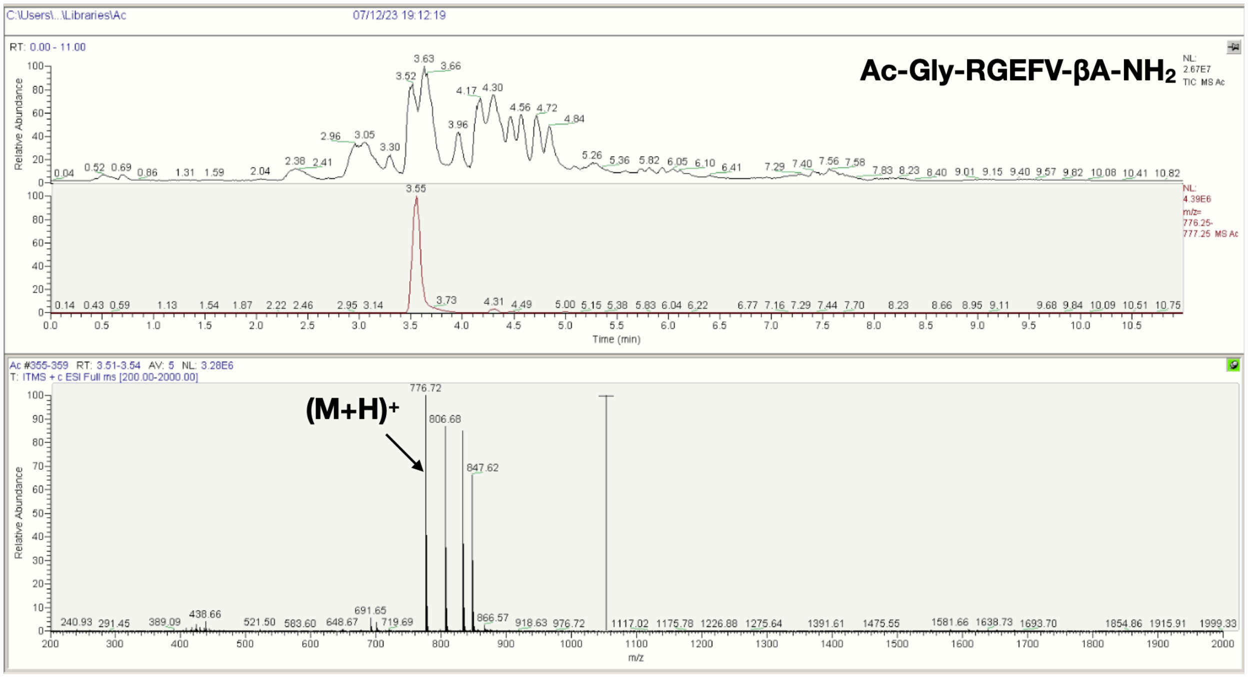
C:\Users\...\Libraries\Ac
07/12/23 19:12:19
RT: 0.00 - 11.00
Ac-Gly-RGEFV-βA-NH2
NL: 2.67E7 TIC MS Ac
NL: 4.39E6 m/z= 776.25-777.25 MS Ac
Relative Abundance
Time (min)
Ac #355-359 RT: 3.51-3.54 AV: 5 NL: 3.28E6
T: ITMS + c ESI Full ms [200.00-2000.00]
(M+H)+
776.72
806.68
847.62
m/z

H

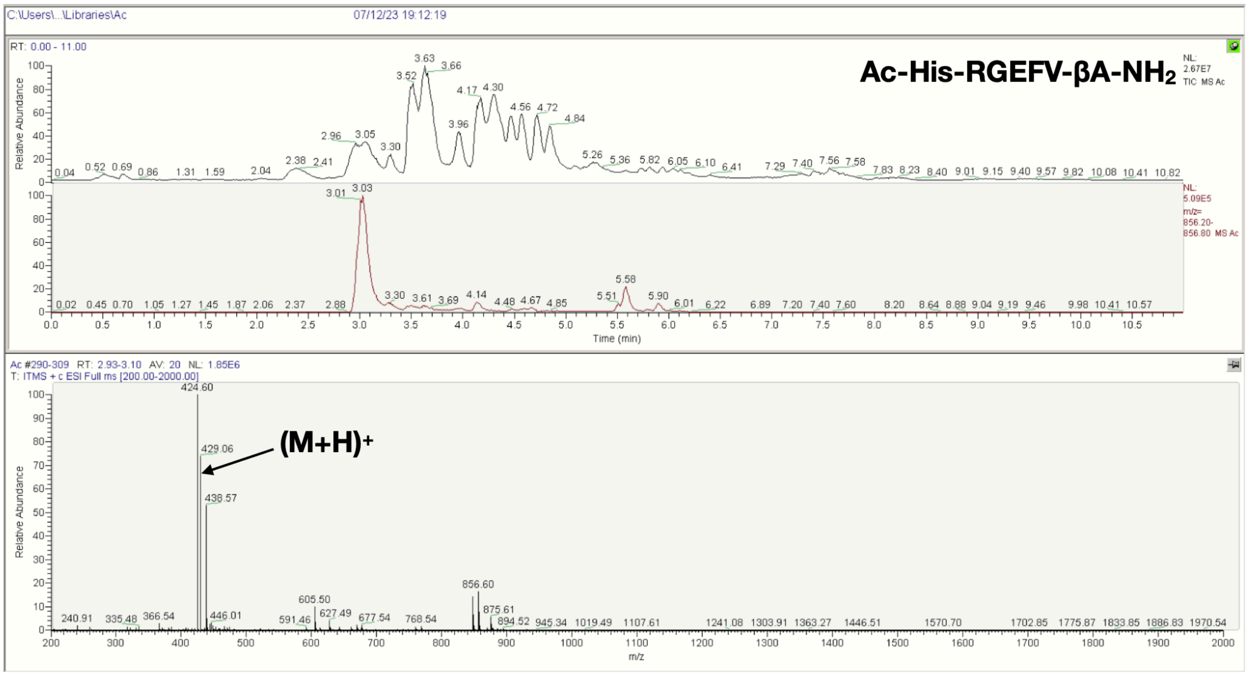
C:\Users\...\Libraries\Ac
07/12/23 19:12:19
RT: 0.00 - 11.00
Ac-His-RGEFV-βA-NH2
NL: 2.67E7 TIC MS Ac
NL: 5.09E5 m/z= 856.20-856.80 MS Ac
Relative Abundance
Time (min)
Ac #290-309 RT: 2.93-3.10 AV: 20 NL: 1.85E6
T: ITMS + c ESI Full ms [200.00-2000.00]
(M+H)+
424.60
429.06
438.57
856.60
m/z

I

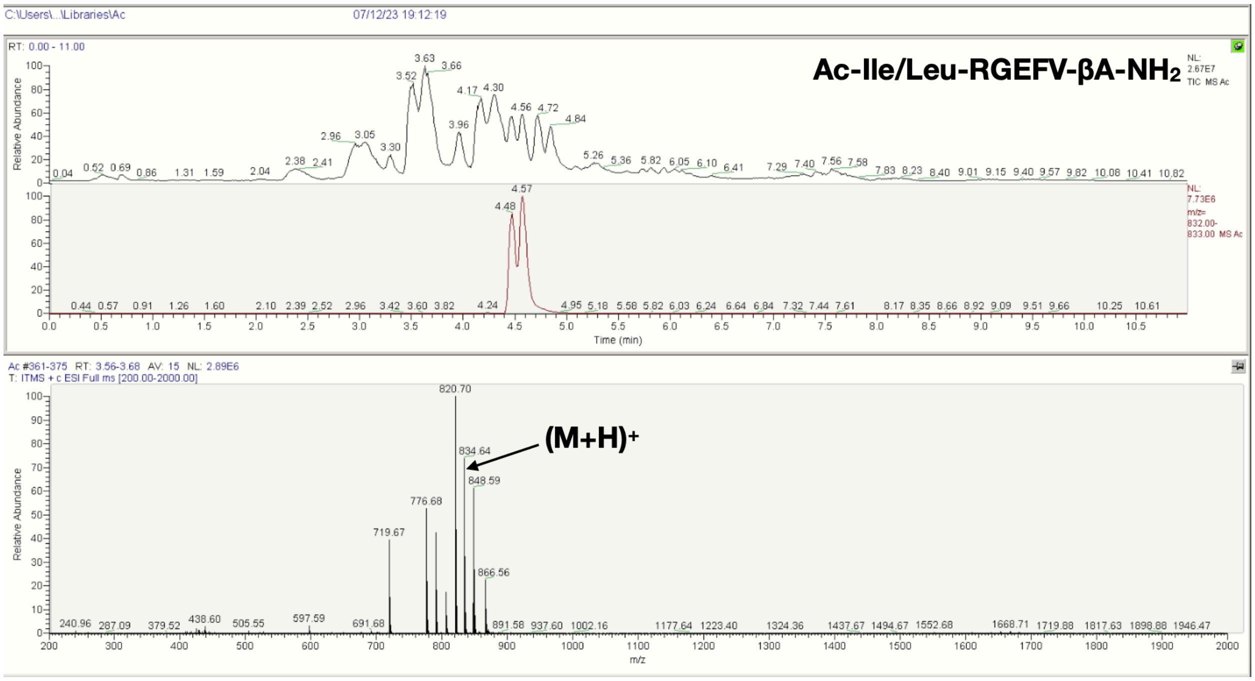
Ac-Ile/Leu-RGEFV-βA-NH2
(M+H)+
820.70
834.64
848.59
776.68
719.67
866.56
Time (min)
m/z
Relative Abundance

J

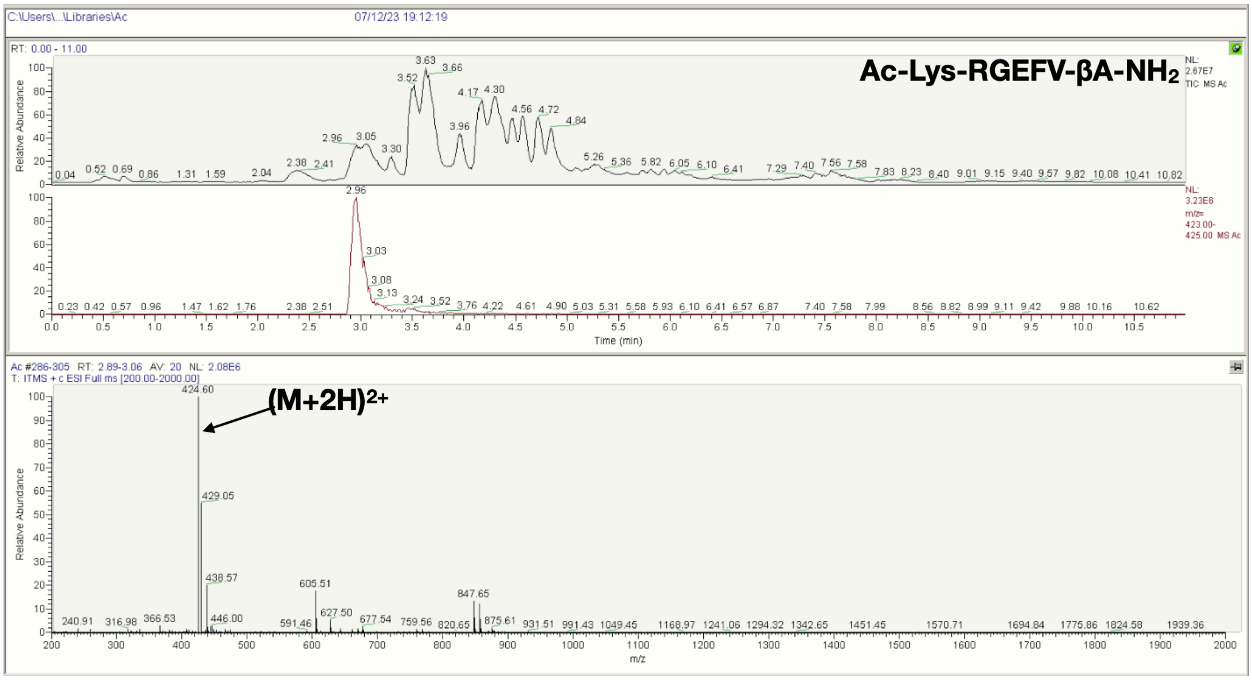
Ac-Lys-RGEFV-βA-NH2
(M+2H)2+
424.60
429.05
438.57
605.51
847.65
Time (min)
m/z
Relative Abundance

**K**

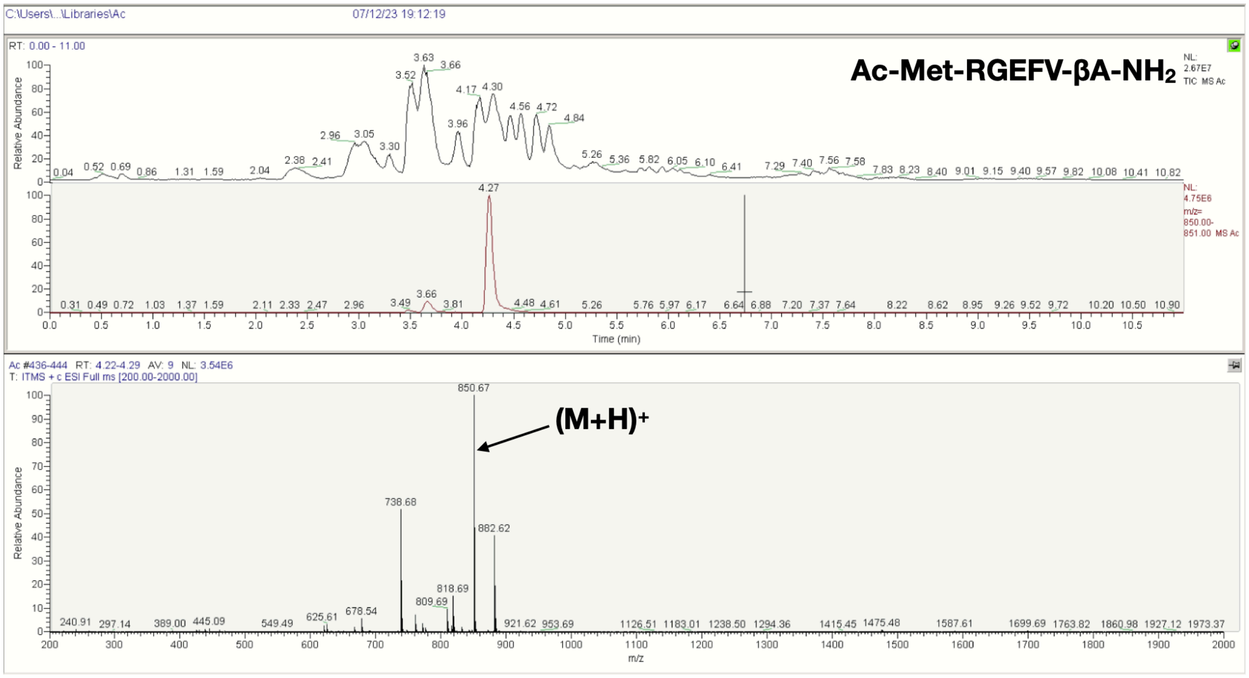
C:\Users\...\Libraries\Ac
07/12/23 19:12:19
RT: 0.00 - 11.00
Ac-Met-RGEFV-βA-NH2
NL: 2.67E7 TIC MS Ac
NL: 4.75E6 m/z= 850.00-851.00 MS Ac
Relative Abundance
Time (min)
Ac #436-444 RT: 4.22-4.29 AV: 9 NL: 3.54E6
T: ITMS + c ESI Full ms [200.00-2000.00]
850.67
(M+H)+
738.68
882.62
818.69
809.69
678.54
m/z

**L**

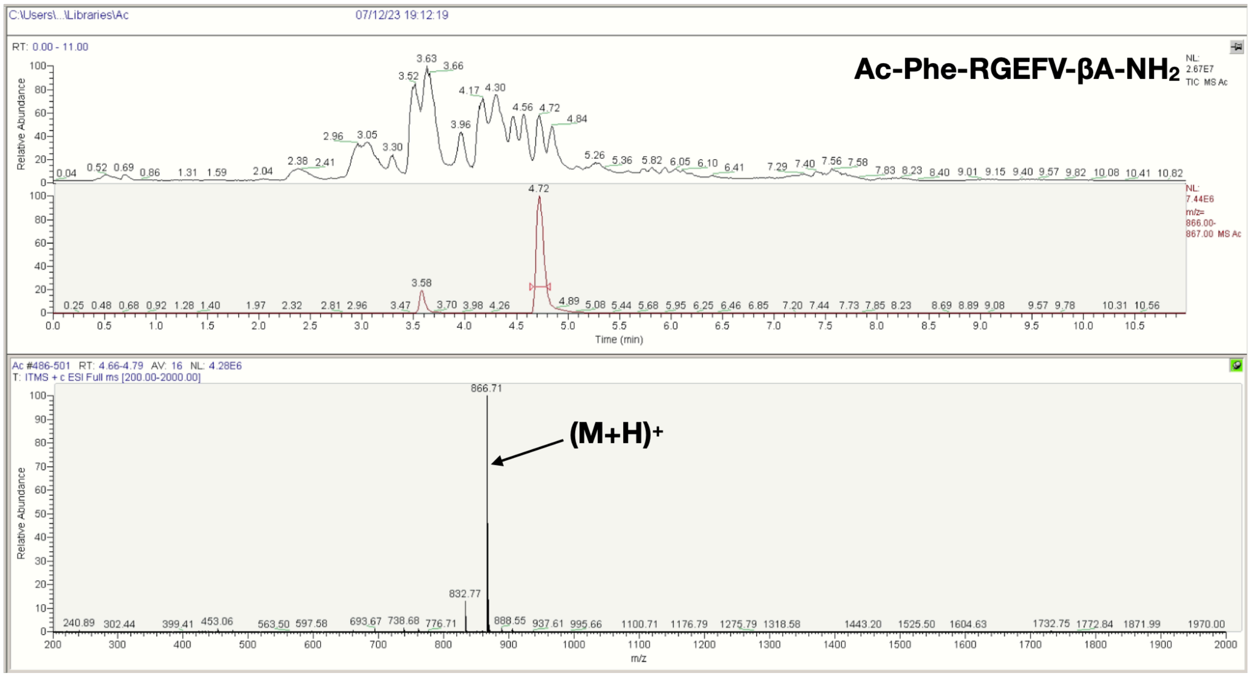
C:\Users\...\Libraries\Ac
07/12/23 19:12:19
RT: 0.00 - 11.00
Ac-Phe-RGEFV-βA-NH2
NL: 2.67E7 TIC MS Ac
NL: 7.44E6 m/z= 866.00-867.00 MS Ac
Relative Abundance
Time (min)
Ac #486-501 RT: 4.66-4.79 AV: 16 NL: 4.28E6
T: ITMS + c ESI Full ms [200.00-2000.00]
866.71
(M+H)+
832.77
m/z

M

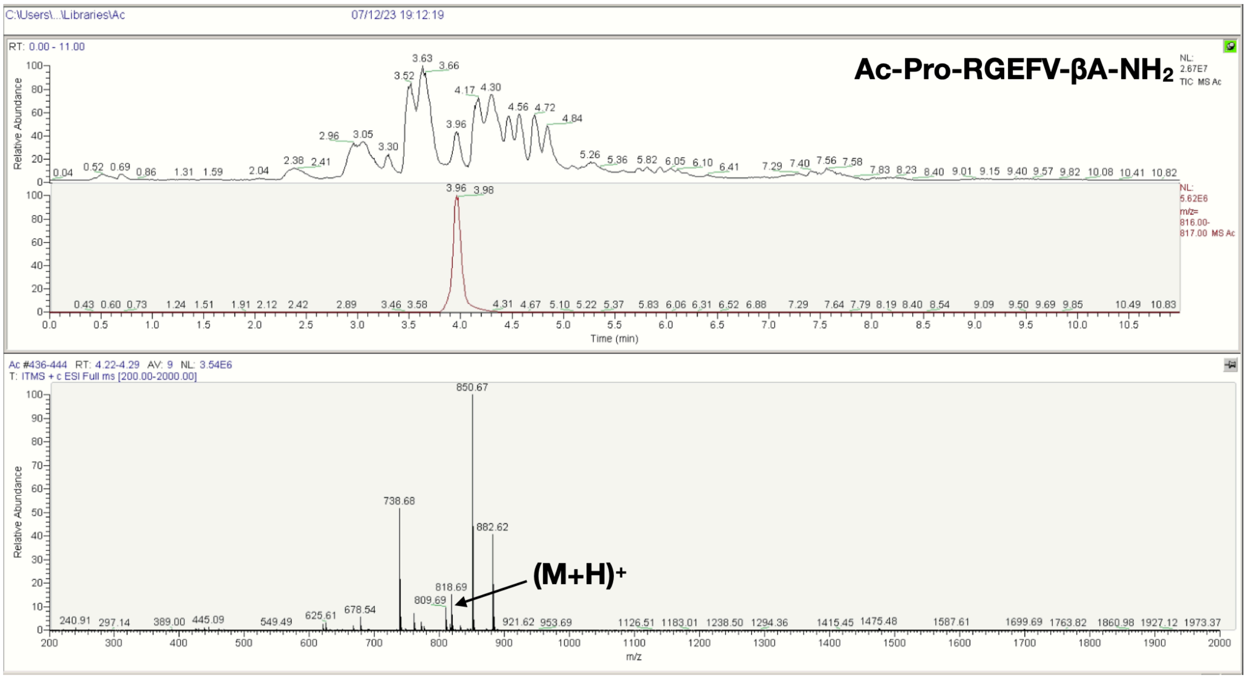
C:\Users\...\Libraries\Ac
07/12/23 19:12:19
RT: 0.00 - 11.00
Ac-Pro-RGEFV-βA-NH2
NL: 2.67E7 TIC MS Ac
NL: 5.62E6 m/z= 816.00-817.00 MS Ac
Relative Abundance
Time (min)
Ac #436-444 RT: 4.22-4.29 AV: 9 NL: 3.54E6
T: ITMS + c ESI Full ms [200.00-2000.00]
850.67
738.68
882.62
818.69
809.69
678.54
(M+H)+
m/z

N

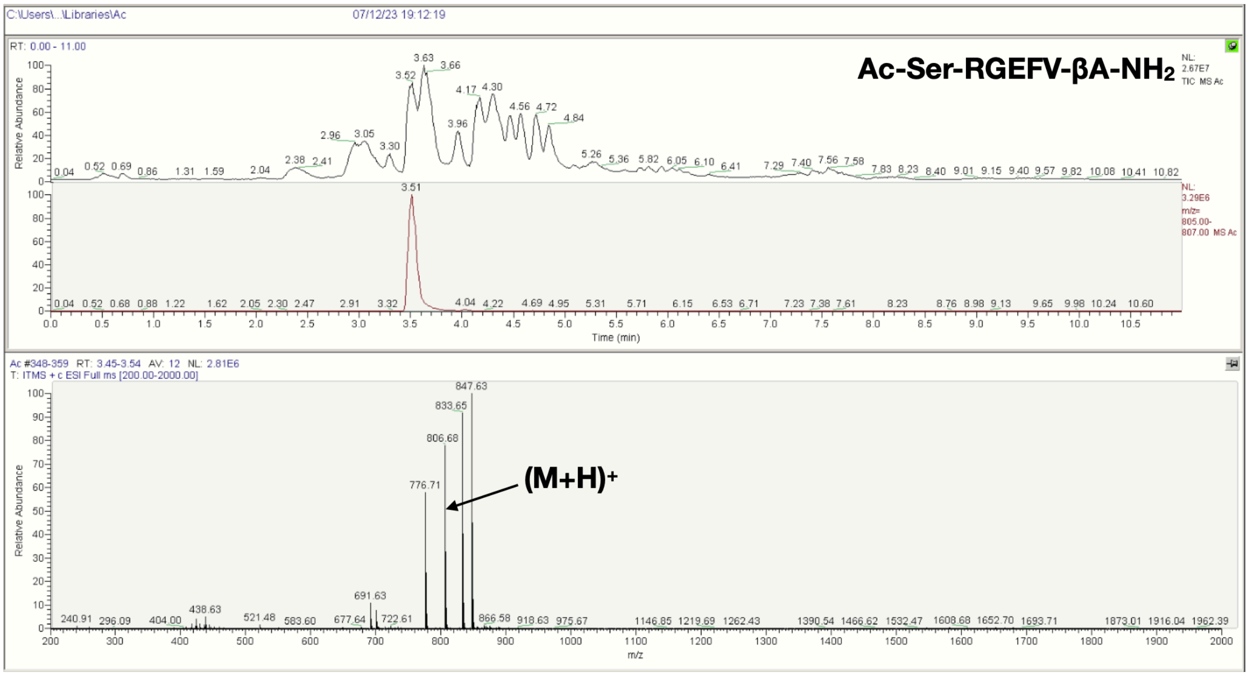
C:\Users\...\Libraries\Ac
07/12/23 19:12:19
RT: 0.00 - 11.00
Ac-Ser-RGEFV-βA-NH2
NL: 2.67E7 TIC MS Ac
NL: 3.29E6 m/z= 805.00-807.00 MS Ac
Relative Abundance
Time (min)
Ac #348-359 RT: 3.45-3.54 AV: 12 NL: 2.81E6
T: ITMS + c ESI Full ms [200.00-2000.00]
847.63
833.65
806.68
776.71
691.63
(M+H)+
m/z

**O**

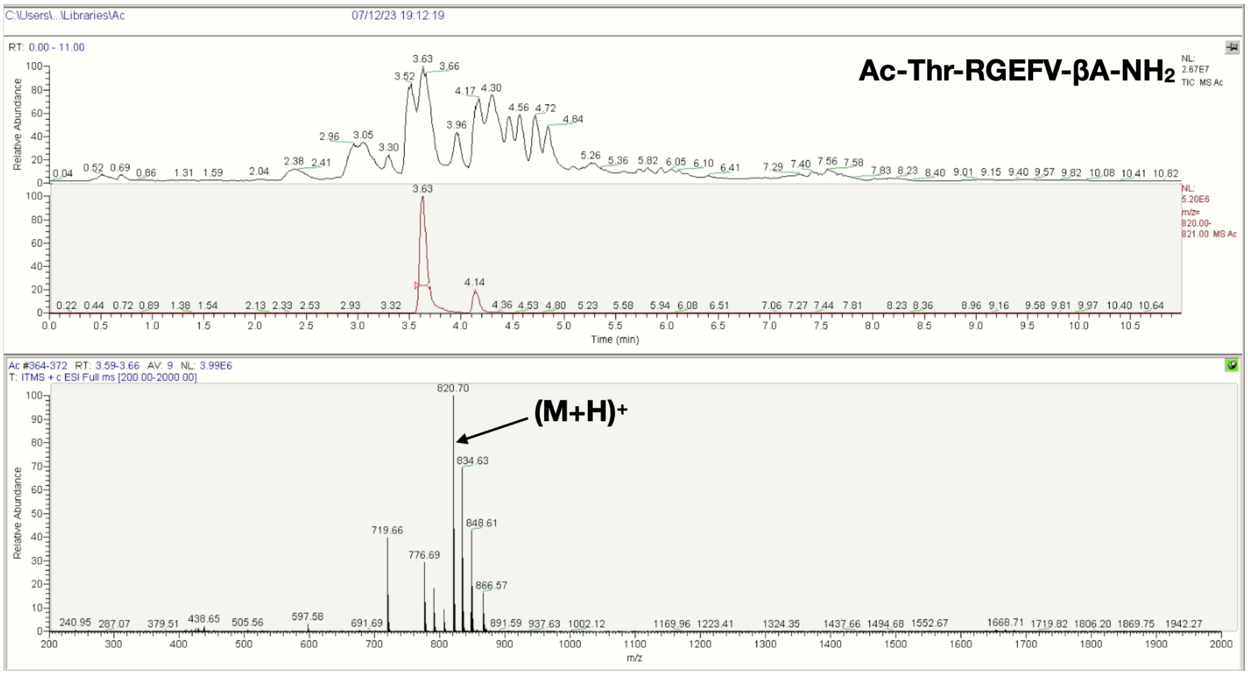
C:\Users\...\Libraries\Ac
07/12/23 19:12:19
RT: 0.00 - 11.00
Ac-Thr-RGEFV-βA-NH2
NL: 2.67E7 TIC MS Ac
NL: 5.20E6 m/z= 820.00-821.00 MS Ac
Relative Abundance
Time (min)
Ac #364-372 RT: 3.59-3.66 AV: 9 NL: 3.99E6
T: ITMS + c ESI Full ms [200.00-2000.00]
820.70
(M+H)+
834.63
848.61
719.66
776.69
866.57
m/z

**P**

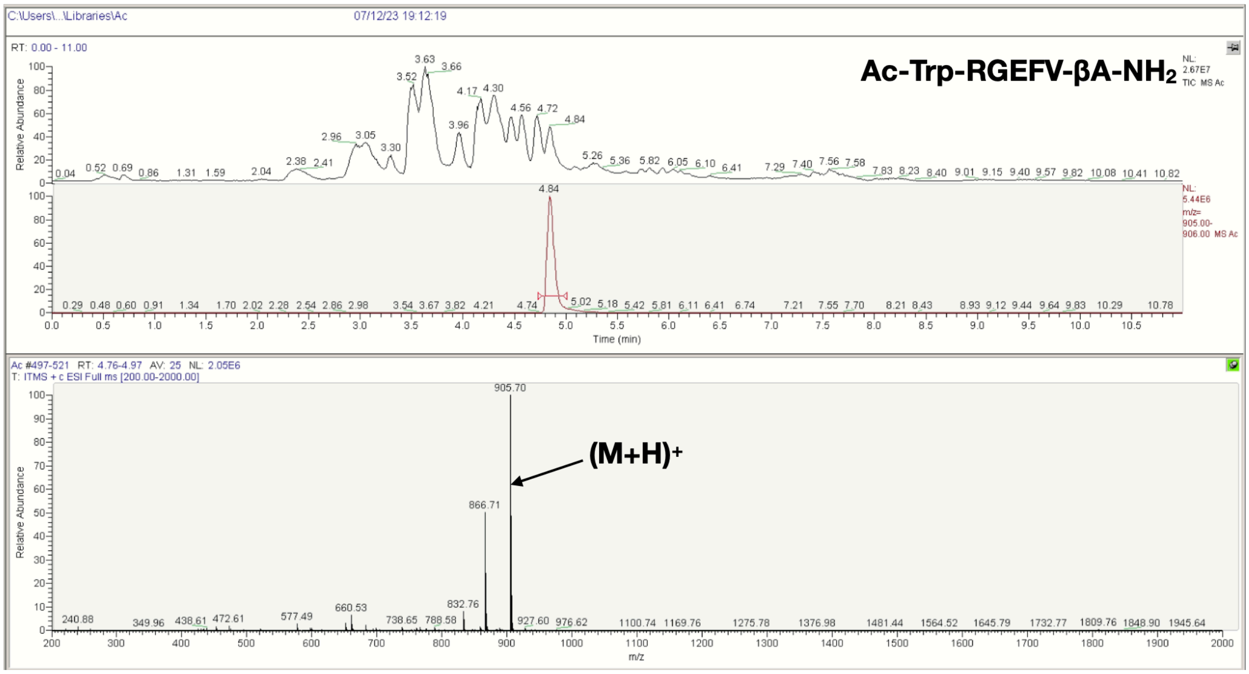
C:\Users\...\Libraries\Ac
07/12/23 19:12:19
RT: 0.00 - 11.00
Ac-Trp-RGEFV-βA-NH2
NL: 2.67E7 TIC MS Ac
NL: 5.44E6 m/z= 905.00-906.00 MS Ac
Relative Abundance
Time (min)
Ac #497-521 RT: 4.76-4.97 AV: 25 NL: 2.05E6
T: ITMS + c ESI Full ms [200.00-2000.00]
905.70
(M+H)+
866.71
832.76
m/z

**Q**

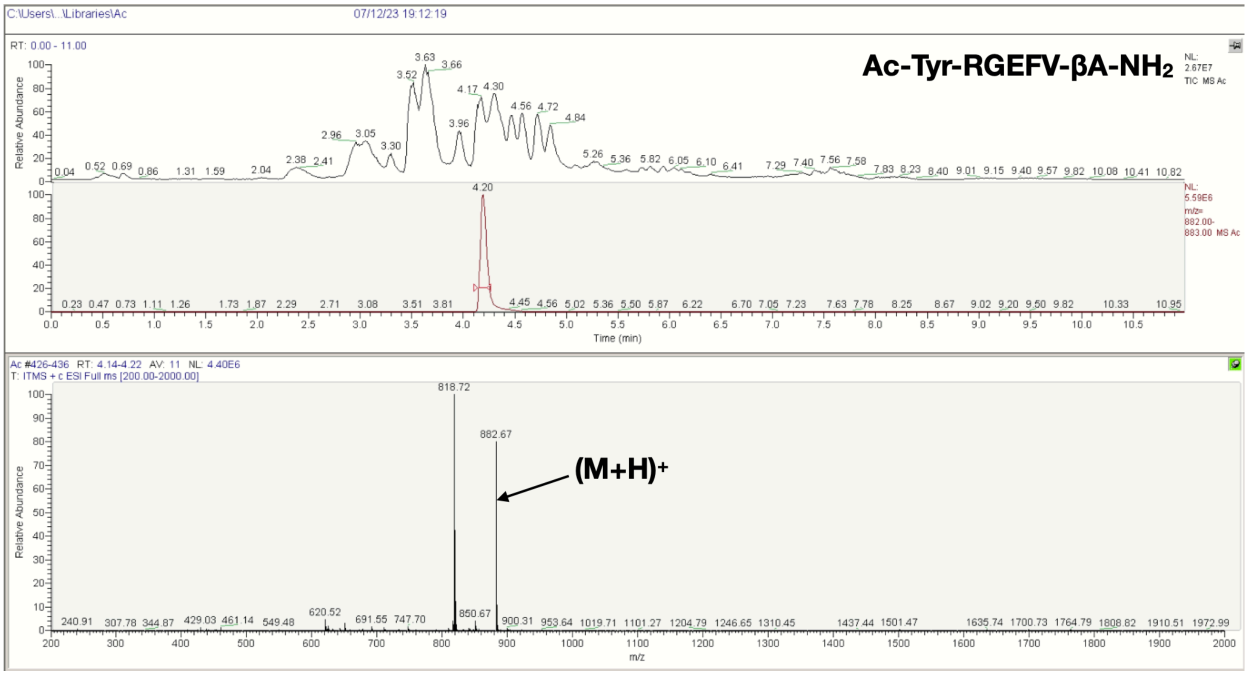


**R**

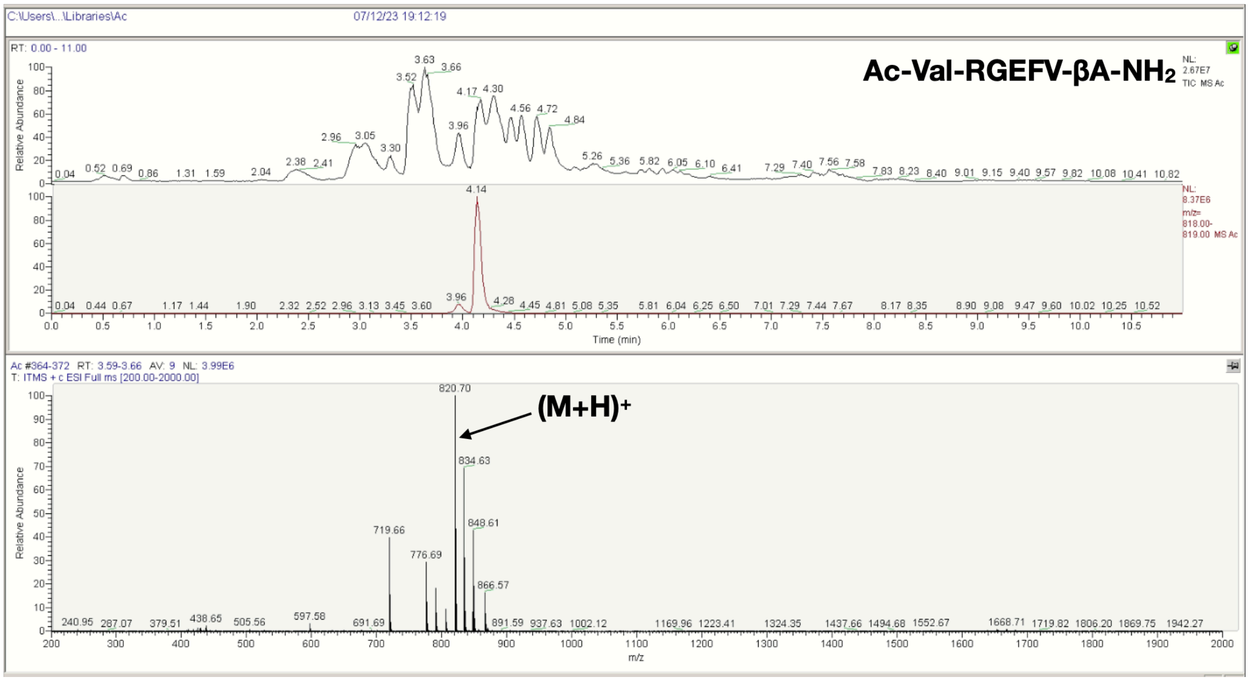


**Figure S12**. LCMS spectra of the Ac-X-RGEFV-βA-$NH_2$ libraries, where X = a) Ala, b) Arg, c) Asn, d) Asp, e) Gln, f) Glu, g) Gly, h) His, i) Ile/Leu, j) Lys, k) Met, l) Phe, m) Pro, n) Ser, o) Thr, p) Trp, q) Tyr, r) Val.

**A**

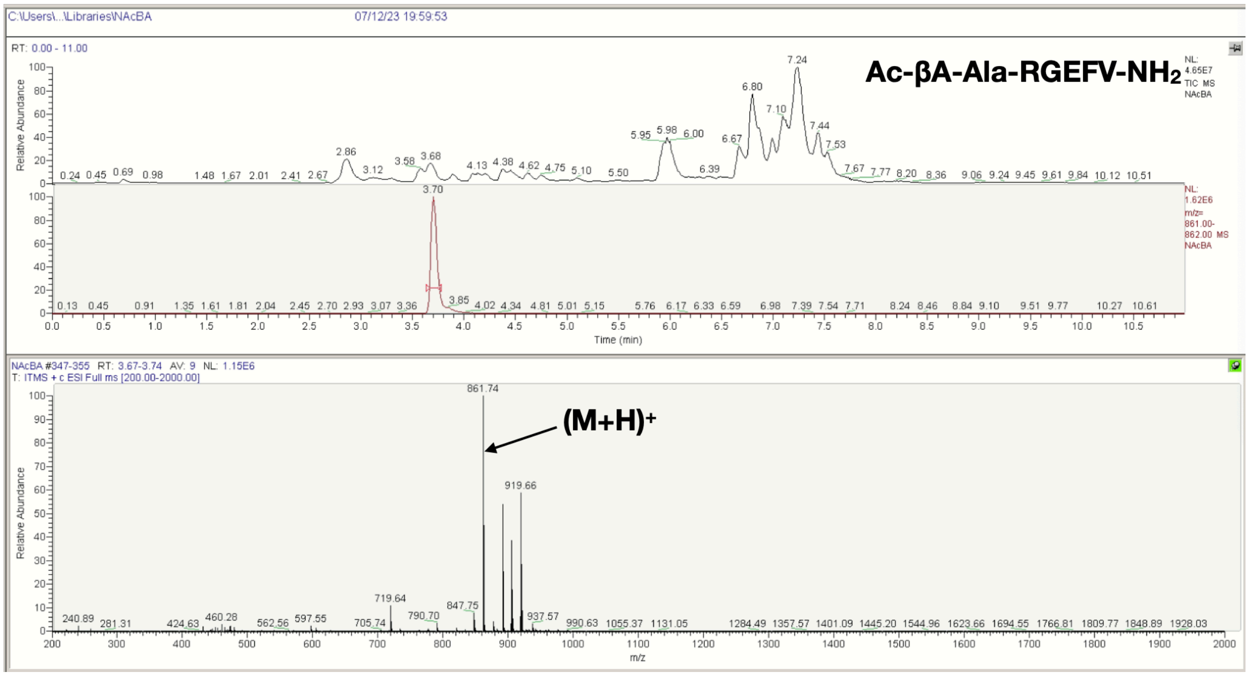

Ac-βA-Ala-RGEFV-NH2
(M+H)+
861.74
919.66
Relative Abundance
Time (min)
m/z


**B**

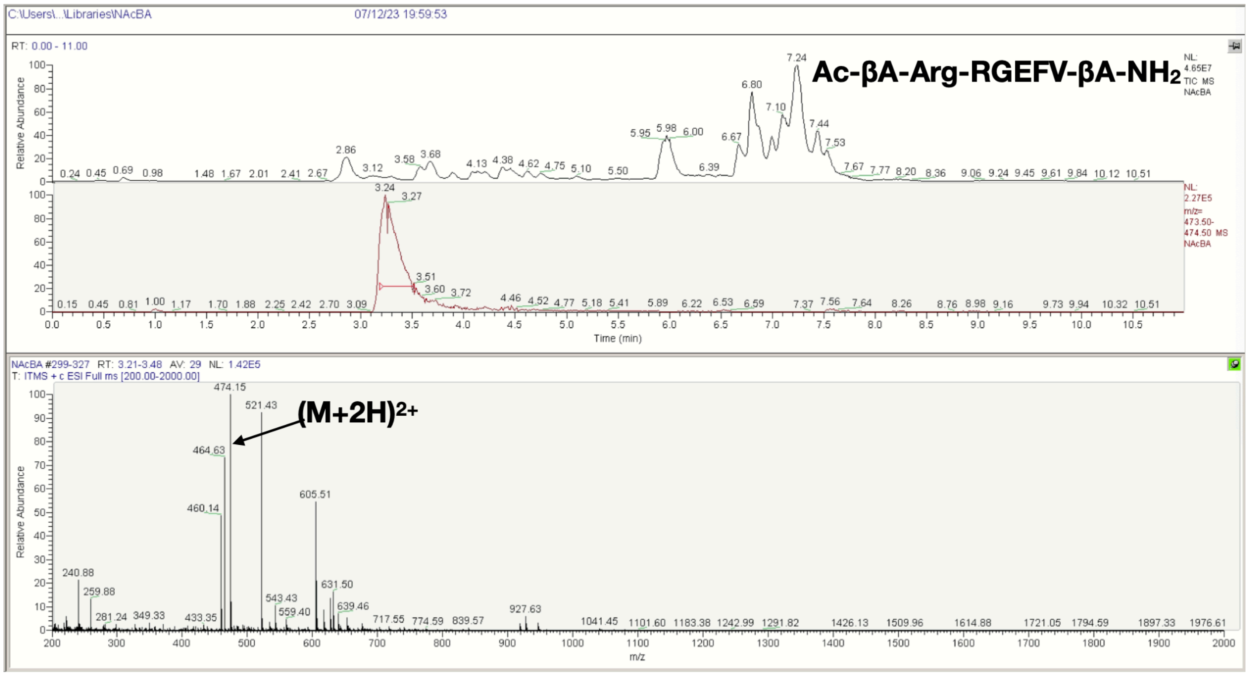

Ac-βA-Arg-RGEFV-βA-NH2
(M+2H)2+
474.15
521.43
464.63
605.51
Relative Abundance
Time (min)
m/z

**C**

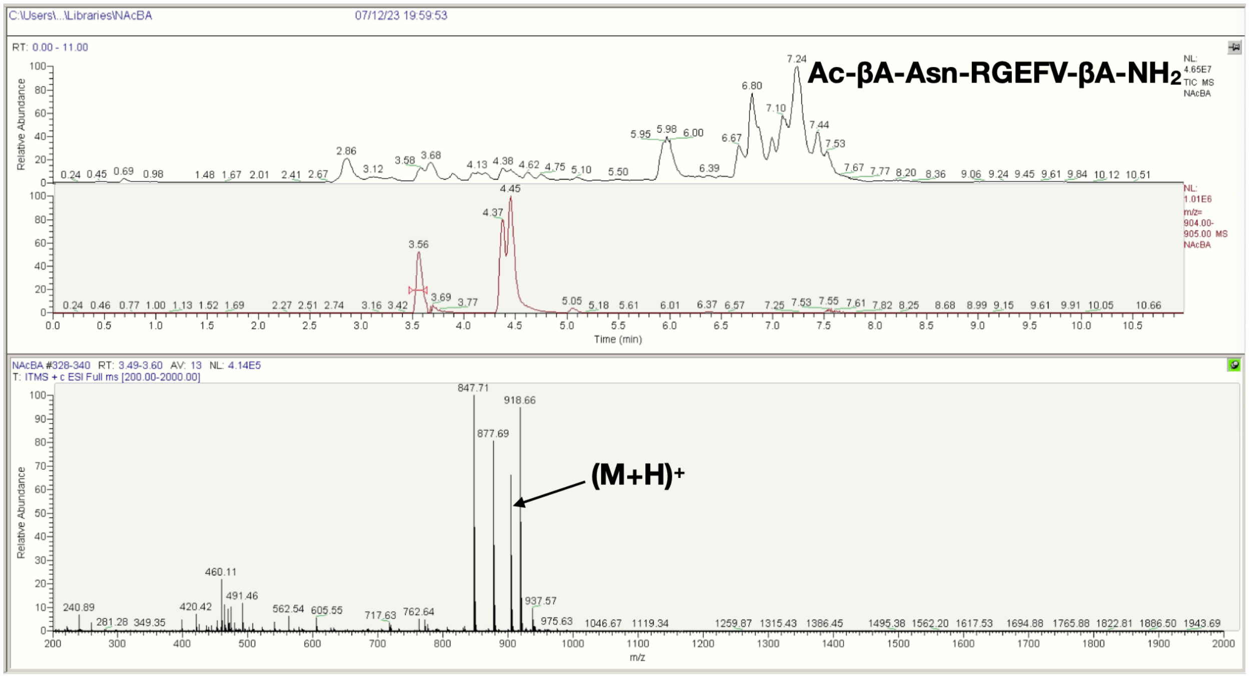
C:\Users\...\Libraries\NAcBA
07/12/23 19:59:53
RT: 0.00 - 11.00
Ac-βA-Asn-RGEFV-βA-NH2
NL: 4.65E7 TIC MS NAcBA
NL: 1.01E6 m/z= 904.00-905.00 MS NAcBA
Relative Abundance
Time (min)
NAcBA #328-340 RT: 3.49-3.60 AV: 13 NL: 4.14E5
T: ITMS + c ESI Full ms [200.00-2000.00]
847.71
918.66
877.69
(M+H)+
m/z

**D**

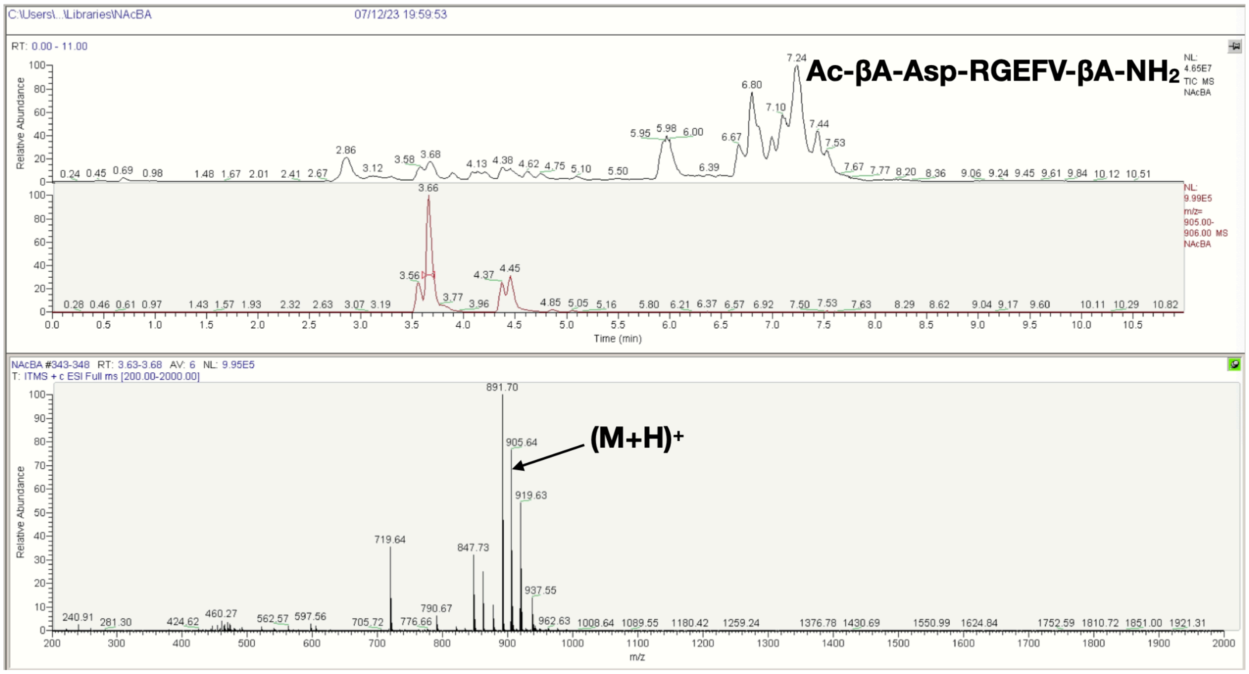
C:\Users\...\Libraries\NAcBA
07/12/23 19:59:53
RT: 0.00 - 11.00
Ac-βA-Asp-RGEFV-βA-NH2
NL: 4.65E7 TIC MS NAcBA
NL: 9.99E5 m/z= 905.00-906.00 MS NAcBA
Relative Abundance
Time (min)
NAcBA #343-348 RT: 3.63-3.68 AV: 6 NL: 9.95E5
T: ITMS + c ESI Full ms [200.00-2000.00]
891.70
905.64
919.63
(M+H)+
m/z

E

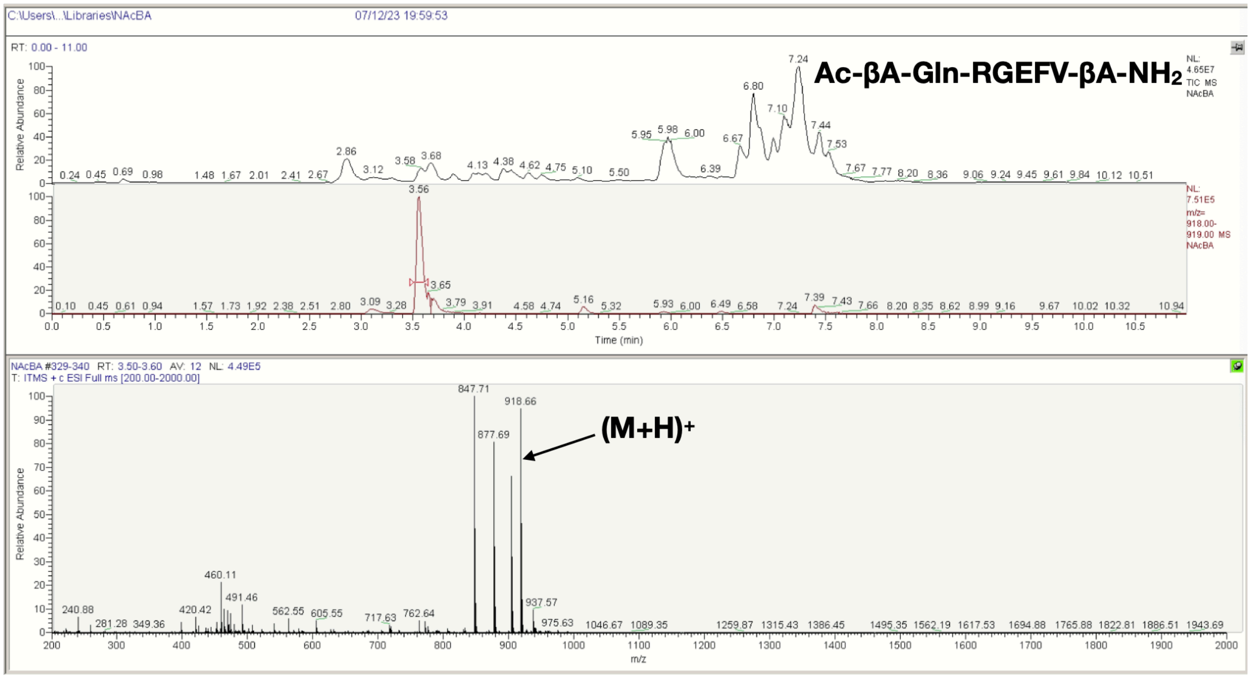


F

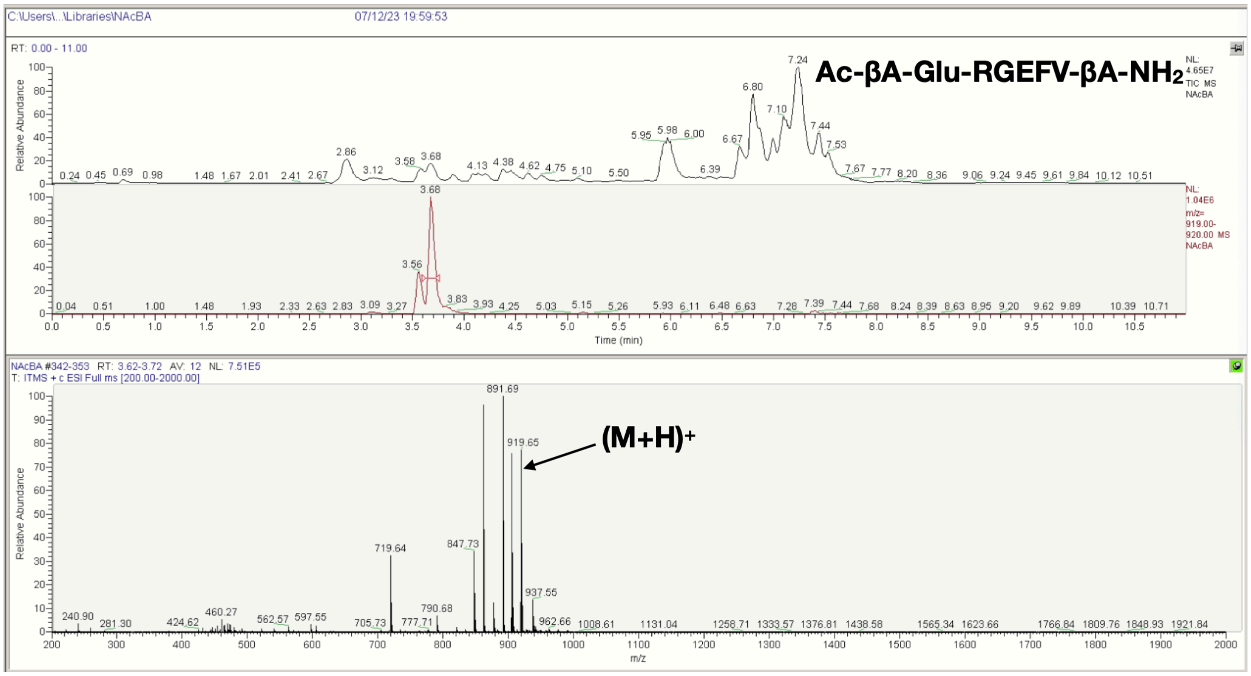

G

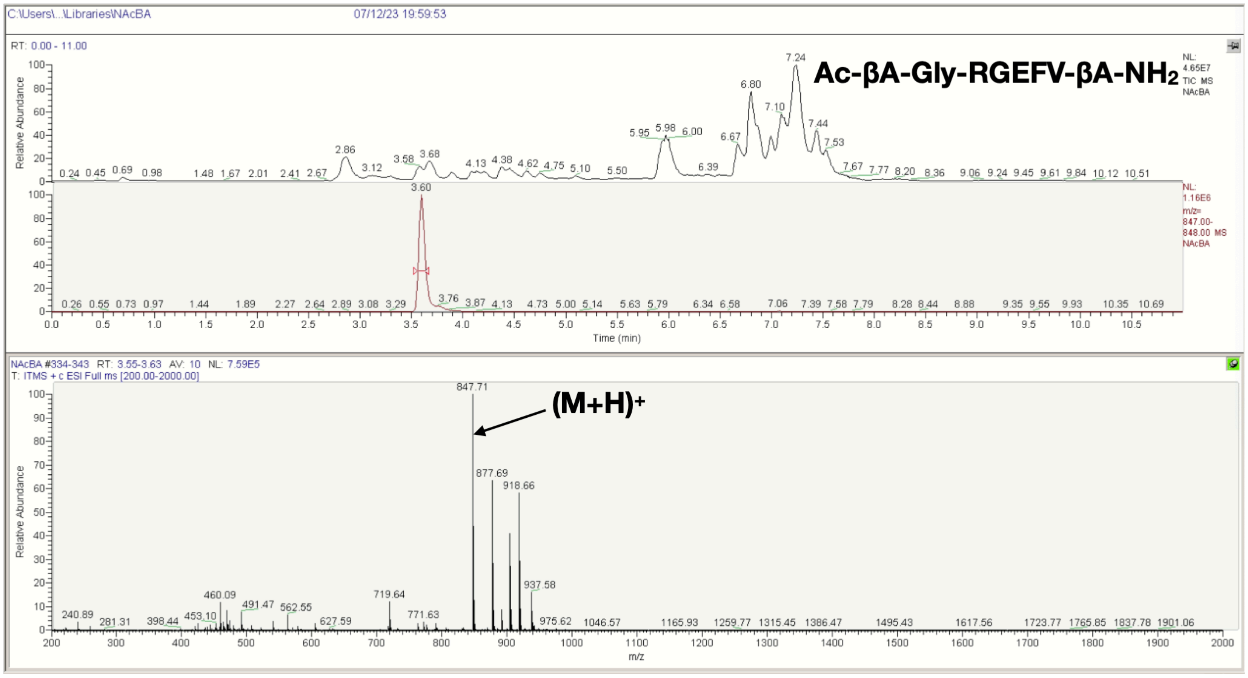
C:\Users\...\Libraries\NAcBA
07/12/23 19:59:53
RT: 0.00 - 11.00
Ac-βA-Gly-RGEFV-βA-NH2
NL: 4.65E7 TIC MS NAcBA
NL: 1.16E6 m/z= 847.00-848.00 MS NAcBA
Relative Abundance
Time (min)
NAcBA #334-343 RT: 3.55-3.63 AV: 10 NL: 7.59E5
T: ITMS + c ESI Full ms [200.00-2000.00]
(M+H)+
847.71
877.69
918.66
937.58
m/z

H

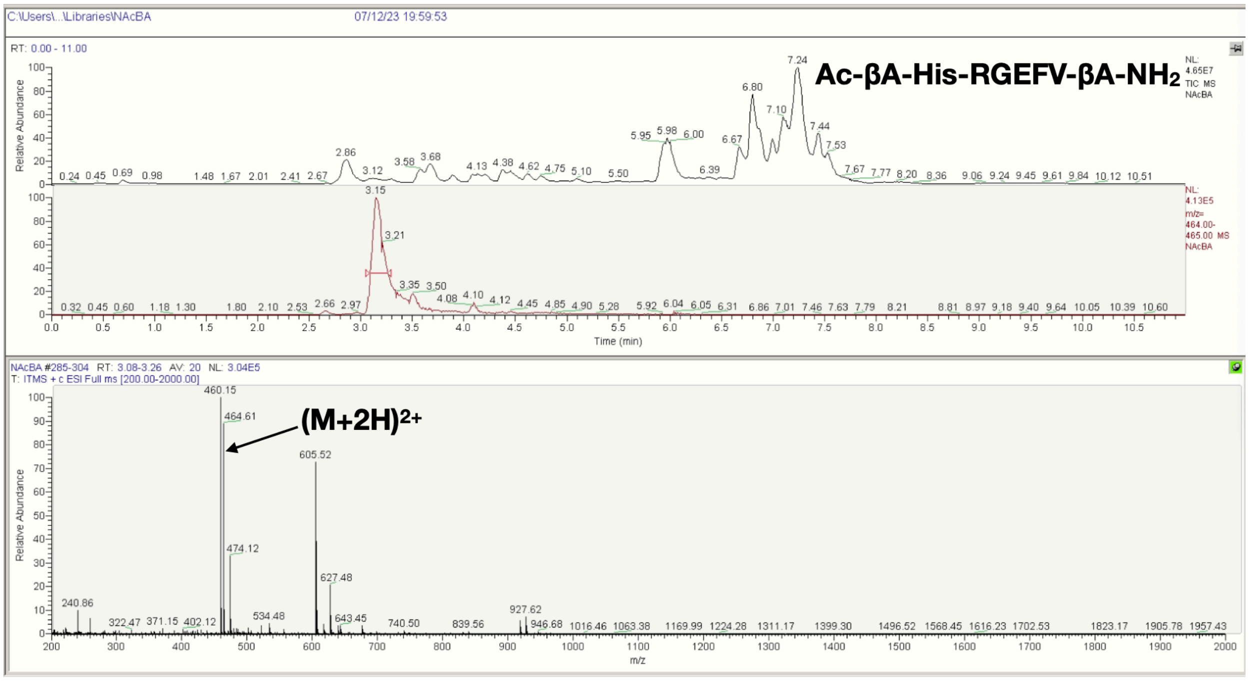
C:\Users\...\Libraries\NAcBA
07/12/23 19:59:53
RT: 0.00 - 11.00
Ac-βA-His-RGEFV-βA-NH2
NL: 4.65E7 TIC MS NAcBA
NL: 4.13E5 m/z= 464.00-465.00 MS NAcBA
Relative Abundance
Time (min)
NAcBA #285-304 RT: 3.08-3.26 AV: 20 NL: 3.04E5
T: ITMS + c ESI Full ms [200.00-2000.00]
(M+2H)2+
460.15
464.61
605.52
474.12
627.48
m/z

**I**

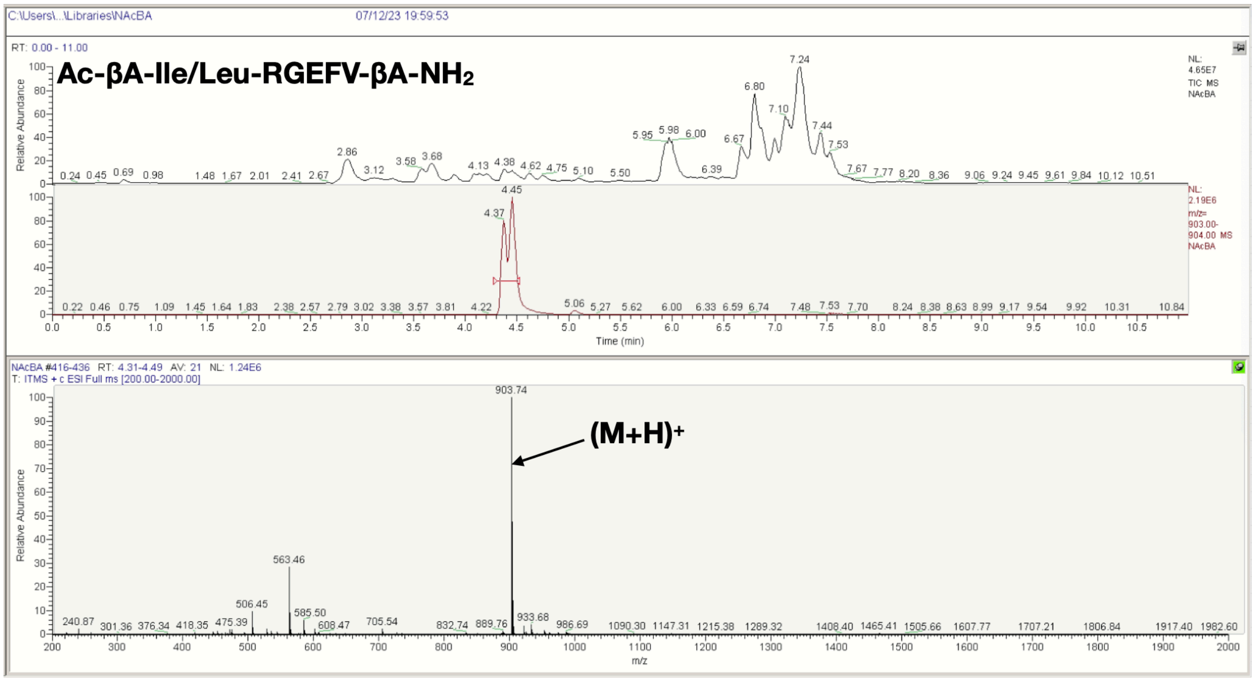
C:\Users\...\Libraries\NAcBA
07/12/23 19:59:53
RT: 0.00 - 11.00
Ac-βA-Ile/Leu-RGEFV-βA-NH2
NL: 4.65E7 TIC MS NAcBA
NL: 2.19E6 m/z= 903.00-904.00 MS NAcBA
Relative Abundance
Time (min)
NAcBA #416-436 RT: 4.31-4.49 AV: 21 NL: 1.24E6
T: ITMS + c ESI Full ms [200.00-2000.00]
903.74
(M+H)+
563.46
m/z

**J**

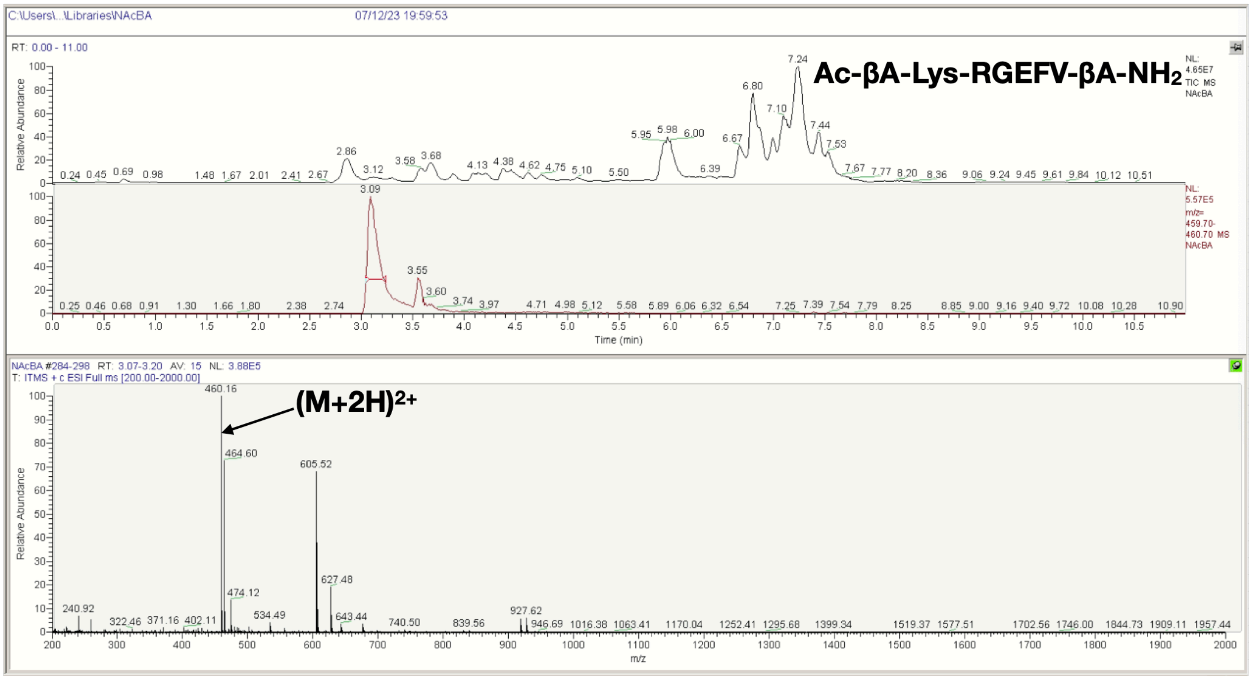
C:\Users\...\Libraries\NAcBA
07/12/23 19:59:53
RT: 0.00 - 11.00
Ac-βA-Lys-RGEFV-βA-NH2
NL: 4.65E7 TIC MS NAcBA
NL: 5.57E5 m/z= 459.70-460.70 MS NAcBA
Relative Abundance
Time (min)
NAcBA #284-298 RT: 3.07-3.20 AV: 15 NL: 3.88E5
T: ITMS + c ESI Full ms [200.00-2000.00]
460.16
(M+2H)2+
464.60
605.52
m/z

**K**

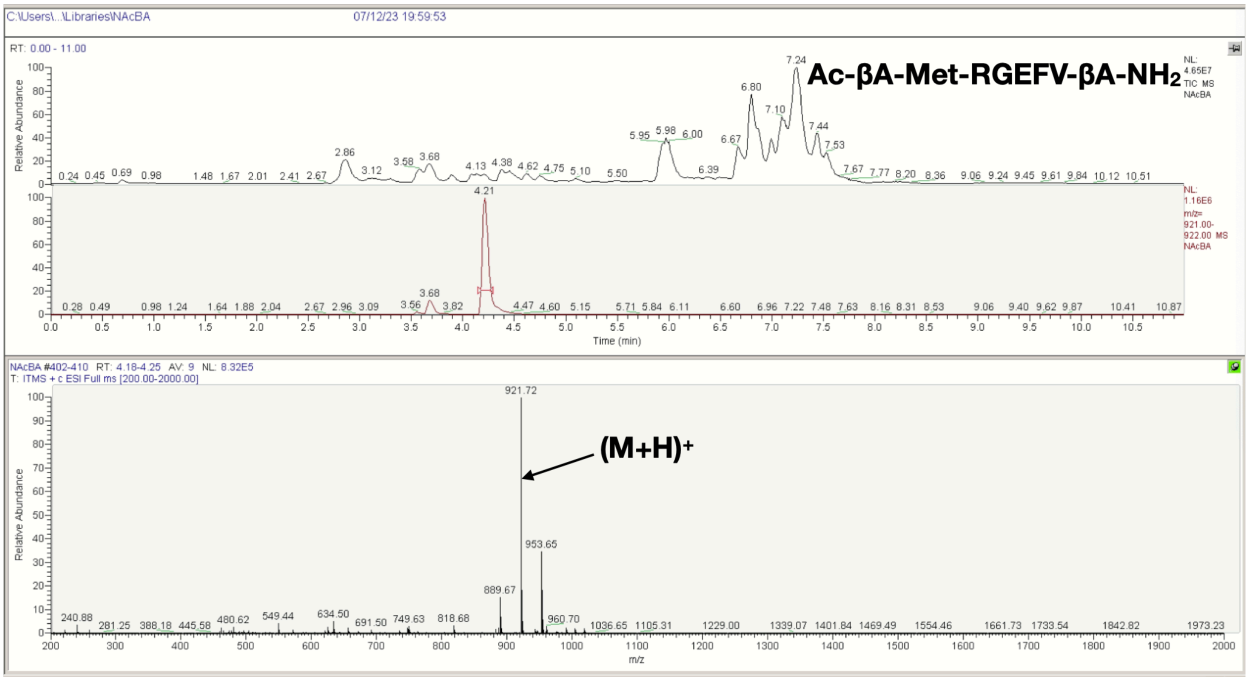

C:\Users\...\Libraries\NAcBA
07/12/23 19:59:53
Ac-βA-Met-RGEFV-βA-NH2
NAcBA #402-410 RT: 4.18-4.25 AV: 9 NL: 8.32E5
T: ITMS + c ESI Full ms [200.00-2000.00]
921.72
(M+H)+
953.65
889.67
Relative Abundance
Time (min)
m/z


**L**

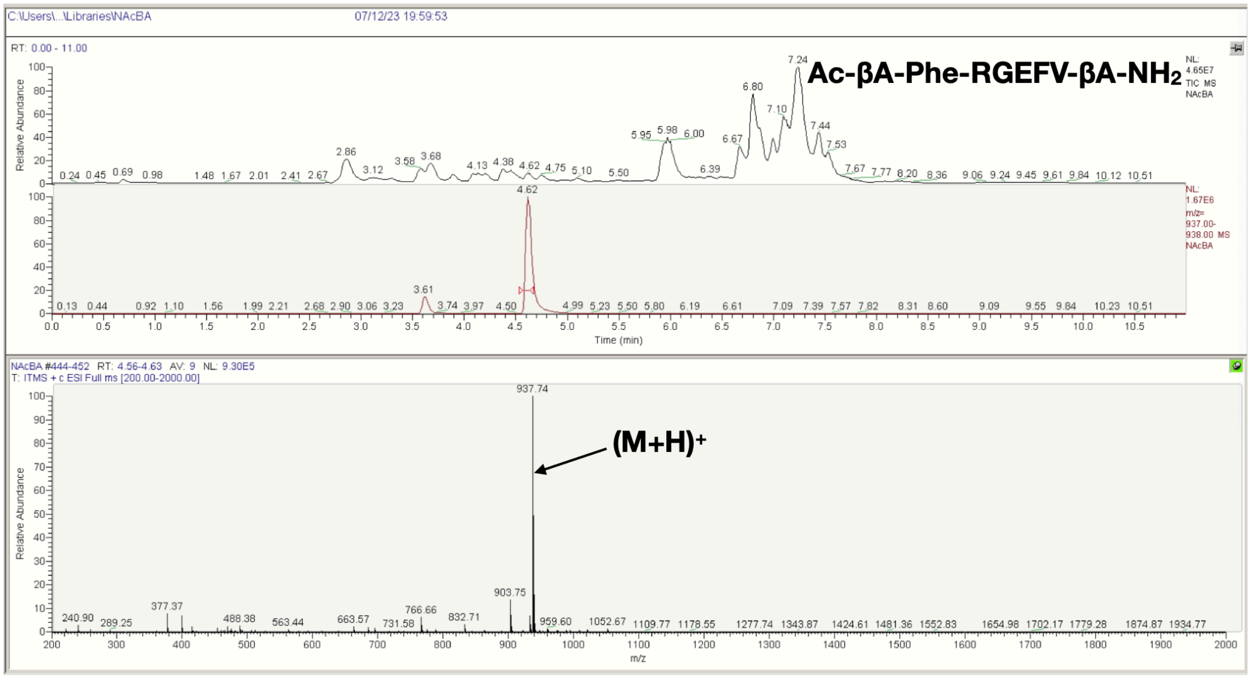

C:\Users\...\Libraries\NAcBA
07/12/23 19:59:53
Ac-βA-Phe-RGEFV-βA-NH2
NAcBA #444-452 RT: 4.56-4.63 AV: 9 NL: 9.30E5
T: ITMS + c ESI Full ms [200.00-2000.00]
937.74
(M+H)+
903.75
Relative Abundance
Time (min)
m/z

M

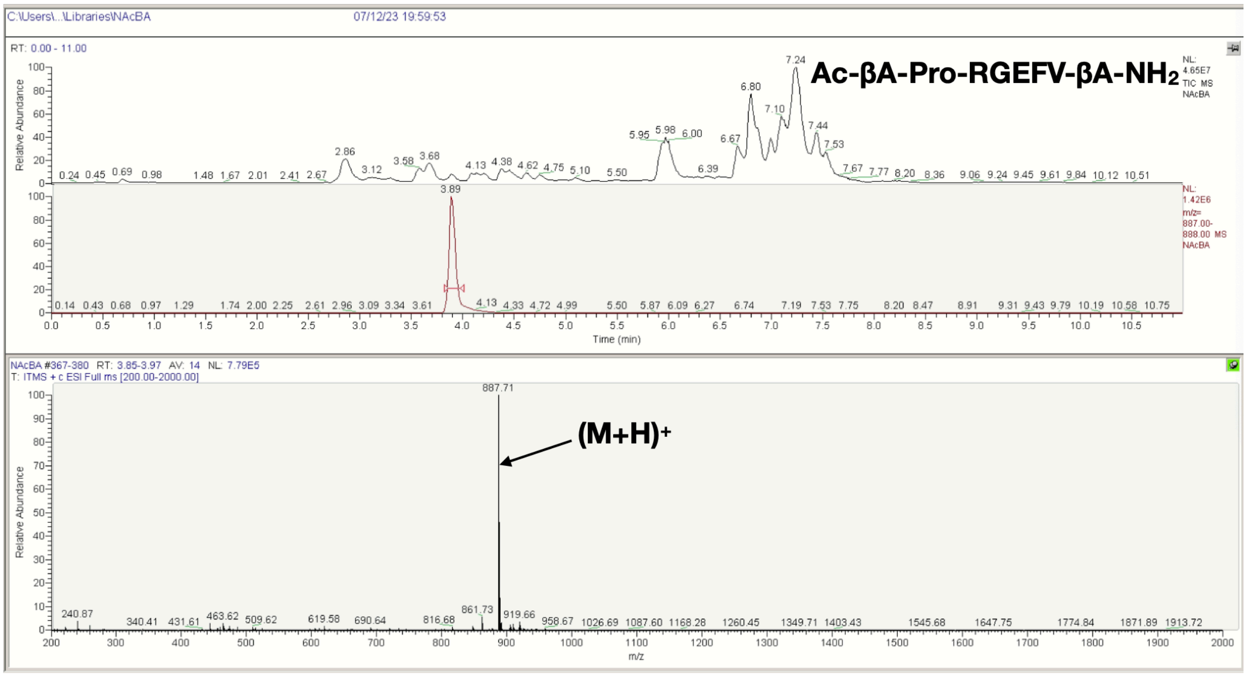
C:\Users\...\Libraries\NAcBA
07/12/23 19:59:53
RT: 0.00 - 11.00
Ac-βA-Pro-RGEFV-βA-NH2
NL: 4.65E7 TIC MS NAcBA
NL: 1.42E6 m/z= 887.00-888.00 MS NAcBA
Relative Abundance
Time (min)
NAcBA #367-380 RT: 3.85-3.97 AV: 14 NL: 7.79E5
T: ITMS + c ESI Full ms [200.00-2000.00]
887.71
(M+H)+
m/z

N

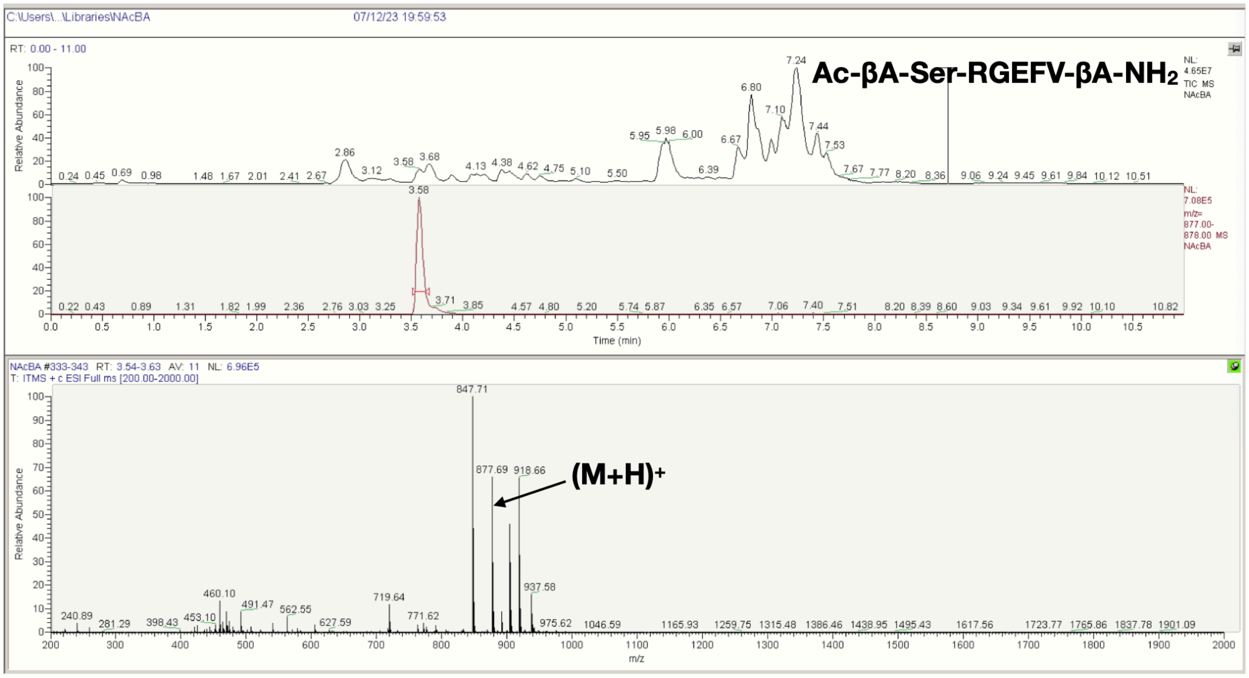
C:\Users\...\Libraries\NAcBA
07/12/23 19:59:53
RT: 0.00 - 11.00
Ac-βA-Ser-RGEFV-βA-NH2
NL: 4.65E7 TIC MS NAcBA
NL: 7.08E5 m/z= 877.00-878.00 MS NAcBA
Relative Abundance
Time (min)
NAcBA #333-343 RT: 3.54-3.63 AV: 11 NL: 6.96E5
T: ITMS + c ESI Full ms [200.00-2000.00]
847.71
877.69
918.66
(M+H)+
m/z

**O**

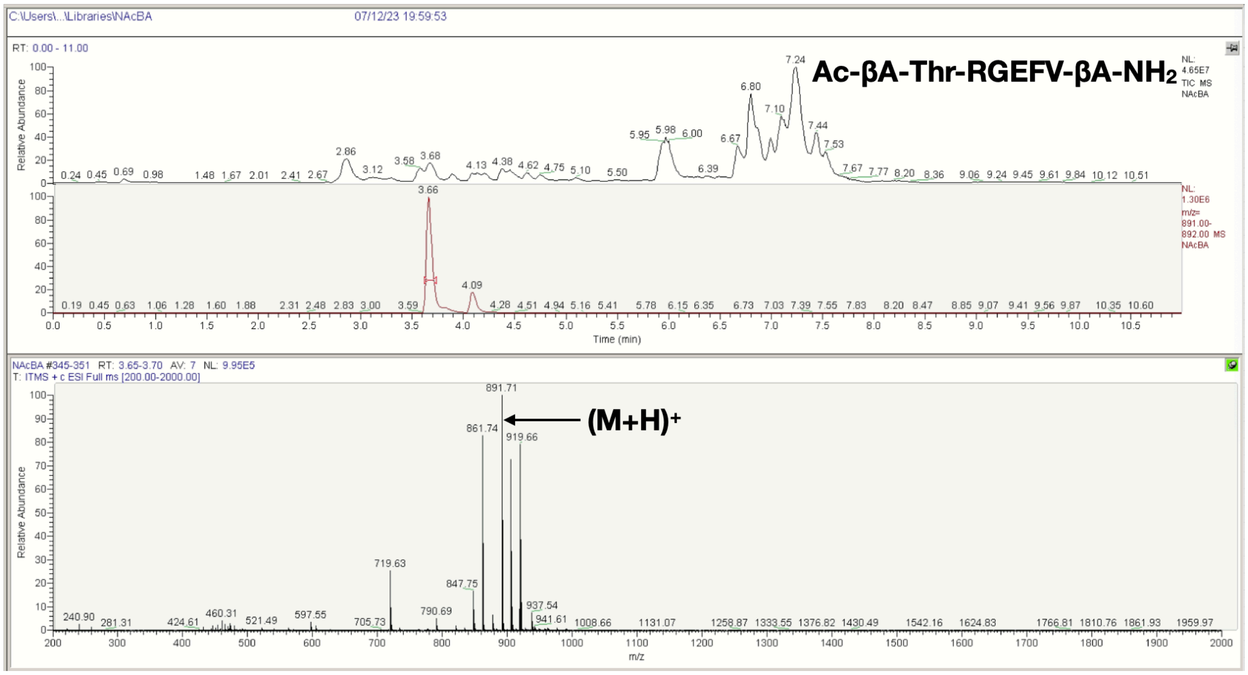

C:\Users\...\Libraries\NAcBA
07/12/23 19:59:53
RT: 0.00 - 11.00
Ac-βA-Thr-RGEFV-βA-NH2
NL: 4.65E7 TIC MS NAcBA
NL: 1.30E6 m/z= 891.00-892.00 MS NAcBA
Relative Abundance
Time (min)
NAcBA #345-351 RT: 3.65-3.70 AV: 7 NL: 9.95E5
T: ITMS + c ESI Full ms [200.00-2000.00]
891.71
(M+H)+
861.74
919.66
719.63
847.75
937.54
m/z


**P**

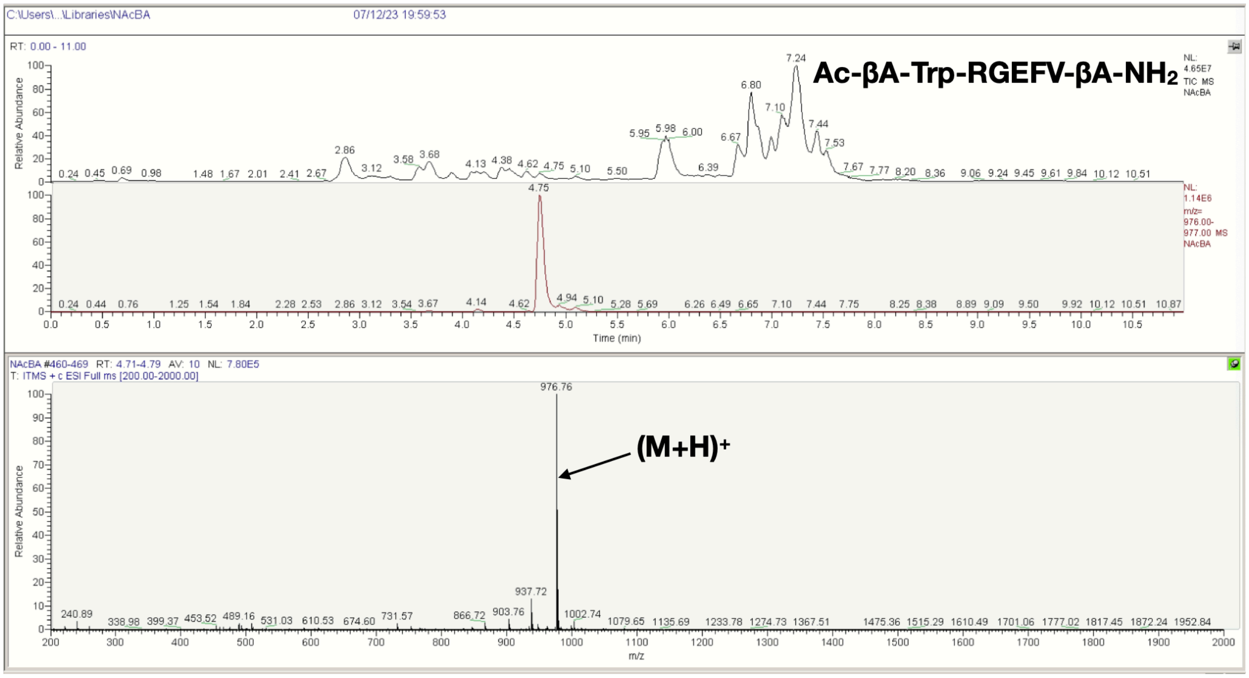

C:\Users\...\Libraries\NAcBA
07/12/23 19:59:53
RT: 0.00 - 11.00
Ac-βA-Trp-RGEFV-βA-NH2
NL: 4.65E7 TIC MS NAcBA
NL: 1.14E6 m/z= 976.00-977.00 MS NAcBA
Relative Abundance
Time (min)
NAcBA #460-469 RT: 4.71-4.79 AV: 10 NL: 7.80E5
T: ITMS + c ESI Full ms [200.00-2000.00]
976.76
(M+H)+
937.72
903.76
1002.74
m/z

**Q**

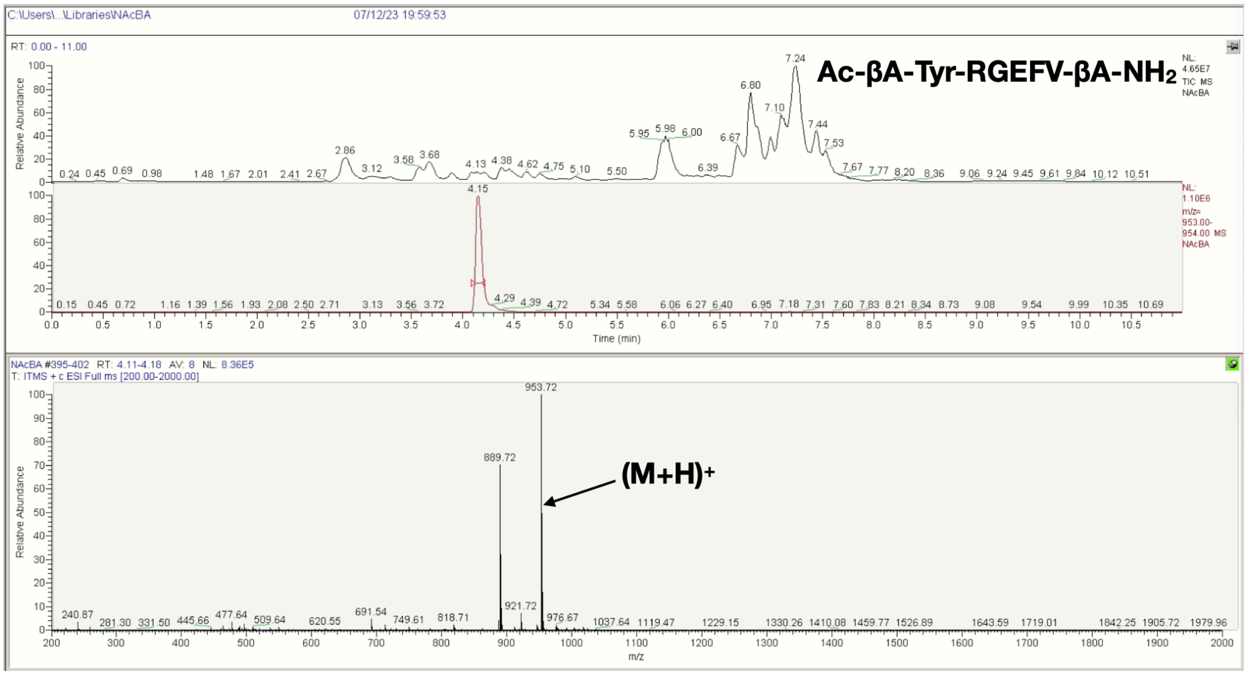


**R**

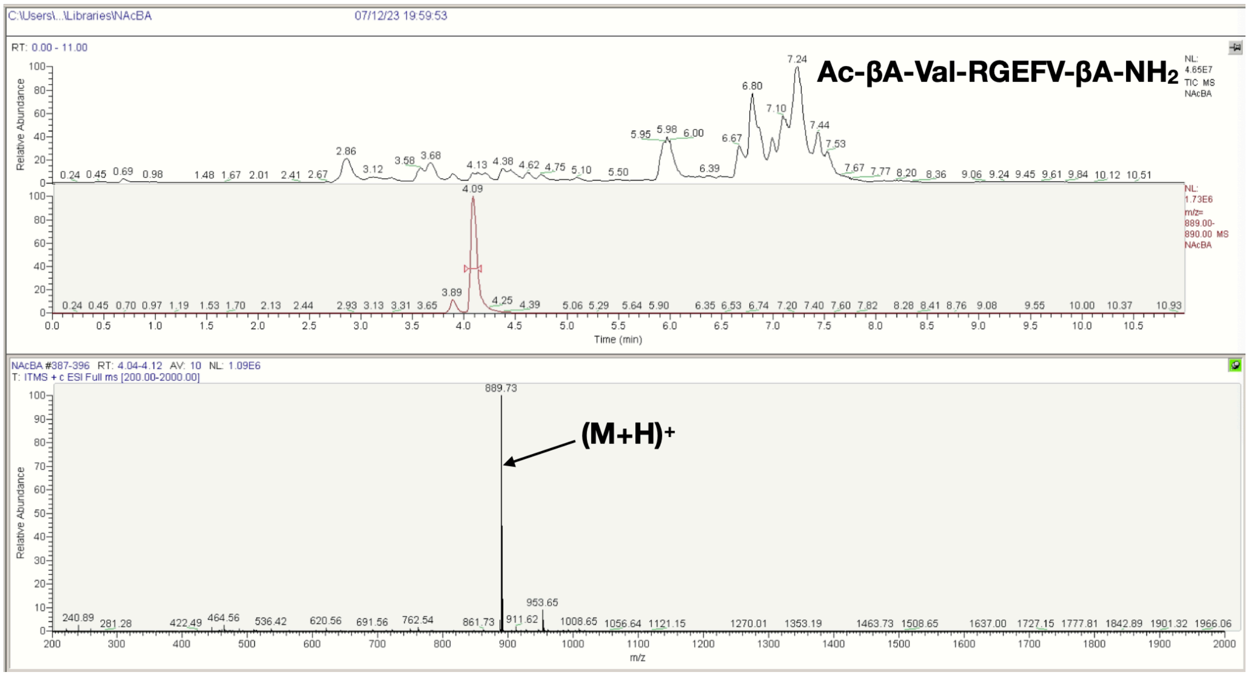


**Figure S13**. LCMS spectra of the Ac-βA-X-RGEFV-βA-$NH_2$ libraries, where X = a) Ala, b) Arg, c) Asn, d) Asp, e) Gln, f) Glu, g) Gly, h) His, i) Ile/Leu, j) Lys, k) Met, l) Phe, m) Pro, n) Ser, o) Thr, p) Trp, q) Tyr, r) Val.

**A**

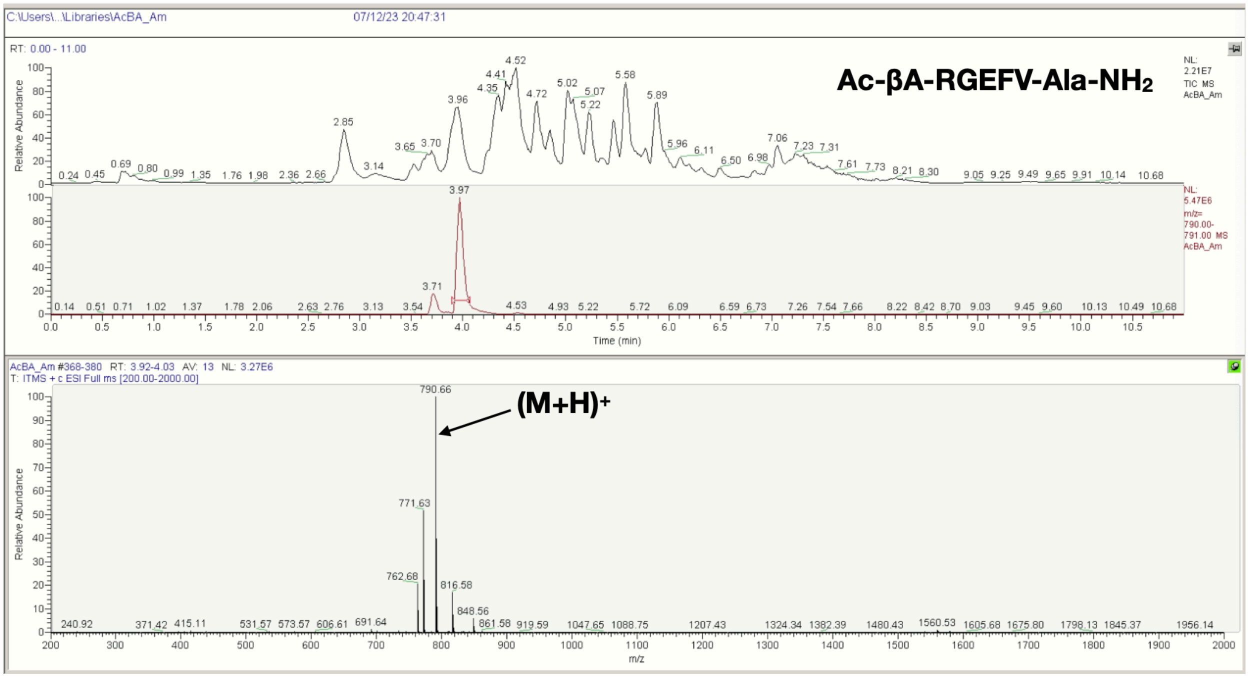

Ac-βA-RGEFV-Ala-NH2
(M+H)+
790.66
Relative Abundance
Time (min)
m/z


**B**

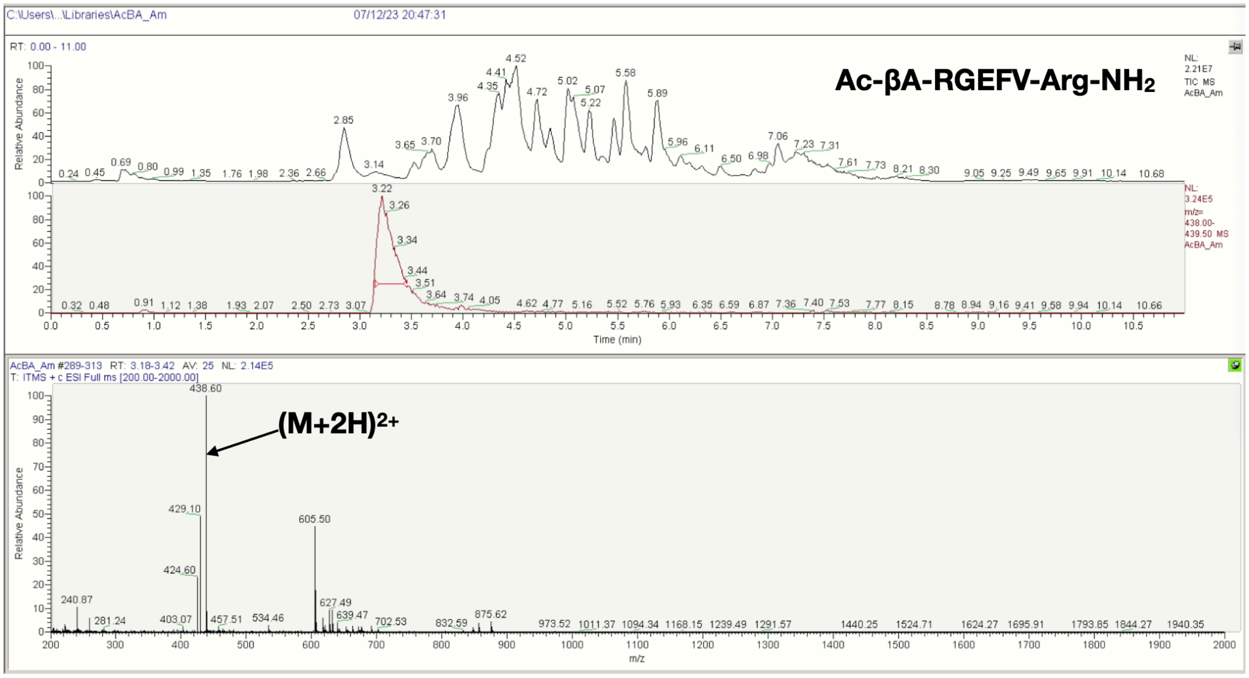

Ac-βA-RGEFV-Arg-NH2
(M+2H)2+
438.60
Relative Abundance
Time (min)
m/z

C

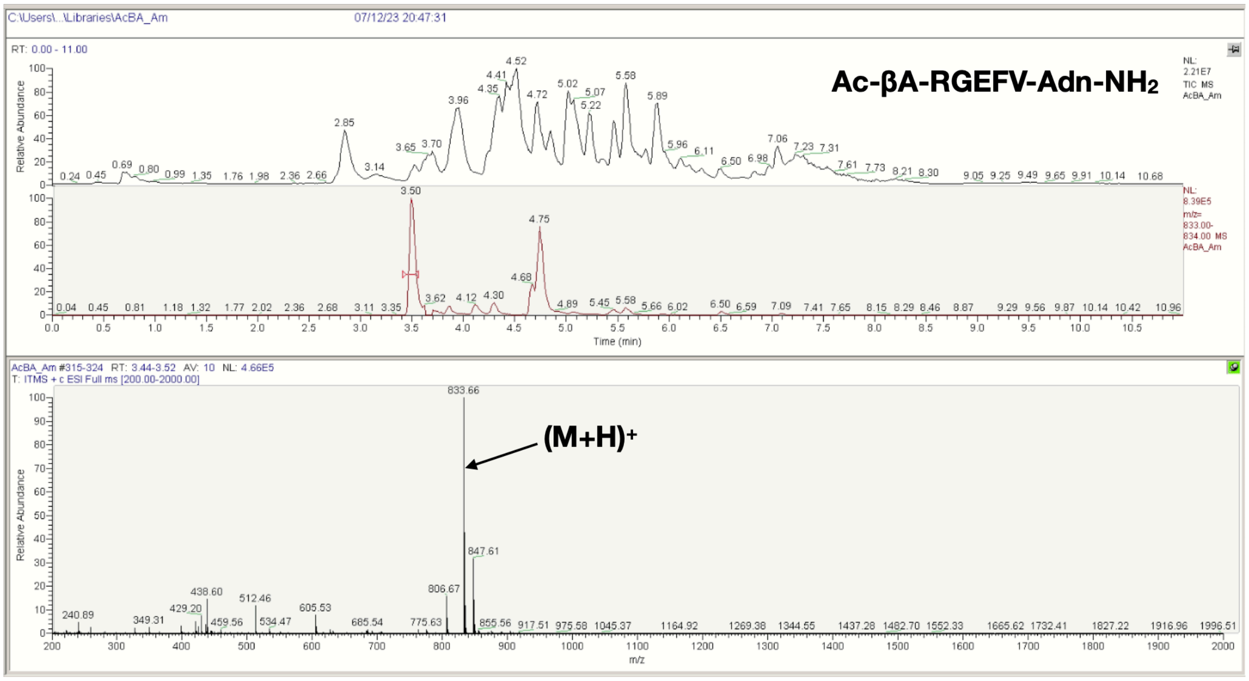

C:\Users\...\Libraries\AcBA_Am
07/12/23 20:47:31
Ac-βA-RGEFV-Adn-NH2
(M+H)+
AcBA_Am #315-324 RT: 3.44-3.52 AV: 10 NL: 4.66E5
T: ITMS + c ESI Full ms [200.00-2000.00]
Relative Abundance
Time (min)
m/z


D

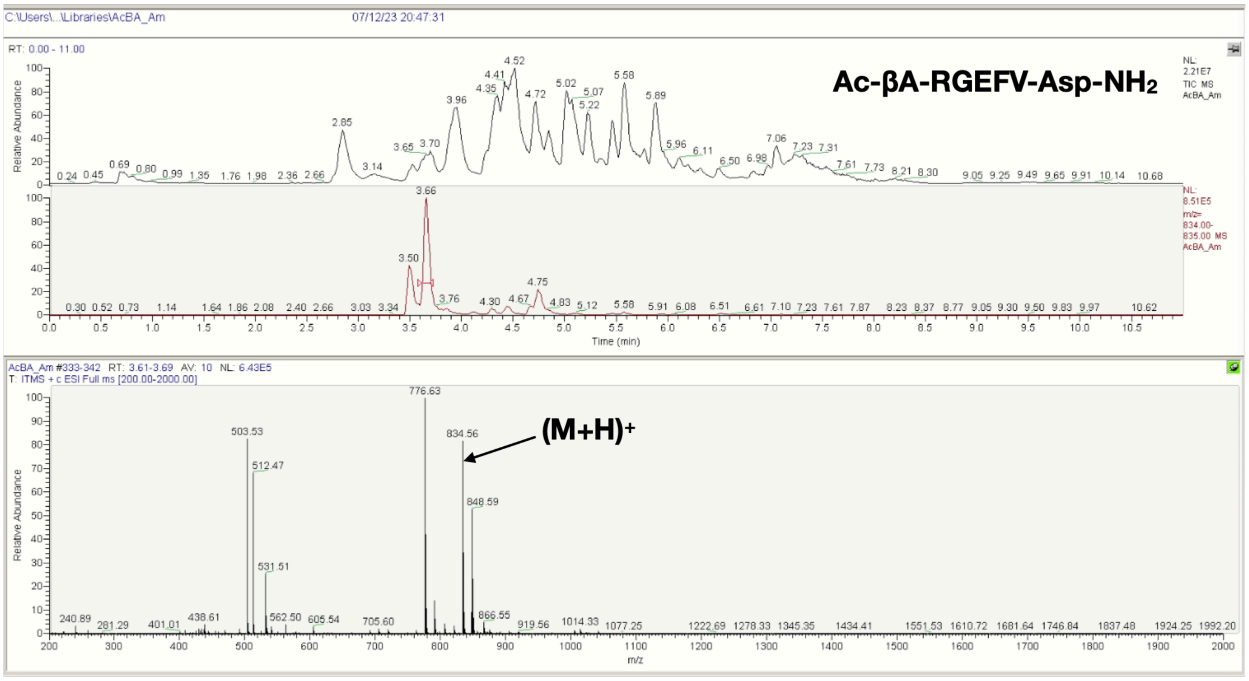

C:\Users\...\Libraries\AcBA_Am
07/12/23 20:47:31
Ac-βA-RGEFV-Asp-NH2
(M+H)+
AcBA_Am #333-342 RT: 3.61-3.69 AV: 10 NL: 6.43E5
T: ITMS + c ESI Full ms [200.00-2000.00]
Relative Abundance
Time (min)
m/z

**E**

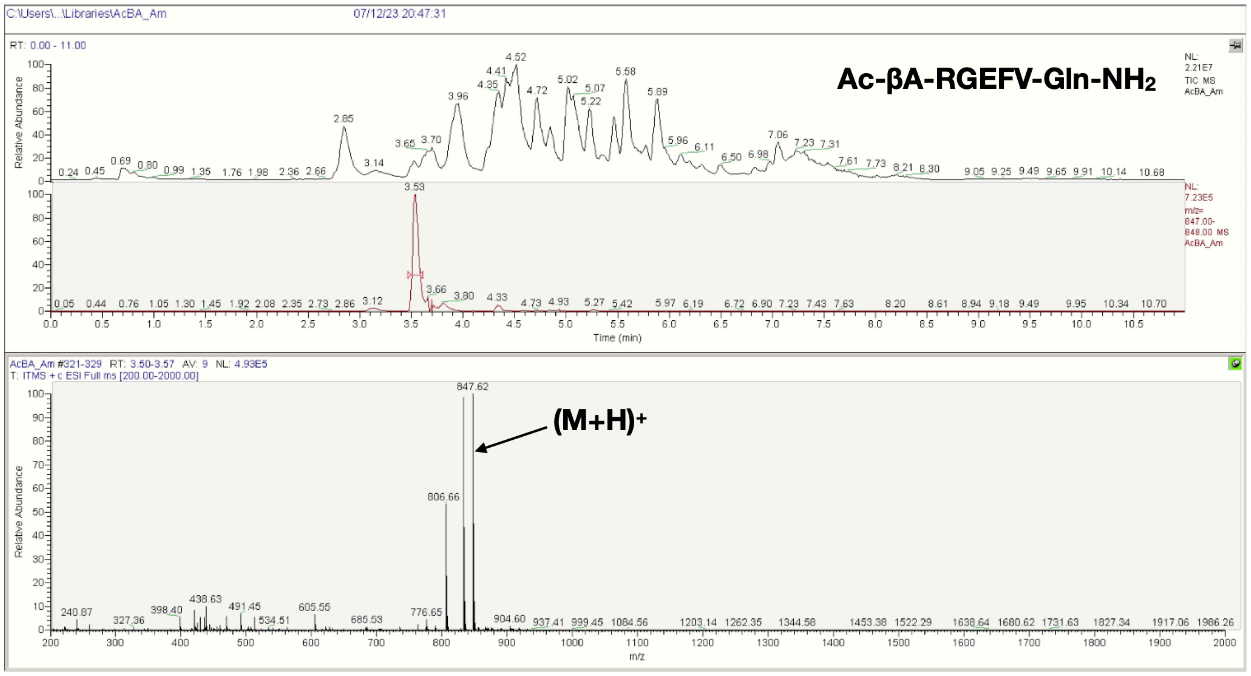
C:\Users\...\Libraries\AcBA_Am
07/12/23 20:47:31
RT: 0.00 - 11.00
Ac-βA-RGEFV-Gln-NH2
NL: 2.21E7 TIC MS AcBA_Am
NL: 7.23E5 m/z= 847.00-848.00 MS AcBA_Am
Relative Abundance
Time (min)
AcBA_Am #321-329 RT: 3.50-3.57 AV: 9 NL: 4.93E5
T: ITMS + c ESI Full ms [200.00-2000.00]
847.62
806.66
(M+H)+
m/z

**F**

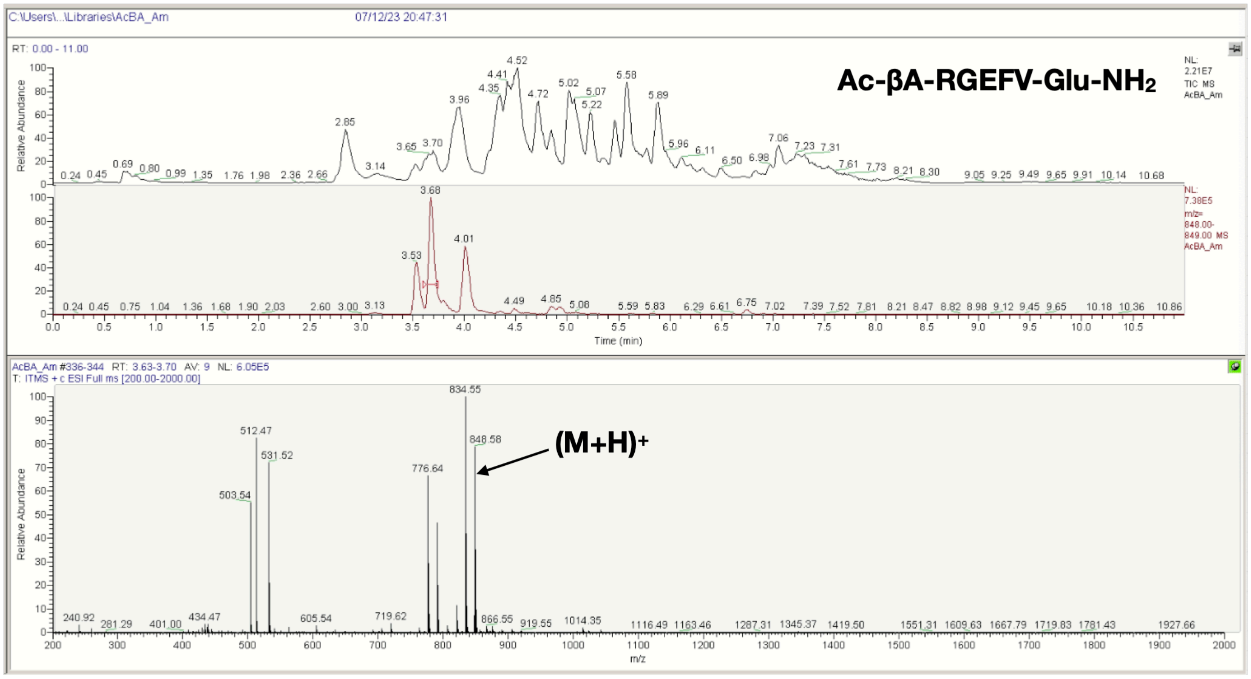
C:\Users\...\Libraries\AcBA_Am
07/12/23 20:47:31
RT: 0.00 - 11.00
Ac-βA-RGEFV-Glu-NH2
NL: 2.21E7 TIC MS AcBA_Am
NL: 7.38E5 m/z= 848.00-849.00 MS AcBA_Am
Relative Abundance
Time (min)
AcBA_Am #336-344 RT: 3.63-3.70 AV: 9 NL: 6.05E5
T: ITMS + c ESI Full ms [200.00-2000.00]
834.55
848.58
776.64
512.47
531.52
503.54
(M+H)+
m/z

**G**

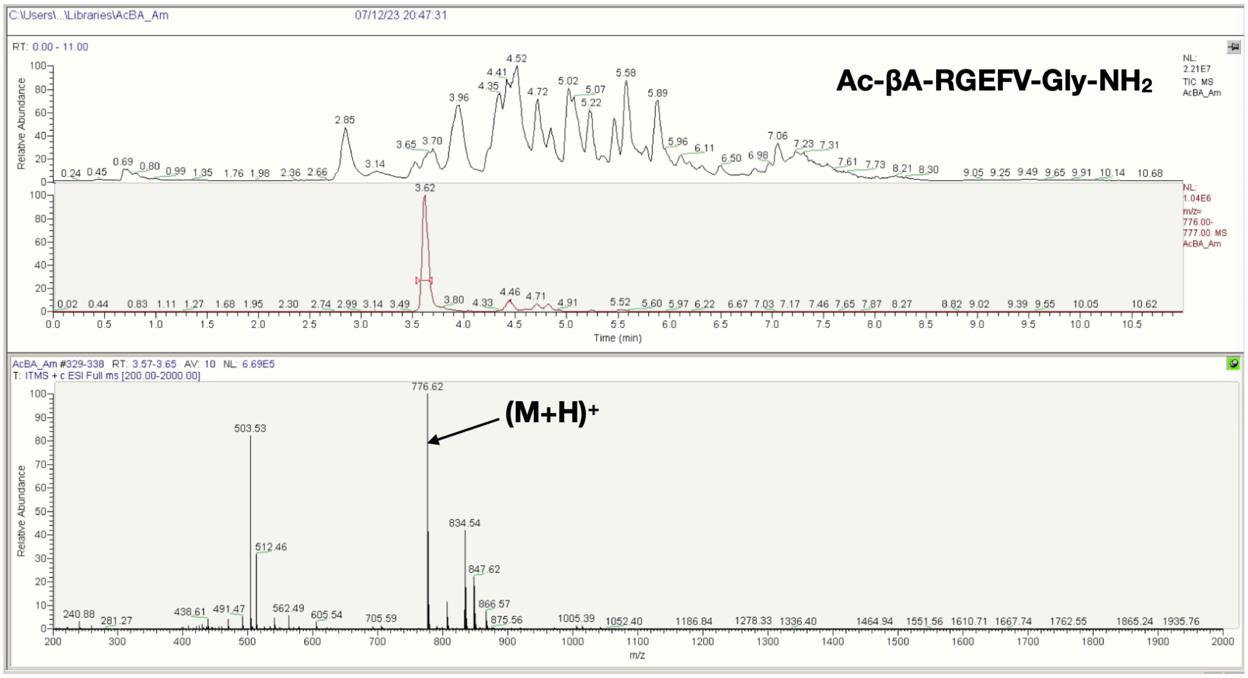
C:\Users\...\Libraries\AcBA_Am
07/12/23 20:47:31
RT: 0.00 - 11.00
Ac-βA-RGEFV-Gly-NH2
NL: 2.21E7 TIC MS AcBA_Am
NL: 1.04E6 m/z= 776.00-777.00 MS AcBA_Am
Relative Abundance
Time (min)
AcBA_Am #329-338 RT: 3.57-3.65 AV: 10 NL: 6.69E5
T: ITMS + c ESI Full ms [200.00-2000.00]
(M+H)+
m/z

**H**

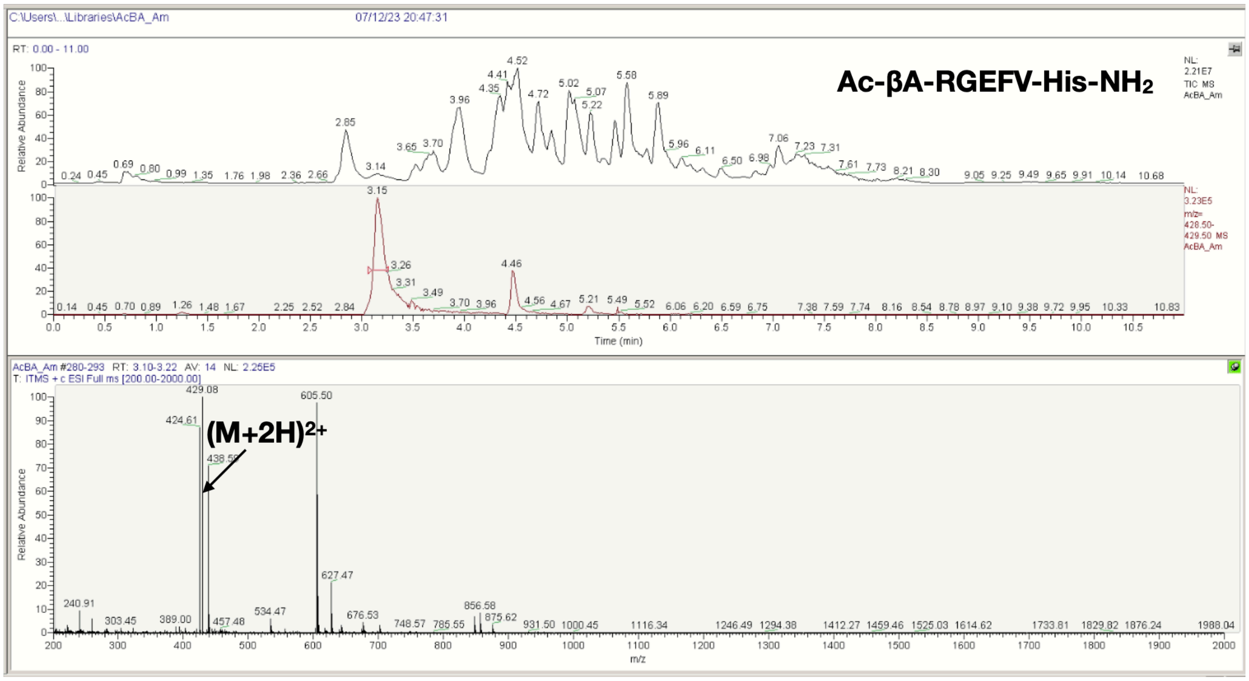
C:\Users\...\Libraries\AcBA_Am
07/12/23 20:47:31
RT: 0.00 - 11.00
Ac-βA-RGEFV-His-NH2
NL: 2.21E7 TIC MS AcBA_Am
NL: 3.23E5 m/z= 428.50-429.50 MS AcBA_Am
Relative Abundance
Time (min)
AcBA_Am #280-293 RT: 3.10-3.22 AV: 14 NL: 2.25E5
T: ITMS + c ESI Full ms [200.00-2000.00]
(M+2H)2+
m/z

**I**

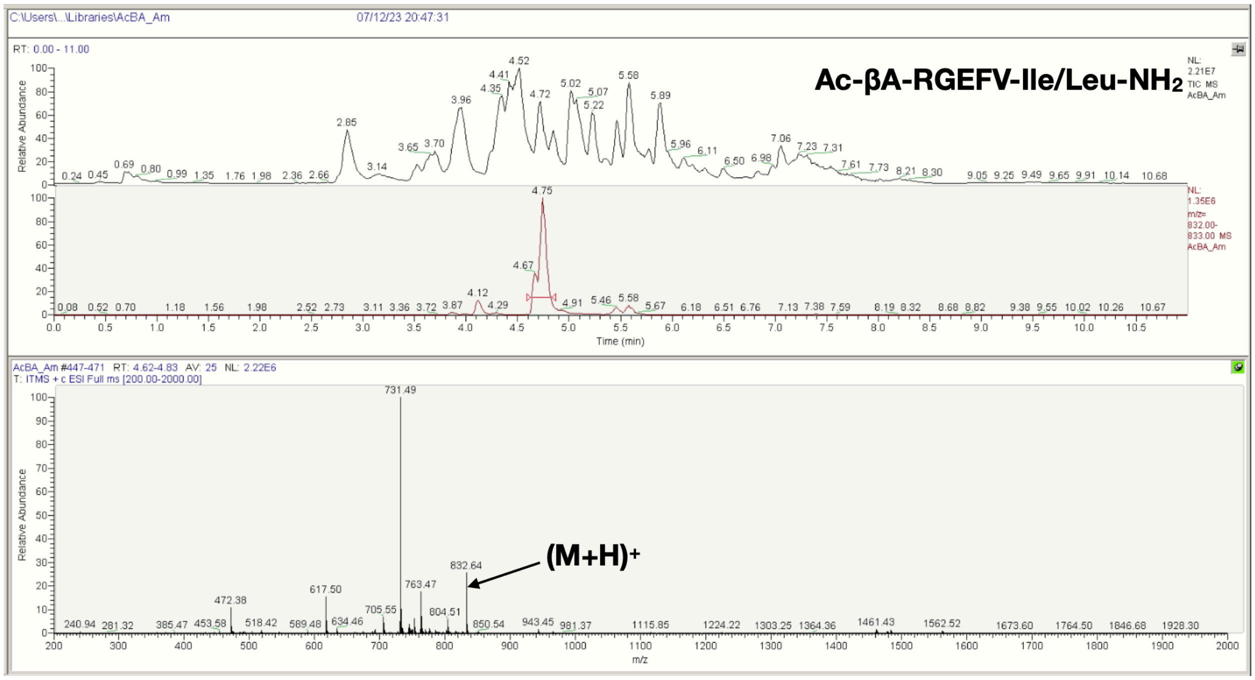
C:\Users\...\Libraries\AcBA_Am
07/12/23 20:47:31
RT: 0.00 - 11.00
Ac-βA-RGEFV-Ile/Leu-NH2
NL: 2.21E7 TIC MS AcBA_Am
NL: 1.35E6 m/z= 832.00-833.00 MS AcBA_Am
Relative Abundance
Time (min)
AcBA_Am #447-471 RT: 4.62-4.83 AV: 25 NL: 2.22E6
T: ITMS + c ESI Full ms [200.00-2000.00]
731.49
832.64
(M+H)+
m/z

**J**

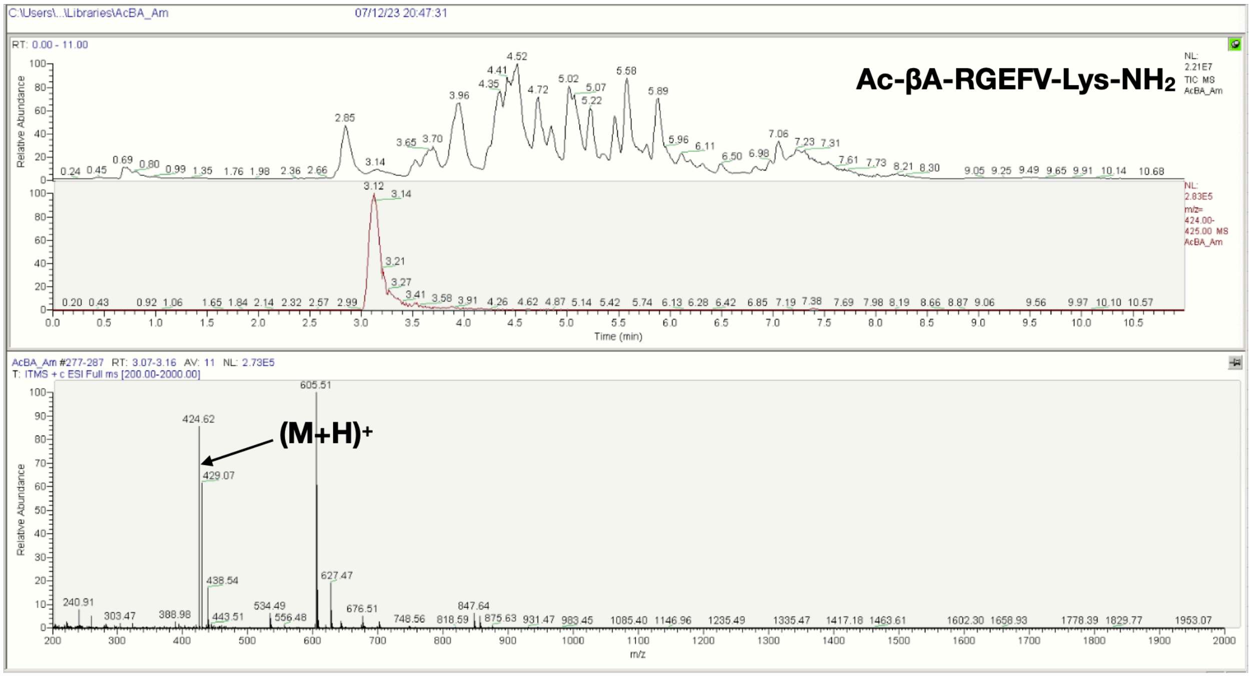
C:\Users\...\Libraries\AcBA_Am
07/12/23 20:47:31
RT: 0.00 - 11.00
Ac-βA-RGEFV-Lys-NH2
NL: 2.21E7 TIC MS AcBA_Am
NL: 2.83E5 m/z= 424.00-425.00 MS AcBA_Am
Relative Abundance
Time (min)
AcBA_Am #277-287 RT: 3.07-3.16 AV: 11 NL: 2.73E5
T: ITMS + c ESI Full ms [200.00-2000.00]
605.51
424.62
(M+H)+
m/z

**K**

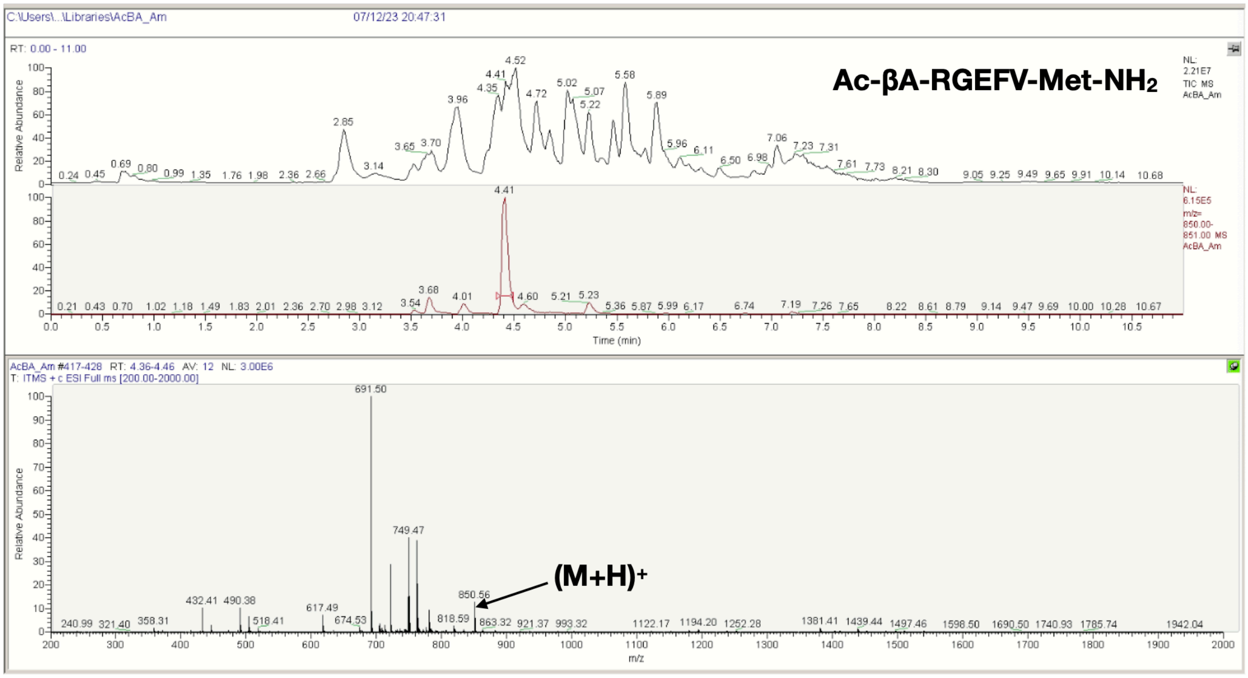


**L**

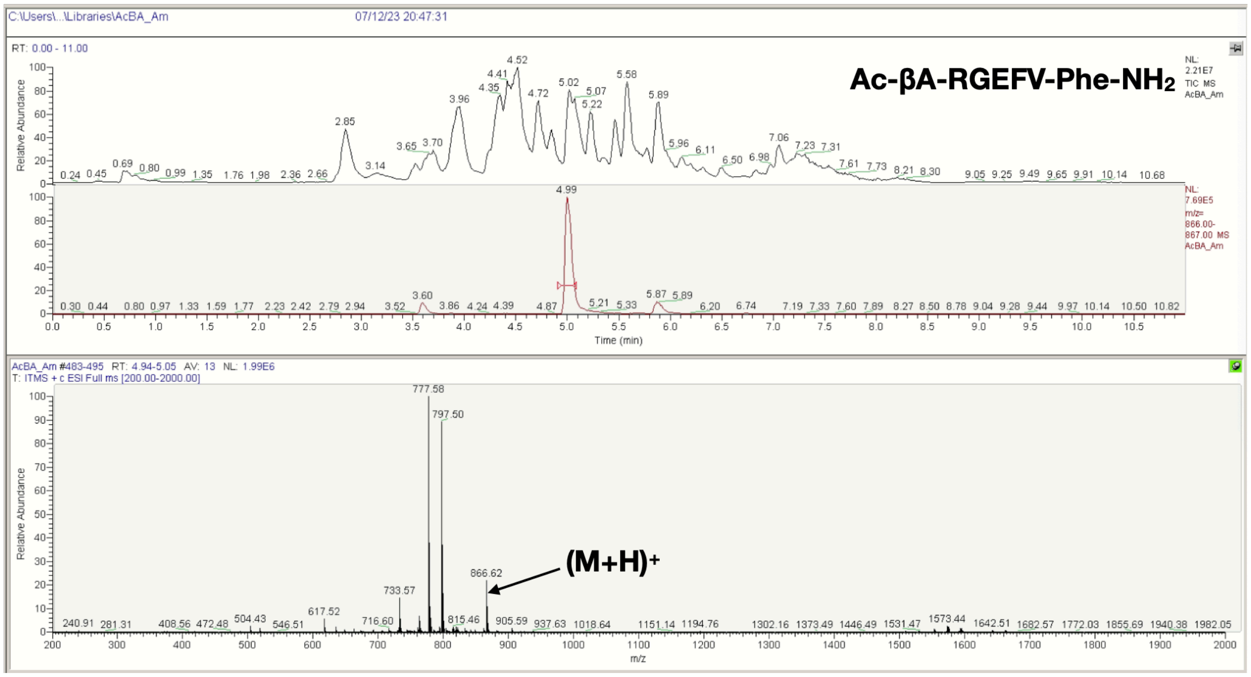

M

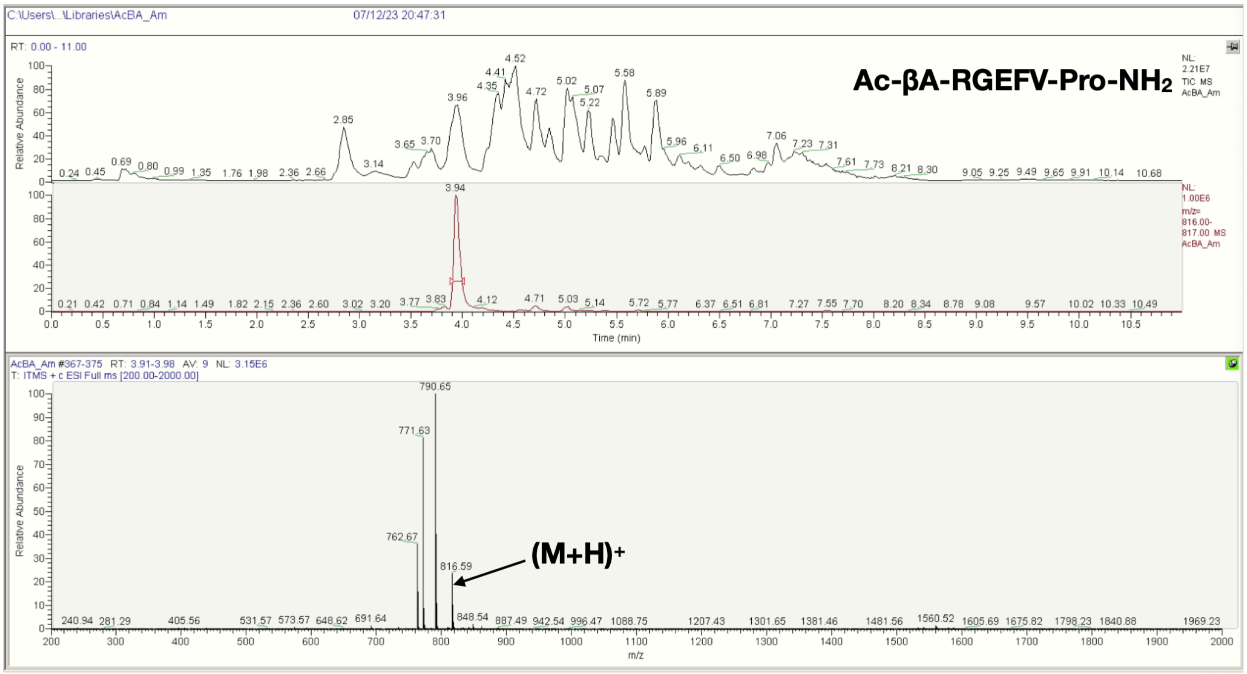
C:\Users\...\Libraries\AcBA_Am
07/12/23 20:47:31
RT: 0.00 - 11.00
Ac-βA-RGEFV-Pro-NH2
NL: 2.21E7 TIC MS AcBA_Am
NL: 1.00E6 m/z= 816.00-817.00 MS AcBA_Am
Relative Abundance
Time (min)
AcBA_Am #367-375 RT: 3.91-3.98 AV: 9 NL: 3.15E6
T: ITMS + c ESI Full ms [200.00-2000.00]
790.65
771.63
762.67
816.59
(M+H)+
m/z

N

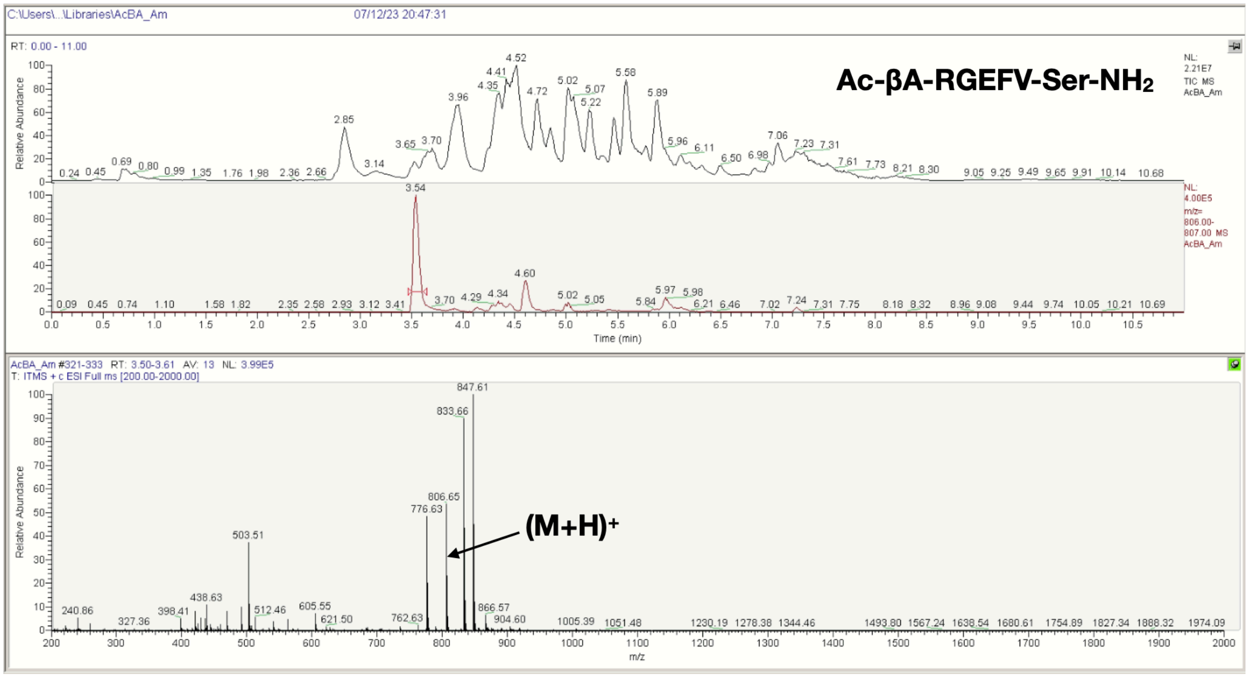
C:\Users\...\Libraries\AcBA_Am
07/12/23 20:47:31
RT: 0.00 - 11.00
Ac-βA-RGEFV-Ser-NH2
NL: 2.21E7 TIC MS AcBA_Am
NL: 4.00E5 m/z= 806.00-807.00 MS AcBA_Am
Relative Abundance
Time (min)
AcBA_Am #321-333 RT: 3.50-3.61 AV: 13 NL: 3.99E5
T: ITMS + c ESI Full ms [200.00-2000.00]
847.61
833.66
806.65
776.63
503.51
(M+H)+
m/z

**O**

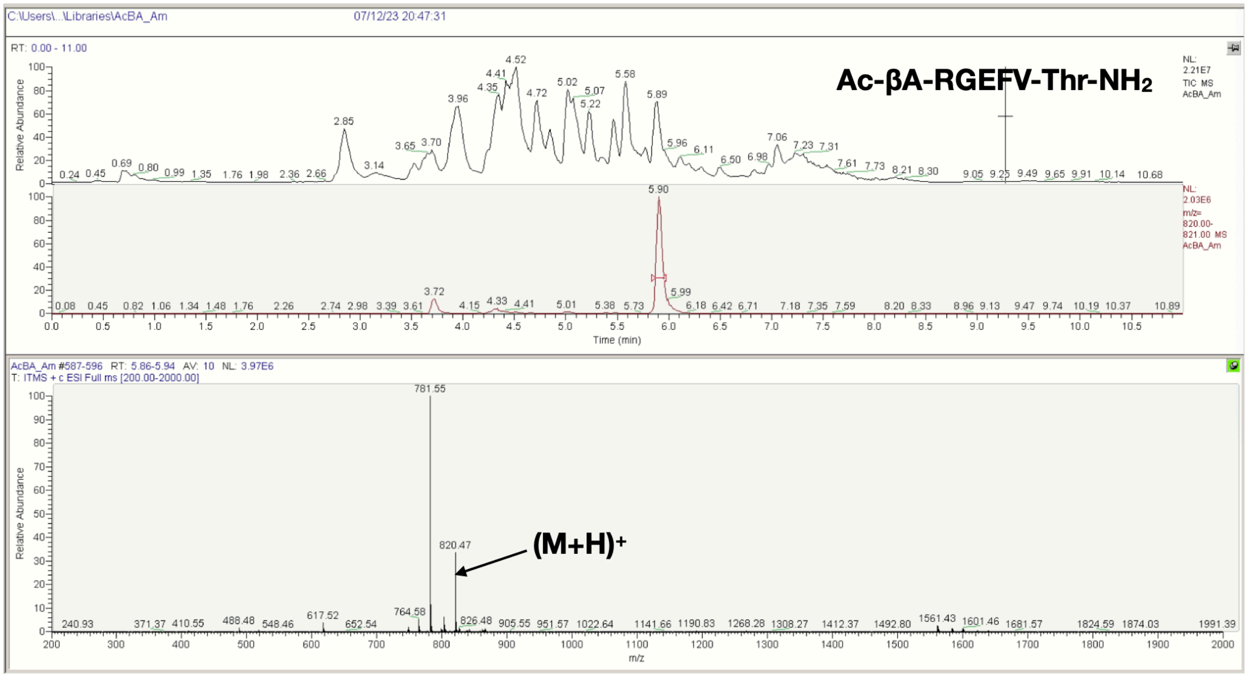
Ac-βA-RGEFV-Thr-NH2
(M+H)+
820.47
781.55

**P**

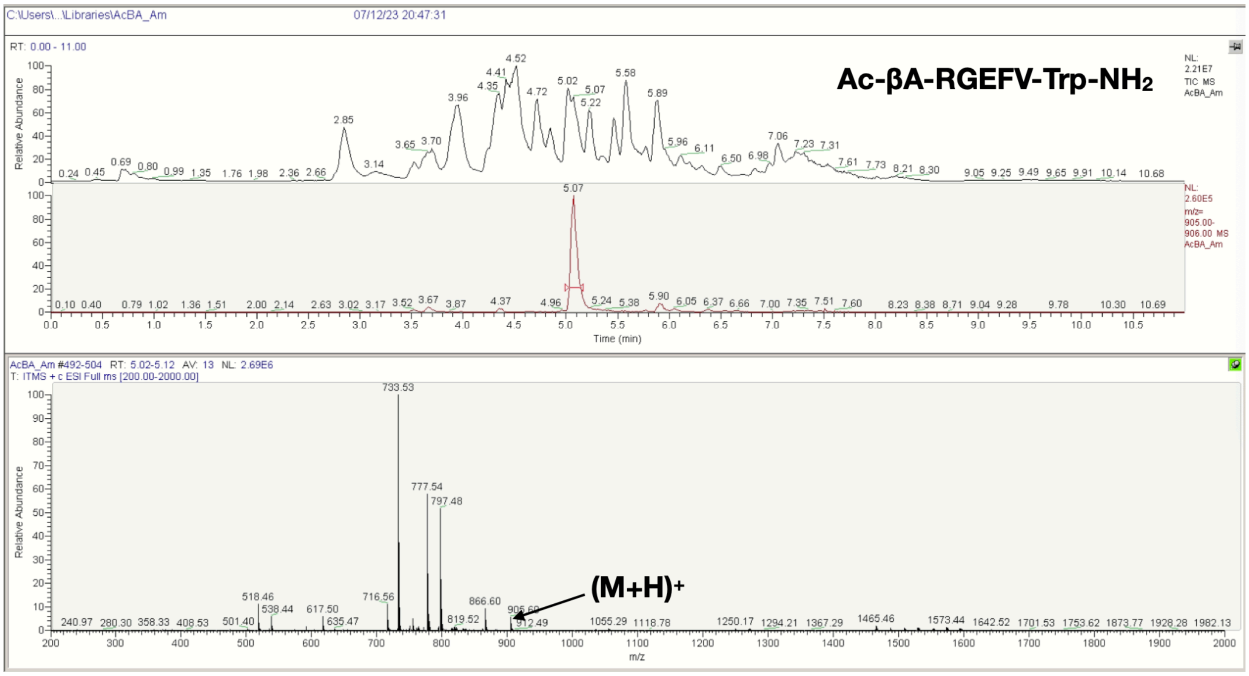
Ac-βA-RGEFV-Trp-NH2
(M+H)+
733.53
777.54
797.48

**Q**

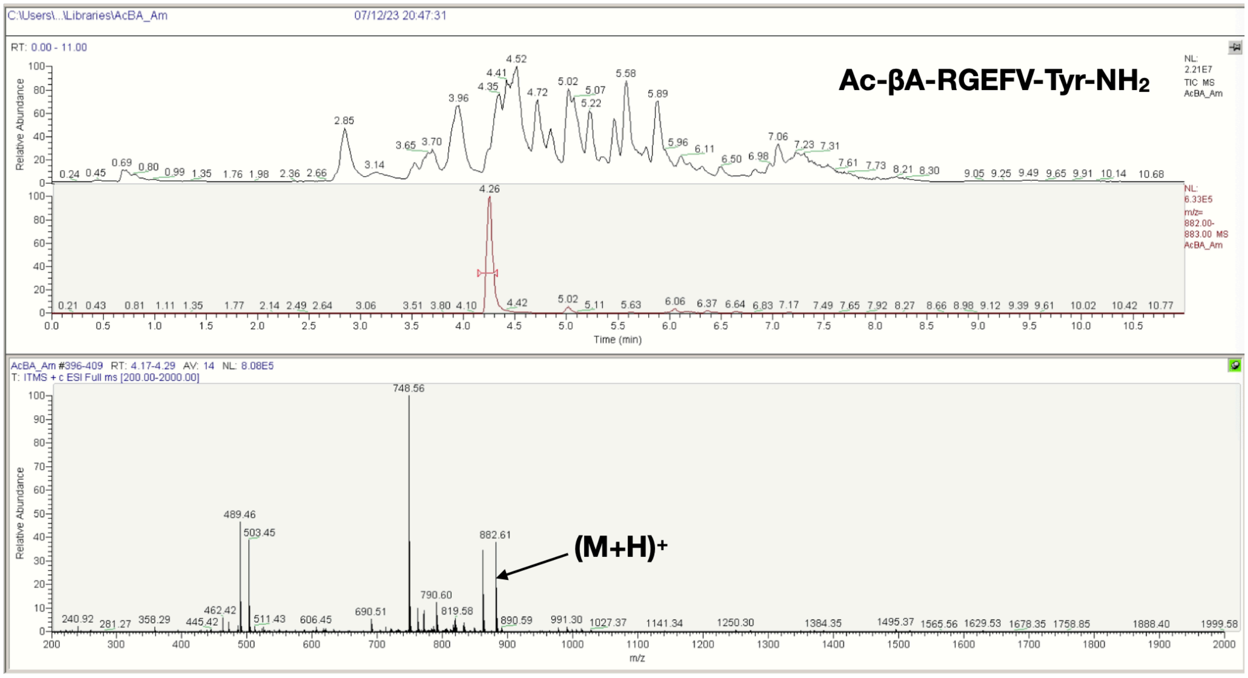


**R**

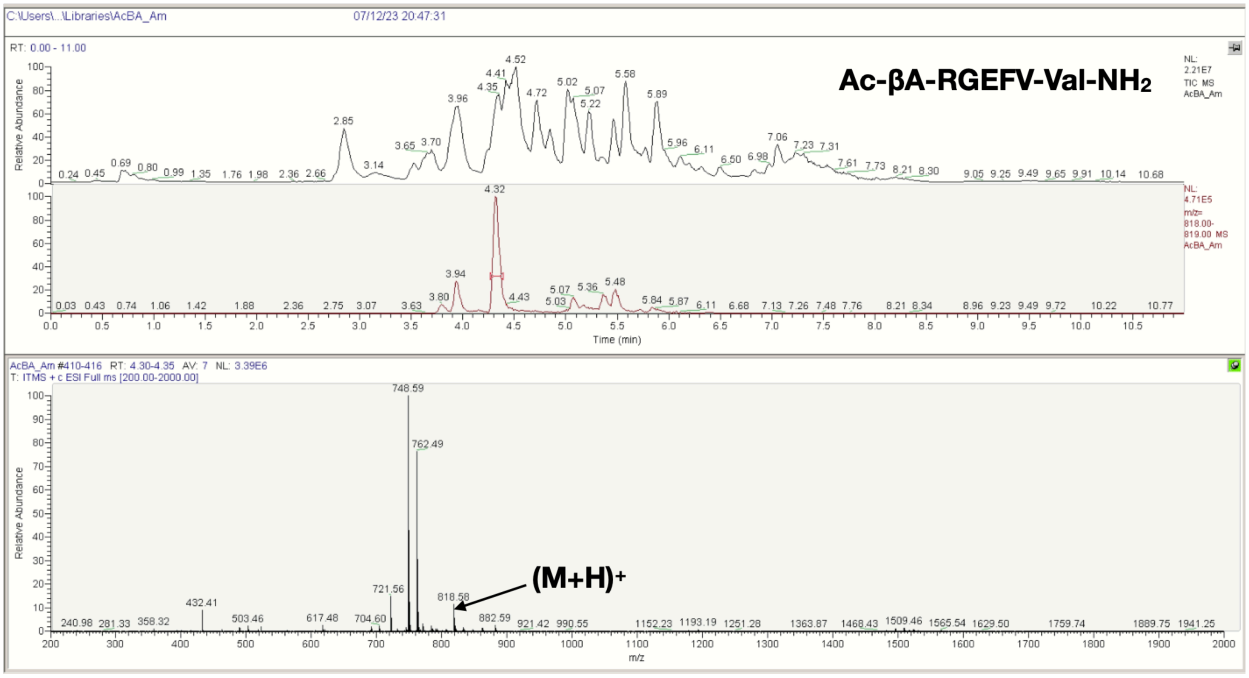


**Figure S14**. LCMS spectra of the Ac-βA-RGEFV-X-$NH_2$ libraries, where X = a) Ala, b) Arg, c) Asn, d) Asp, e) Gln, f) Glu, g) Gly, h) His, i) Ile/Leu, j) Lys, k) Met, l) Phe, m) Pro, n) Ser, o) Thr, p) Trp, q) Tyr, r) Val.

A

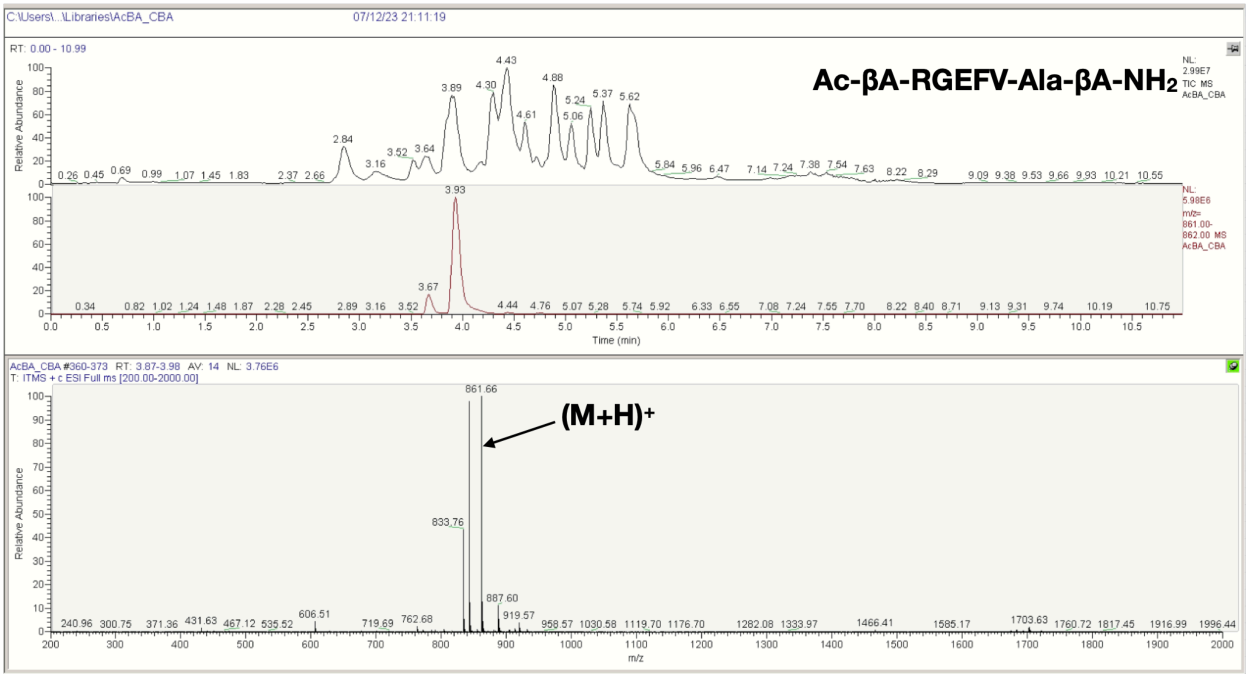

C:\Users\...\Libraries\AcBA_CBA
07/12/23 21:11:19
RT: 0.00 - 10.99
Ac-βA-RGEFV-Ala-βA-NH2
NL: 2.99E7 TIC MS AcBA_CBA
NL: 5.98E6 m/z= 861.00-862.00 MS AcBA_CBA
Relative Abundance
Time (min)
AcBA_CBA #360-373 RT: 3.87-3.98 AV: 14 NL: 3.76E6
T: ITMS + c ESI Full ms [200.00-2000.00]
861.66
(M+H)+
833.76
887.60
m/z


B

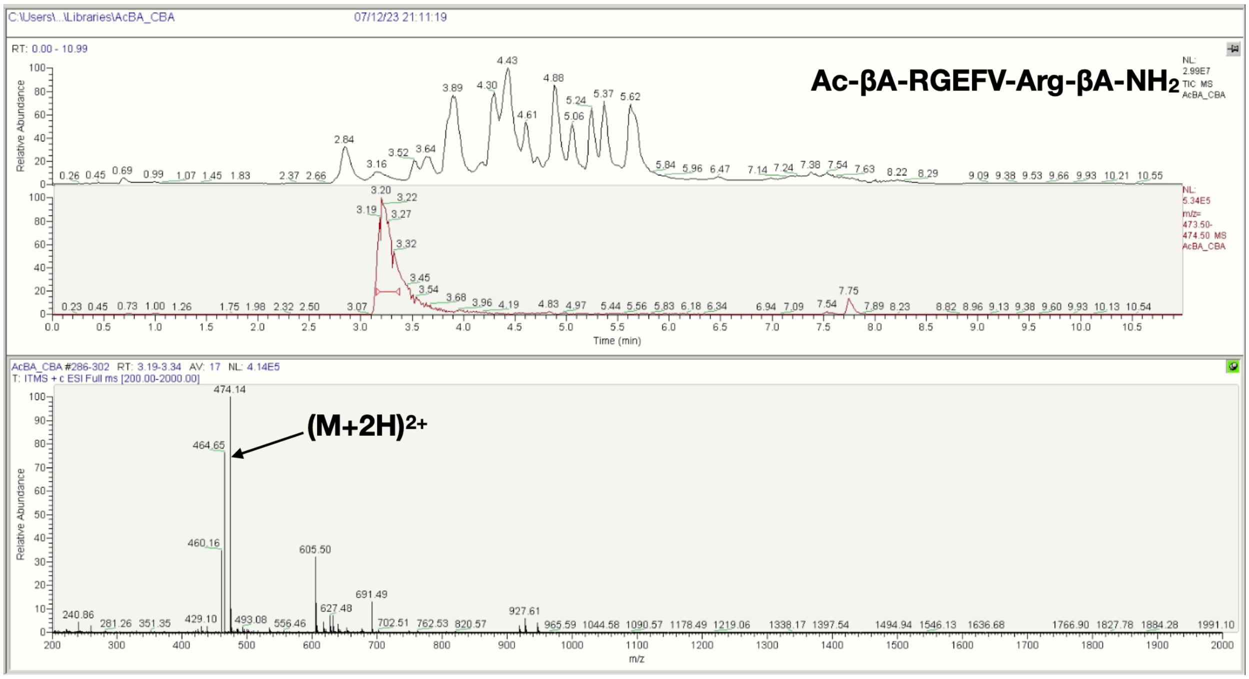

C:\Users\...\Libraries\AcBA_CBA
07/12/23 21:11:19
RT: 0.00 - 10.99
Ac-βA-RGEFV-Arg-βA-NH2
NL: 2.99E7 TIC MS AcBA_CBA
NL: 5.34E5 m/z= 473.50-474.50 MS AcBA_CBA
Relative Abundance
Time (min)
AcBA_CBA #286-302 RT: 3.19-3.34 AV: 17 NL: 4.14E5
T: ITMS + c ESI Full ms [200.00-2000.00]
474.14
(M+2H)2+
464.65
460.16
605.50
691.49
m/z

**C**

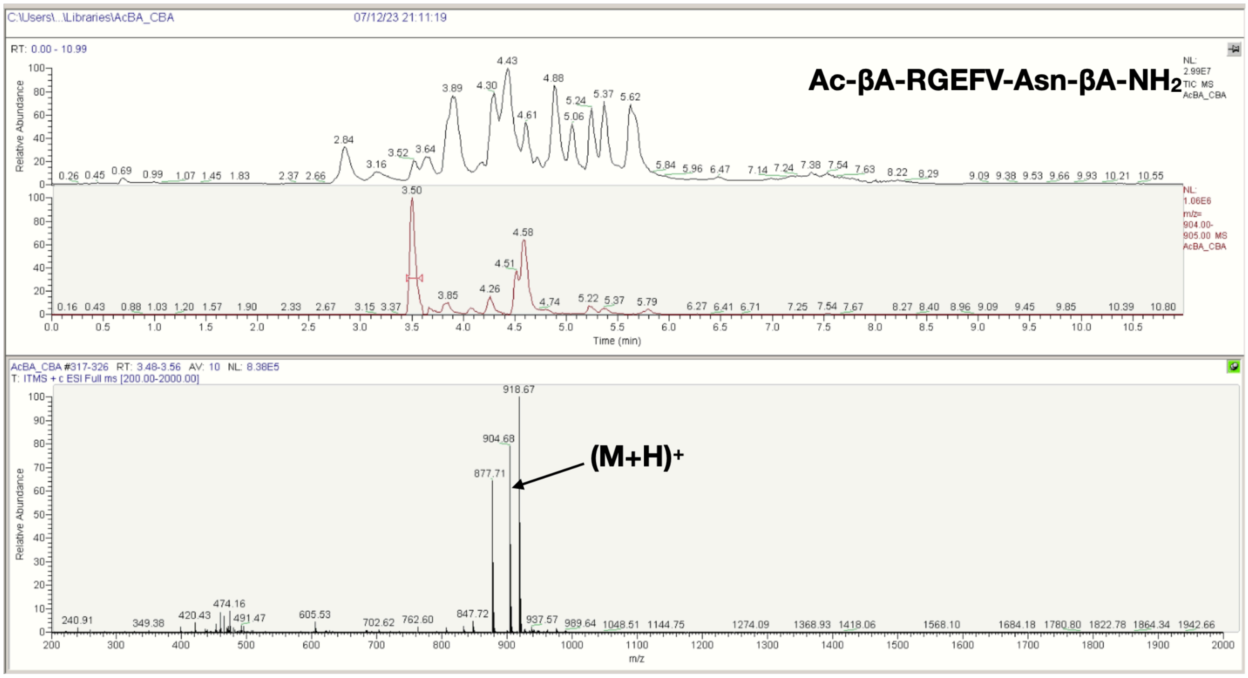
C:\Users\..\Libraries\AcBA_CBA
07/12/23 21:11:19
RT: 0.00 - 10.99
Ac-βA-RGEFV-Asn-βA-NH2
NL: 2.99E7 TIC MS AcBA_CBA
NL: 1.06E6 m/z= 904.00-905.00 MS AcBA_CBA
Relative Abundance
Time (min)
AcBA_CBA #317-326 RT: 3.48-3.56 AV: 10 NL: 8.38E5
T: ITMS + c ESI Full ms [200.00-2000.00]
918.67
904.68
877.71
(M+H)+
m/z

**D**

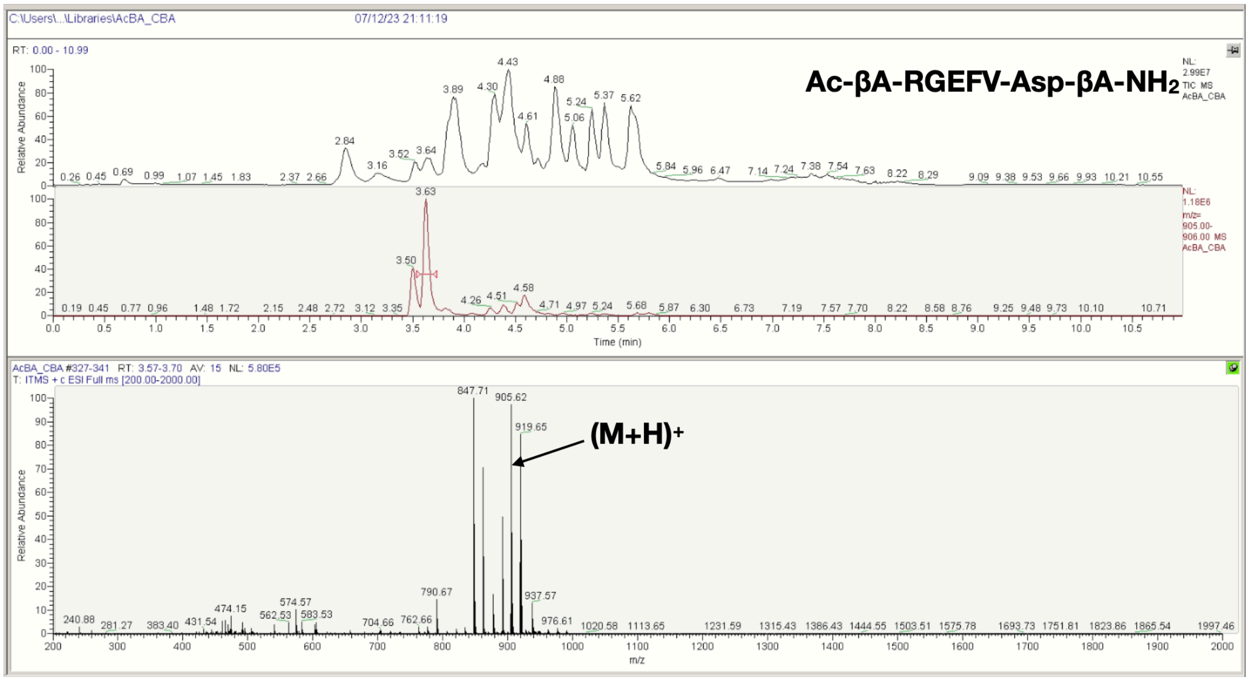
C:\Users\..\Libraries\AcBA_CBA
07/12/23 21:11:19
RT: 0.00 - 10.99
Ac-βA-RGEFV-Asp-βA-NH2
NL: 2.99E7 TIC MS AcBA_CBA
NL: 1.18E6 m/z= 905.00-906.00 MS AcBA_CBA
Relative Abundance
Time (min)
AcBA_CBA #327-341 RT: 3.57-3.70 AV: 15 NL: 5.80E5
T: ITMS + c ESI Full ms [200.00-2000.00]
847.71
905.62
919.65
(M+H)+
m/z

E

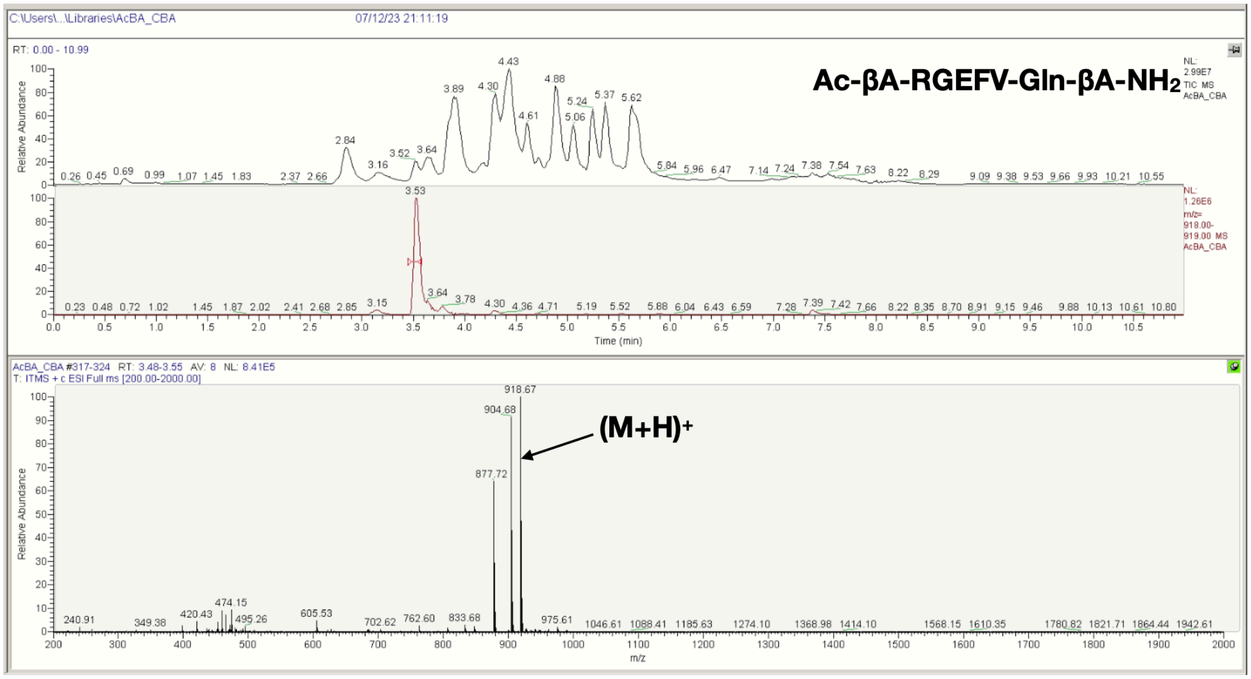
C:\Users\...\Libraries\AcBA_CBA
07/12/23 21:11:19
RT: 0.00 - 10.99
Ac-βA-RGEFV-Gln-βA-NH2
NL: 2.99E7 TIC MS AcBA_CBA
NL: 1.26E6 m/z= 918.00-919.00 MS AcBA_CBA
Relative Abundance
Time (min)
AcBA_CBA #317-324 RT: 3.48-3.55 AV: 8 NL: 8.41E5
T: ITMS + c ESI Full ms [200.00-2000.00]
918.67
904.68
877.72
(M+H)+
m/z

F

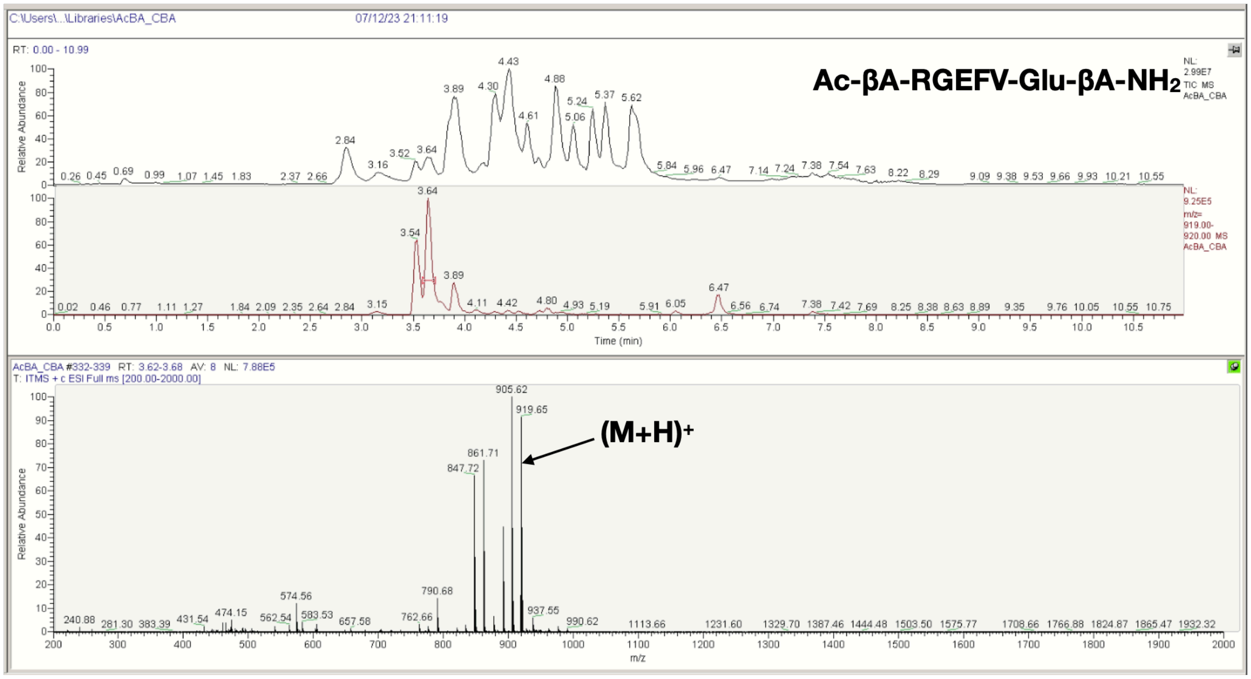
C:\Users\...\Libraries\AcBA_CBA
07/12/23 21:11:19
RT: 0.00 - 10.99
Ac-βA-RGEFV-Glu-βA-NH2
NL: 2.99E7 TIC MS AcBA_CBA
NL: 9.25E5 m/z= 919.00-920.00 MS AcBA_CBA
Relative Abundance
Time (min)
AcBA_CBA #332-339 RT: 3.62-3.68 AV: 8 NL: 7.88E5
T: ITMS + c ESI Full ms [200.00-2000.00]
905.62
919.65
861.71
847.72
(M+H)+
m/z

G

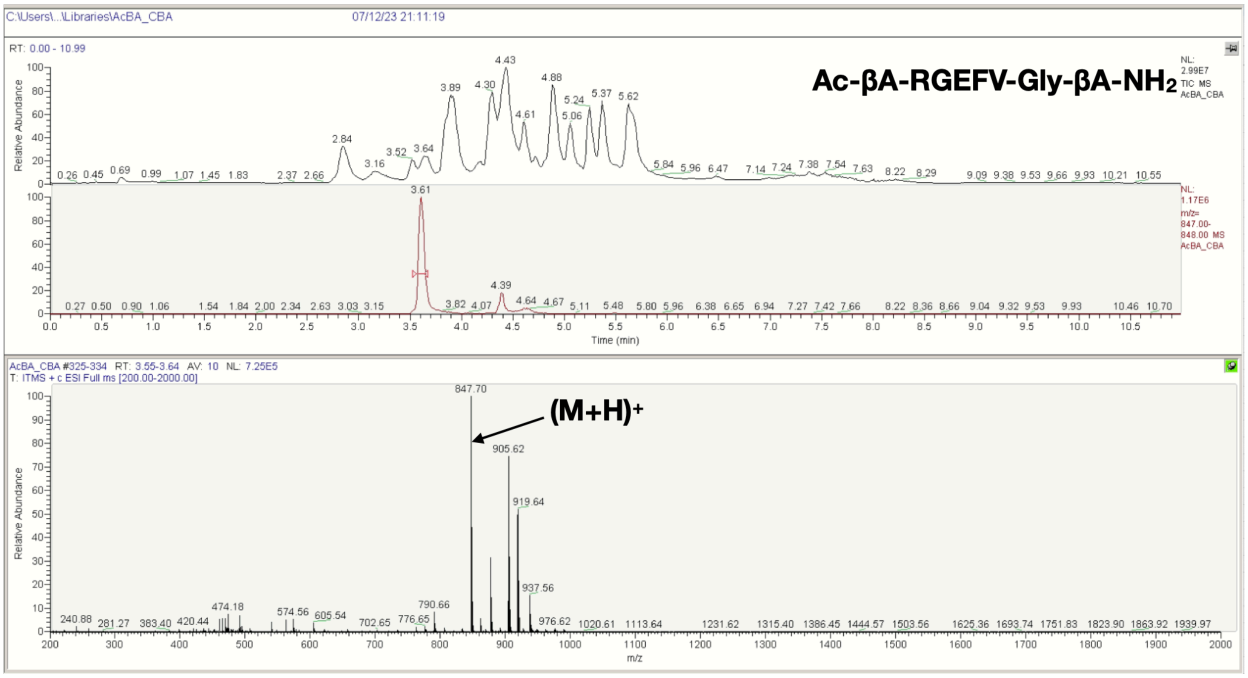
Ac-βA-RGEFV-Gly-βA-NH2
(M+H)+
847.70
905.62
919.64
Time (min)
m/z
Relative Abundance

H

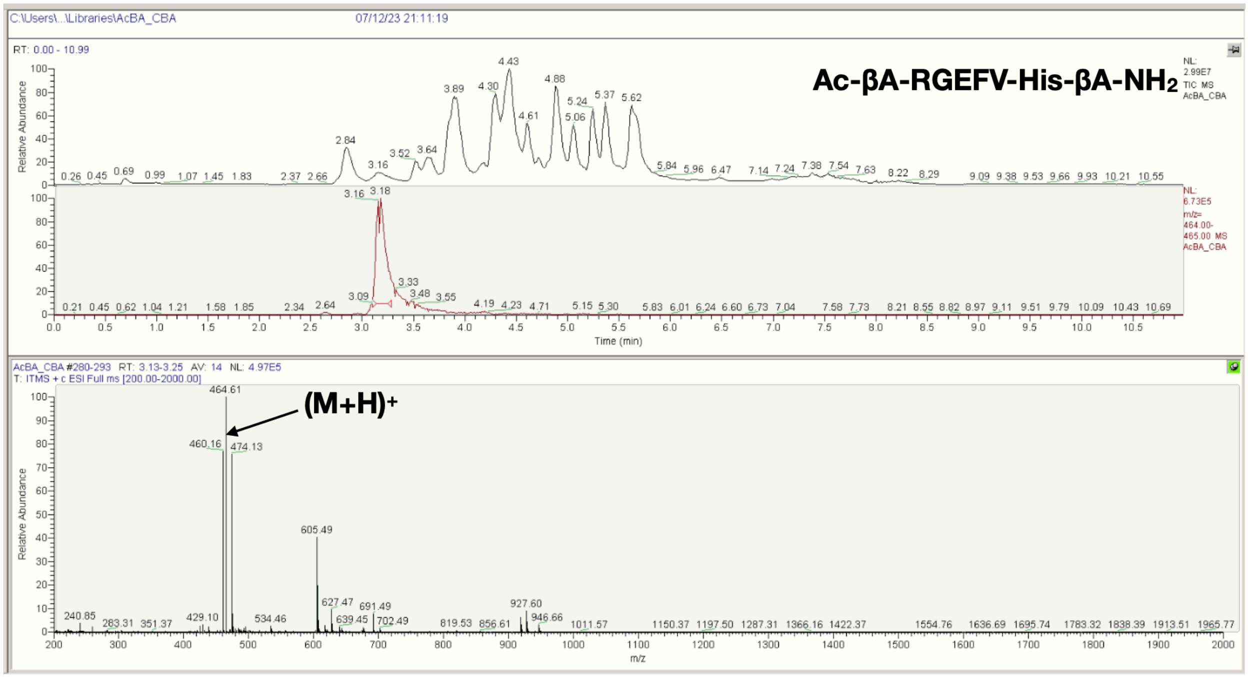
Ac-βA-RGEFV-His-βA-NH2
(M+H)+
464.61
460.16
474.13
605.49
Time (min)
m/z
Relative Abundance

I

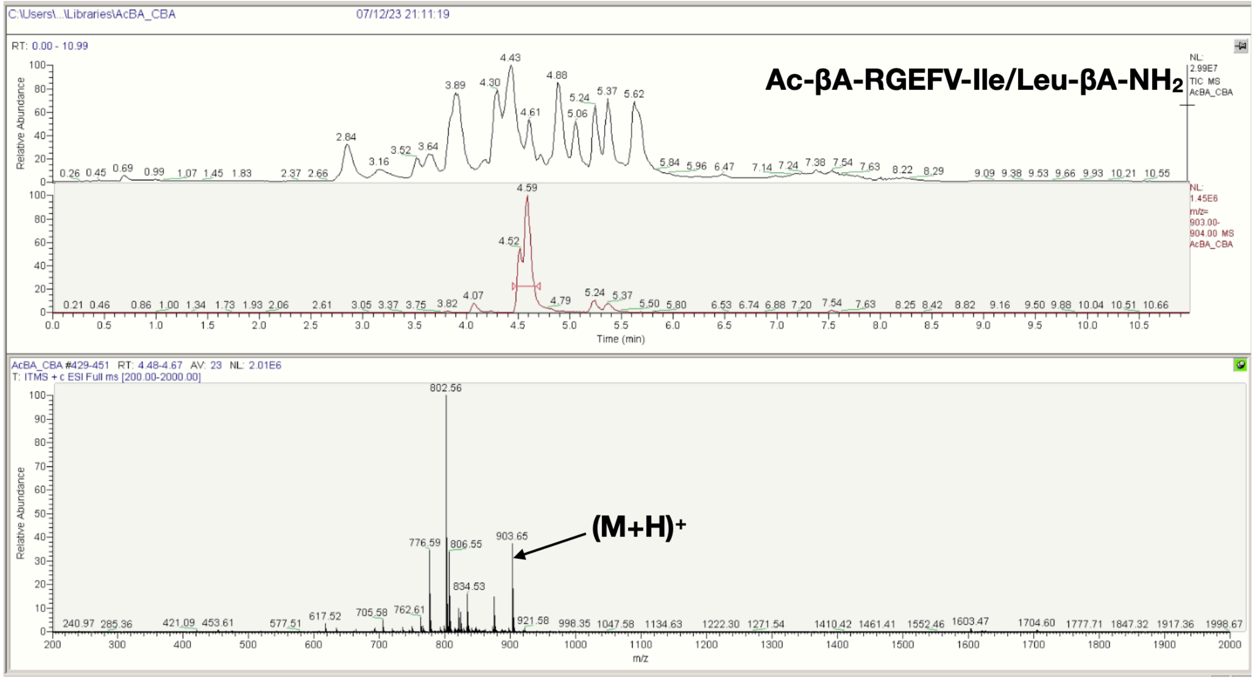
Ac-βA-RGEFV-Ile/Leu-βA-NH2
(M+H)+
802.56
903.65
776.59
806.55
834.53
RT: 0.00 - 10.99
Time (min)
m/z
Relative Abundance

J

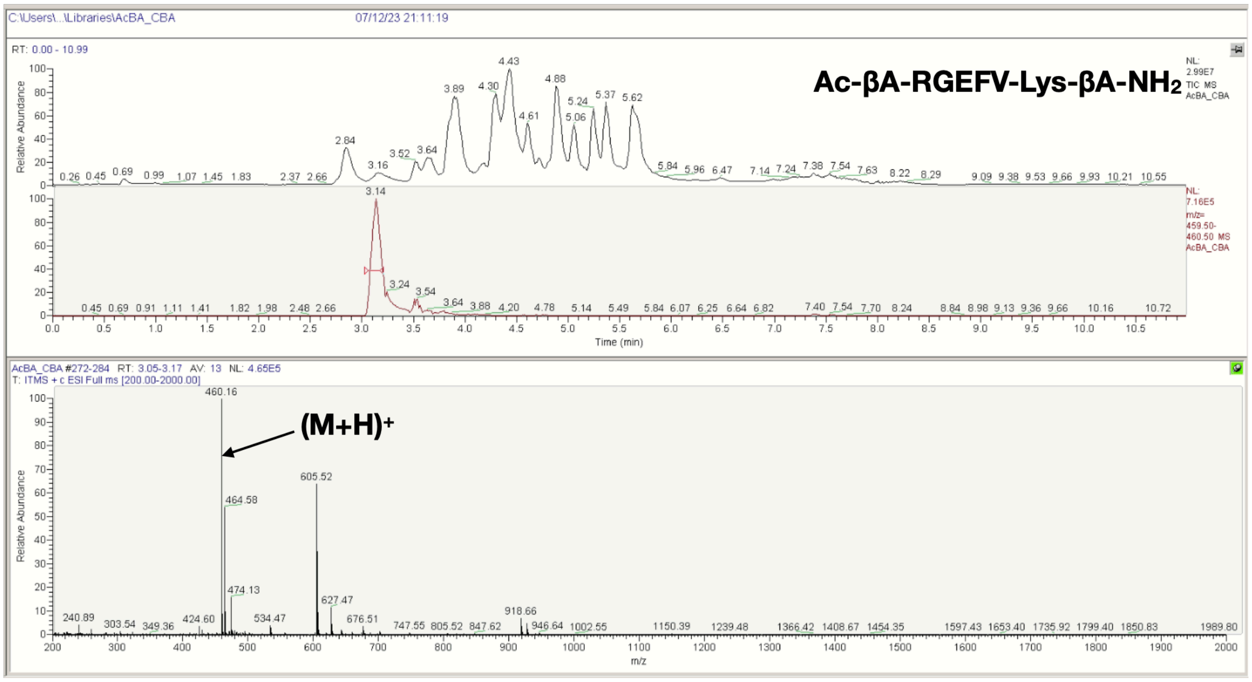
Ac-βA-RGEFV-Lys-βA-NH2
(M+H)+
460.16
605.52
464.58
474.13
627.47
RT: 0.00 - 10.99
Time (min)
m/z
Relative Abundance

**K**

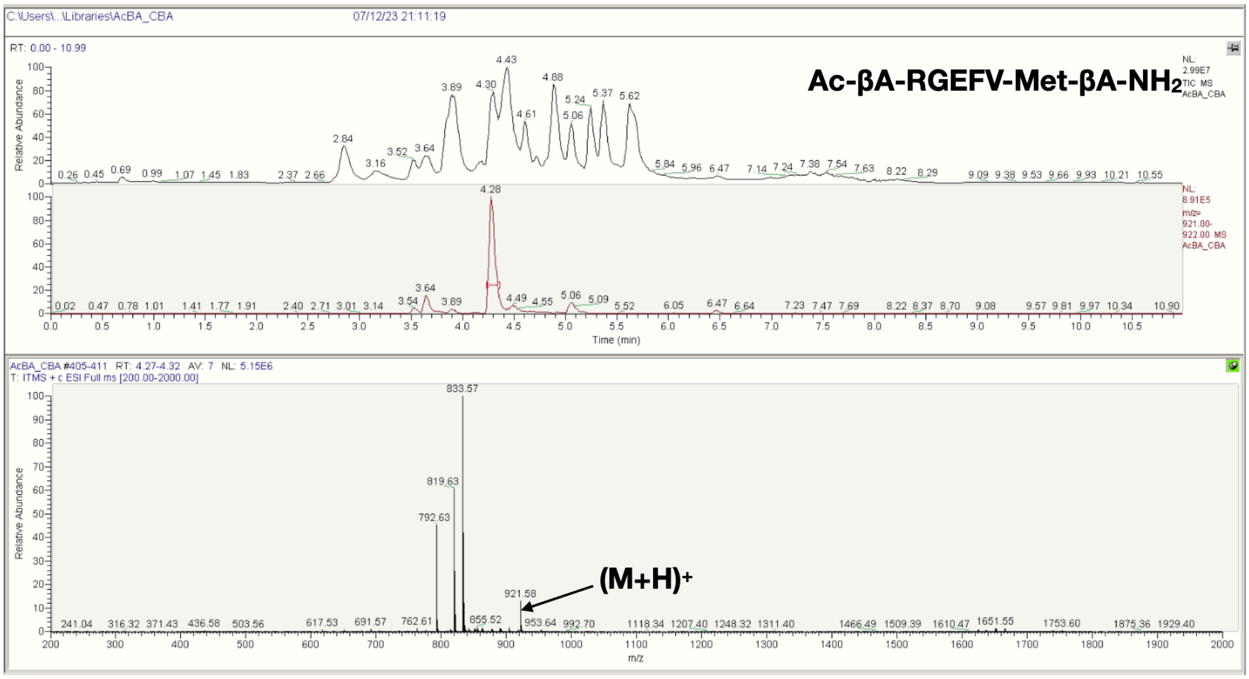
Ac-βA-RGEFV-Met-βA-NH2
(M+H)+
921.58
833.57
819.63
792.63

**L**

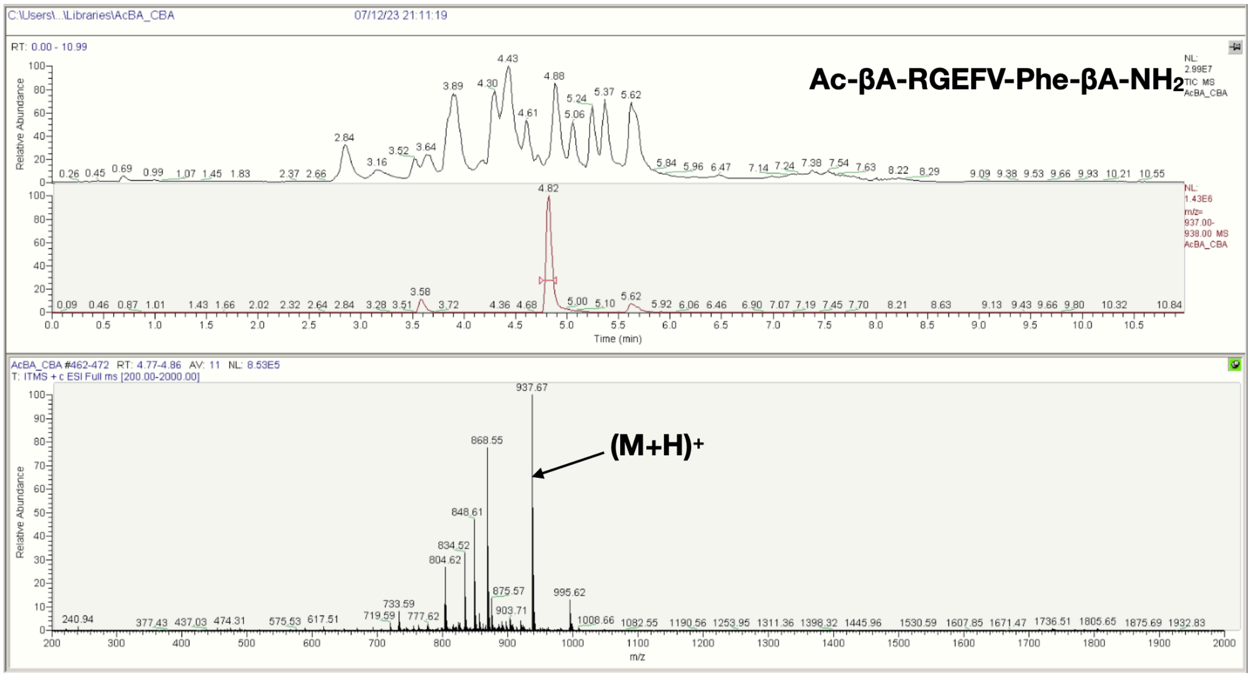
Ac-βA-RGEFV-Phe-βA-NH2
(M+H)+
937.67
868.55
848.61

**M**

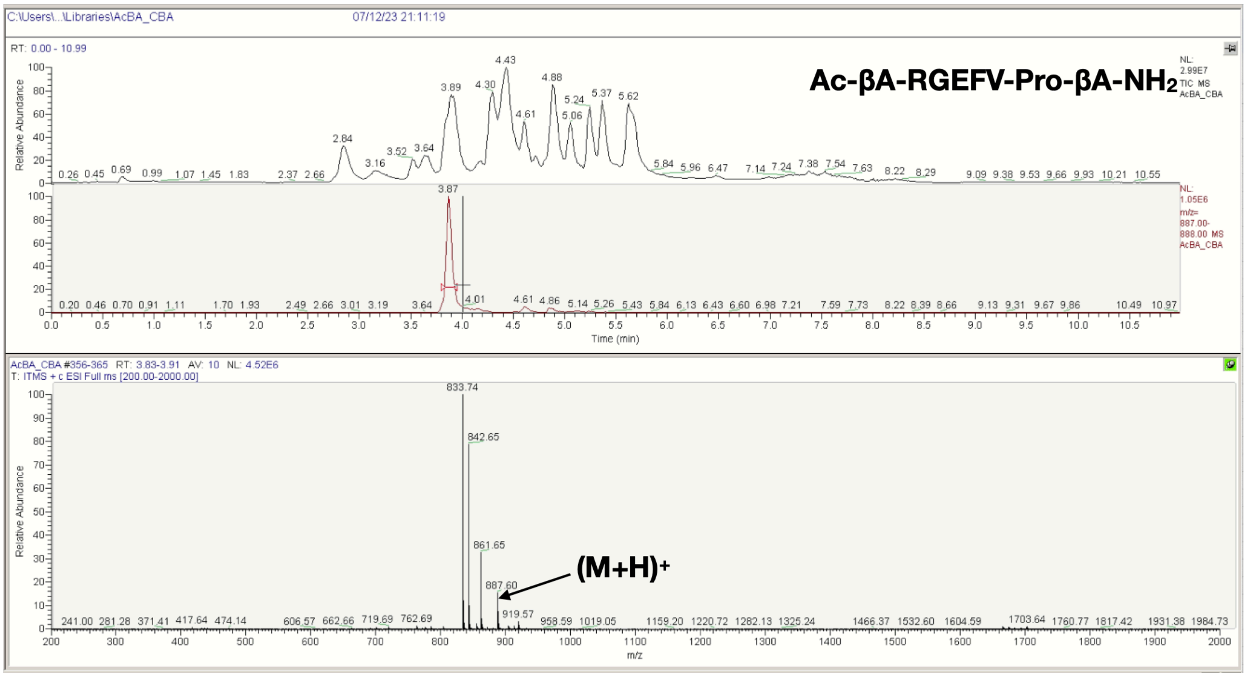
C:\Users\...\Libraries\AcBA_CBA
07/12/23 21:11:19
RT: 0.00 - 10.99
Ac-βA-RGEFV-Pro-βA-NH2
NL: 2.99E7 TIC MS AcBA_CBA
NL: 1.05E6 m/z= 887.00-888.00 MS AcBA_CBA
Time (min)
Relative Abundance
AcBA_CBA #356-365 RT: 3.83-3.91 AV: 10 NL: 4.52E6
T: ITMS + c ESI Full ms [200.00-2000.00]
833.74
842.65
861.65
887.60
919.57
(M+H)+
m/z

**N**

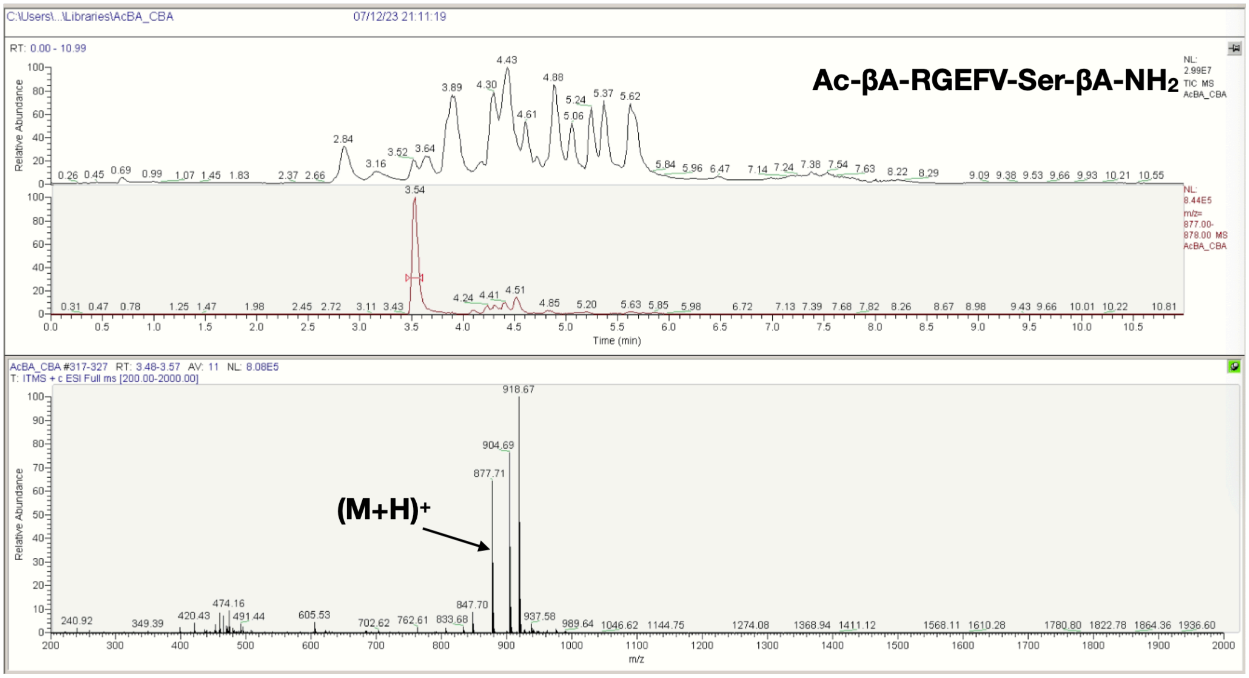
C:\Users\...\Libraries\AcBA_CBA
07/12/23 21:11:19
RT: 0.00 - 10.99
Ac-βA-RGEFV-Ser-βA-NH2
NL: 2.99E7 TIC MS AcBA_CBA
NL: 8.44E5 m/z= 877.00-878.00 MS AcBA_CBA
Time (min)
Relative Abundance
AcBA_CBA #317-327 RT: 3.48-3.57 AV: 11 NL: 8.08E5
T: ITMS + c ESI Full ms [200.00-2000.00]
918.67
904.69
877.71
(M+H)+
m/z

**O**

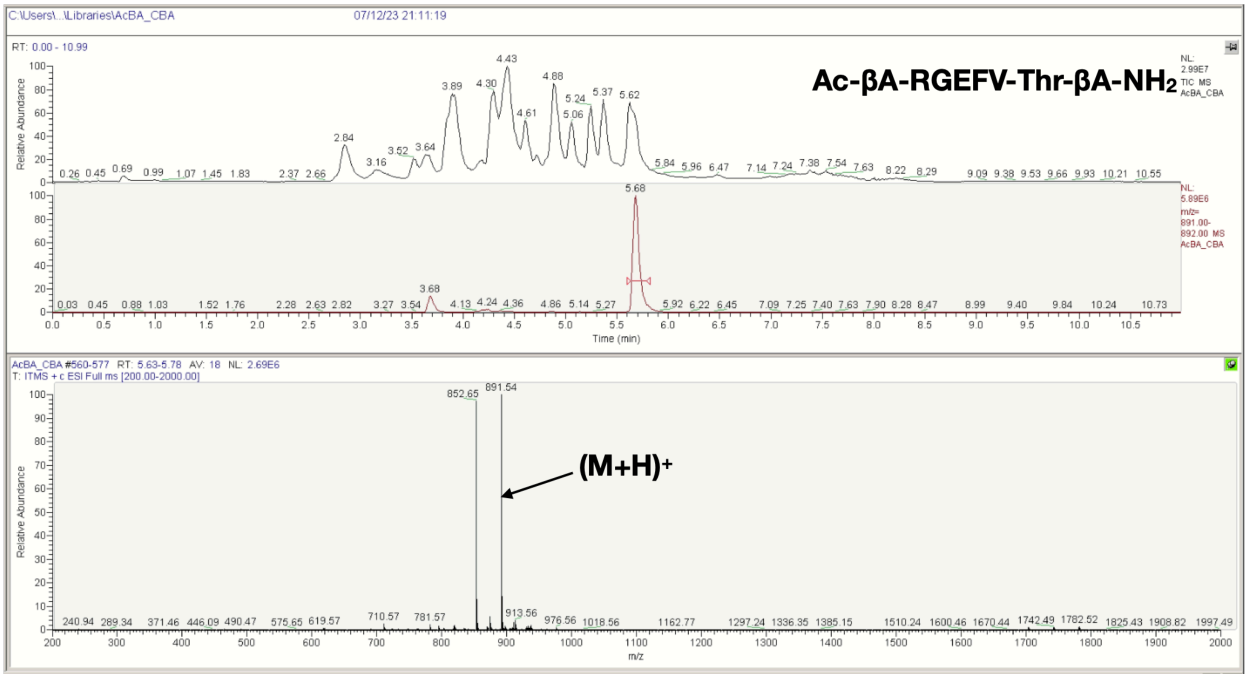
C:\Users\...\Libraries\AcBA_CBA
07/12/23 21:11:19
RT: 0.00 - 10.99
Ac-βA-RGEFV-Thr-βA-NH2
NL: 2.99E7 TIC MS AcBA_CBA
NL: 5.89E6 m/z= 891.00-892.00 MS AcBA_CBA
Relative Abundance
Time (min)
AcBA_CBA #560-577 RT: 5.63-5.78 AV: 18 NL: 2.69E6
T: ITMS + c ESI Full ms [200.00-2000.00]
852.65
891.54
(M+H)+
m/z

**P**

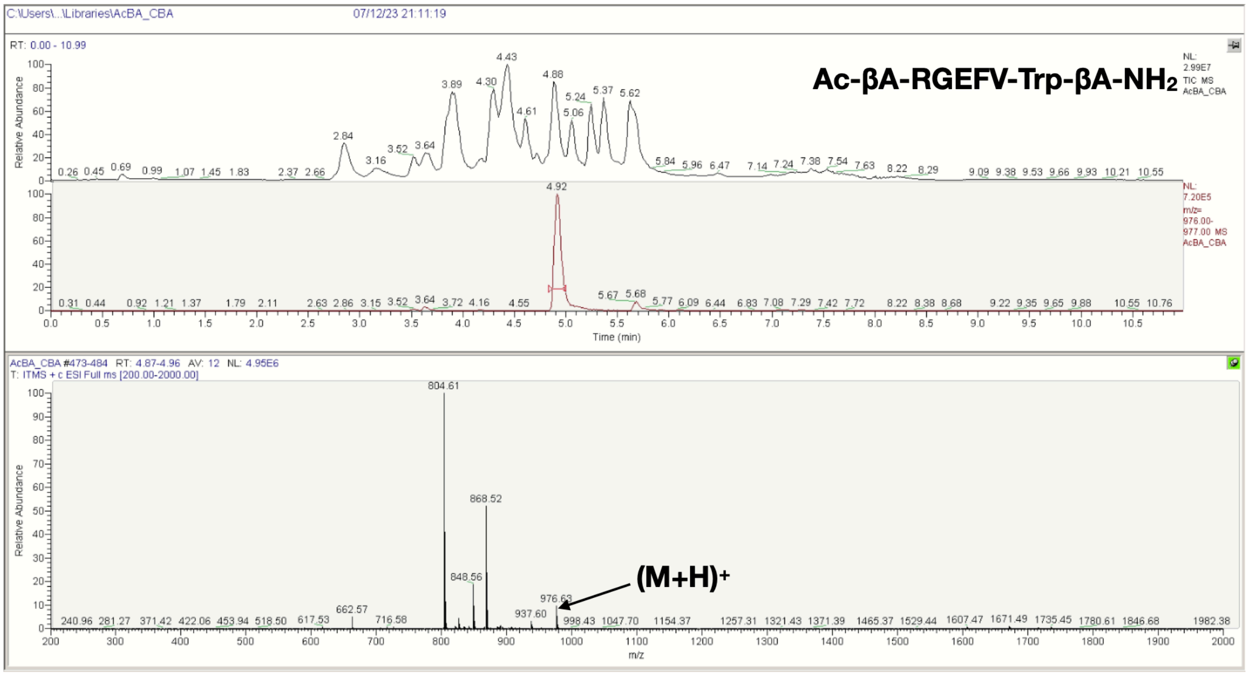
C:\Users\...\Libraries\AcBA_CBA
07/12/23 21:11:19
RT: 0.00 - 10.99
Ac-βA-RGEFV-Trp-βA-NH2
NL: 2.99E7 TIC MS AcBA_CBA
NL: 7.20E5 m/z= 976.00-977.00 MS AcBA_CBA
Relative Abundance
Time (min)
AcBA_CBA #473-484 RT: 4.87-4.96 AV: 12 NL: 4.95E6
T: ITMS + c ESI Full ms [200.00-2000.00]
804.61
868.52
848.56
976.63
(M+H)+
m/z

**Q**

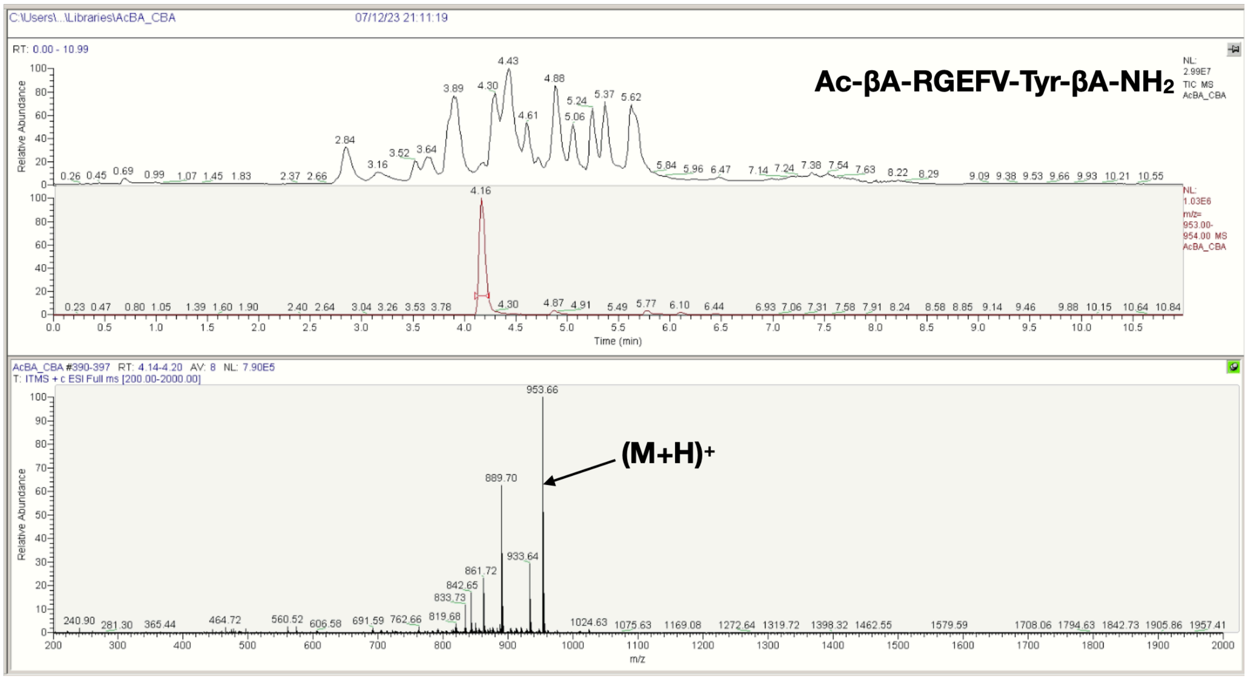


**R**

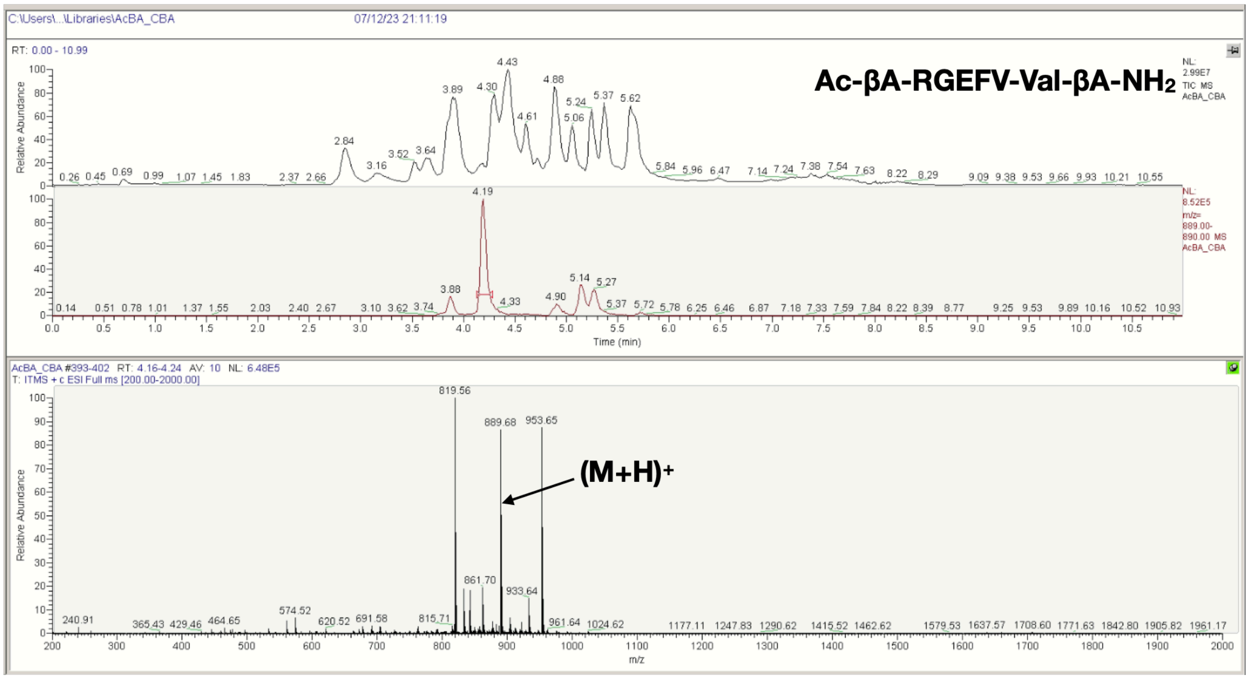


**Figure S15**. LCMS spectra of the Ac-βA-RGEFV-X-βA-$NH_2$ libraries, where X = a) Ala, b) Arg, c) Asn, d) Asp, e) Gln, f) Glu, g) Gly, h) His, i) Ile/Leu, j) Lys, k) Met, l) Phe, m) Pro, n) Ser, o) Thr, p) Trp, q) Tyr, r) Val.

**A**

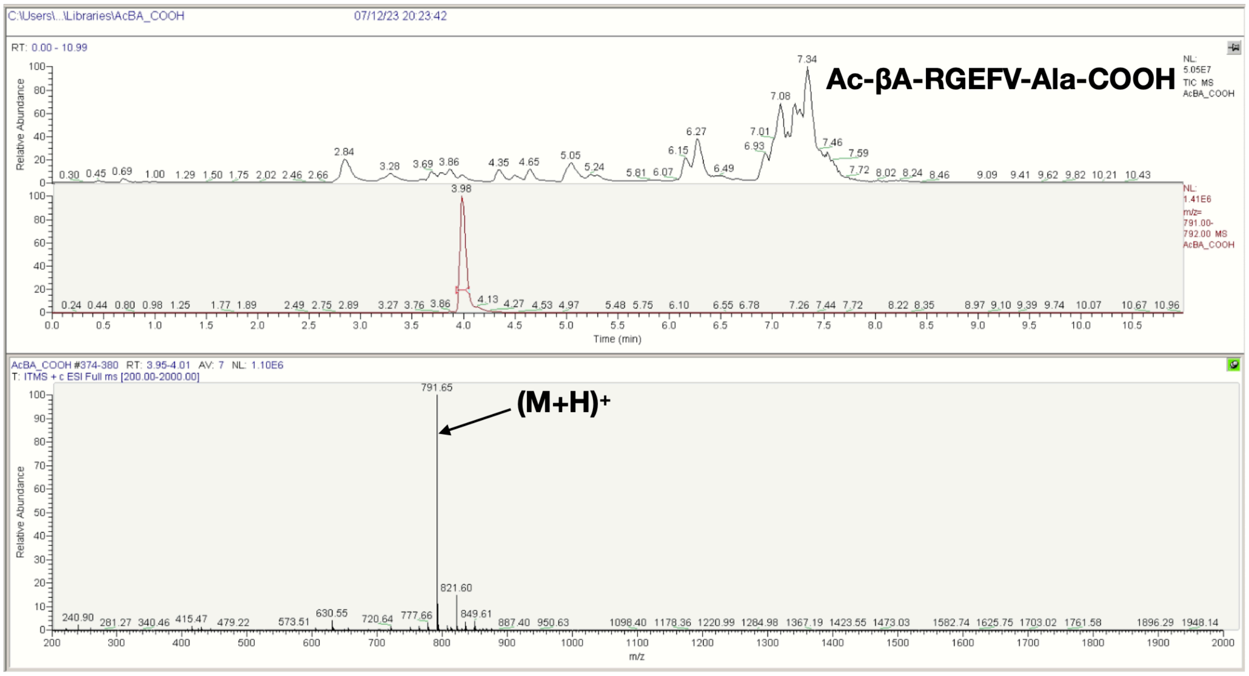
C:\Users\...\Libraries\AcBA_COOH
07/12/23 20:23:42
RT: 0.00 - 10.99
Ac-βA-RGEFV-Ala-COOH
Relative Abundance
Time (min)
AcBA_COOH #374-380 RT: 3.95-4.01 AV: 7 NL: 1.10E6
T: ITMS + c ESI Full ms [200.00-2000.00]
791.65
(M+H)+
821.60
m/z

**B**

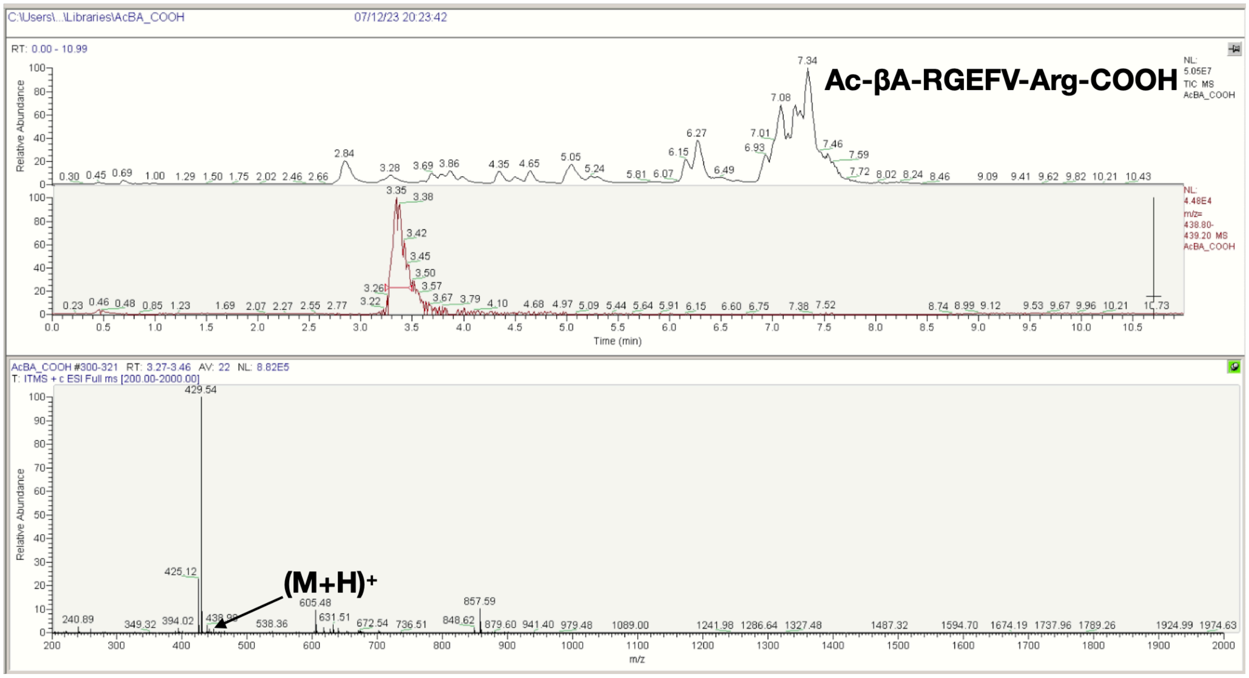
C:\Users\...\Libraries\AcBA_COOH
07/12/23 20:23:42
RT: 0.00 - 10.99
Ac-βA-RGEFV-Arg-COOH
Relative Abundance
Time (min)
AcBA_COOH #300-321 RT: 3.27-3.46 AV: 22 NL: 8.82E5
T: ITMS + c ESI Full ms [200.00-2000.00]
429.54
425.12
(M+H)+
605.48
857.59
m/z

**C**

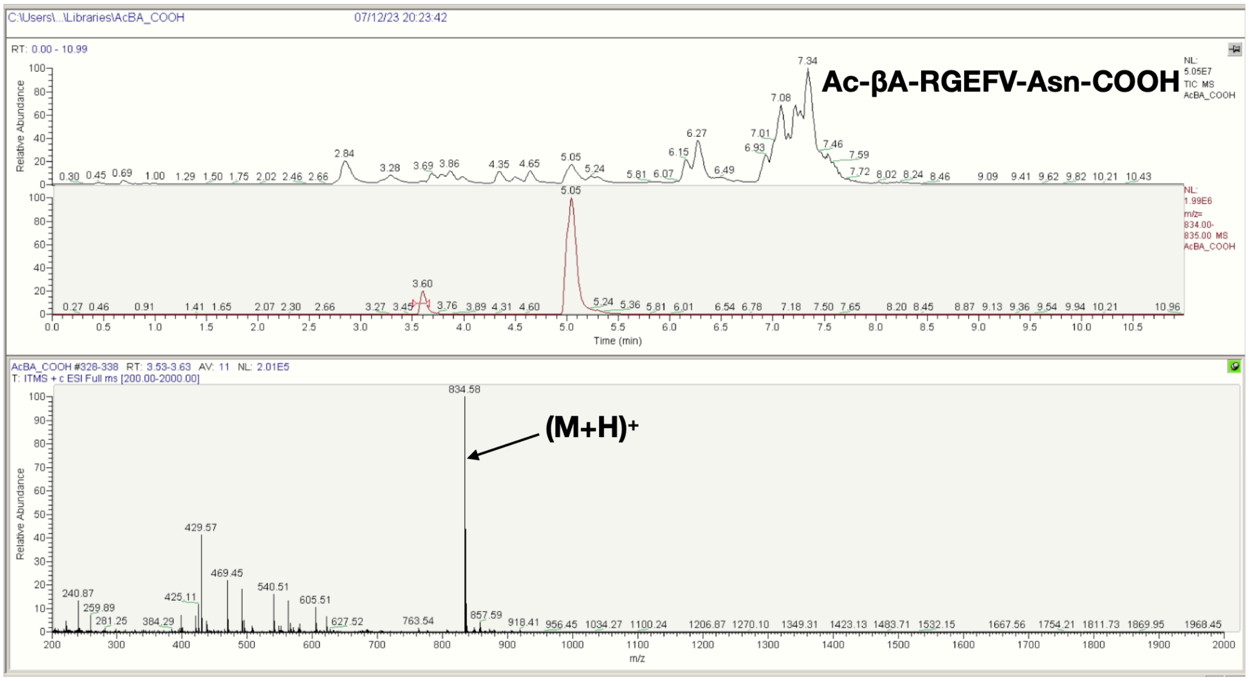

C:\Users\...\Libraries\AcBA_COOH
07/12/23 20:23:42
Ac-βA-RGEFV-Asn-COOH
Relative Abundance
Time (min)
AcBA_COOH #328-338 RT: 3.53-3.63 AV: 11 NL: 2.01E5
T: ITMS + c ESI Full ms [200.00-2000.00]
(M+H)+
834.58
m/z


**D**

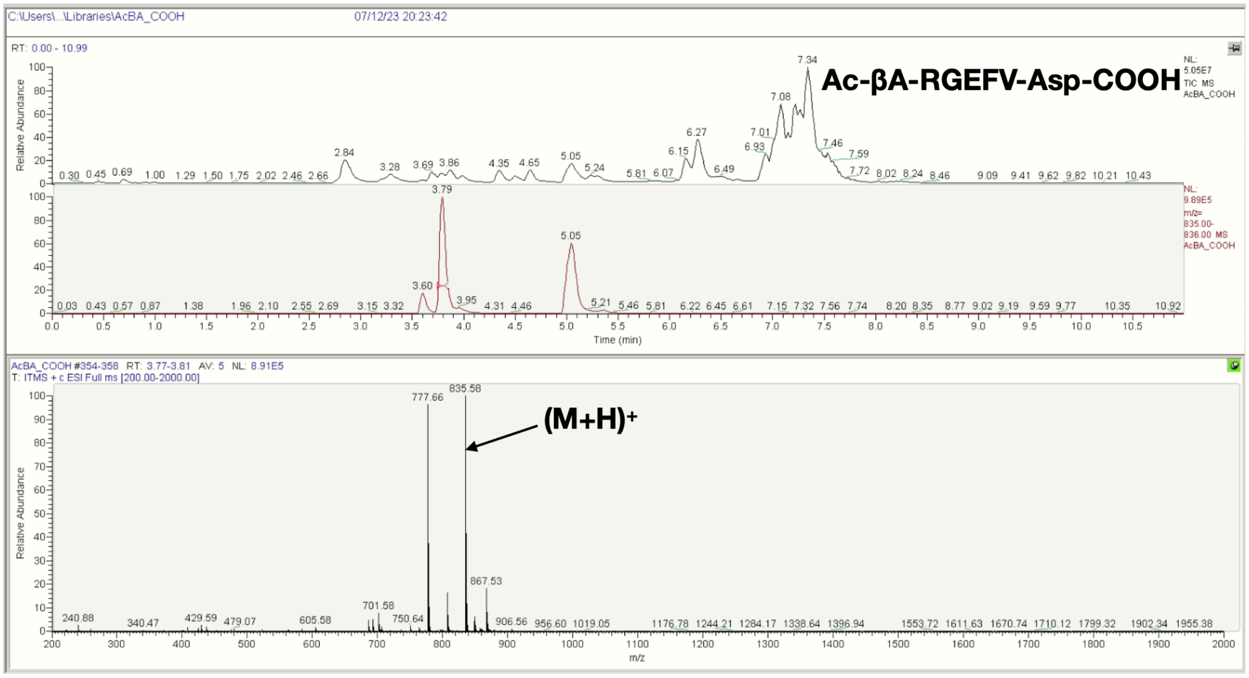

C:\Users\...\Libraries\AcBA_COOH
07/12/23 20:23:42
Ac-βA-RGEFV-Asp-COOH
Relative Abundance
Time (min)
AcBA_COOH #354-358 RT: 3.77-3.81 AV: 5 NL: 8.91E5
T: ITMS + c ESI Full ms [200.00-2000.00]
(M+H)+
835.58
777.66
m/z

E

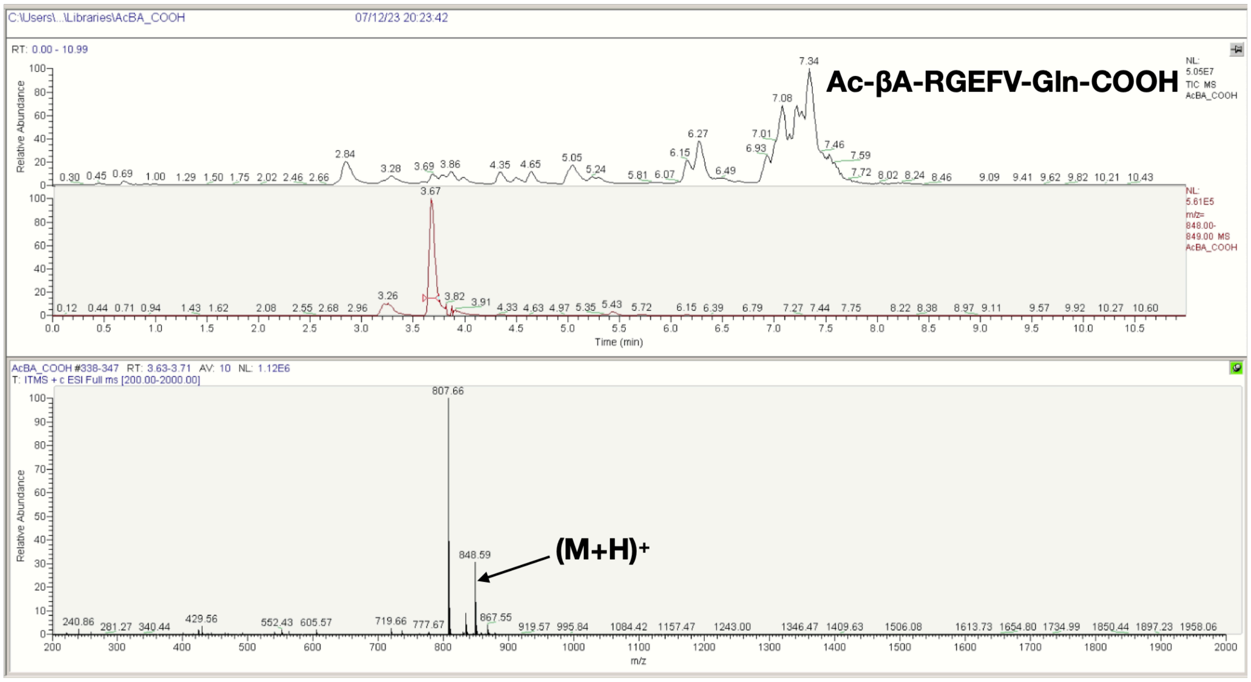
C:\Users\...\Libraries\AcBA_COOH
07/12/23 20:23:42
RT: 0.00 - 10.99
Ac-βA-RGEFV-Gln-COOH
NL: 5.05E7 TIC MS AcBA_COOH
NL: 5.61E5 m/z= 848.00-849.00 MS AcBA_COOH
Relative Abundance
Time (min)
AcBA_COOH #338-347 RT: 3.63-3.71 AV: 10 NL: 1.12E6
T: ITMS + c ESI Full ms [200.00-2000.00]
807.66
848.59
(M+H)+
m/z

F

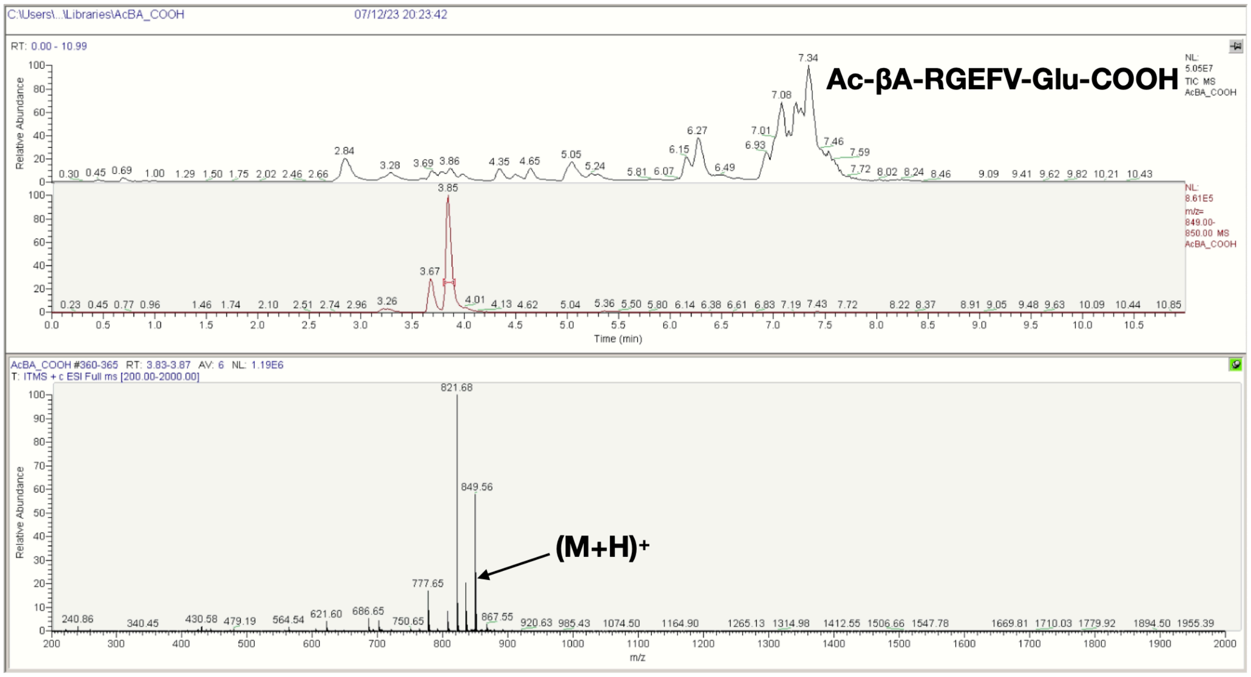
C:\Users\...\Libraries\AcBA_COOH
07/12/23 20:23:42
RT: 0.00 - 10.99
Ac-βA-RGEFV-Glu-COOH
NL: 5.05E7 TIC MS AcBA_COOH
NL: 8.61E5 m/z= 849.00-850.00 MS AcBA_COOH
Relative Abundance
Time (min)
AcBA_COOH #360-365 RT: 3.83-3.87 AV: 6 NL: 1.19E6
T: ITMS + c ESI Full ms [200.00-2000.00]
821.68
849.56
(M+H)+
m/z

G

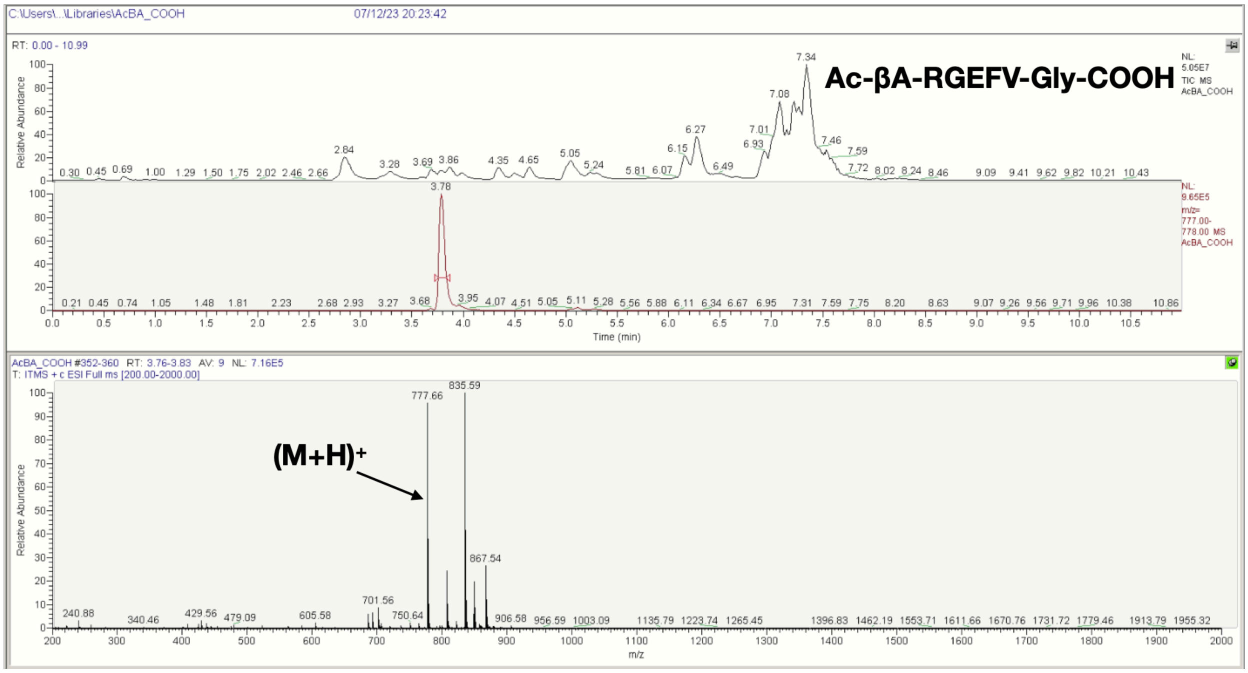
C:\Users\...\Libraries\AcBA_COOH
07/12/23 20:23:42
RT: 0.00 - 10.99
Ac-βA-RGEFV-Gly-COOH
NL: 5.05E7 TIC MS AcBA_COOH
NL: 9.65E5 m/z= 777.00-778.00 MS AcBA_COOH
Relative Abundance
Time (min)
AcBA_COOH #352-360 RT: 3.76-3.83 AV: 9 NL: 7.16E5
T: ITMS + c ESI Full ms [200.00-2000.00]
(M+H)+
777.66
835.59
867.54
m/z

H

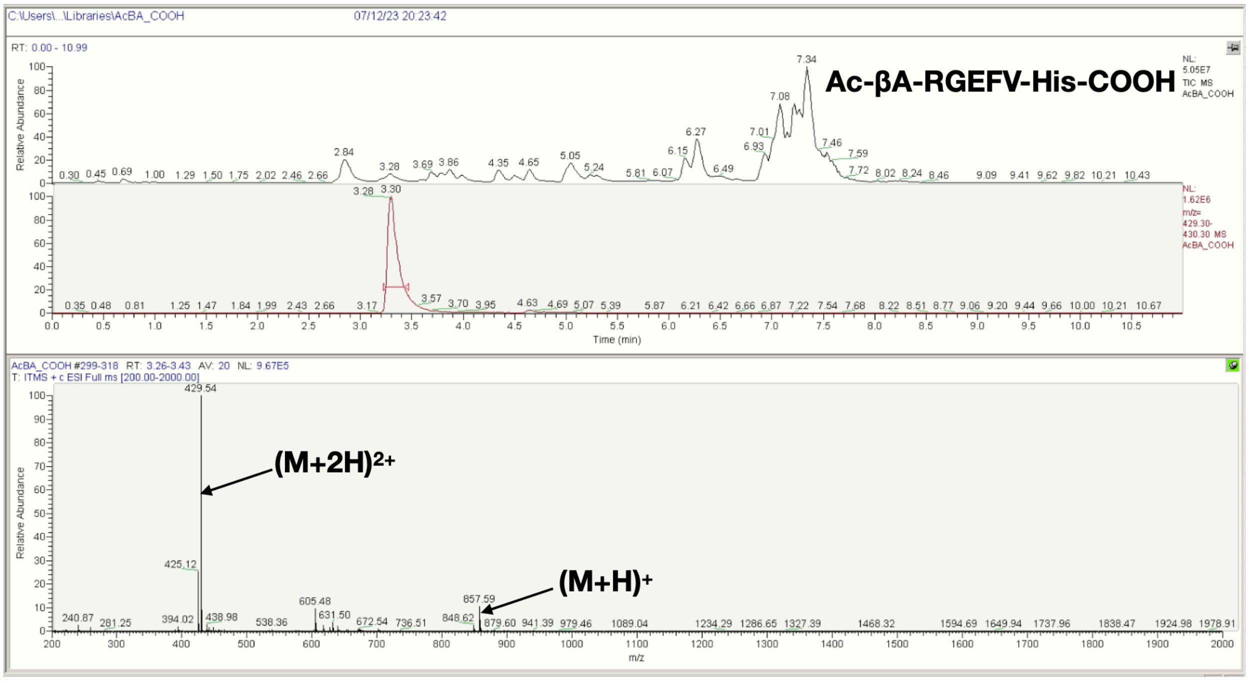
C:\Users\...\Libraries\AcBA_COOH
07/12/23 20:23:42
RT: 0.00 - 10.99
Ac-βA-RGEFV-His-COOH
NL: 5.05E7 TIC MS AcBA_COOH
NL: 1.62E6 m/z= 429.30-430.30 MS AcBA_COOH
Relative Abundance
Time (min)
AcBA_COOH #299-318 RT: 3.26-3.43 AV: 20 NL: 9.67E5
T: ITMS + c ESI Full ms [200.00-2000.00]
(M+2H)2+
429.54
(M+H)+
857.59
m/z

I

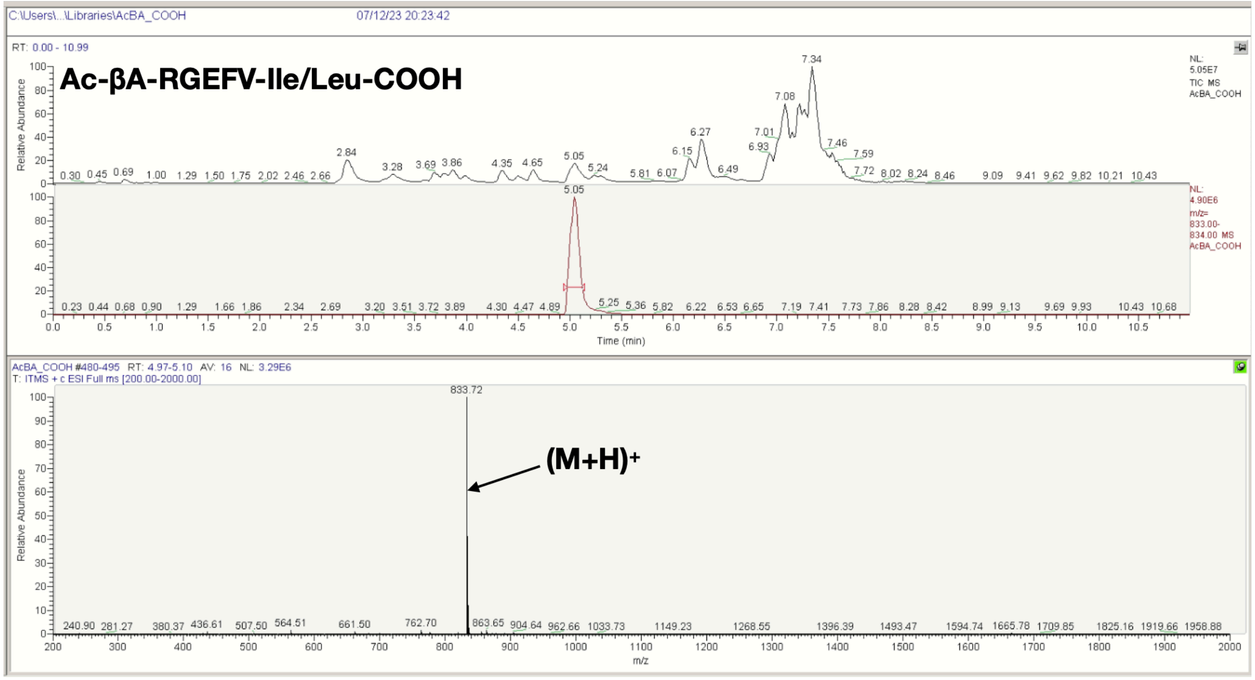


J

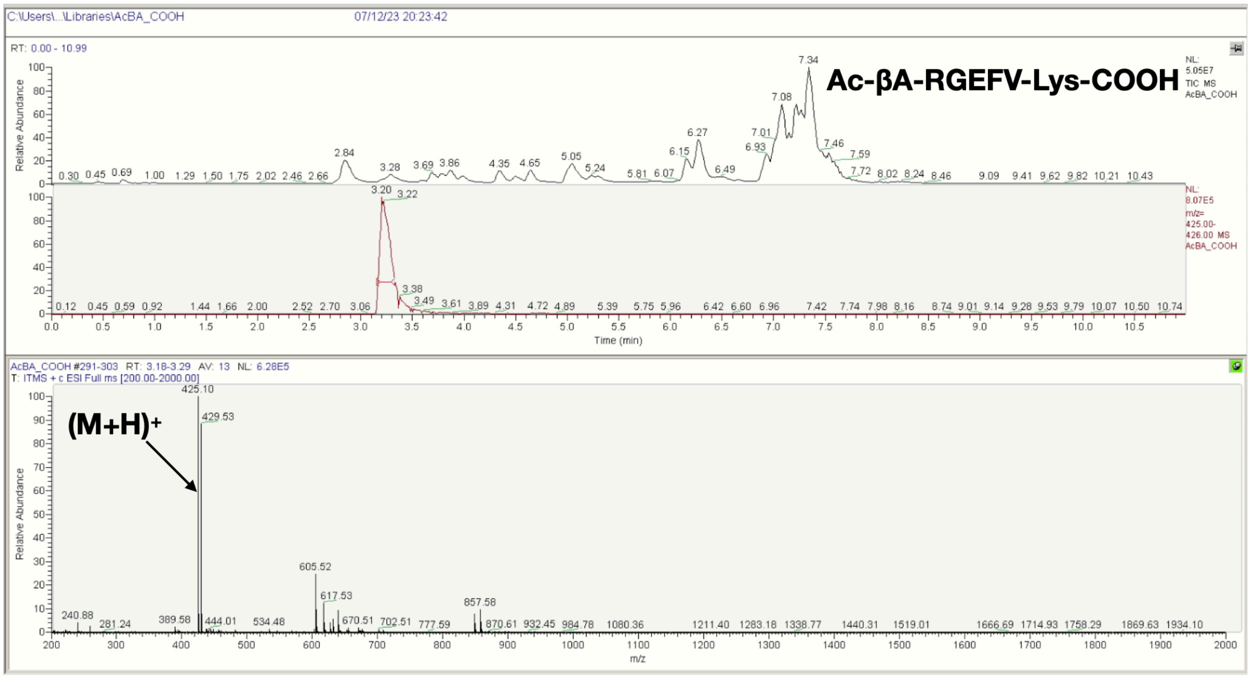

K

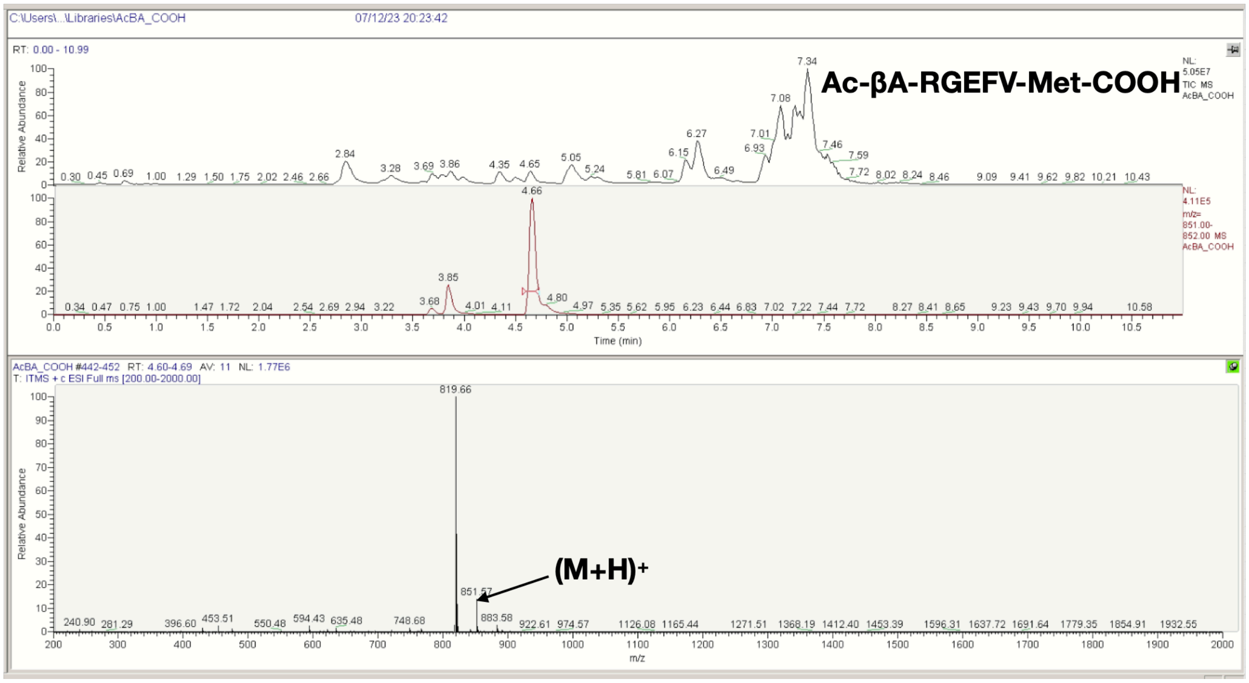
C:\Users\...\Libraries\AcBA_COOH
07/12/23 20:23:42
RT: 0.00 - 10.99
Ac-βA-RGEFV-Met-COOH
NL: 5.05E7 TIC MS AcBA_COOH
NL: 4.11E5 m/z= 851.00-852.00 MS AcBA_COOH
Relative Abundance
Time (min)
AcBA_COOH #442-452 RT: 4.60-4.69 AV: 11 NL: 1.77E6
T: ITMS + c ESI Full ms [200.00-2000.00]
819.66
(M+H)+
851.57
m/z

L

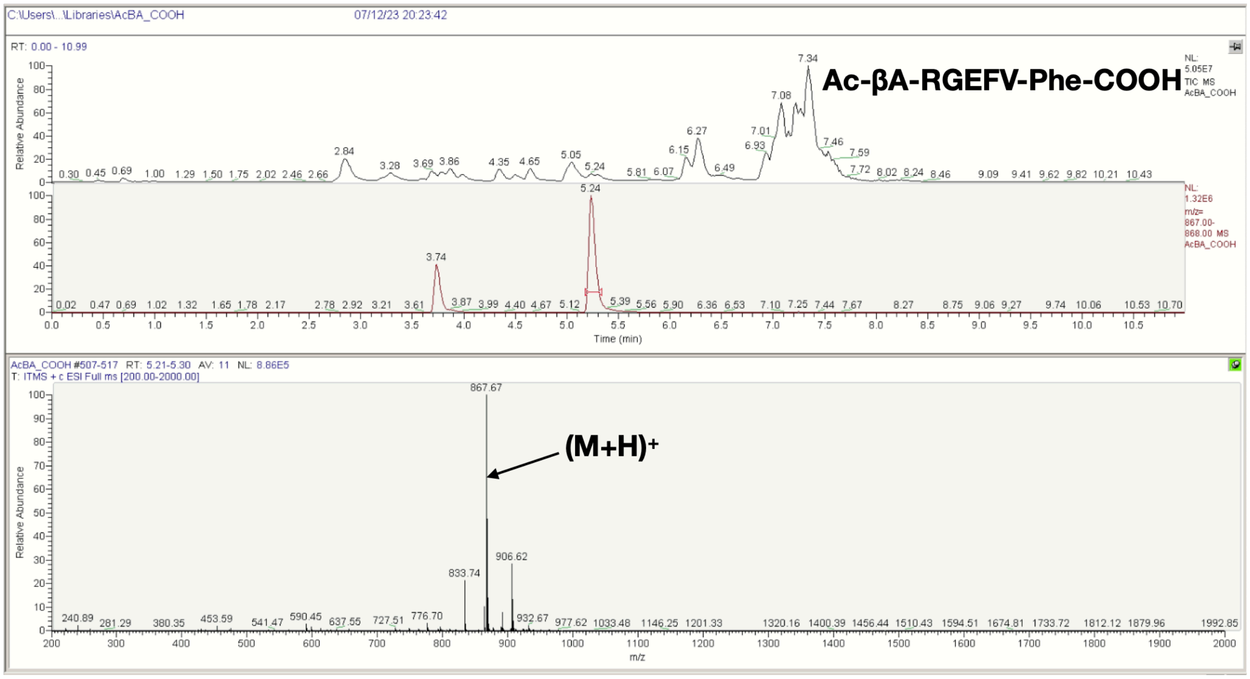
C:\Users\...\Libraries\AcBA_COOH
07/12/23 20:23:42
RT: 0.00 - 10.99
Ac-βA-RGEFV-Phe-COOH
NL: 5.05E7 TIC MS AcBA_COOH
NL: 1.32E6 m/z= 867.00-868.00 MS AcBA_COOH
Relative Abundance
Time (min)
AcBA_COOH #507-517 RT: 5.21-5.30 AV: 11 NL: 8.86E5
T: ITMS + c ESI Full ms [200.00-2000.00]
867.67
(M+H)+
906.62
833.74
m/z

**M**

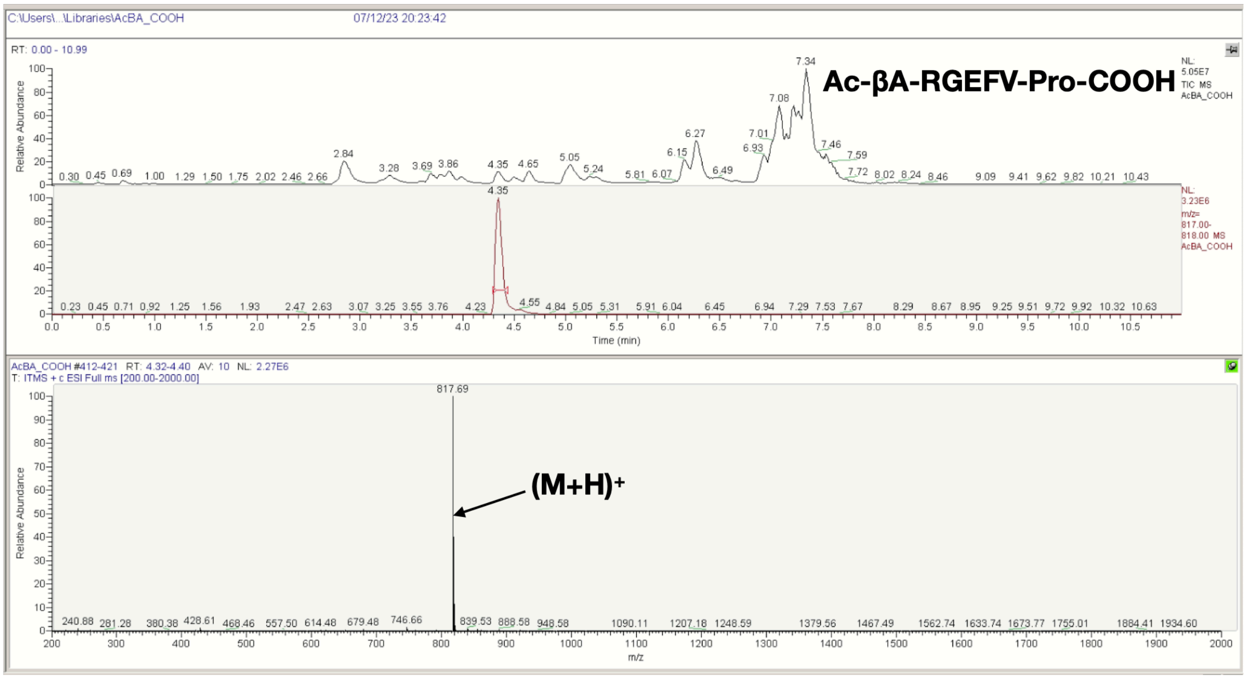
C:\Users\...\Libraries\AcBA_COOH
07/12/23 20:23:42
RT: 0.00 - 10.99
Ac-βA-RGEFV-Pro-COOH
NL: 5.05E7 TIC MS AcBA_COOH
NL: 3.23E6 m/z= 817.00-818.00 MS AcBA_COOH
Relative Abundance
Time (min)
4.35
7.34
AcBA_COOH #412-421 RT: 4.32-4.40 AV: 10 NL: 2.27E6
T: ITMS + c ESI Full ms [200.00-2000.00]
817.69
(M+H)+
m/z

**N**

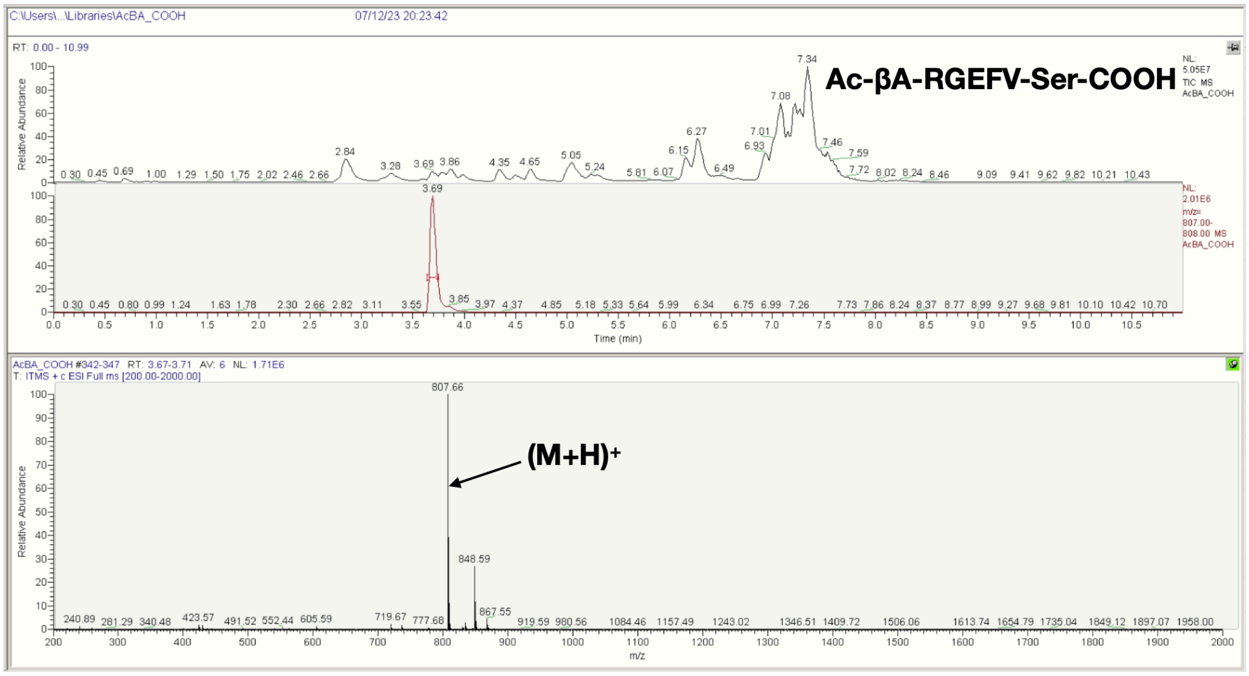
C:\Users\...\Libraries\AcBA_COOH
07/12/23 20:23:42
RT: 0.00 - 10.99
Ac-βA-RGEFV-Ser-COOH
NL: 5.05E7 TIC MS AcBA_COOH
NL: 2.01E6 m/z= 807.00-808.00 MS AcBA_COOH
Relative Abundance
Time (min)
3.69
7.34
AcBA_COOH #342-347 RT: 3.67-3.71 AV: 6 NL: 1.71E6
T: ITMS + c ESI Full ms [200.00-2000.00]
807.66
848.59
867.55
(M+H)+
m/z

**O**

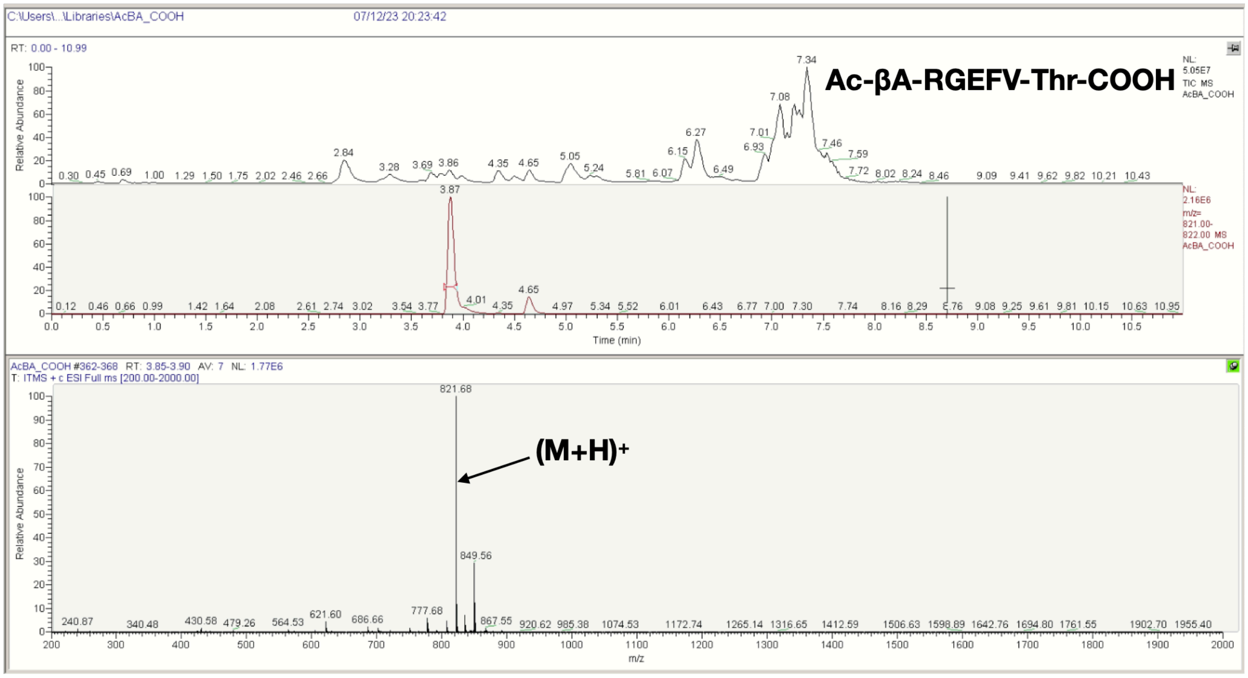
Ac-βA-RGEFV-Thr-COOH
(M+H)+
821.68
849.56

**P**

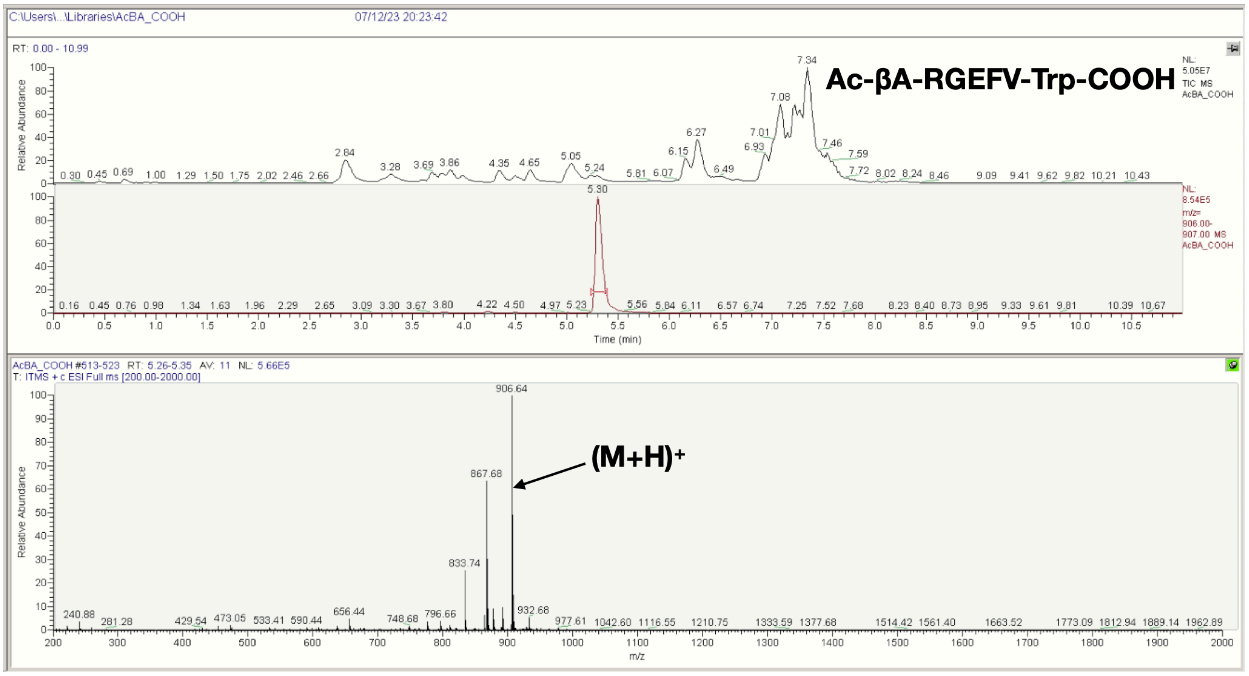
Ac-βA-RGEFV-Trp-COOH
(M+H)+
906.64
867.68

**Q**

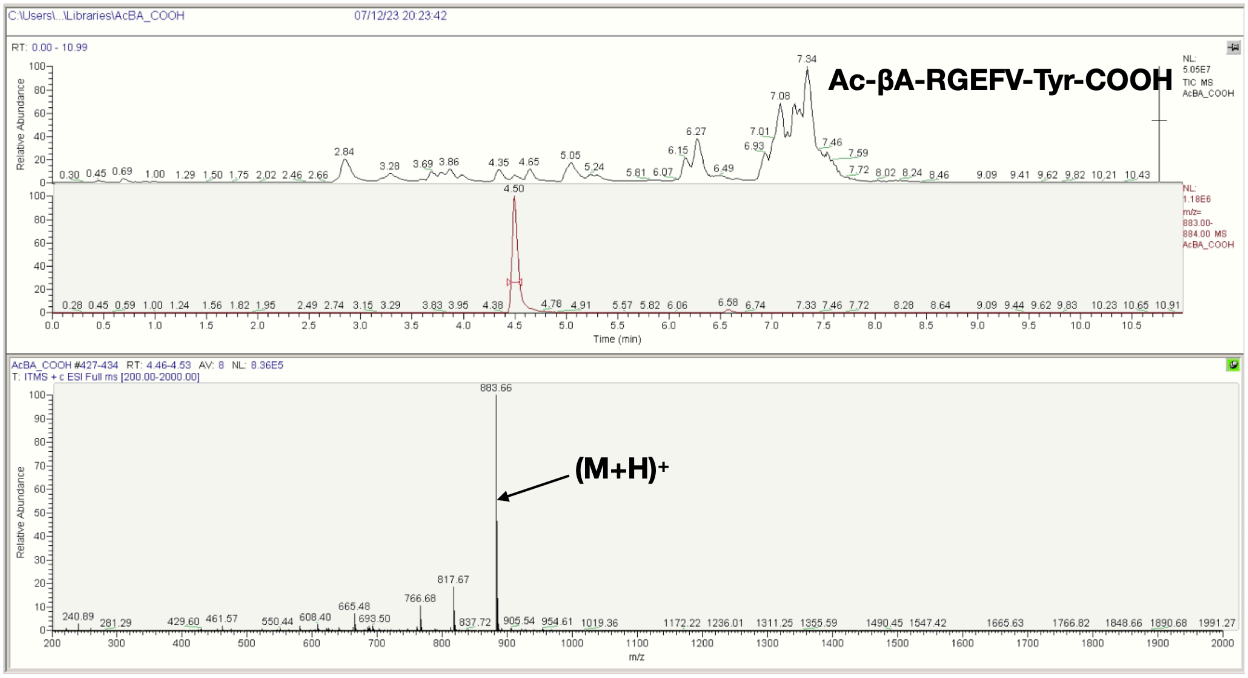


**R**

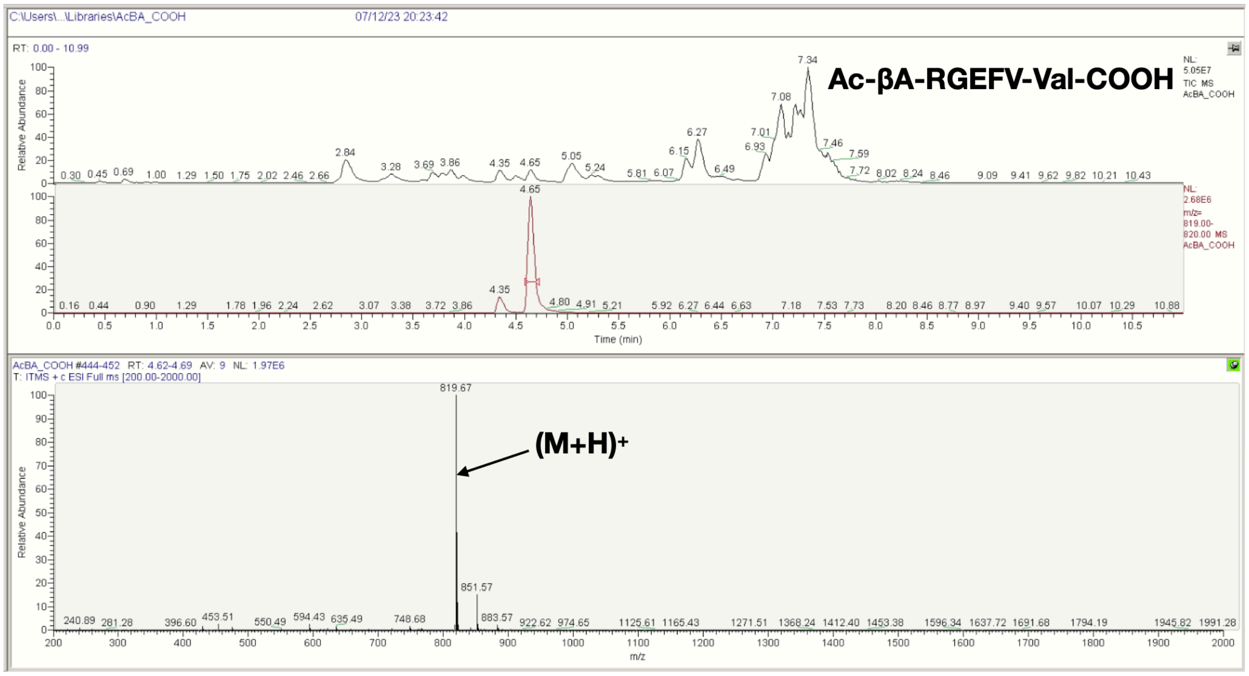


**Figure S16**. LCMS spectra of the Ac-βA-RGEFV-X-COOH libraries, where X = a) Ala, b) Arg, c) Asn, d) Asp, e) Gln, f) Glu, g) Gly, h) His, i) Ile/Leu, j) Lys, k) Met, l) Phe, m) Pro, n) Ser, o) Thr, p) Trp, q) Tyr, r) Val.

**A**

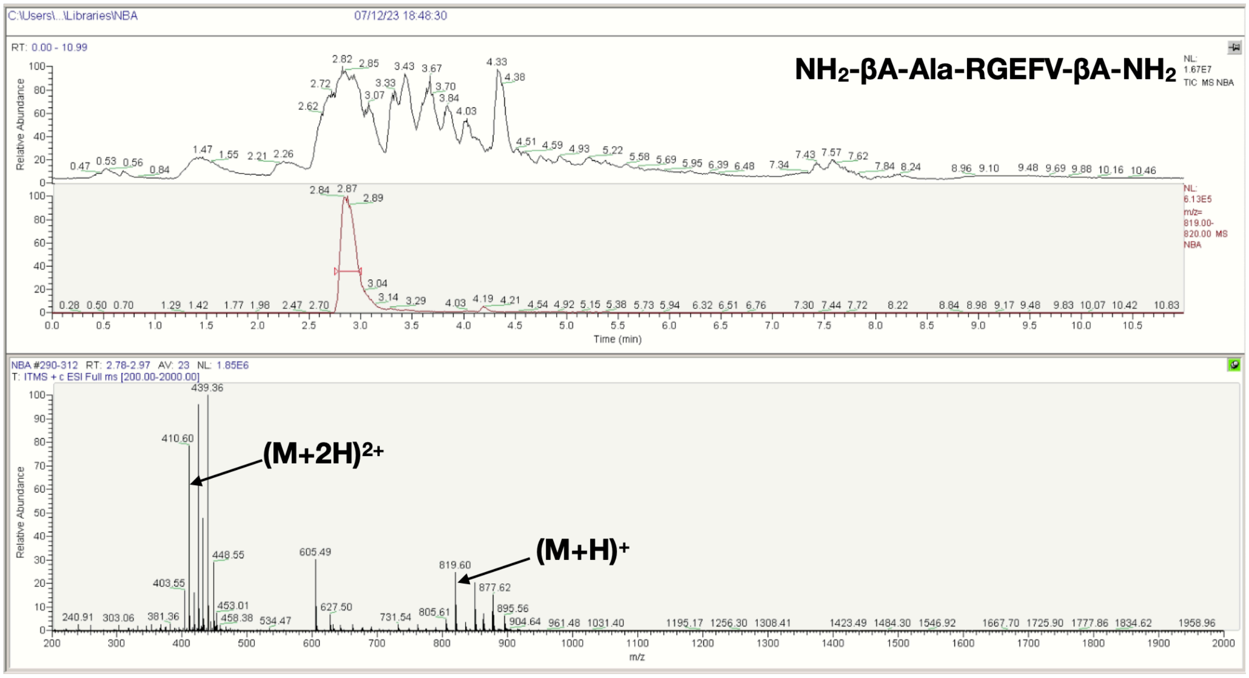

C:\Users\...\Libraries\NBA
07/12/23 18:48:30
RT: 0.00 - 10.99
NH2-βA-Ala-RGEFV-βA-NH2
NL: 1.67E7 TIC MS NBA
NL: 6.13E5 m/z= 819.00-820.00 MS NBA
Relative Abundance
Time (min)
NBA #290-312 RT: 2.78-2.97 AV: 23 NL: 1.85E6
T: ITMS + c ESI Full ms [200.00-2000.00]
(M+2H)2+
(M+H)+
m/z


**B**

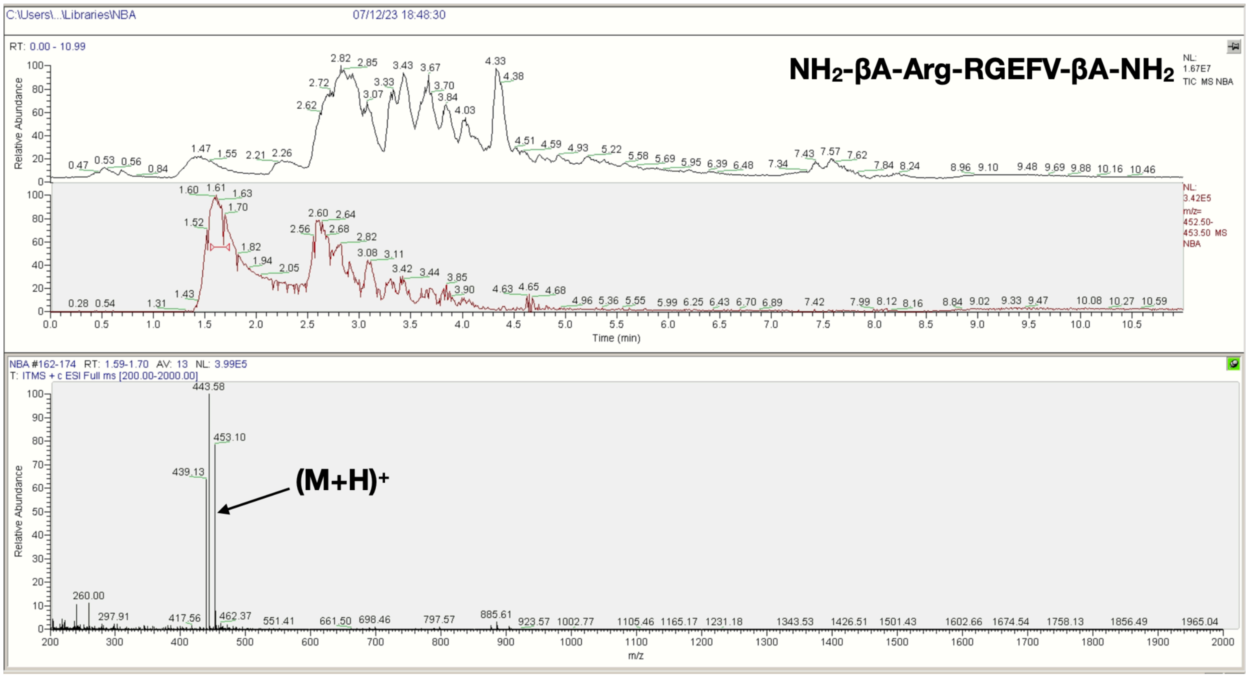

C:\Users\...\Libraries\NBA
07/12/23 18:48:30
RT: 0.00 - 10.99
NH2-βA-Arg-RGEFV-βA-NH2
NL: 1.67E7 TIC MS NBA
NL: 3.42E5 m/z= 452.50-453.50 MS NBA
Relative Abundance
Time (min)
NBA #162-174 RT: 1.59-1.70 AV: 13 NL: 3.99E5
T: ITMS + c ESI Full ms [200.00-2000.00]
(M+H)+
m/z

C

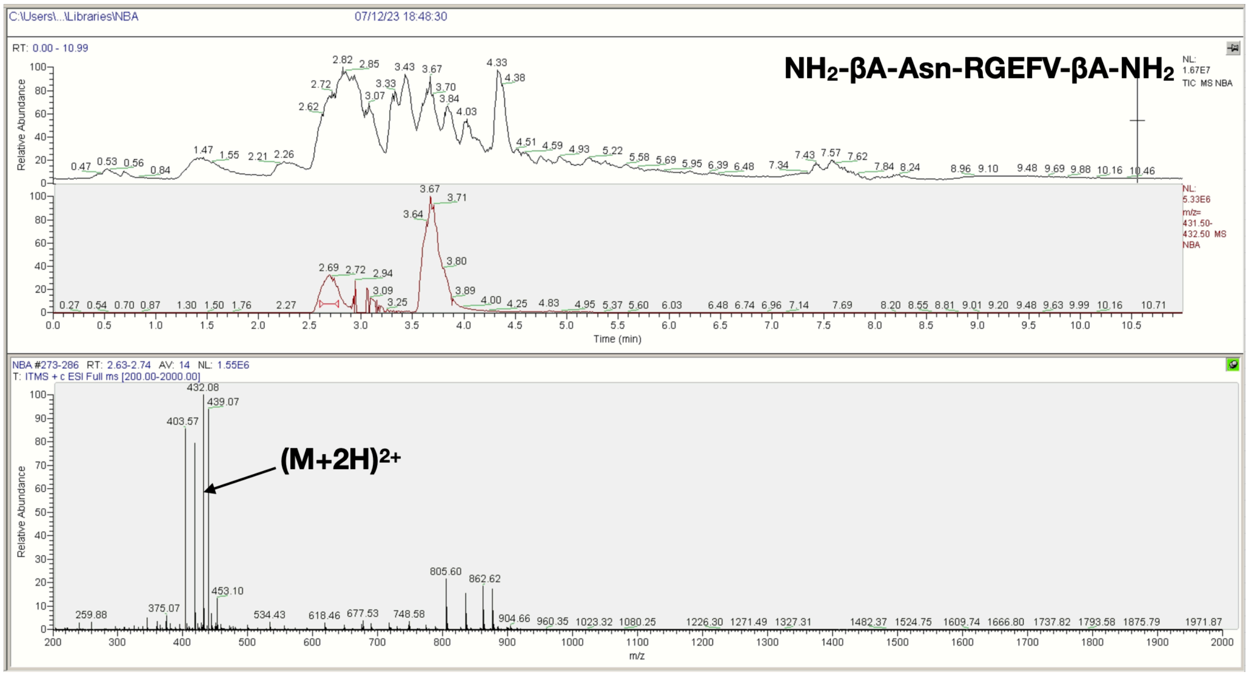
C:\Users\...\Libraries\NBA
07/12/23 18:48:30
RT: 0.00 - 10.99
NH2-βA-Asn-RGEFV-βA-NH2
NL: 1.67E7 TIC MS NBA
NL: 5.33E6 m/z= 431.50-432.50 MS NBA
Relative Abundance
Time (min)
NBA #273-286 RT: 2.63-2.74 AV: 14 NL: 1.55E6
T: ITMS + c ESI Full ms [200.00-2000.00]
(M+2H)2+
432.08
439.07
403.57
805.60
862.62
m/z

D

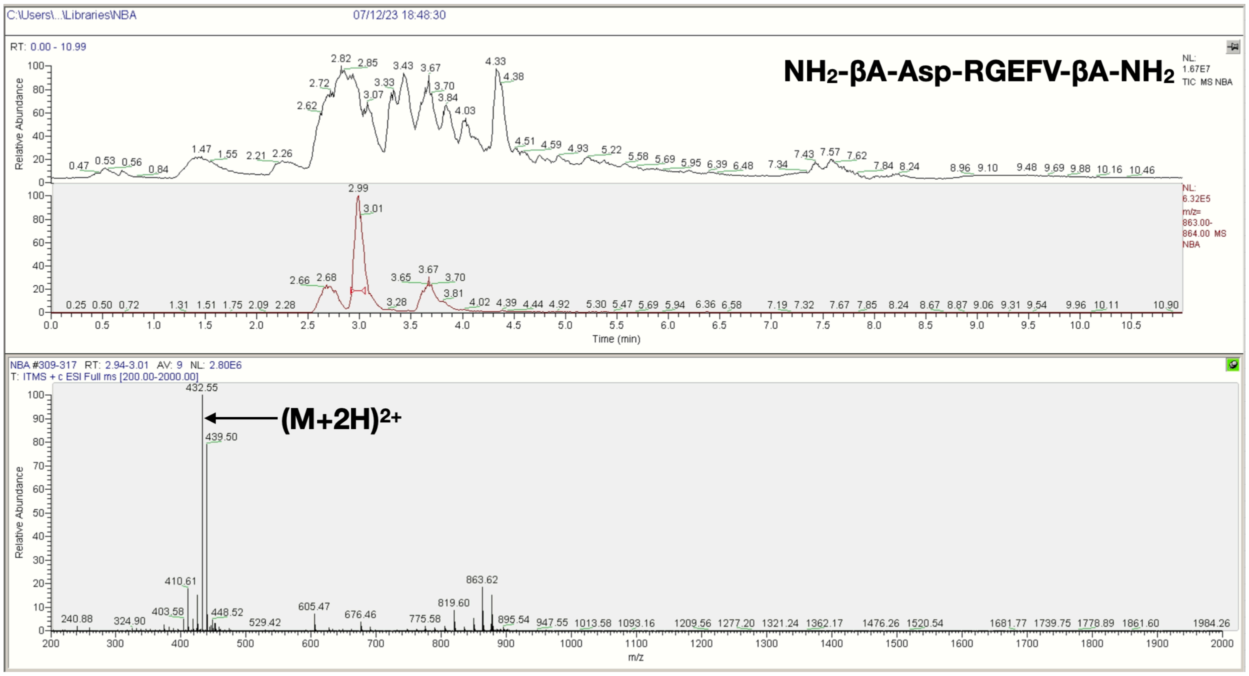
C:\Users\...\Libraries\NBA
07/12/23 18:48:30
RT: 0.00 - 10.99
NH2-βA-Asp-RGEFV-βA-NH2
NL: 1.67E7 TIC MS NBA
NL: 6.32E5 m/z= 863.00-864.00 MS NBA
Relative Abundance
Time (min)
NBA #309-317 RT: 2.94-3.01 AV: 9 NL: 2.80E6
T: ITMS + c ESI Full ms [200.00-2000.00]
(M+2H)2+
432.55
439.50
410.61
863.62
819.60
m/z

E

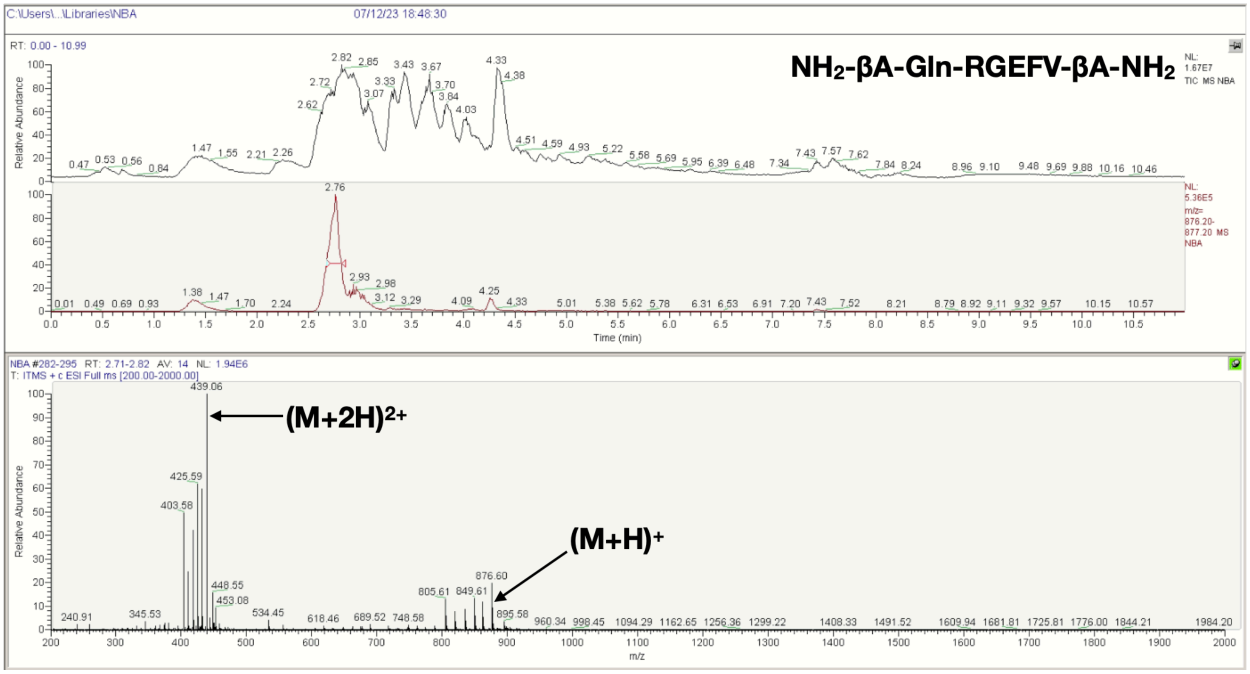
NH2-βA-Gln-RGEFV-βA-NH2
(M+2H)2+
(M+H)+
Relative Abundance
Time (min)
m/z

F

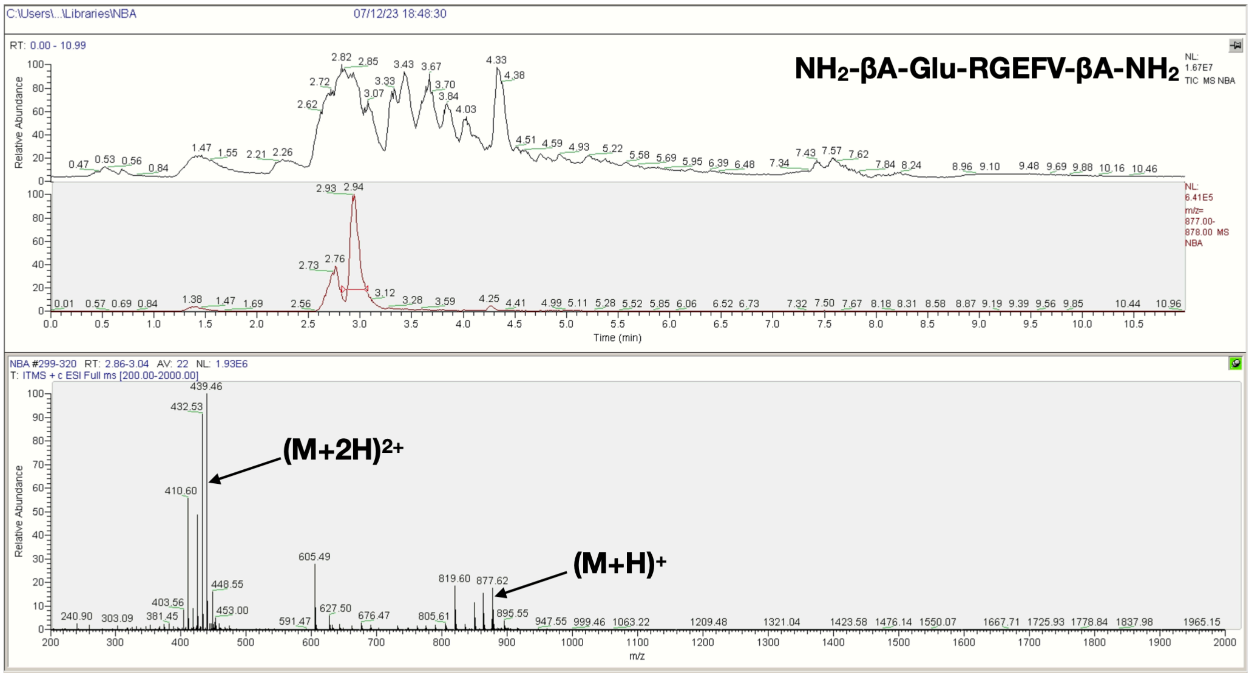
NH2-βA-Glu-RGEFV-βA-NH2
(M+2H)2+
(M+H)+
Relative Abundance
Time (min)
m/z

G

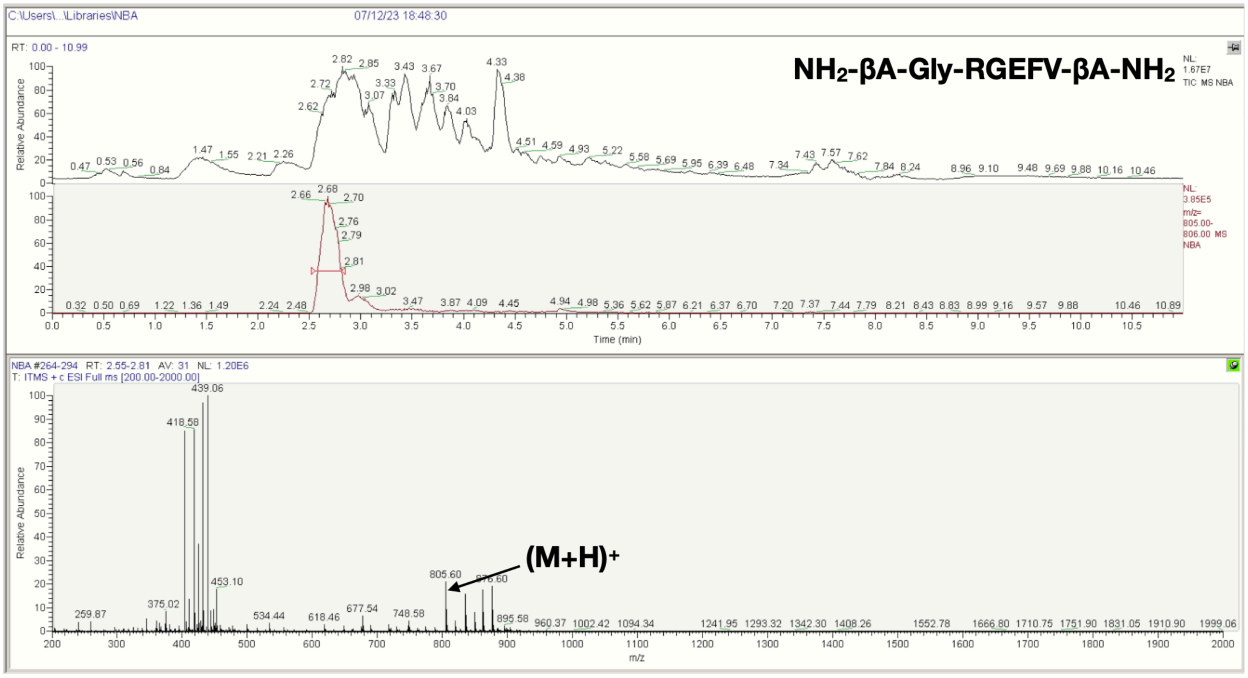
C:\Users\...\Libraries\NBA
07/12/23 18:48:30
RT: 0.00 - 10.99
NH2-βA-Gly-RGEFV-βA-NH2
NL: 1.67E7 TIC MS NBA
NL: 3.85E5 m/z= 805.00-806.00 MS NBA
Relative Abundance
Time (min)
NBA #264-294 RT: 2.55-2.81 AV: 31 NL: 1.20E6
T: ITMS + c ESI Full ms [200.00-2000.00]
439.06
418.58
453.10
805.60
876.60
(M+H)+
m/z

H

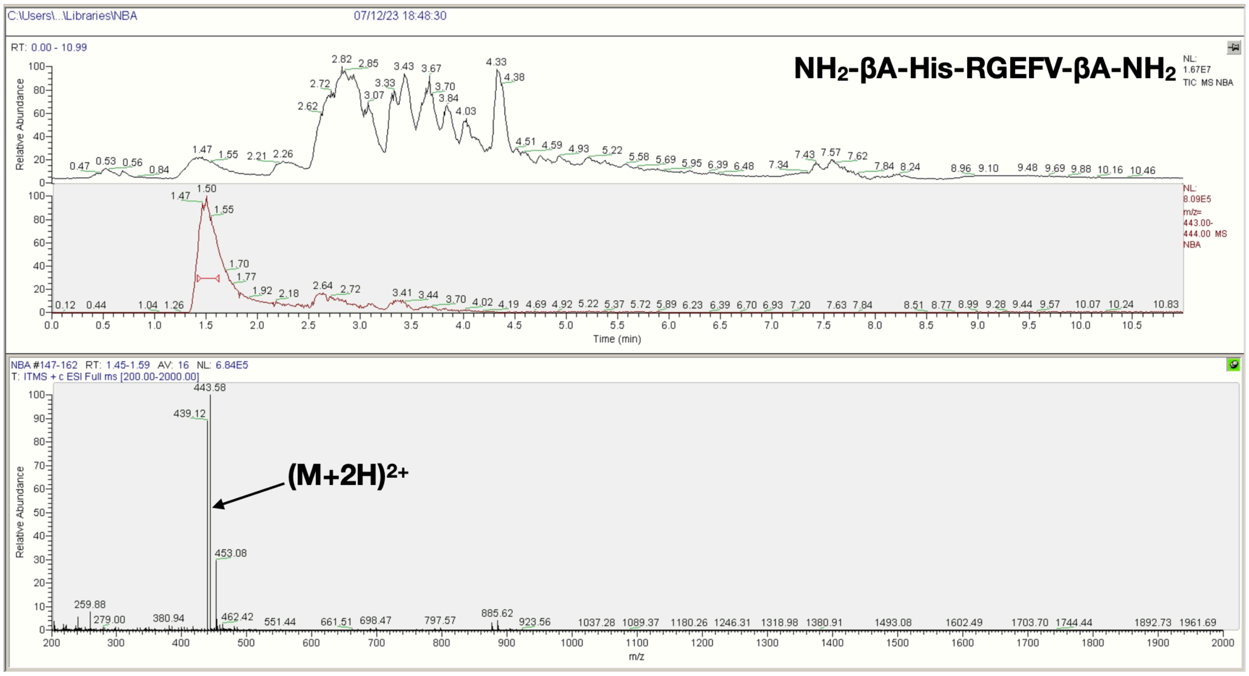
C:\Users\...\Libraries\NBA
07/12/23 18:48:30
RT: 0.00 - 10.99
NH2-βA-His-RGEFV-βA-NH2
NL: 1.67E7 TIC MS NBA
NL: 8.09E5 m/z= 443.00-444.00 MS NBA
Relative Abundance
Time (min)
NBA #147-162 RT: 1.45-1.59 AV: 16 NL: 6.84E5
T: ITMS + c ESI Full ms [200.00-2000.00]
443.58
439.12
453.08
(M+2H)2+
m/z

I

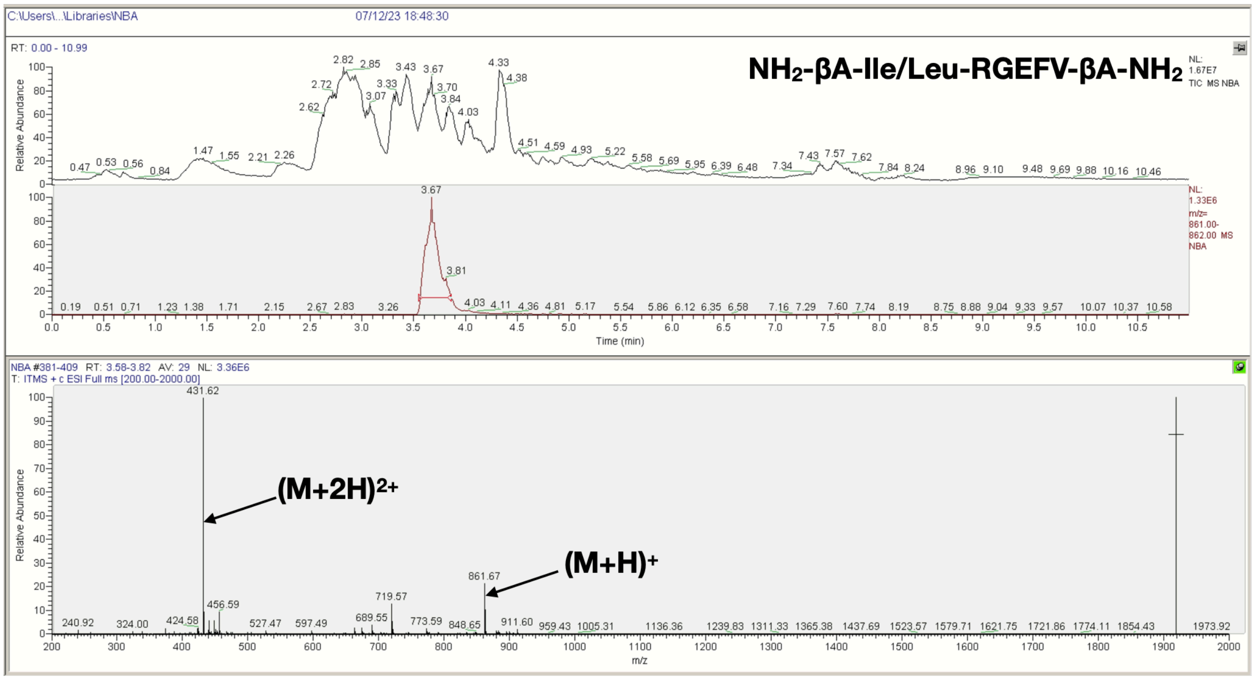
NH2-βA-Ile/Leu-RGEFV-βA-NH2
(M+2H)2+
(M+H)+
431.62
861.67
Relative Abundance
Time (min)
m/z

J

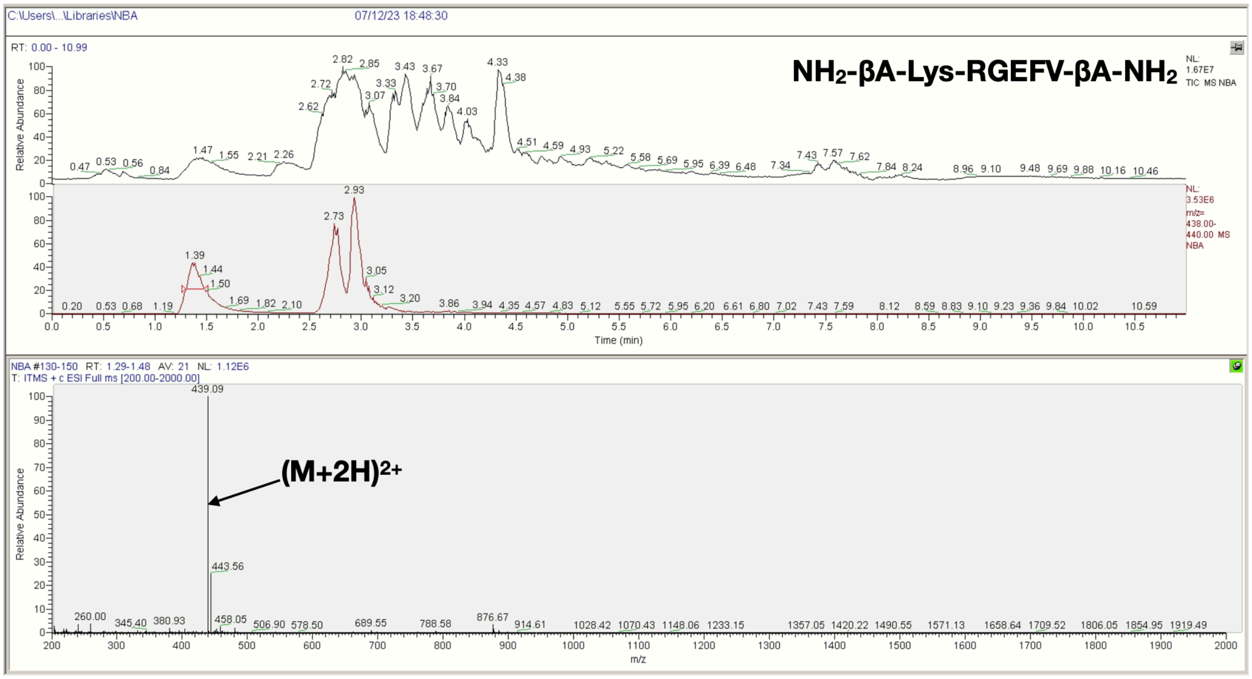
NH2-βA-Lys-RGEFV-βA-NH2
(M+2H)2+
439.09
443.56
Relative Abundance
Time (min)
m/z

K

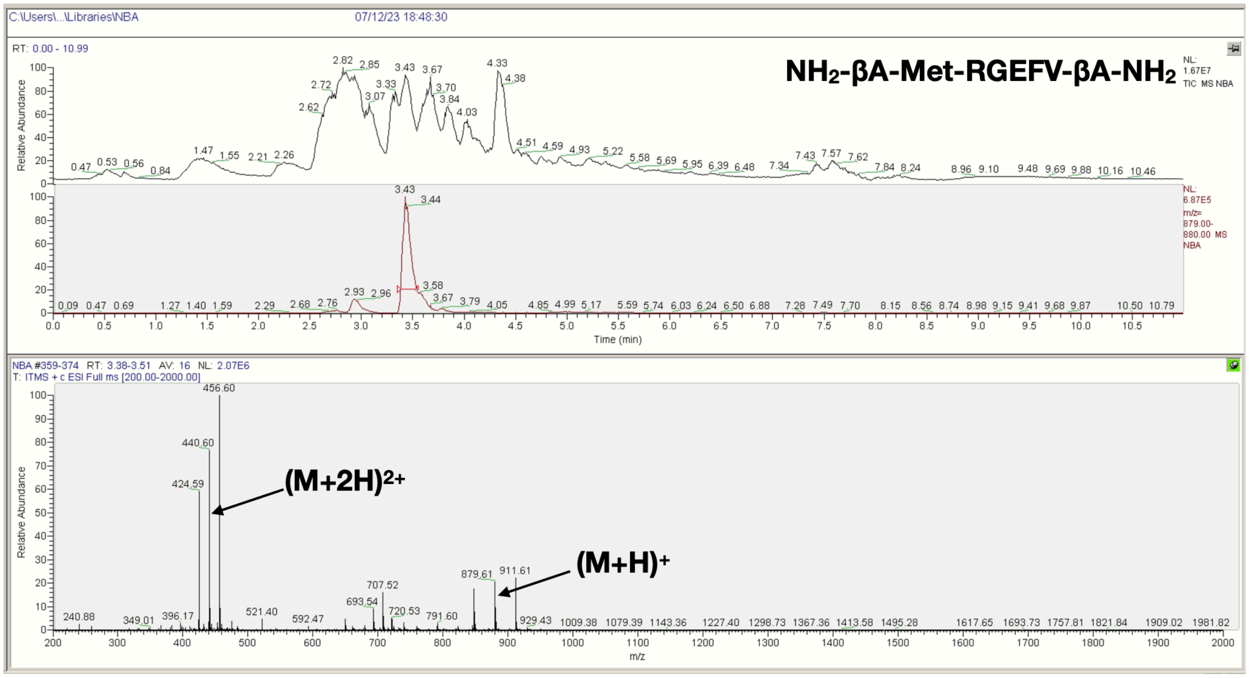


L

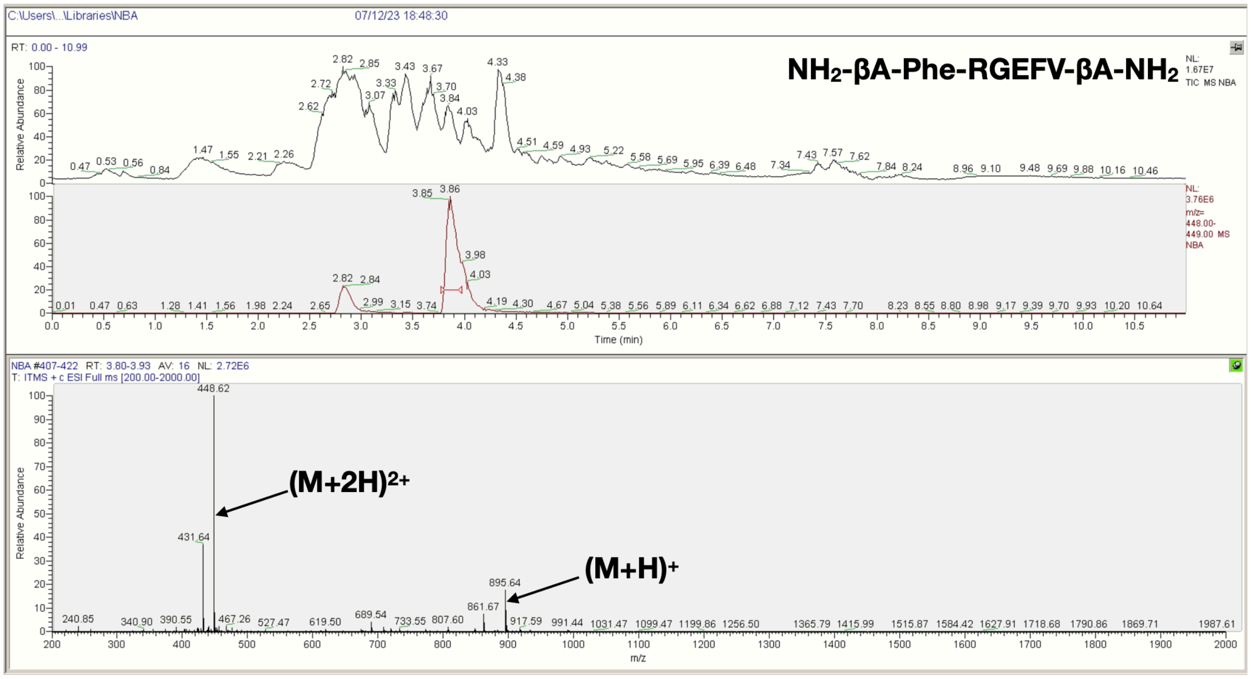

M

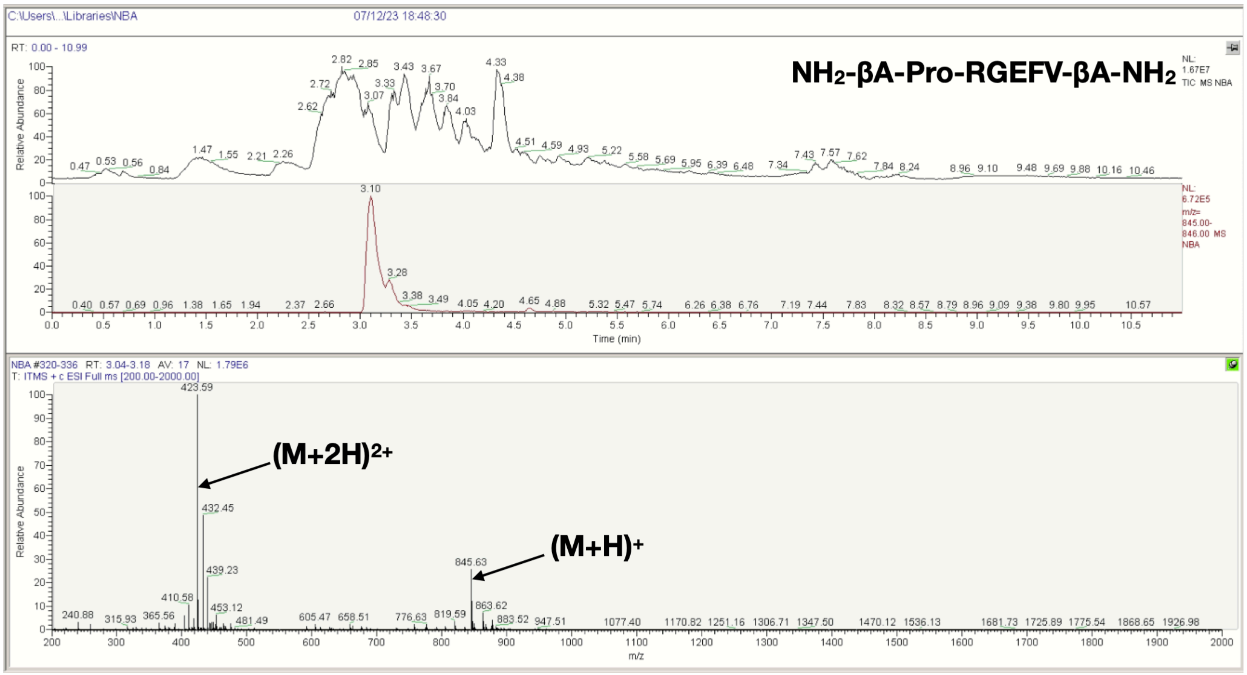
C:\Users\...\Libraries\NBA
07/12/23 18:48:30
RT: 0.00 - 10.99
NH2-βA-Pro-RGEFV-βA-NH2
NL: 1.67E7 TIC MS NBA
NL: 6.72E5 m/z= 845.00-846.00 MS NBA
Relative Abundance
Time (min)
NBA #320-336 RT: 3.04-3.18 AV: 17 NL: 1.79E6
T: ITMS + c ESI Full ms [200.00-2000.00]
423.59
(M+2H)2+
432.45
439.23
845.63
(M+H)+
863.62
m/z

N

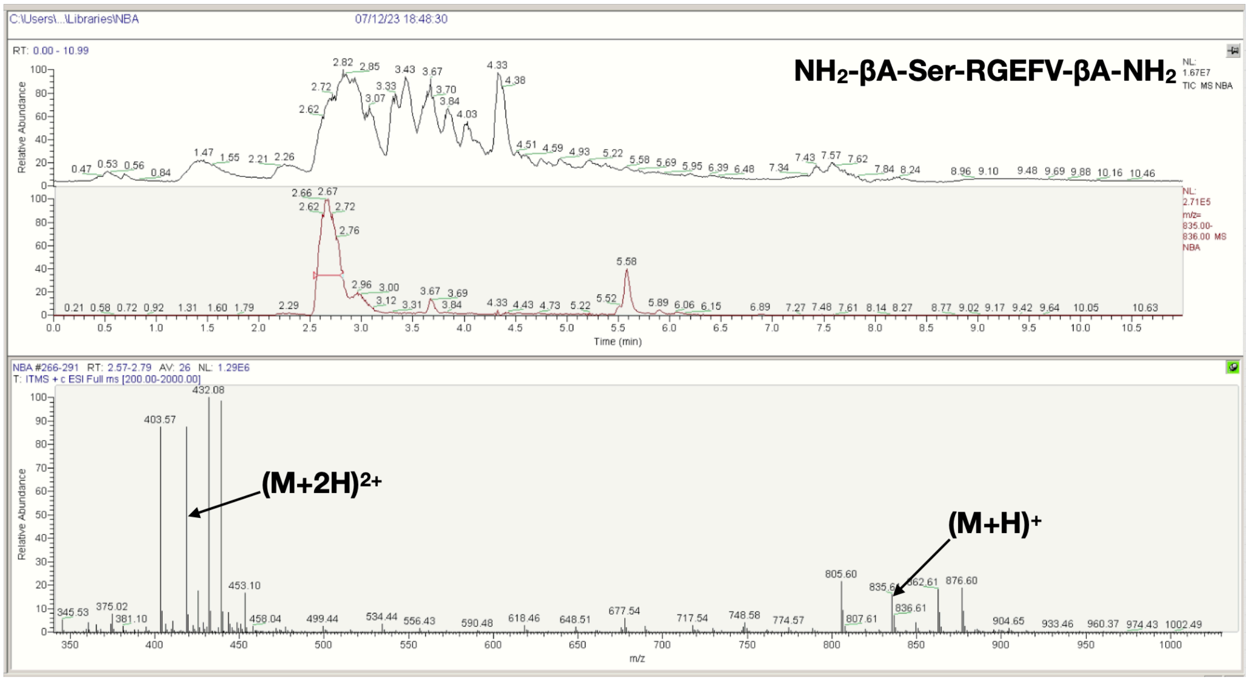
C:\Users\...\Libraries\NBA
07/12/23 18:48:30
RT: 0.00 - 10.99
NH2-βA-Ser-RGEFV-βA-NH2
NL: 1.67E7 TIC MS NBA
NL: 2.71E5 m/z= 835.00-836.00 MS NBA
Relative Abundance
Time (min)
NBA #266-291 RT: 2.57-2.79 AV: 26 NL: 1.29E6
T: ITMS + c ESI Full ms [200.00-2000.00]
432.08
403.57
(M+2H)2+
453.10
805.60
(M+H)+
835.61
862.61
876.60
m/z

**O**

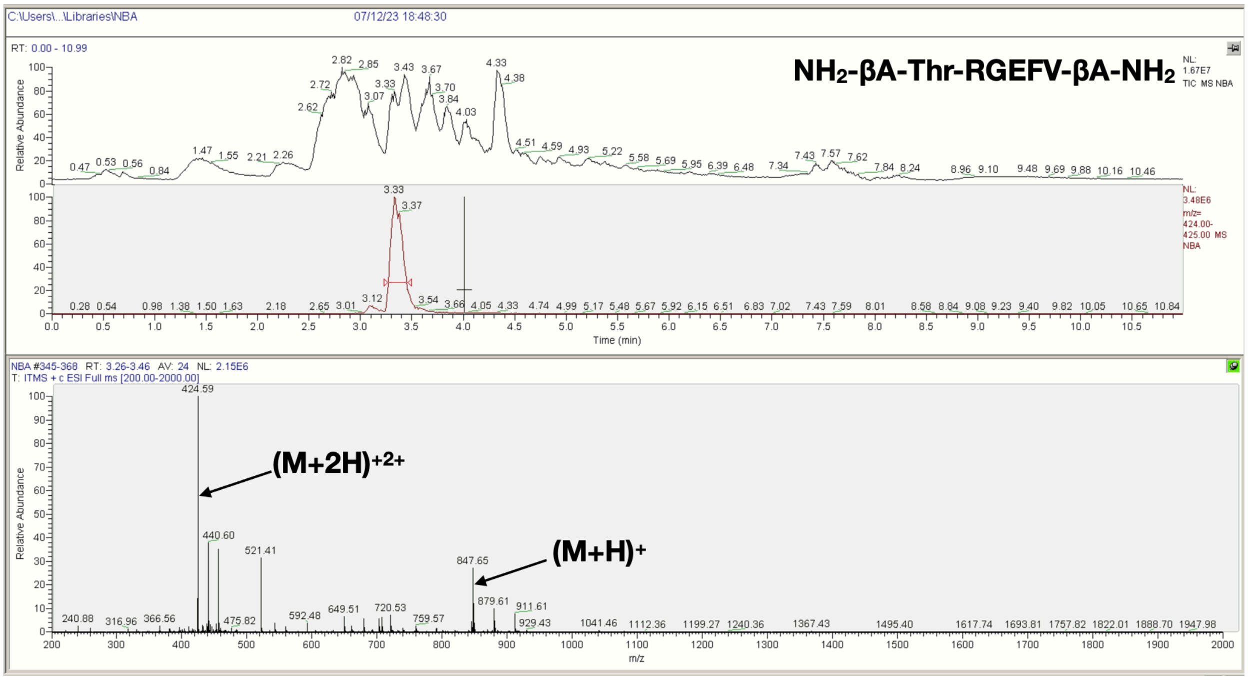


**P**

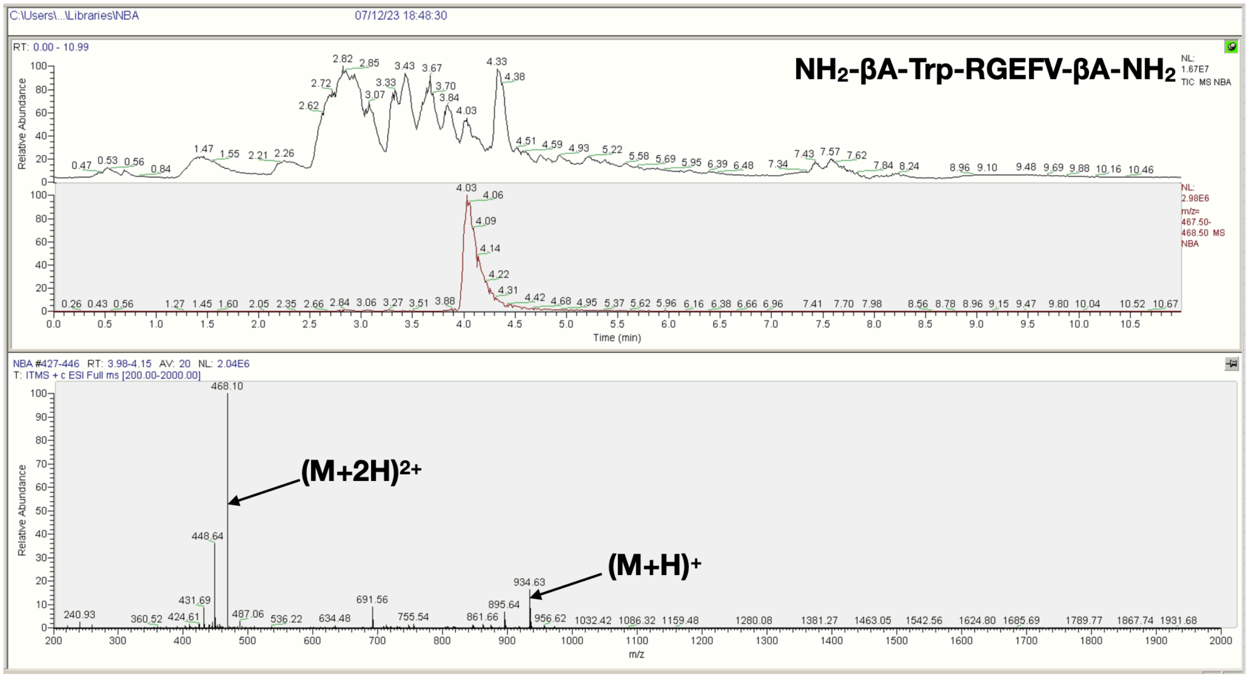

**Q**

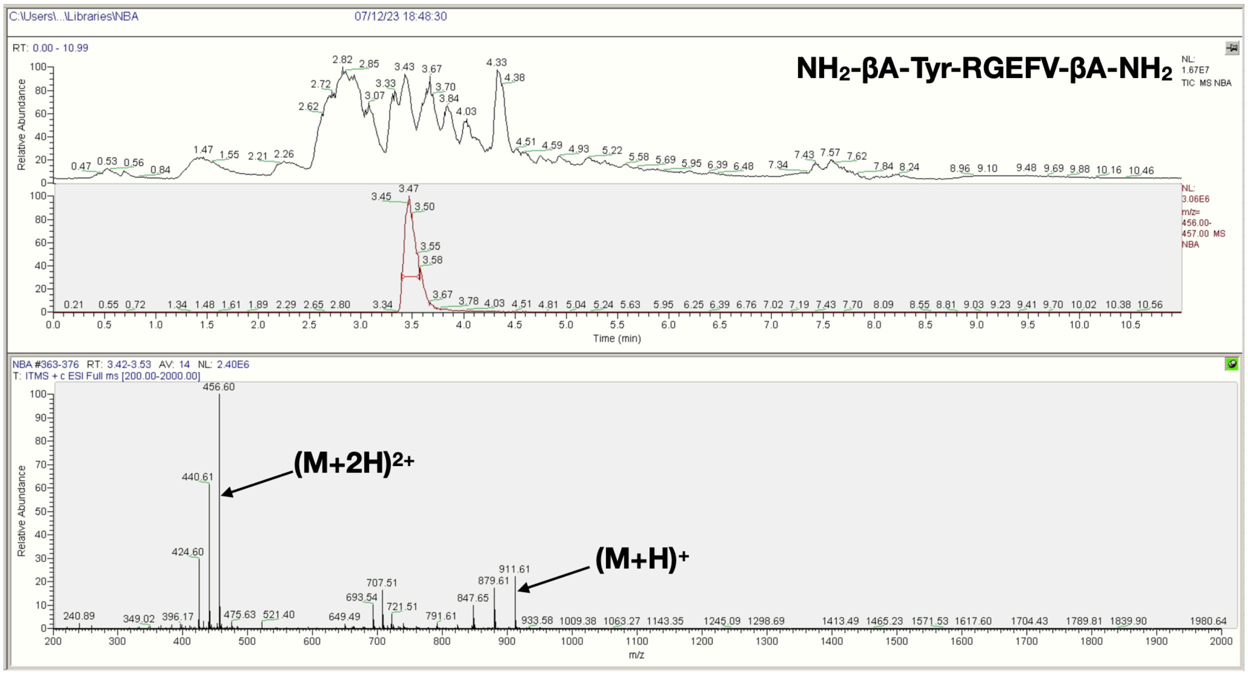


**R**

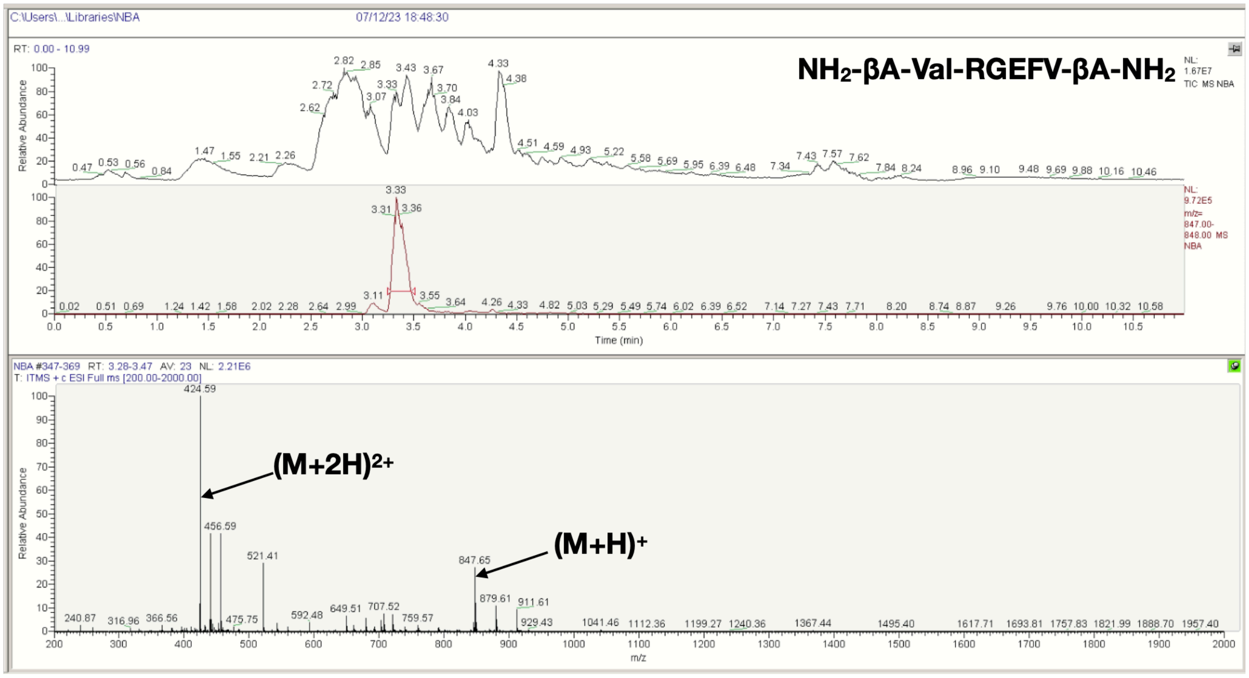


**Figure S17**. LCMS spectra of the $NH_2$-βA-X-RGEFV-βA-$NH_2$ libraries, where X = a) Ala, b) Arg, c) Asn, d) Asp, e) Gln, f) Glu, g) Gly, h) His, i) Ile/Leu, j) Lys, k) Met, l) Phe, m) Pro, n) Ser, o) Thr, p) Trp, q) Tyr, r) Val.

**A**

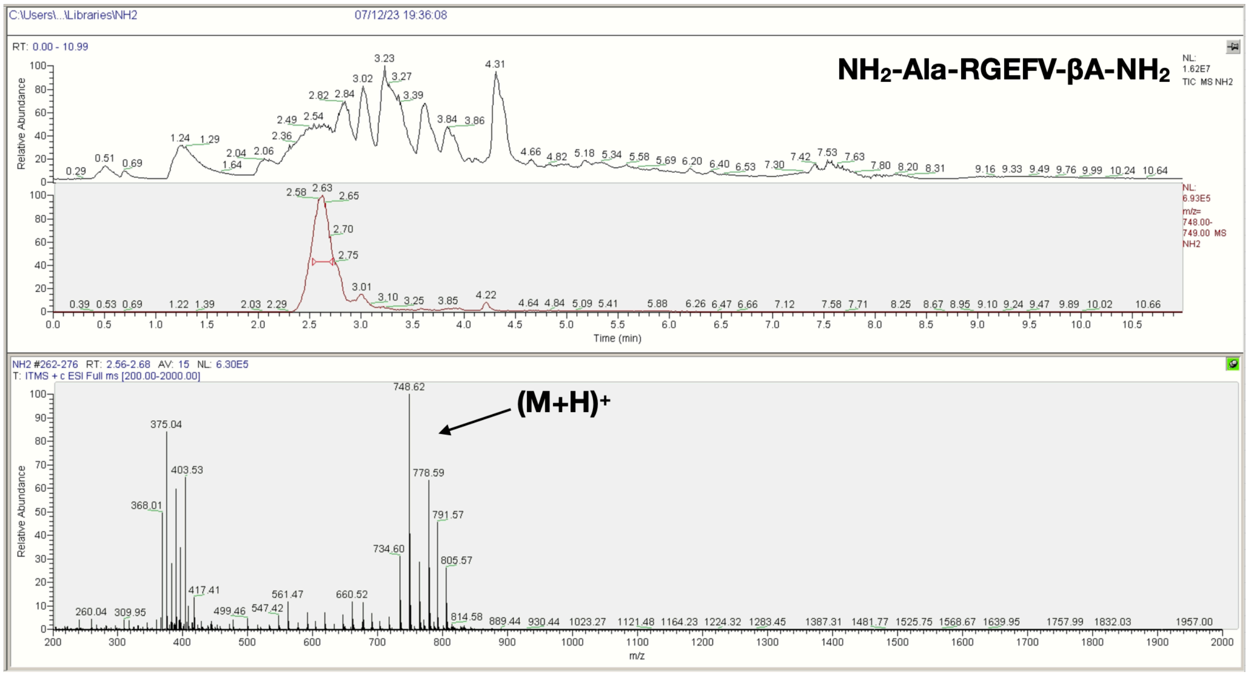

C:\Users\...\Libraries\NH2
07/12/23 19:36:08
RT: 0.00 - 10.99
NH2-Ala-RGEFV-βA-NH2
NL: 1.62E7 TIC MS NH2
NL: 6.93E5 m/z= 748.00-749.00 MS NH2
Relative Abundance
Time (min)
NH2 #262-276 RT: 2.56-2.68 AV: 15 NL: 6.30E5
T: ITMS + c ESI Full ms [200.00-2000.00]
748.62
(M+H)+
375.04
403.53
368.01
778.59
791.57
734.60
805.57
m/z


**B**

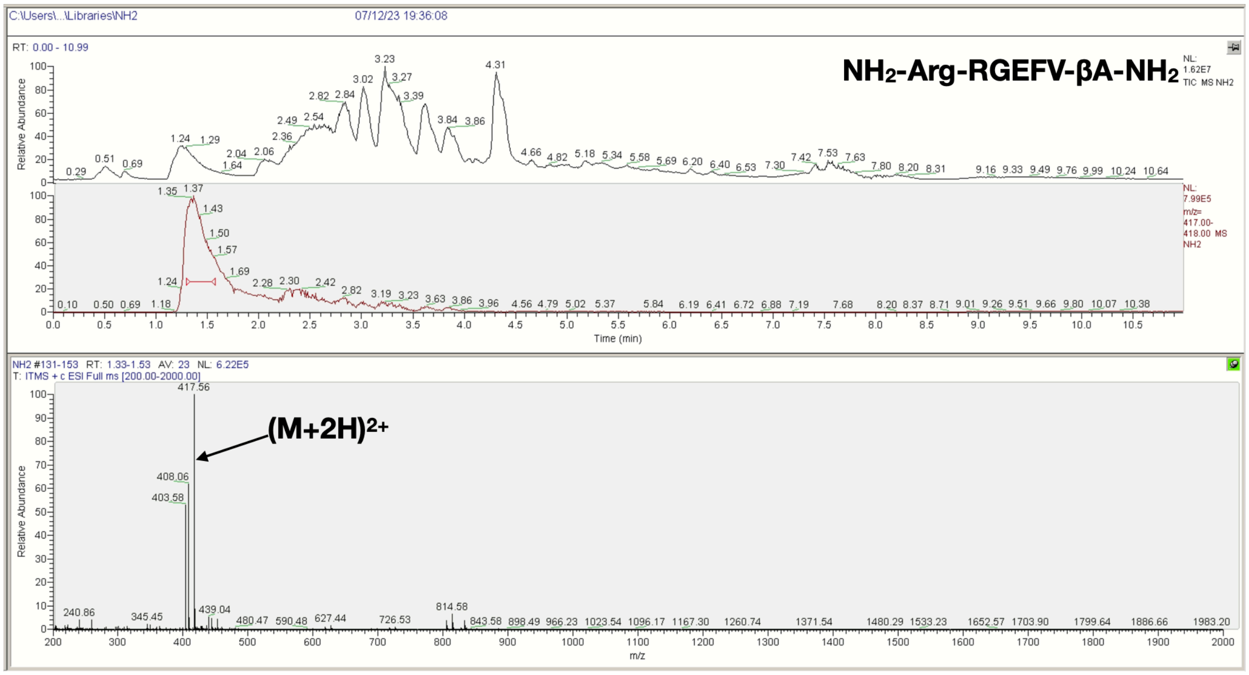

C:\Users\...\Libraries\NH2
07/12/23 19:36:08
RT: 0.00 - 10.99
NH2-Arg-RGEFV-βA-NH2
NL: 1.62E7 TIC MS NH2
NL: 7.99E5 m/z= 417.00-418.00 MS NH2
Relative Abundance
Time (min)
NH2 #131-153 RT: 1.33-1.53 AV: 23 NL: 6.22E5
T: ITMS + c ESI Full ms [200.00-2000.00]
417.56
(M+2H)2+
408.06
403.58
814.58
m/z

**C**

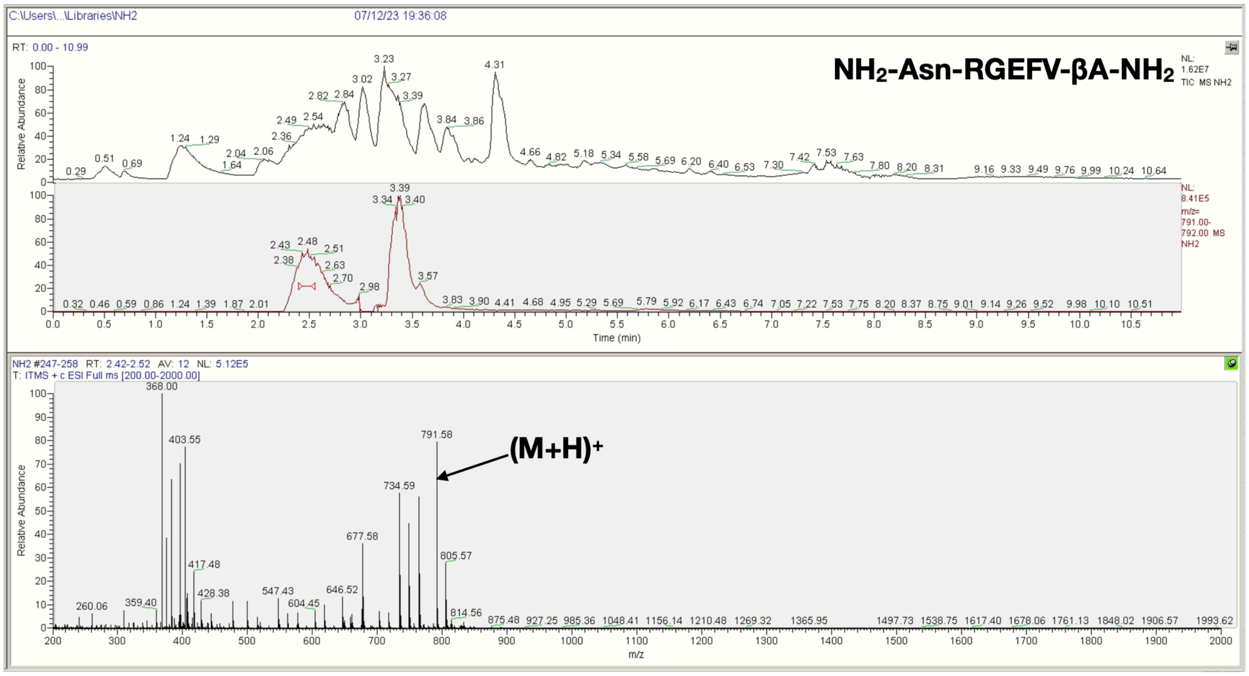
C:\Users\...\Libraries\NH2
07/12/23 19:36:08
RT: 0.00 - 10.99
NH2-Asn-RGEFV-βA-NH2
NL: 1.62E7 TIC MS NH2
NL: 8.41E5 m/z= 791.00-792.00 MS NH2
Relative Abundance
Time (min)
NH2 #247-258 RT: 2.42-2.52 AV: 12 NL: 5.12E5
T: ITMS + c ESI Full ms [200.00-2000.00]
368.00
403.55
791.58
(M+H)+
734.59
677.58
805.57
m/z

**D**

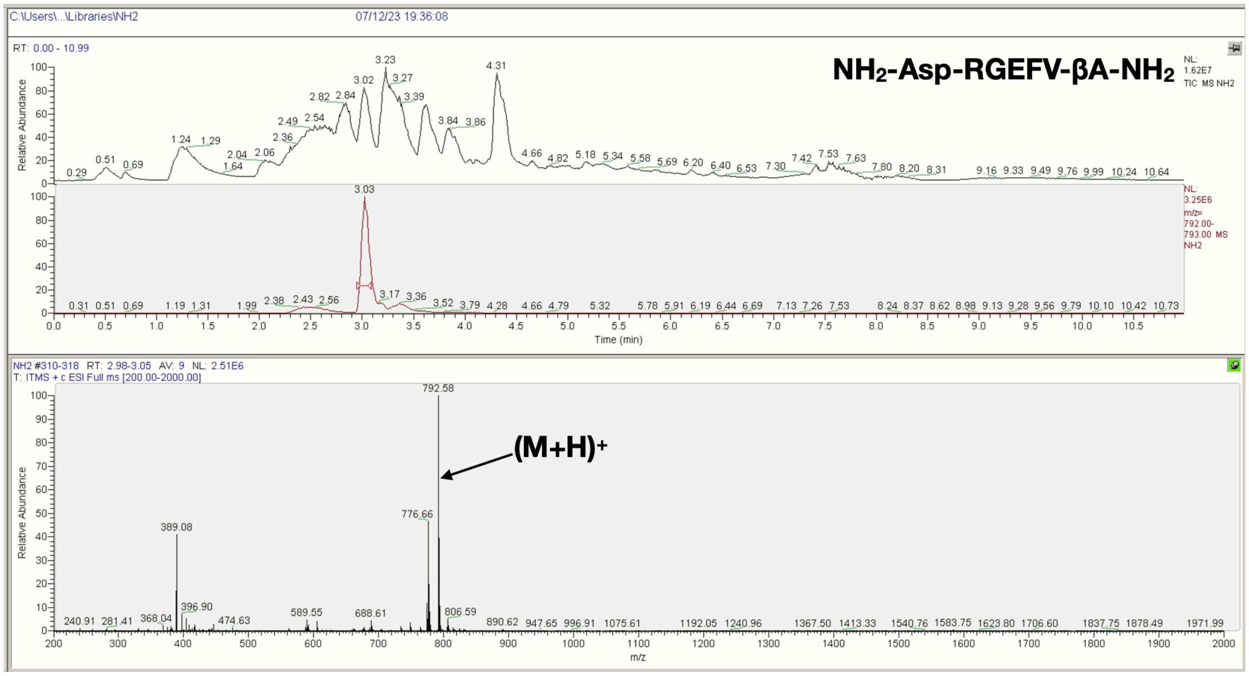
C:\Users\...\Libraries\NH2
07/12/23 19:36:08
RT: 0.00 - 10.99
NH2-Asp-RGEFV-βA-NH2
NL: 1.62E7 TIC MS NH2
NL: 3.25E6 m/z= 792.00-793.00 MS NH2
Relative Abundance
Time (min)
NH2 #310-318 RT: 2.98-3.05 AV: 9 NL: 2.51E6
T: ITMS + c ESI Full ms [200.00-2000.00]
792.58
(M+H)+
776.66
389.08
806.59
m/z

E

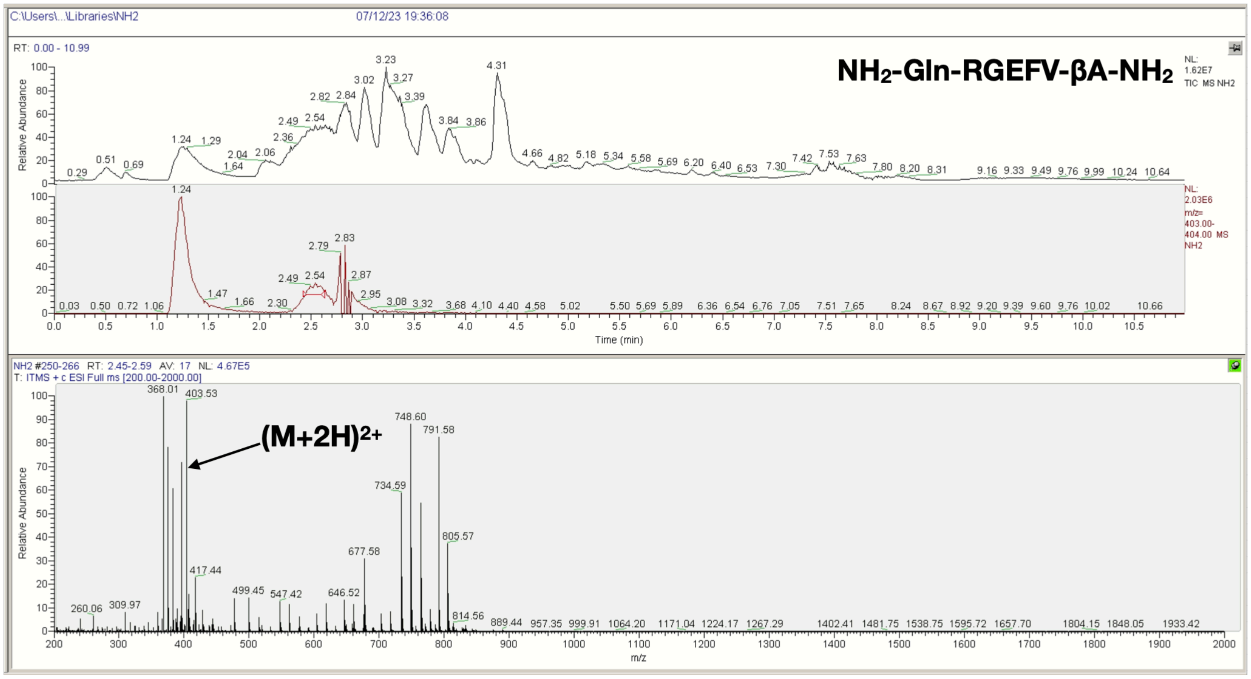
C:\Users\...\Libraries\NH2
07/12/23 19:36:08
NH2-Gln-RGEFV-βA-NH2
RT: 0.00 - 10.99
NL: 1.62E7 TIC MS NH2
NL: 2.03E6 m/z= 403.00-404.00 MS NH2
Relative Abundance
Time (min)
NH2 #250-266 RT: 2.45-2.59 AV: 17 NL: 4.67E5
T: ITMS + c ESI Full ms [200.00-2000.00]
(M+2H)2+
m/z

F

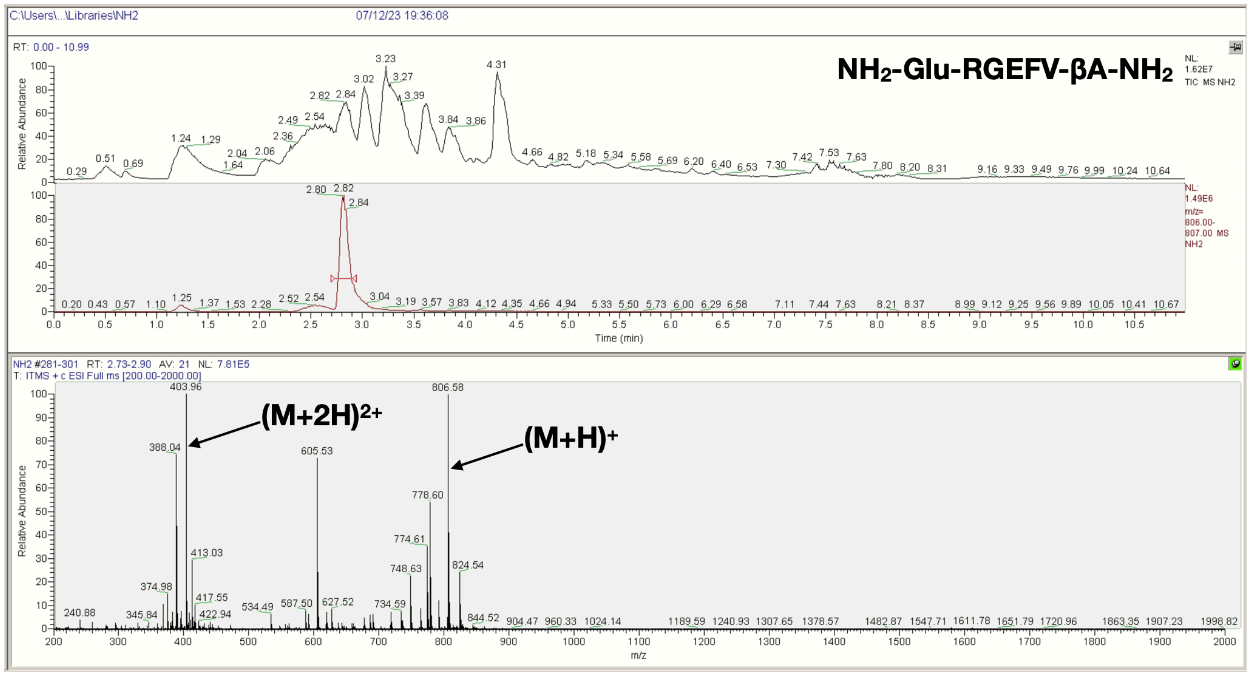
C:\Users\...\Libraries\NH2
07/12/23 19:36:08
NH2-Glu-RGEFV-βA-NH2
RT: 0.00 - 10.99
NL: 1.62E7 TIC MS NH2
NL: 1.49E6 m/z= 806.00-807.00 MS NH2
Relative Abundance
Time (min)
NH2 #281-301 RT: 2.73-2.90 AV: 21 NL: 7.81E5
T: ITMS + c ESI Full ms [200.00-2000.00]
(M+2H)2+
(M+H)+
m/z

G

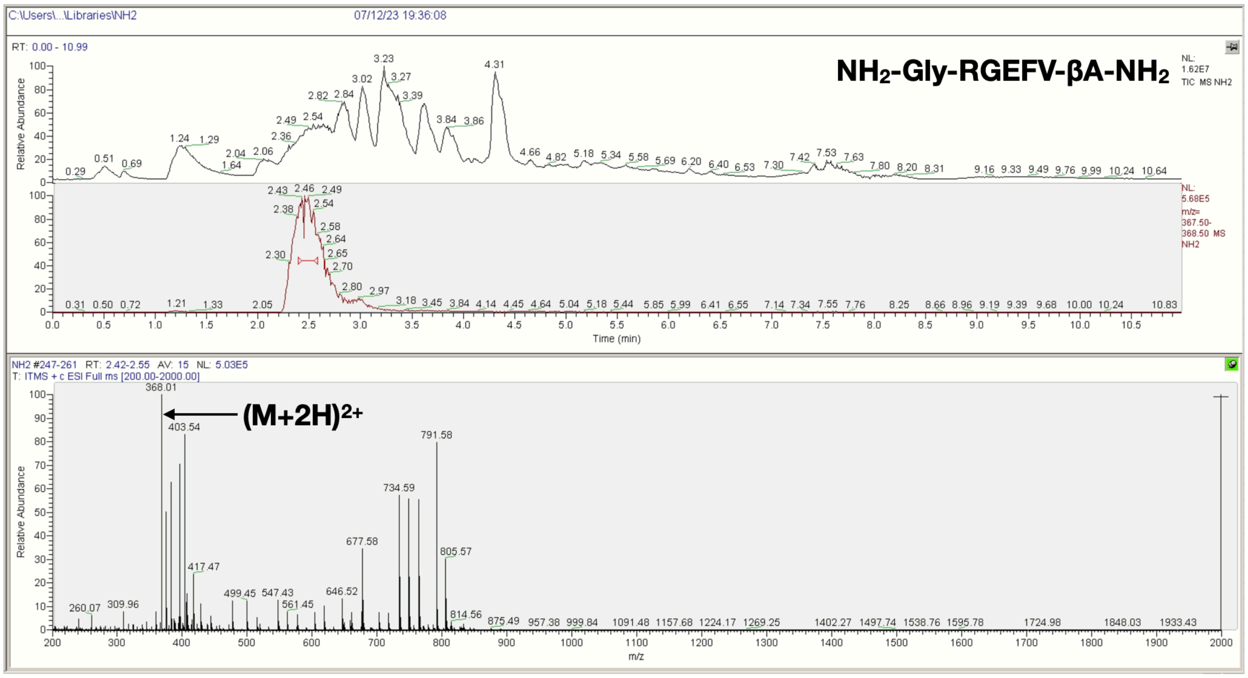


H

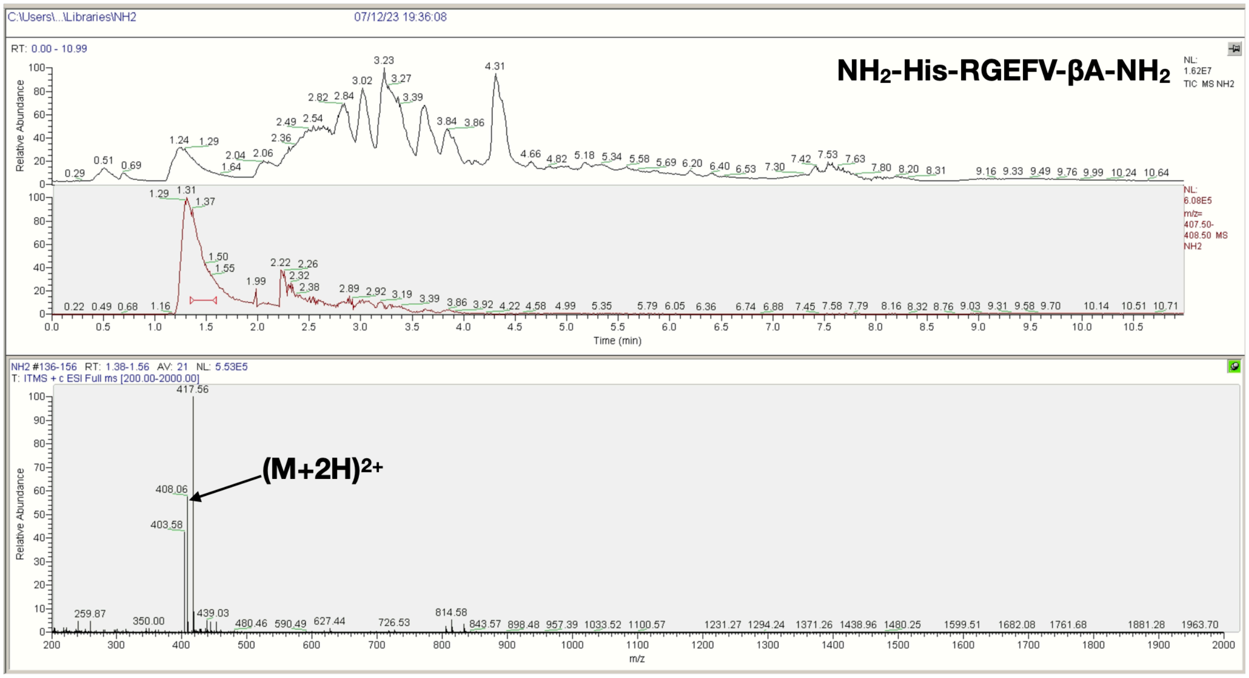

**I**

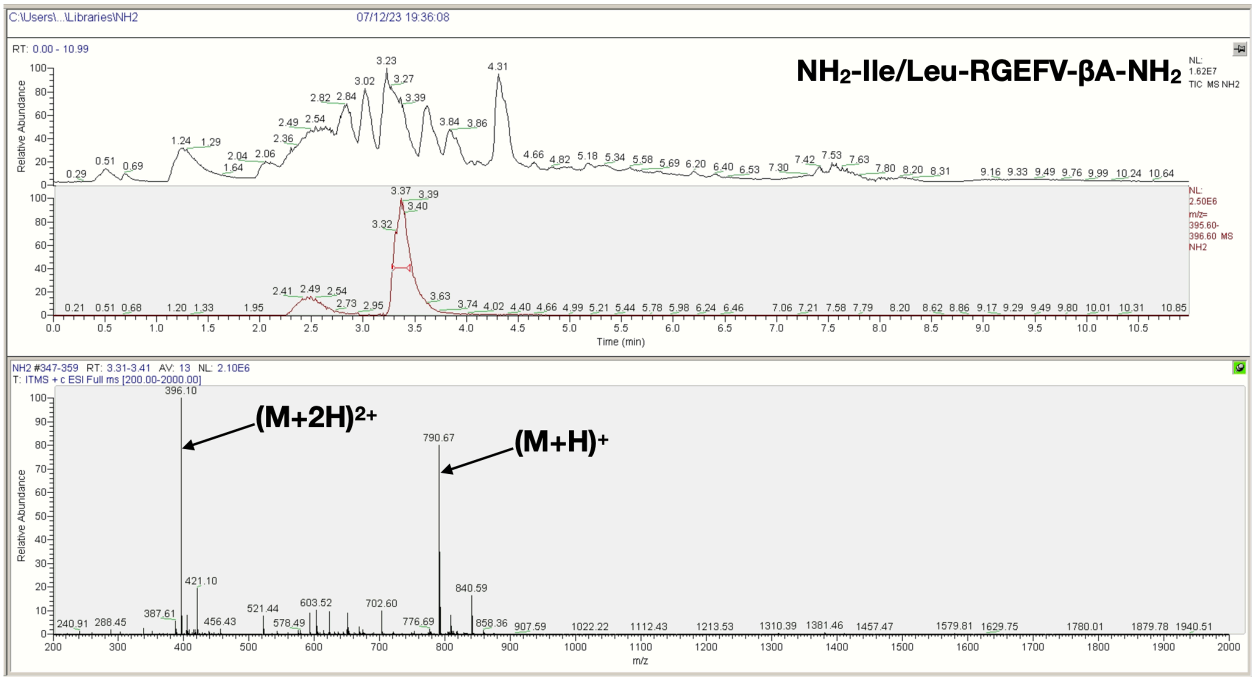
NH2-Ile/Leu-RGEFV-βA-NH2
396.10
(M+2H)2+
790.67
(M+H)+
Time (min)
m/z
Relative Abundance

**J**

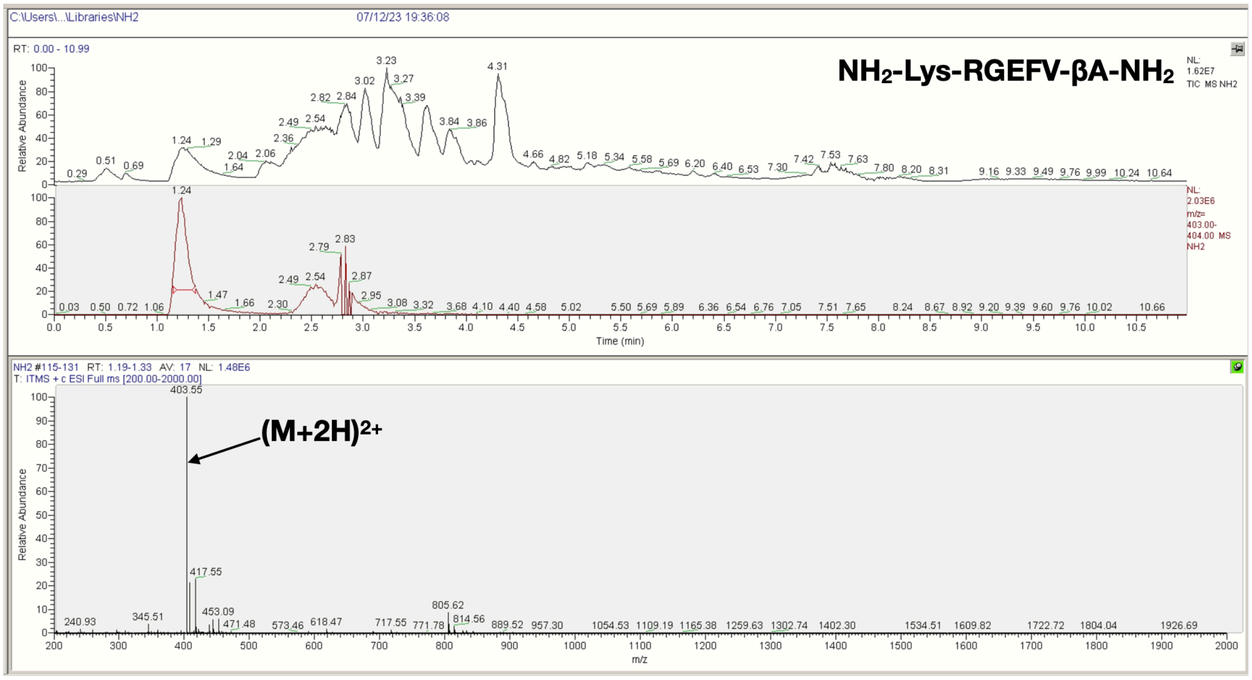
NH2-Lys-RGEFV-βA-NH2
403.55
(M+2H)2+
Time (min)
m/z
Relative Abundance

K

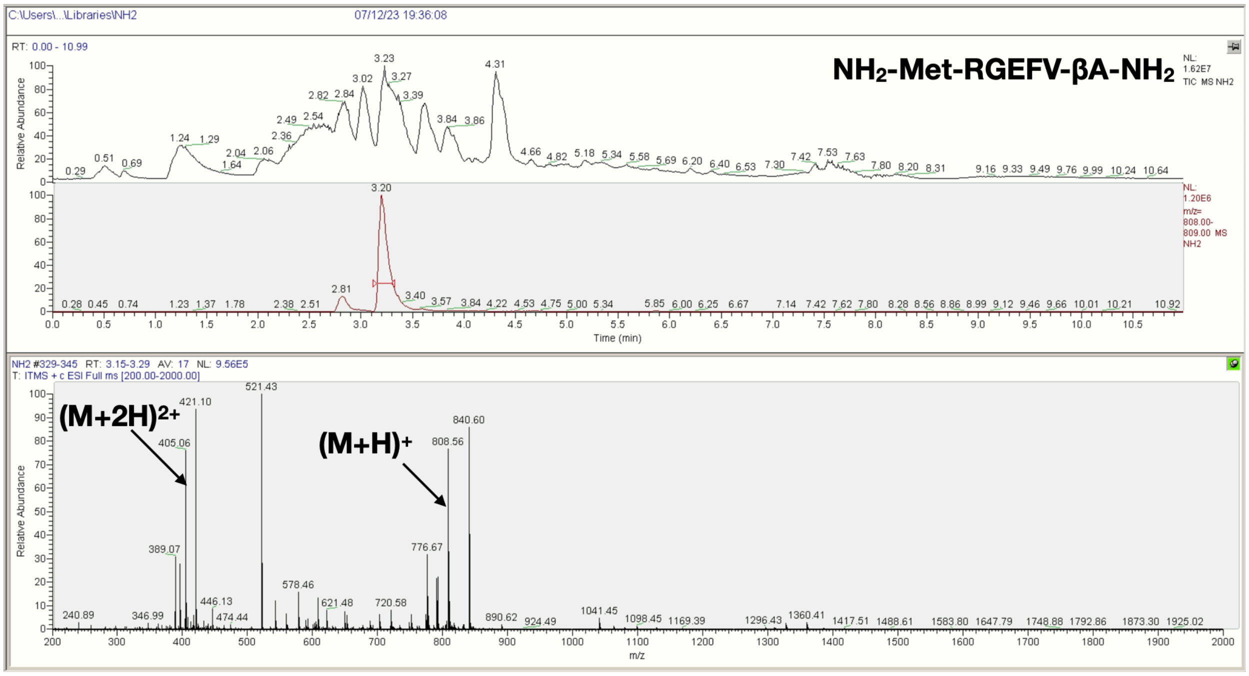
C:\Users\...\Libraries\NH2
07/12/23 19:36:08
RT: 0.00 - 10.99
NH2-Met-RGEFV-βA-NH2
NL: 1.62E7 TIC MS NH2
NL: 1.20E6 m/z= 808.00-809.00 MS NH2
Relative Abundance
Time (min)
NH2 #329-345 RT: 3.15-3.29 AV: 17 NL: 9.56E5
T: ITMS + c ESI Full ms [200.00-2000.00]
(M+2H)2+
(M+H)+
m/z

L

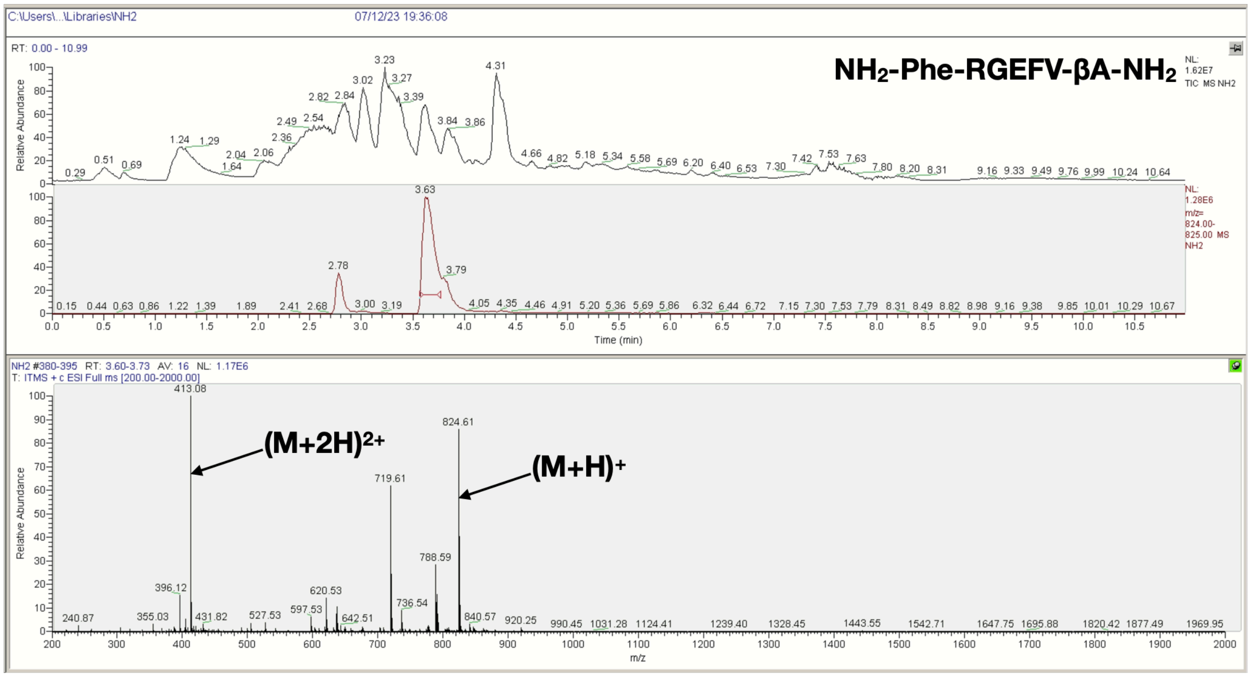
C:\Users\...\Libraries\NH2
07/12/23 19:36:08
RT: 0.00 - 10.99
NH2-Phe-RGEFV-βA-NH2
NL: 1.62E7 TIC MS NH2
NL: 1.28E6 m/z= 824.00-825.00 MS NH2
Relative Abundance
Time (min)
NH2 #380-395 RT: 3.60-3.73 AV: 16 NL: 1.17E6
T: ITMS + c ESI Full ms [200.00-2000.00]
(M+2H)2+
(M+H)+
m/z

**M**

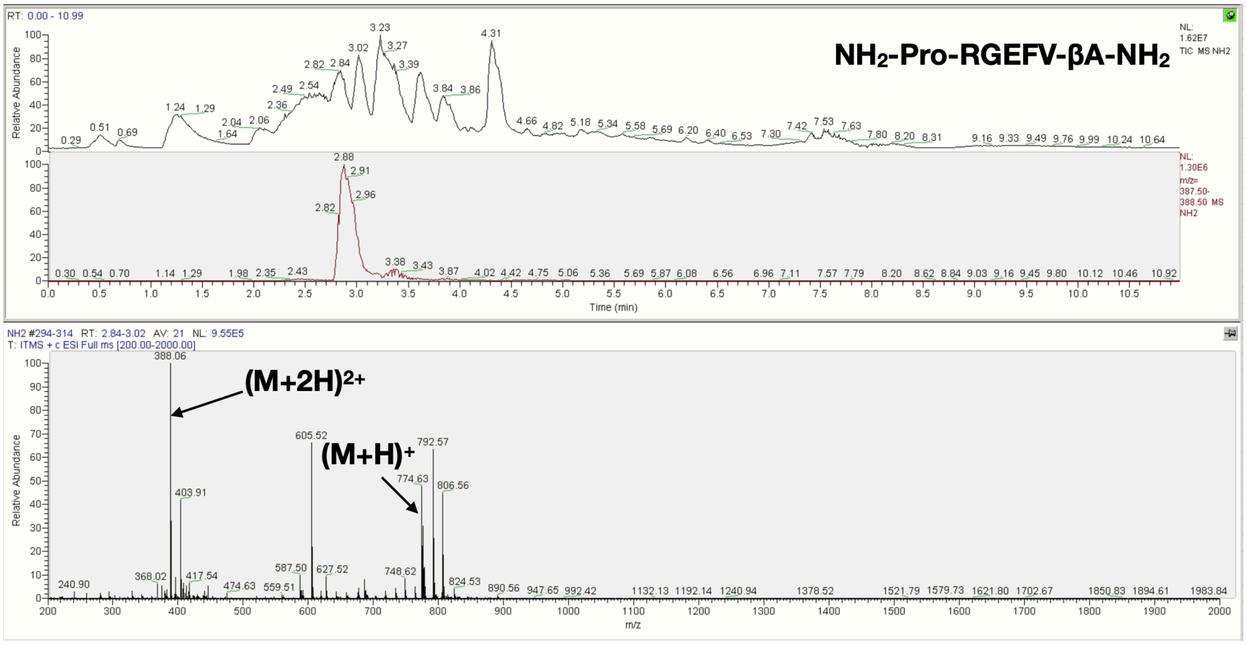
RT: 0.00 - 10.99
NH2-Pro-RGEFV-βA-NH2
NL: 1.62E7 TIC MS NH2
NL: 1.30E6 m/z= 387.50-388.50 MS NH2
Relative Abundance
Time (min)
NH2 #294-314 RT: 2.84-3.02 AV: 21 NL: 9.55E5
T: ITMS + c ESI Full ms [200.00-2000.00]
(M+2H)2+
(M+H)+
388.06
605.52
792.57
774.63
806.56
403.91
m/z

**N**

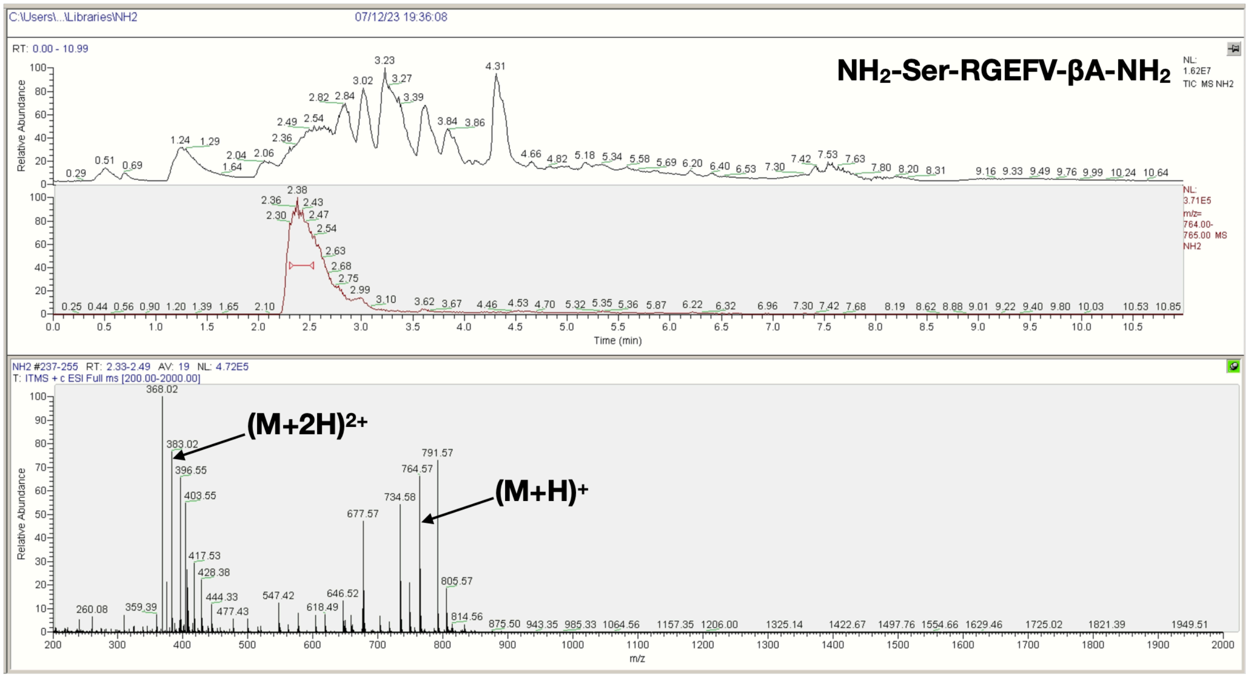
C:\Users\...\Libraries\NH2
07/12/23 19:36:08
RT: 0.00 - 10.99
NH2-Ser-RGEFV-βA-NH2
NL: 1.62E7 TIC MS NH2
NL: 3.71E5 m/z= 764.00-765.00 MS NH2
Relative Abundance
Time (min)
NH2 #237-255 RT: 2.33-2.49 AV: 19 NL: 4.72E5
T: ITMS + c ESI Full ms [200.00-2000.00]
(M+2H)2+
(M+H)+
368.02
383.02
396.55
403.55
677.57
734.58
764.57
791.57
805.57
m/z

O

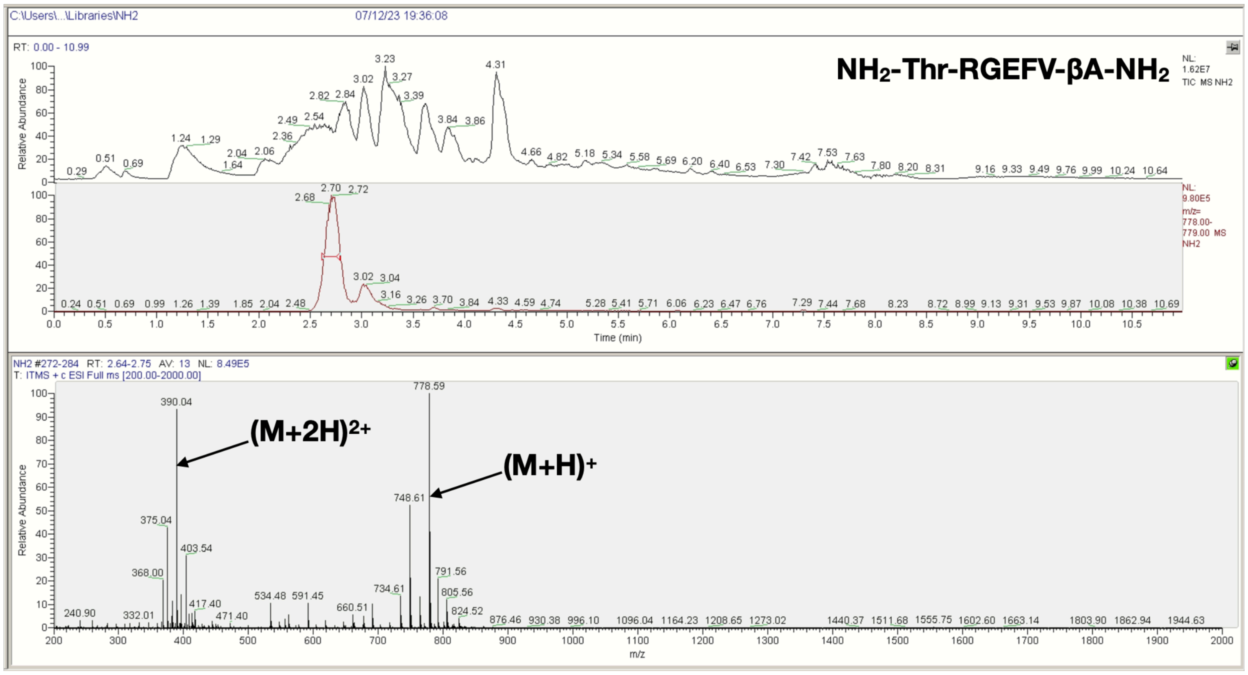
C:\Users\...\Libraries\NH2
07/12/23 19:36:08
RT: 0.00 - 10.99
NH2-Thr-RGEFV-βA-NH2
NL: 1.62E7 TIC MS NH2
NL: 9.80E5 m/z= 778.00-779.00 MS NH2
Relative Abundance
Time (min)
NH2 #272-284 RT: 2.64-2.75 AV: 13 NL: 8.49E5
T: ITMS + c ESI Full ms [200.00-2000.00]
390.04
778.59
748.61
375.04
403.54
368.00
791.56
805.56
734.61
(M+2H)2+
(M+H)+
m/z

P

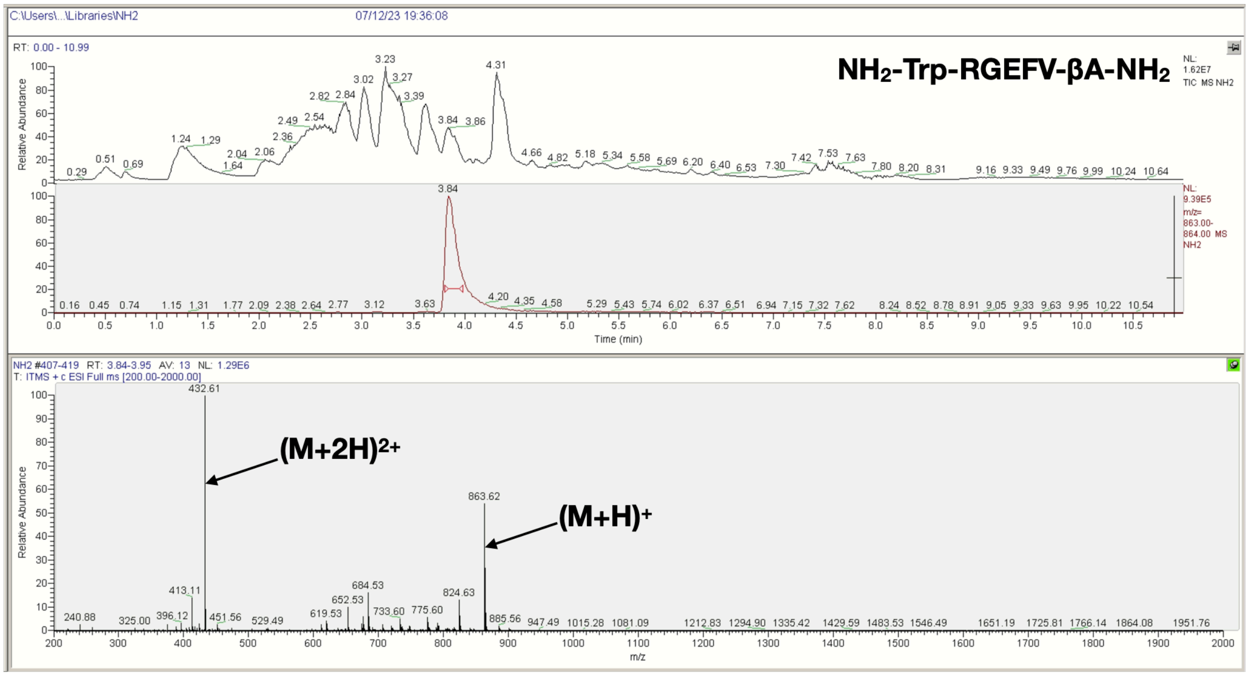
C:\Users\...\Libraries\NH2
07/12/23 19:36:08
RT: 0.00 - 10.99
NH2-Trp-RGEFV-βA-NH2
NL: 1.62E7 TIC MS NH2
NL: 9.39E5 m/z= 863.00-864.00 MS NH2
Relative Abundance
Time (min)
NH2 #407-419 RT: 3.84-3.95 AV: 13 NL: 1.29E6
T: ITMS + c ESI Full ms [200.00-2000.00]
432.61
863.62
413.11
684.53
652.53
824.63
(M+2H)2+
(M+H)+
m/z

Q

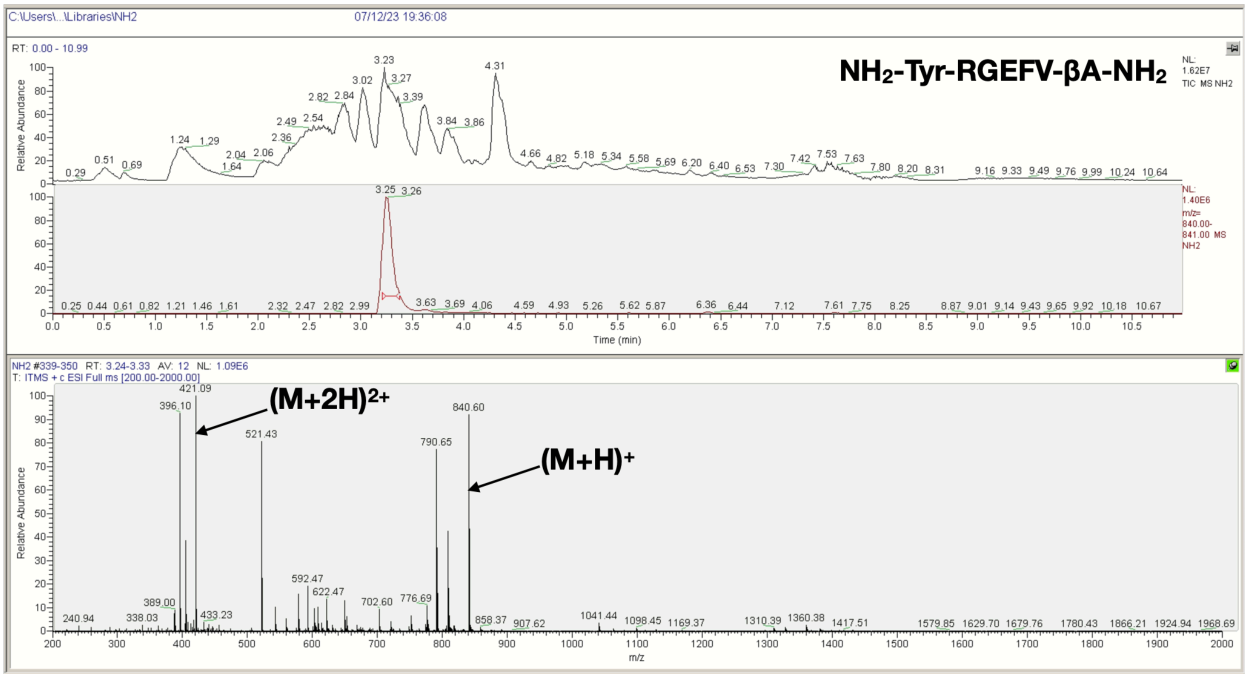


R

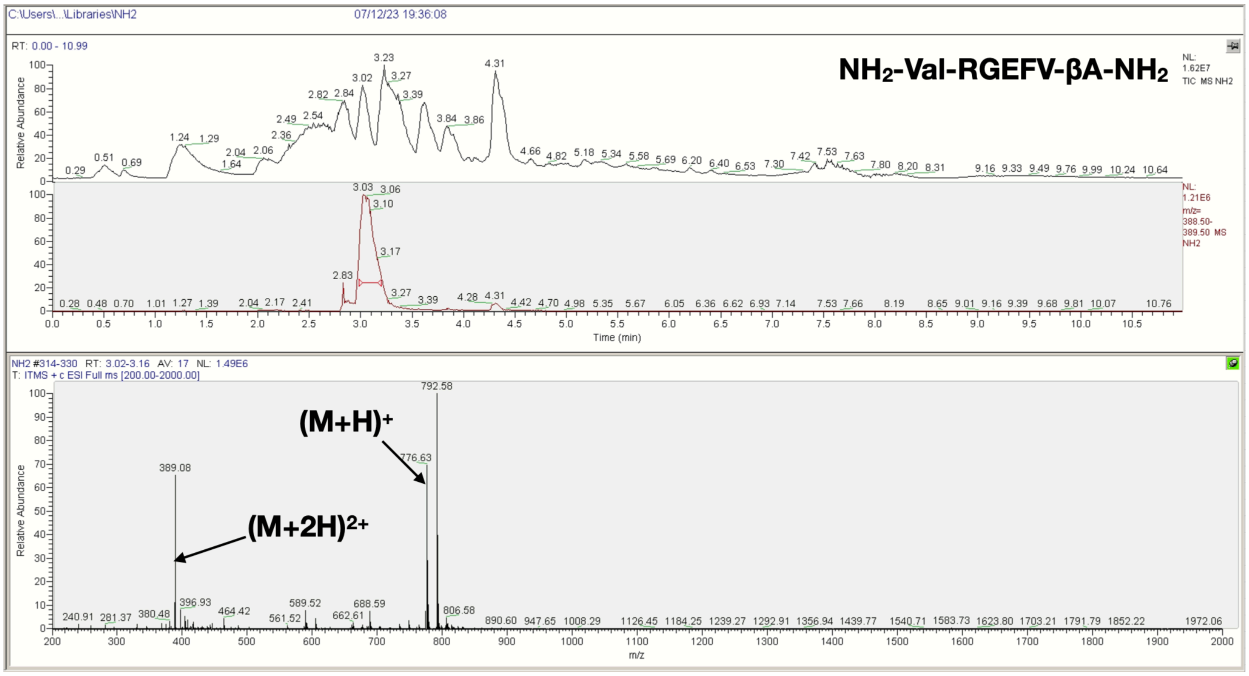


**Figure S18**. LCMS spectra of the $NH_2$-X-RGEFV-βA-$NH_2$ libraries, where X = a) Ala, b) Arg, c) Asn, d) Asp, e) Gln, f) Glu, g) Gly, h) His, i) Ile/Leu, j) Lys, k) Met, l) Phe, m) Pro, n) Ser, o) Thr, p) Trp, q) Tyr, r) Val.

### A Ac-βA-Gly-RGEFV-βA-Am

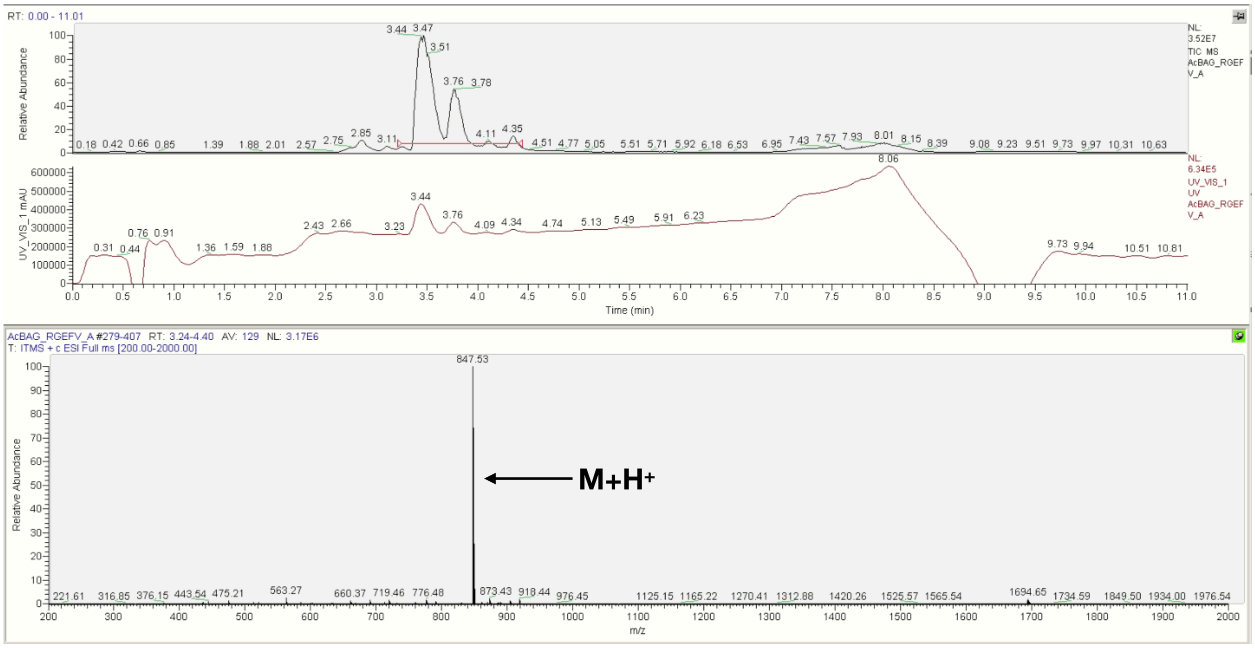


### B Ac-Gly-RGEFV-βA-Am

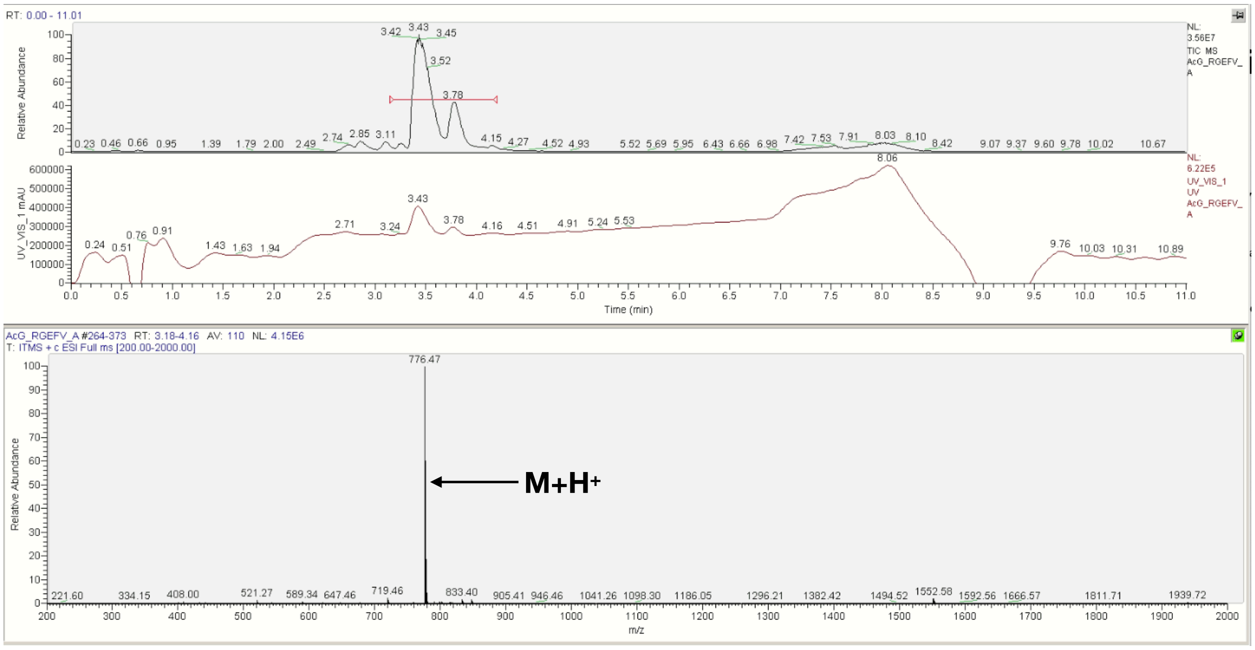

**C NH$_2$-βA-Gly-RGEFV-βA-Am**

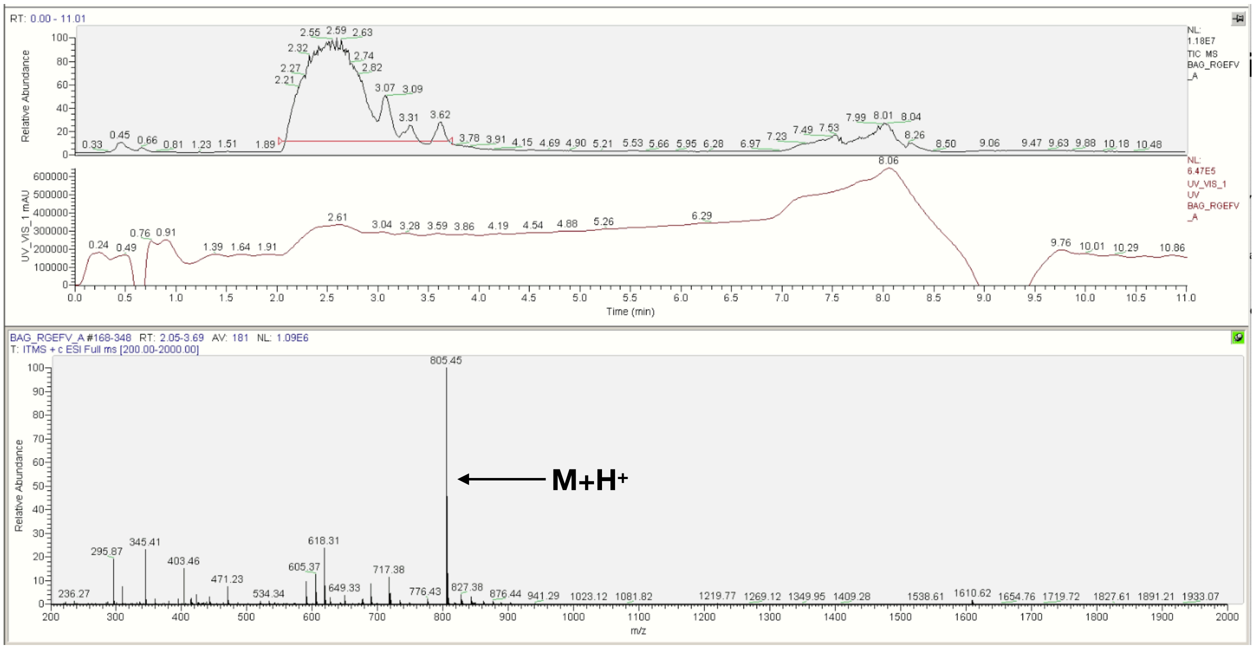


**D NH$_2$-Gly-RGEFV-βA-Am**

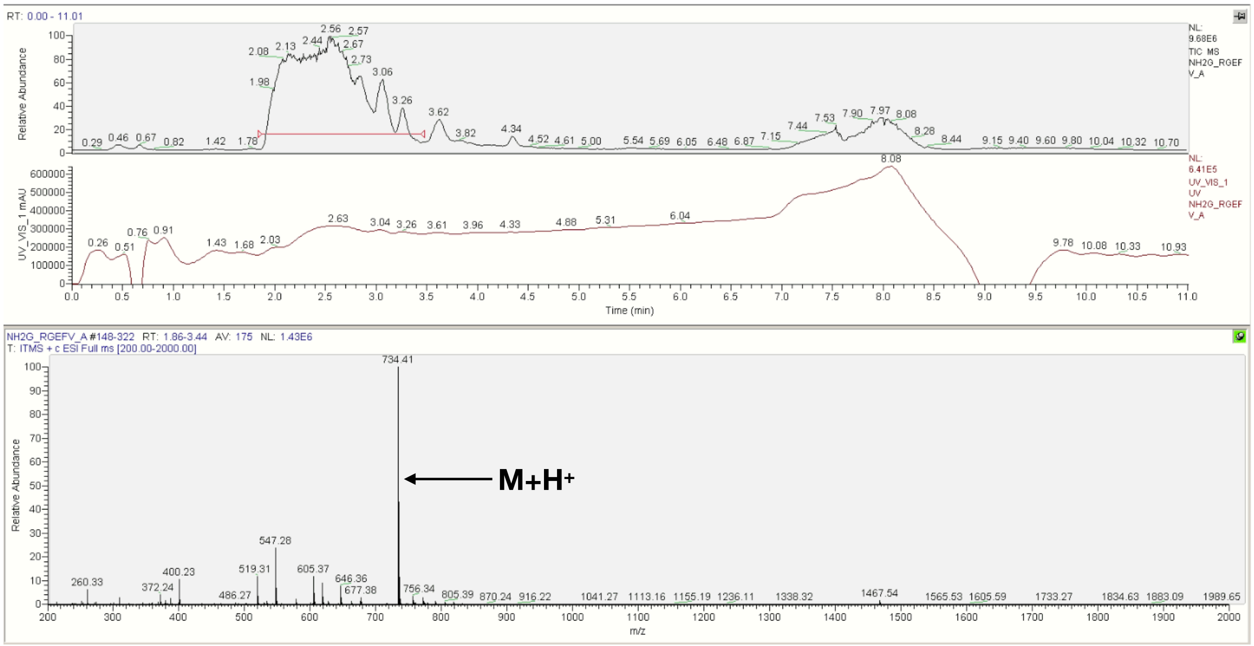

**E Ac-βA-RGEFV-Gly-βA-Am**

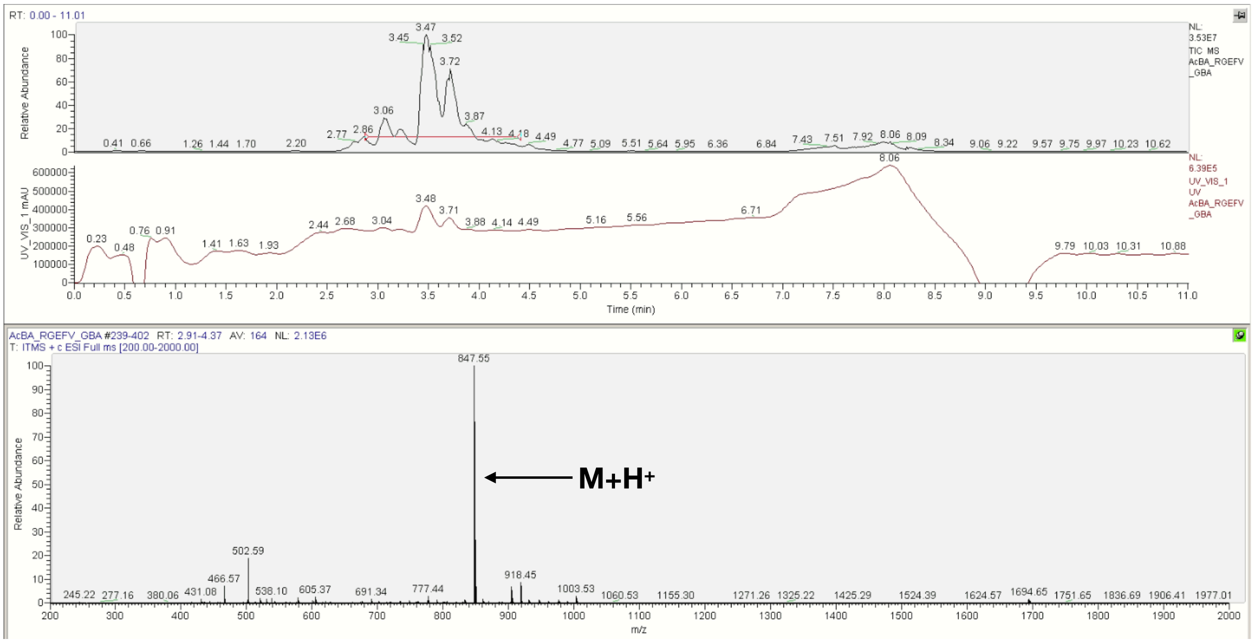


**F Ac-βA-RGEFV-Gly-Am**

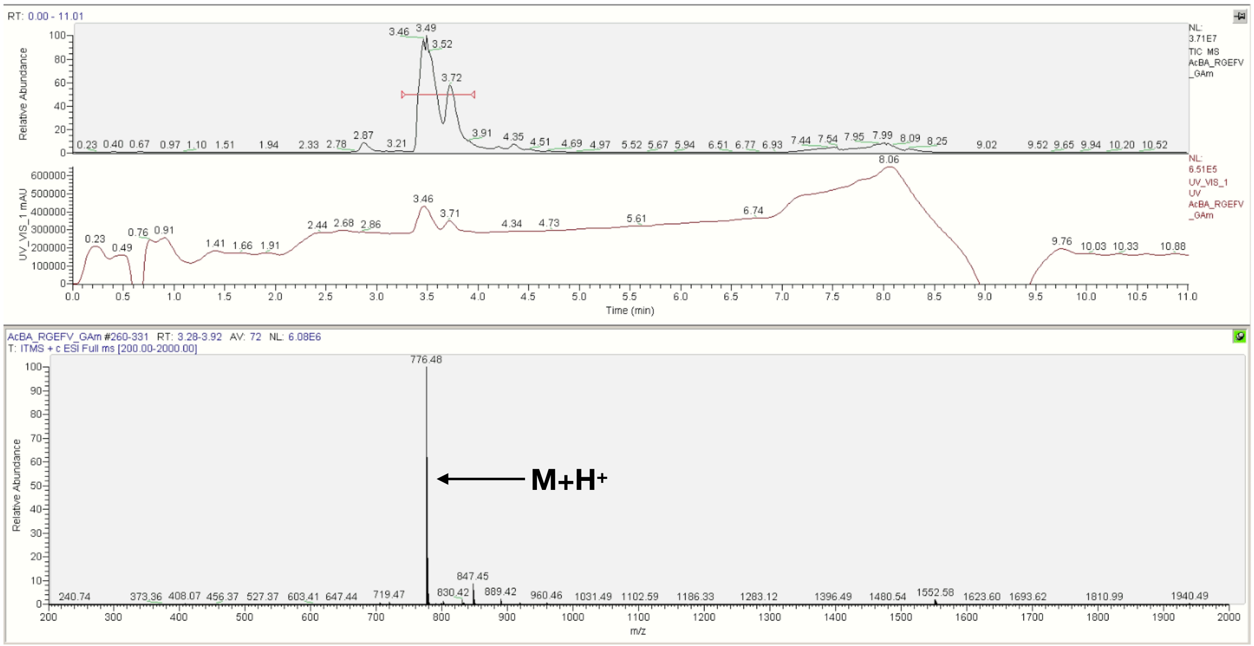

### G Ac-βA-RGEFV-Gly-COOH

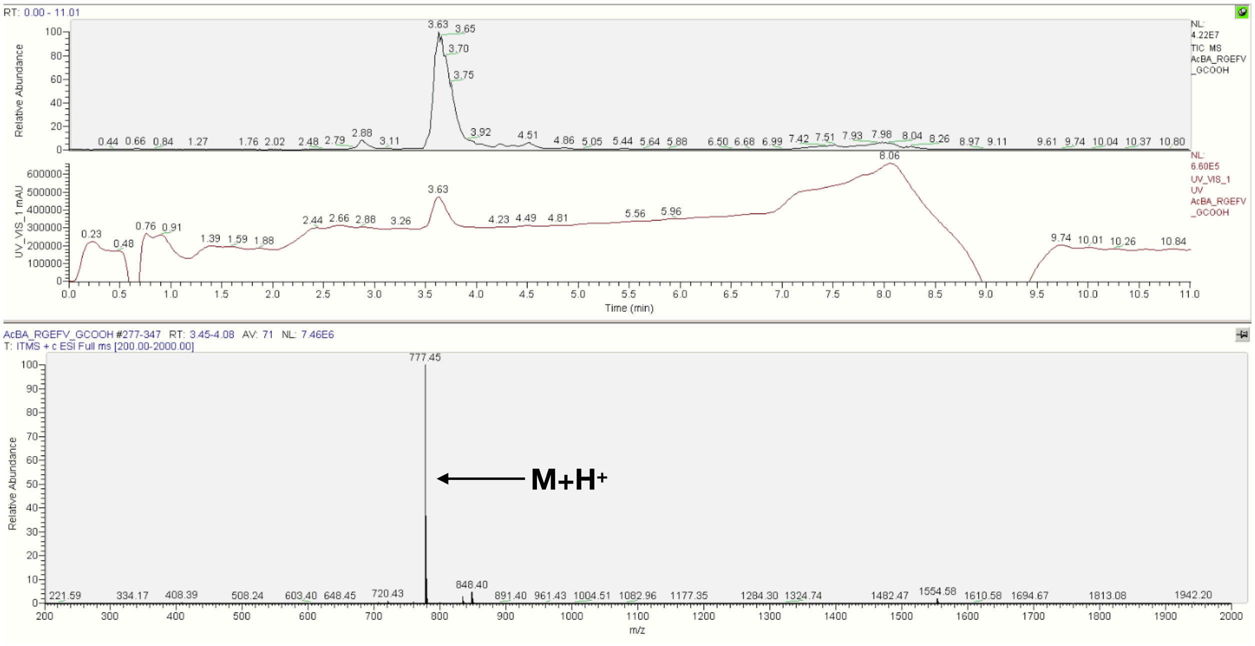


**Figure S19**. LCMS spectra of the peptides used for the concentration studies in which a glycine was placed on the N-terminus for N-terminal libraries, and the C-terminus for C-terminal libraries. a) Ac-βA, b) Ac, c) N-βA, d) $NH_2$, e) C-βA, f) Am, g) COOH.

### A Ac-βA-Gly-LIAANK-βA-Am

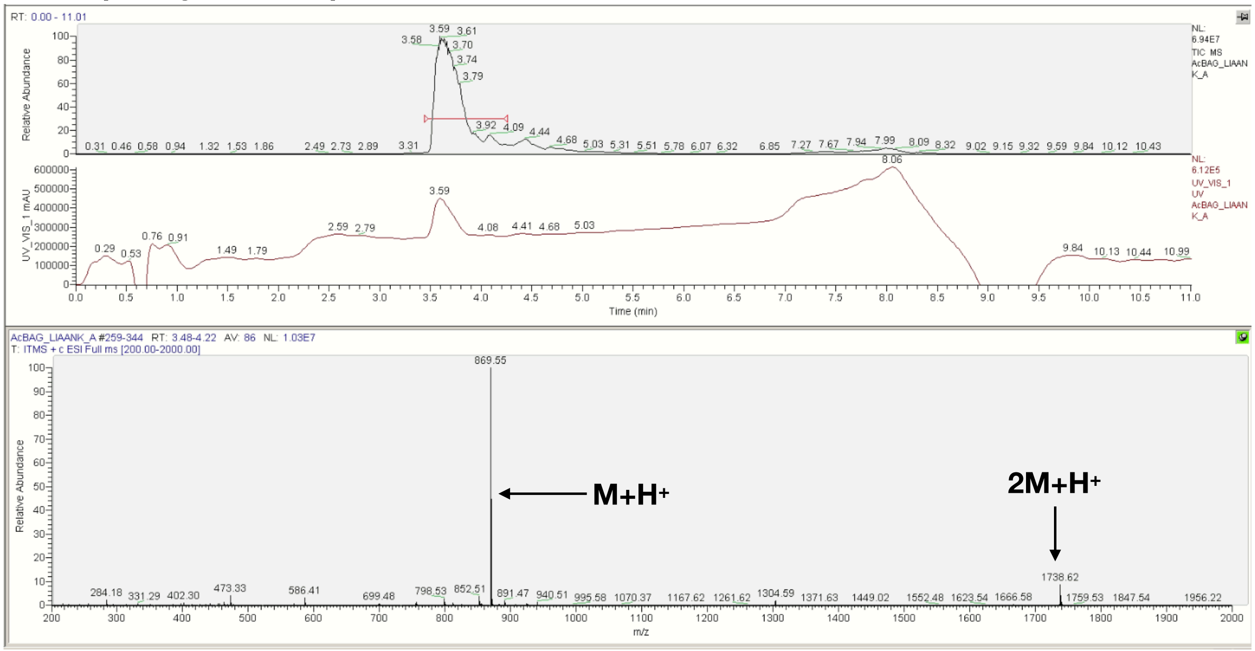

**B Ac-Gly-LIAANK-βA-Am**

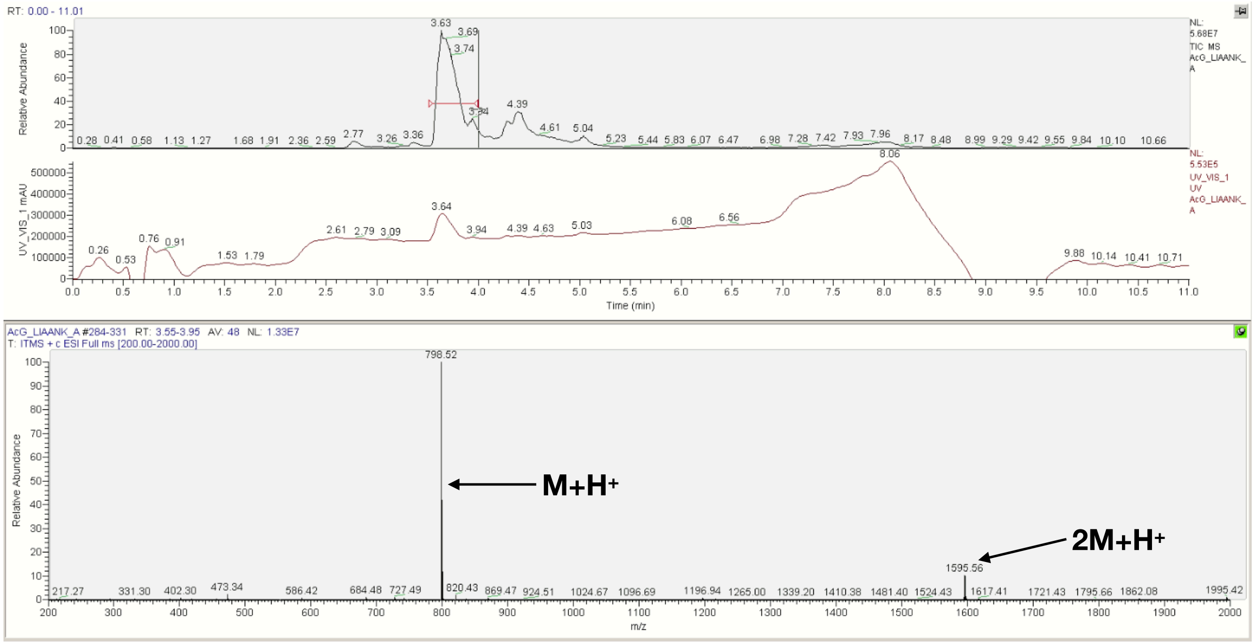


**C NH$_2$-βA-Gly-LIAANK-βA-Am**

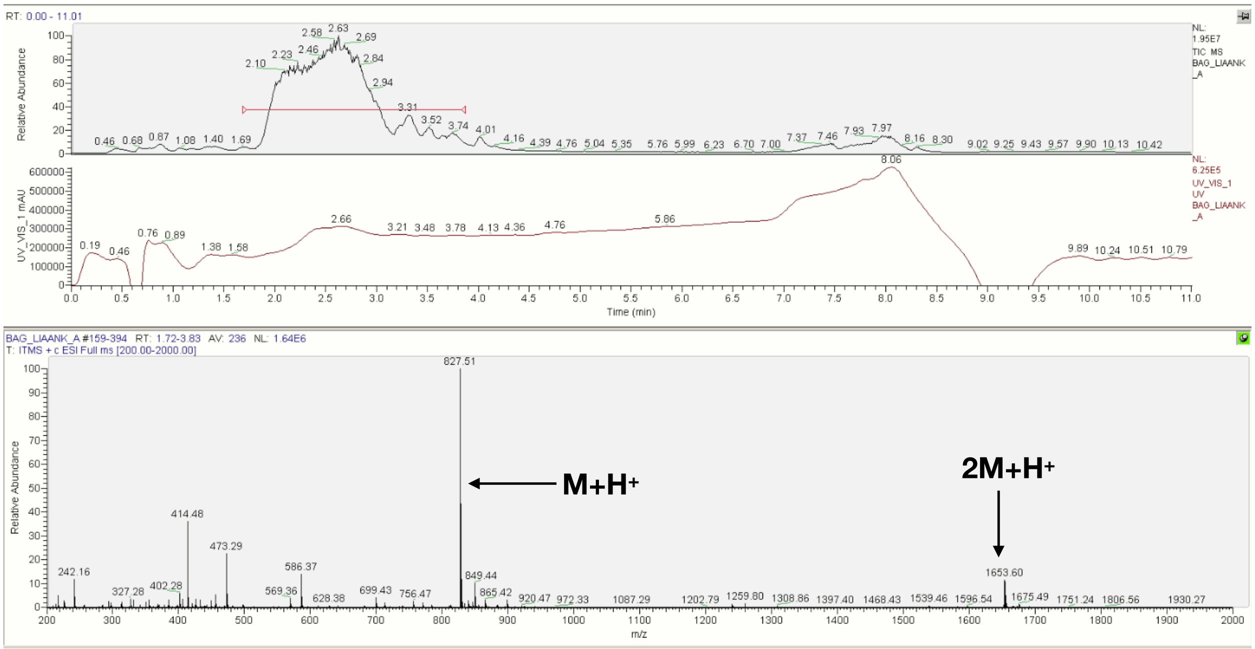

**D NH$_2$-Gly-LIAANK-βA-Am**

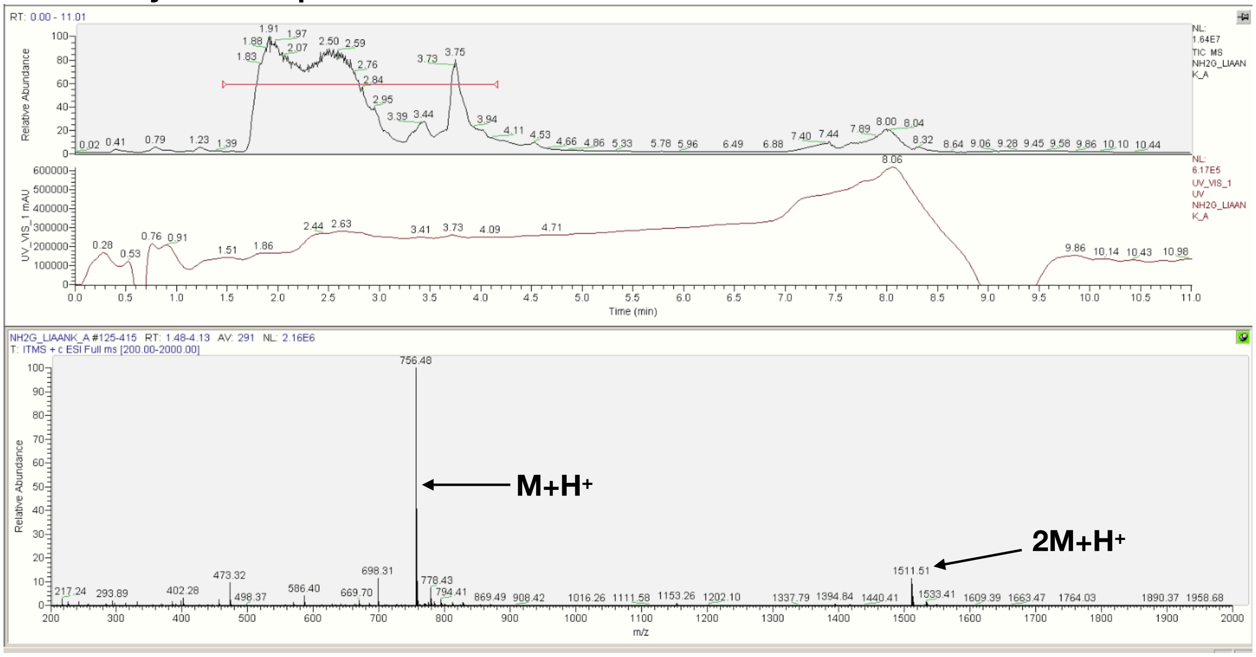


**E Ac-βA-LIAANK-Gly-βA-Am**

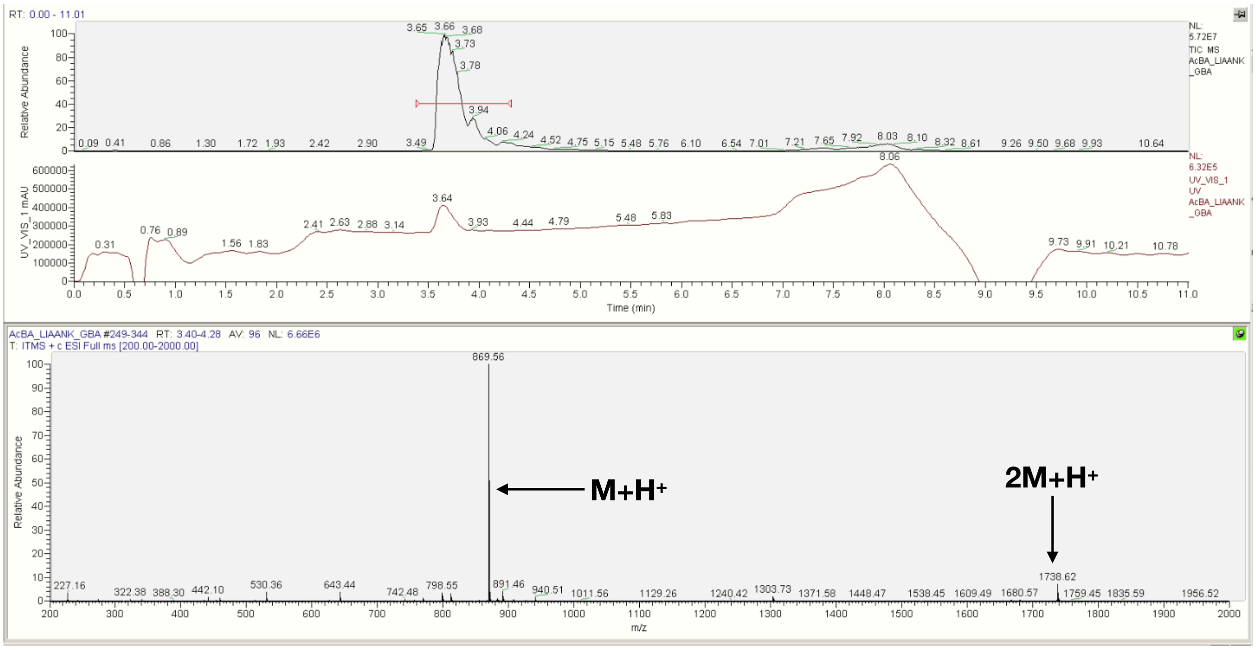

**F Ac-βA-LIAANK-Gly-Am**

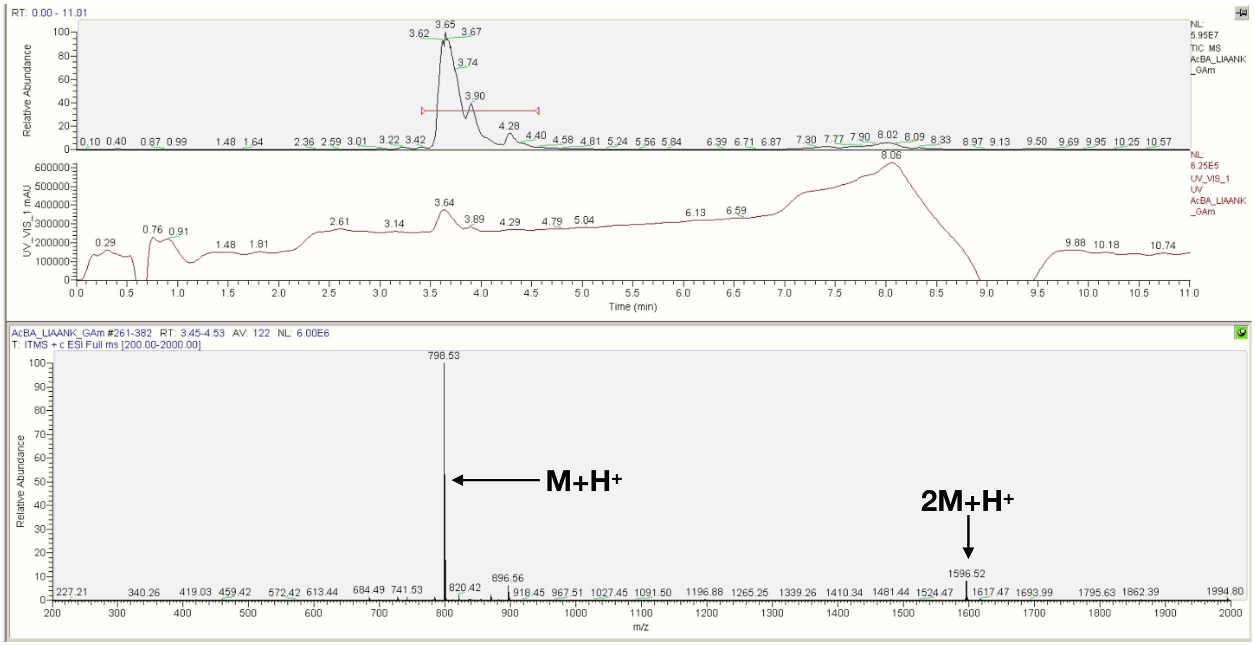


**G Ac-βA-LIAANK-Gly-COOH**

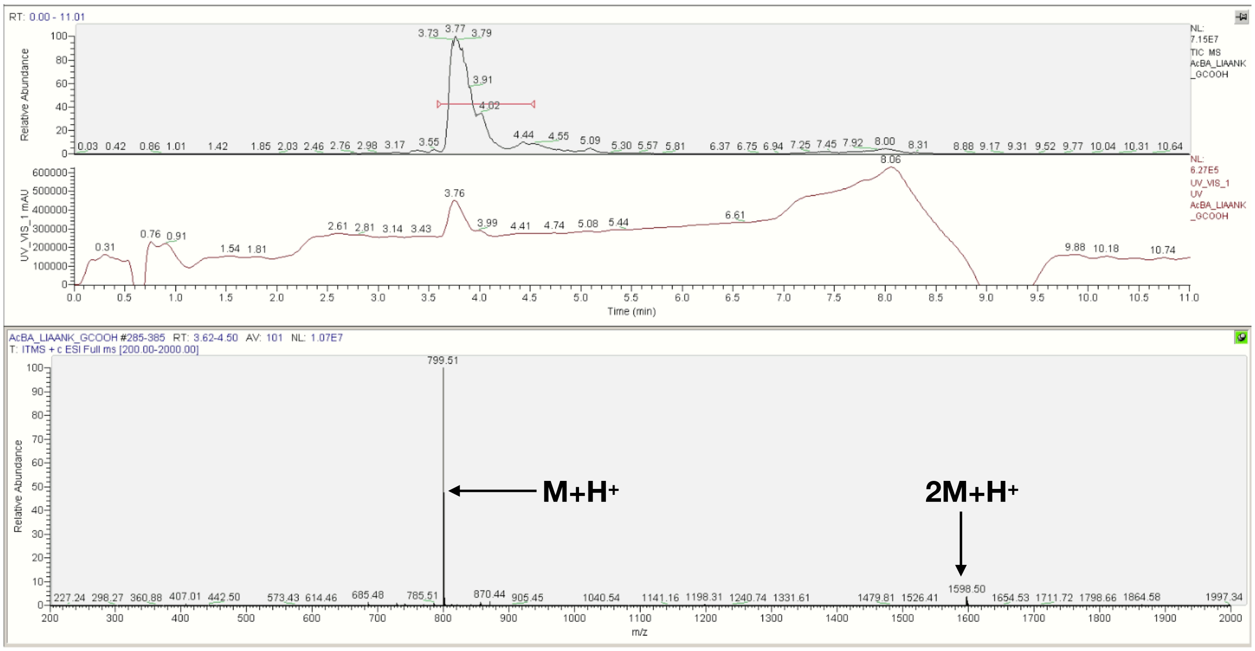


**Figure S20**. LCMS spectra of the LIAANK peptides in which a glycine was placed on the N-terminus for N-terminal libraries, and the C-terminus for C-terminal libraries. a) Ac-βA, b) Ac, c) N-βA, d) $NH_2$, e) C-βA, f) Am, g) COOH.

**A Ac-βA-Gly-IVKVA-βA-Am**

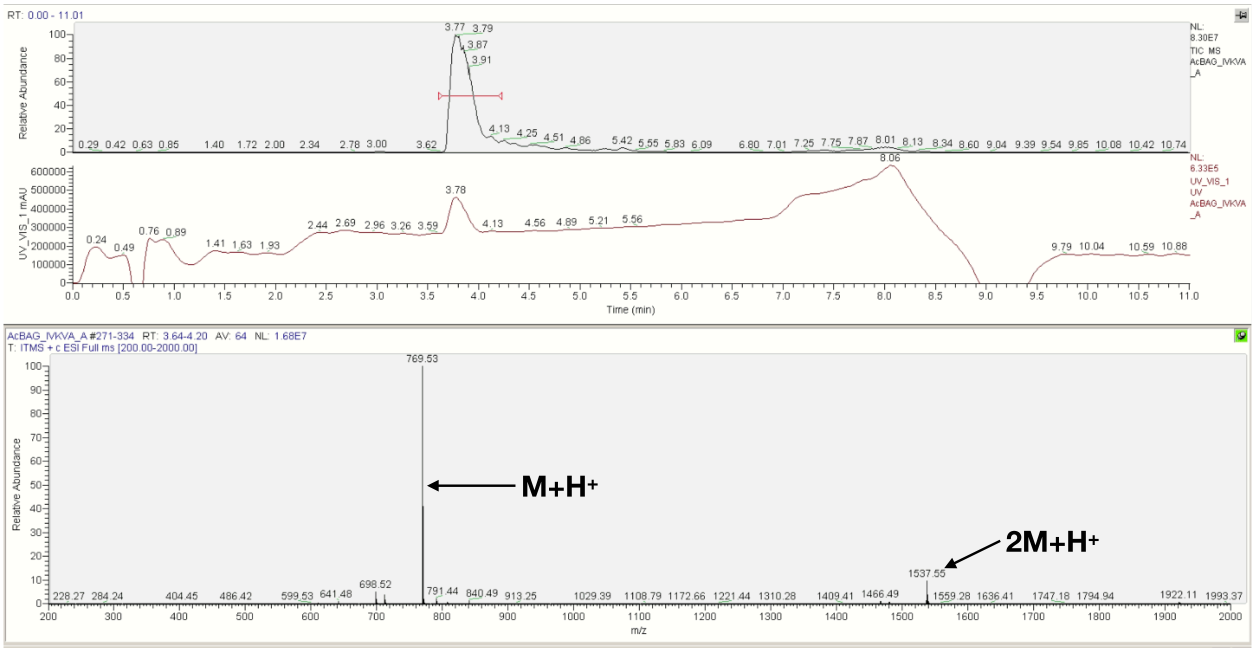


**B Ac-Gly-IVKVA-βA-Am**

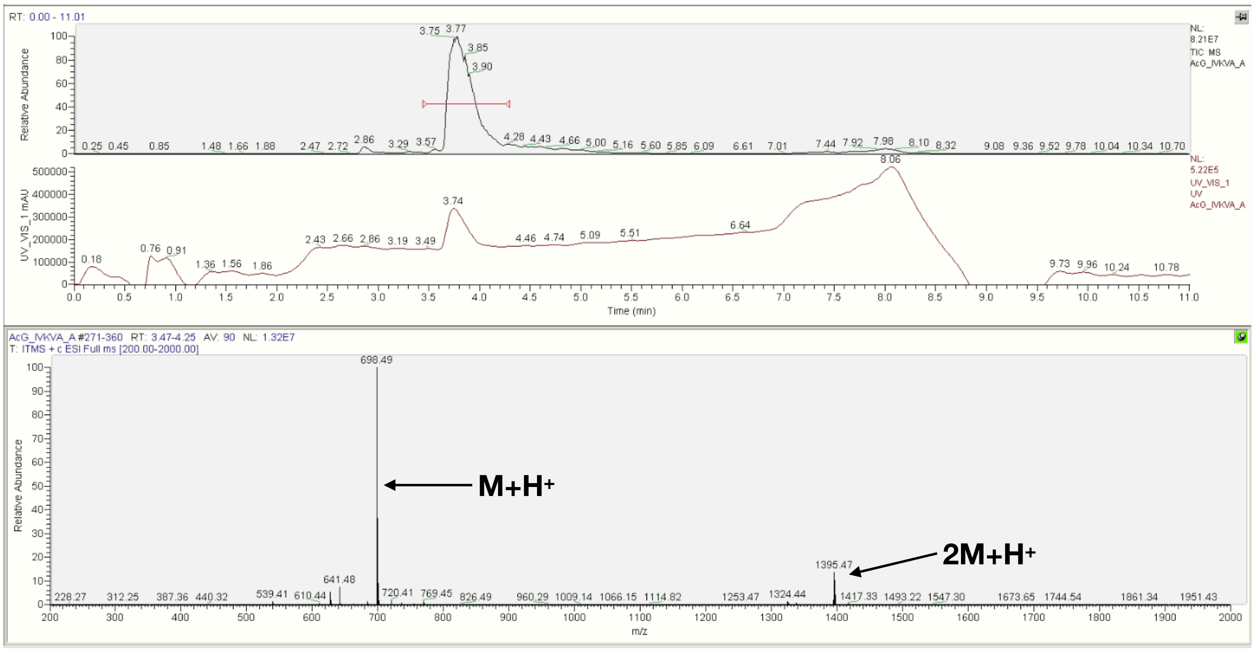

### C NH$_2$-βA-Gly-IVKVA-βA-Am

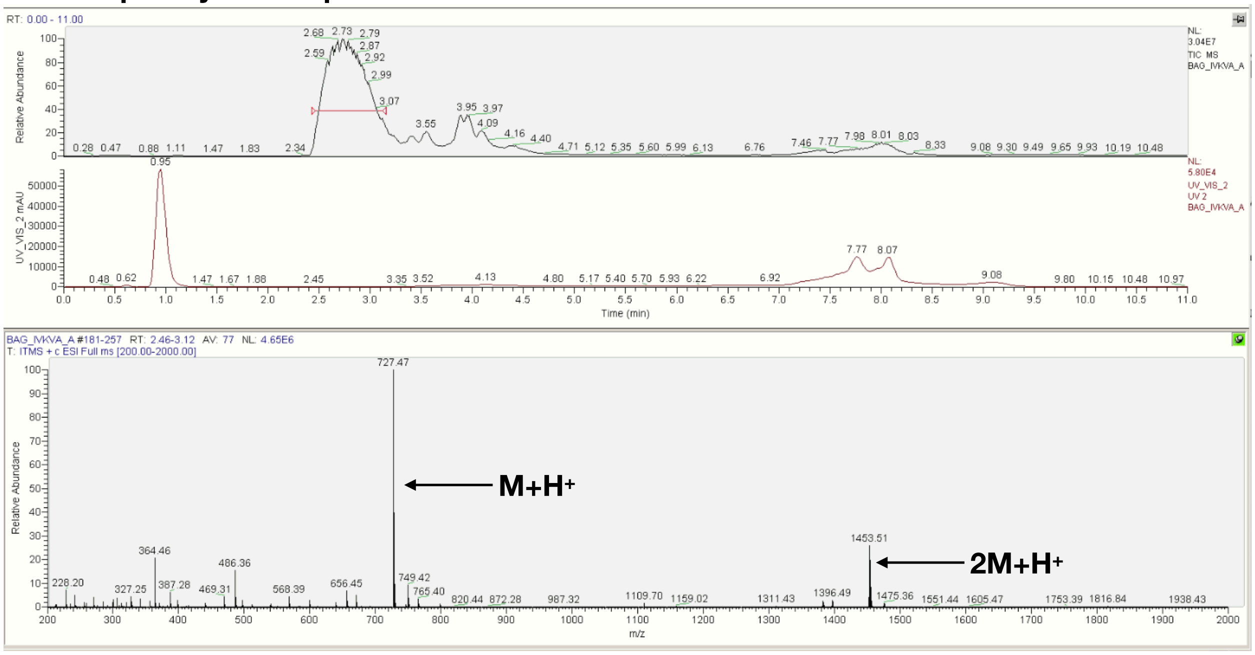


### D NH$_2$-Gly-IVKVA-βA-Am

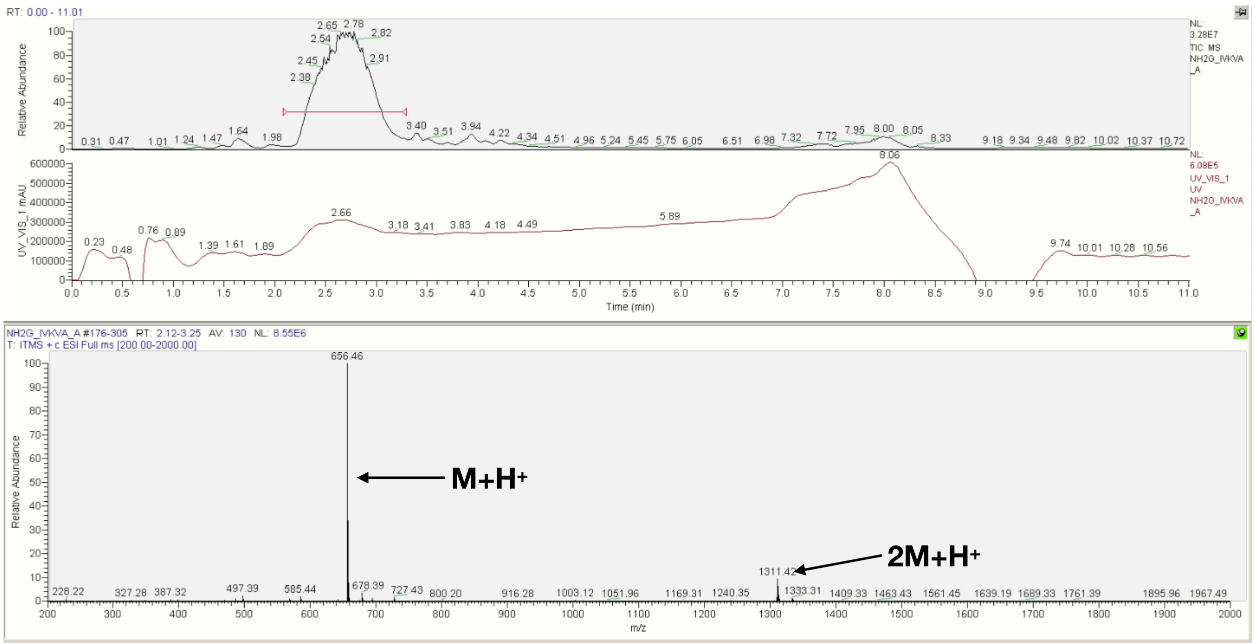

### E Ac-βA-IVKVA-Gly-βA-Am

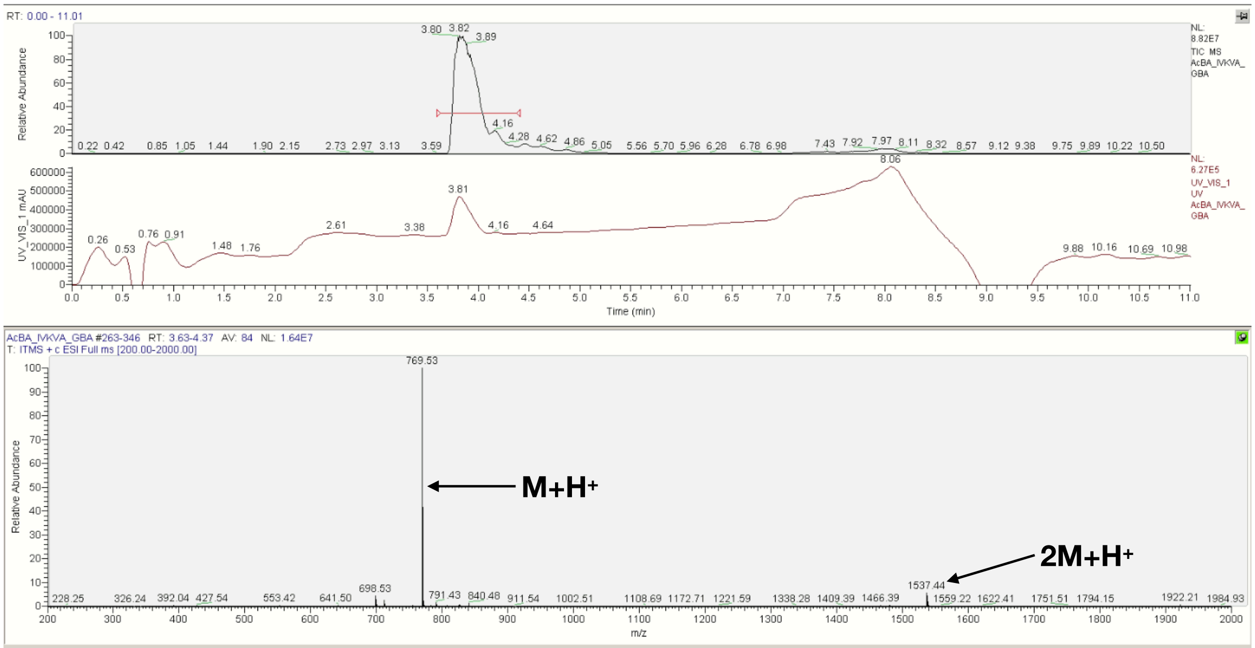


### F Ac-βA-IVKVA-Gly-Am

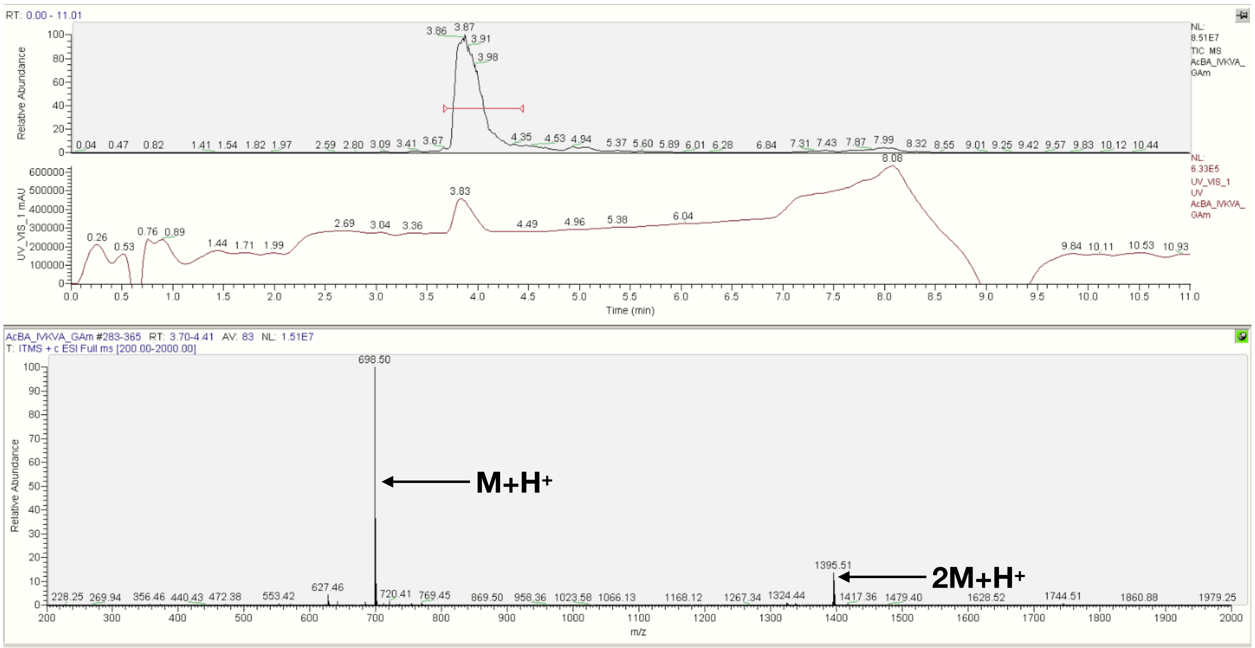

**G Ac-βA-IVKVA-Gly-COOH**

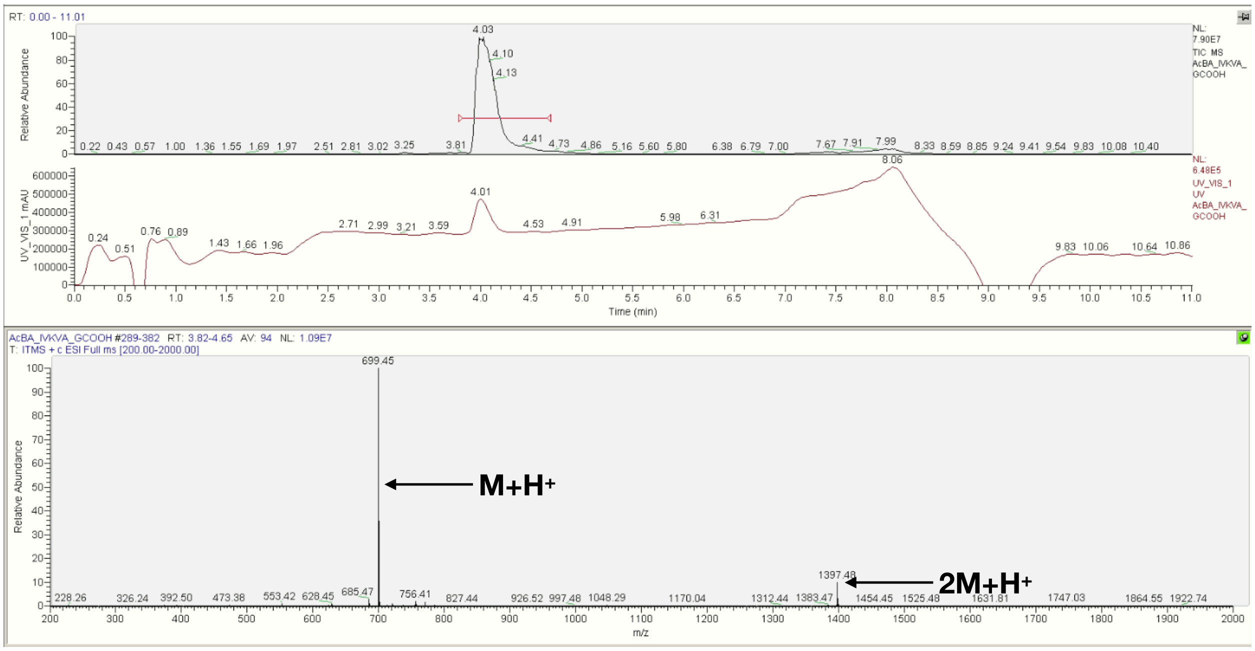


**Figure S21**. LCMS spectra of the IVKVA peptides in which a glycine was placed on the N-terminus for N-terminal libraries, and the C-terminus for C-terminal libraries. a) Ac-βA, b) Ac, c) N-βA, d) $NH_2$, e) C-βA, f) Am, g) COOH.

**A**

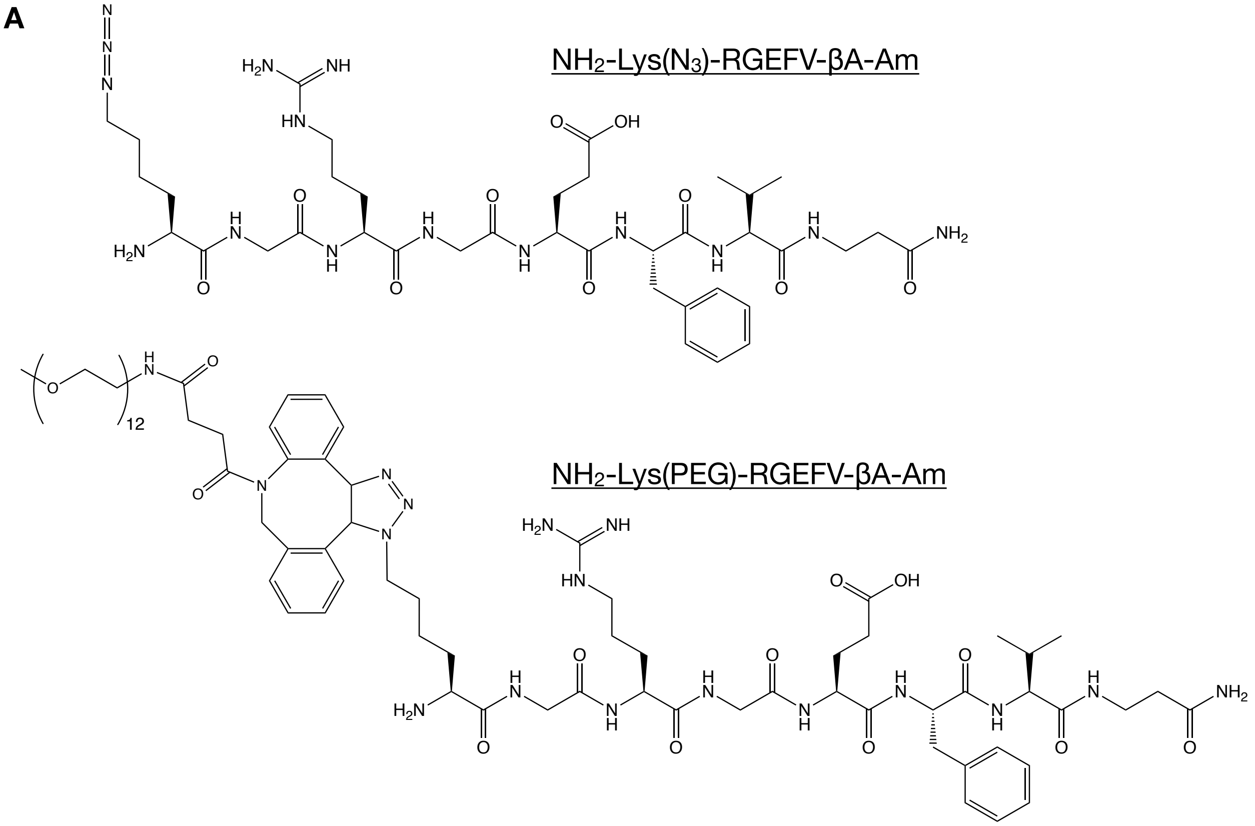

**B Ac-βA-Lys($N_3$)-RGEFV-βA-Am**

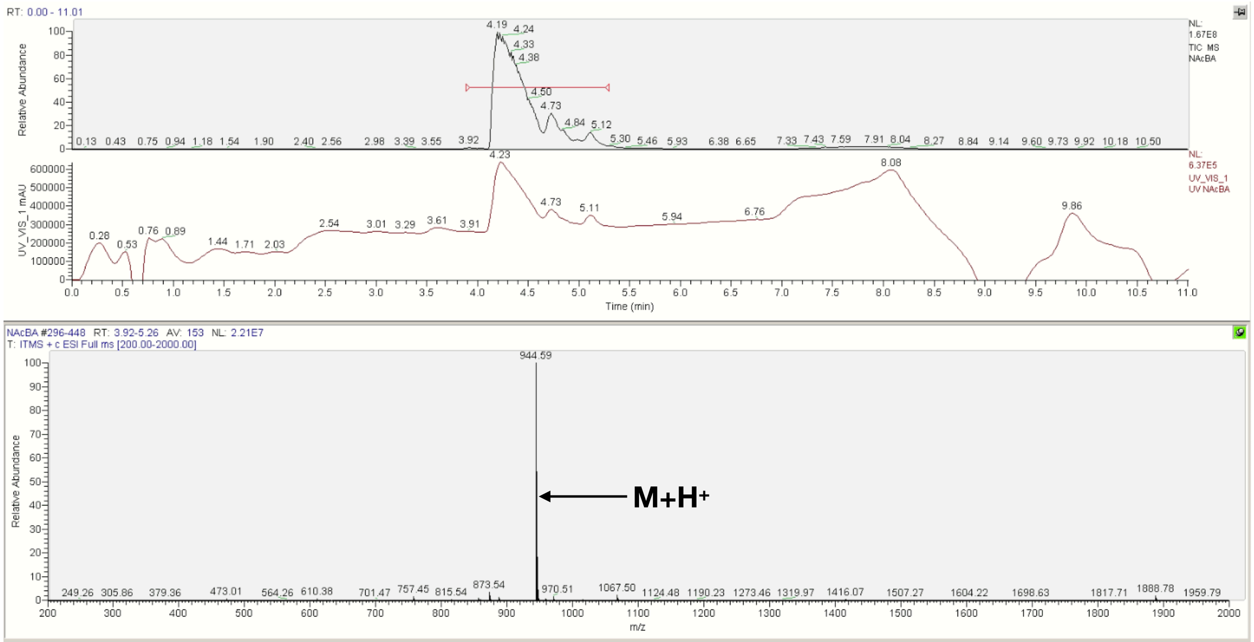


**C Ac-Lys($N_3$)-RGEFV-βA-Am**

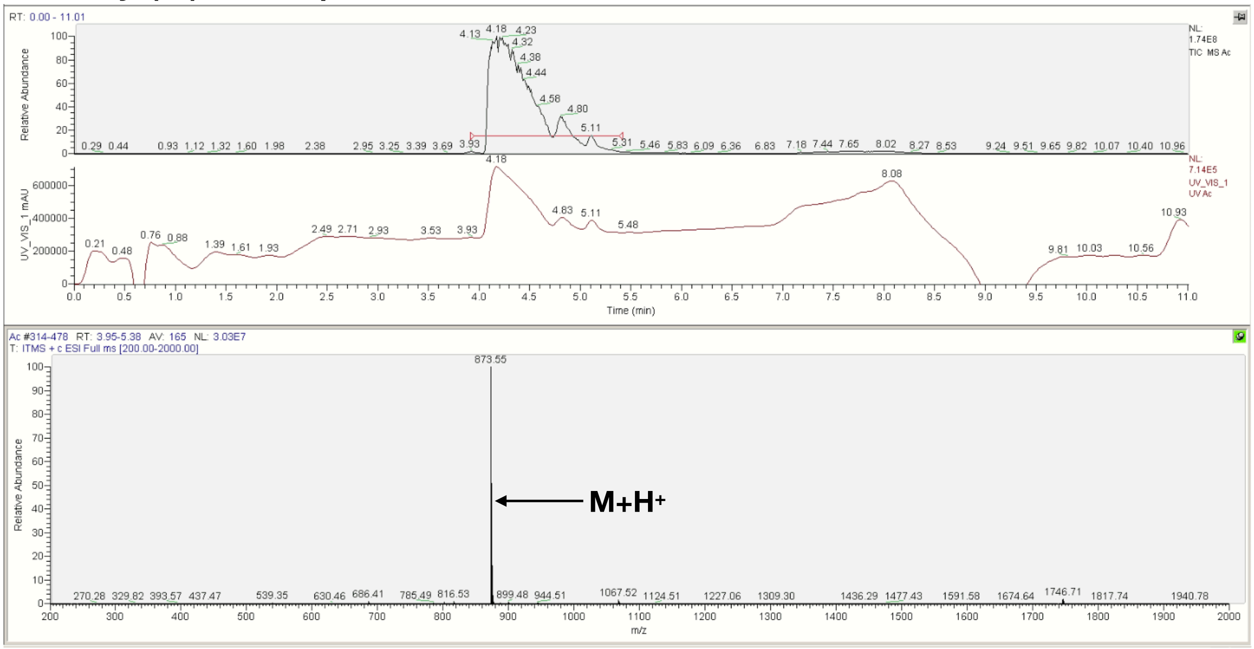

**D NH$_2$-βA-Lys(N$_3$)-RGEFV-βA-Am**

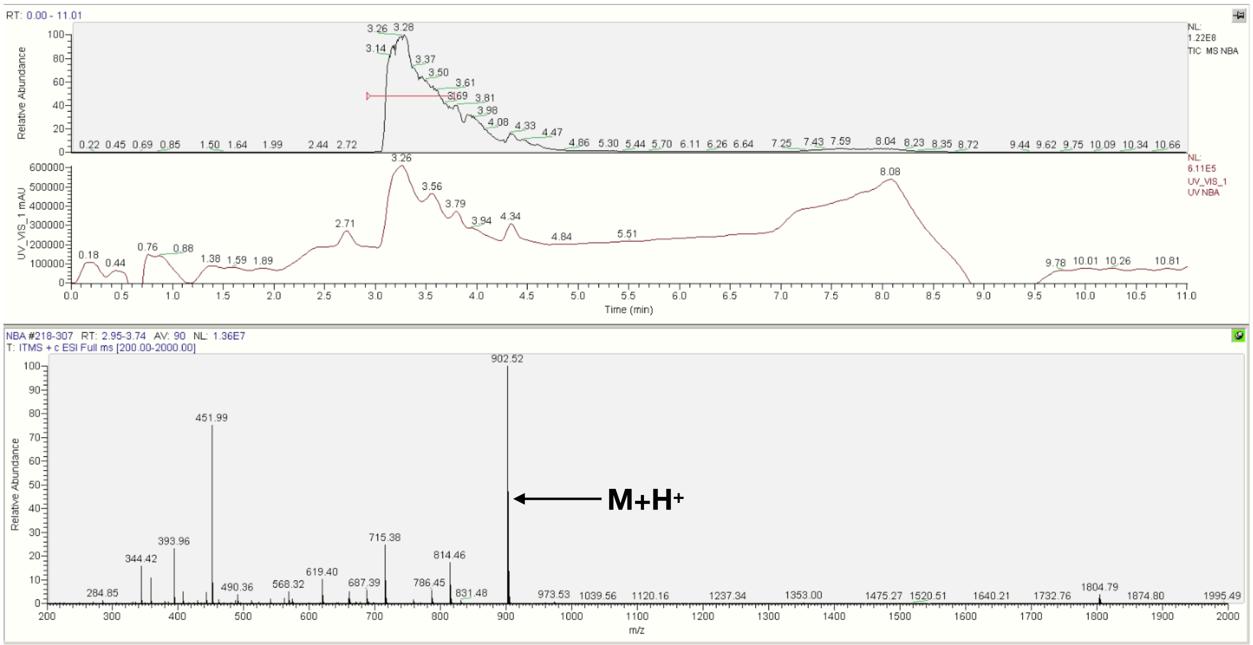


**E NH$_2$-Lys(N$_3$)-RGEFV-βA-Am**

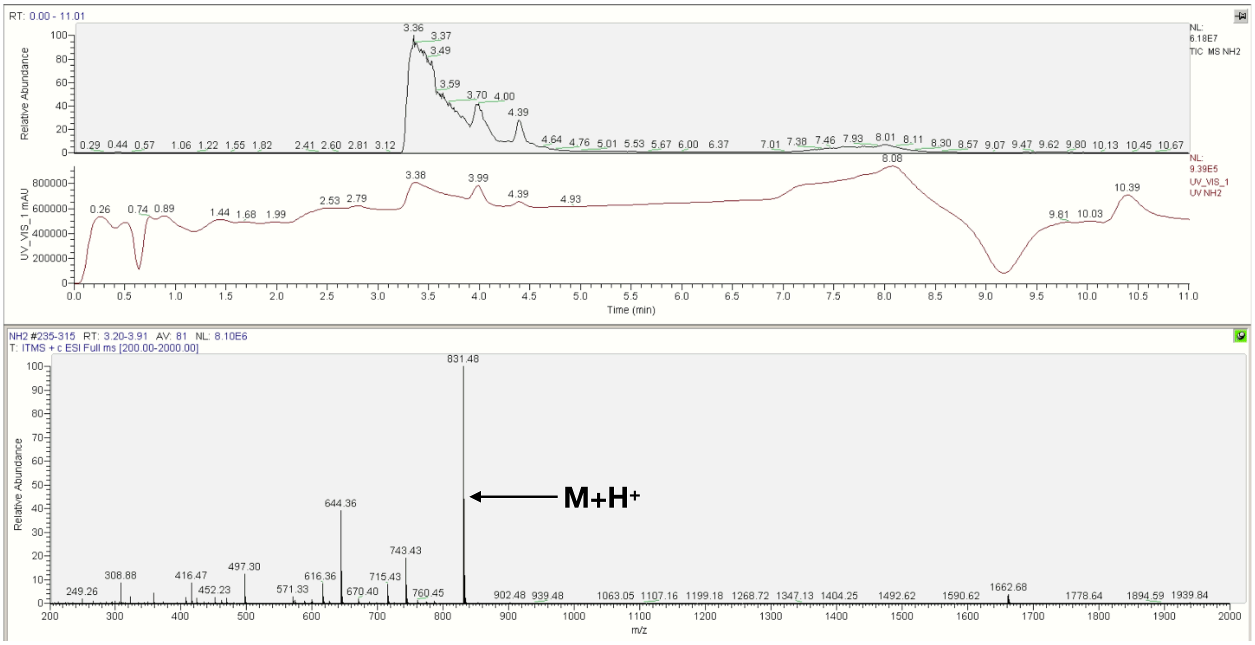

**F Ac-βA-RGEFV-Lys(N$_3$)-βA-Am**

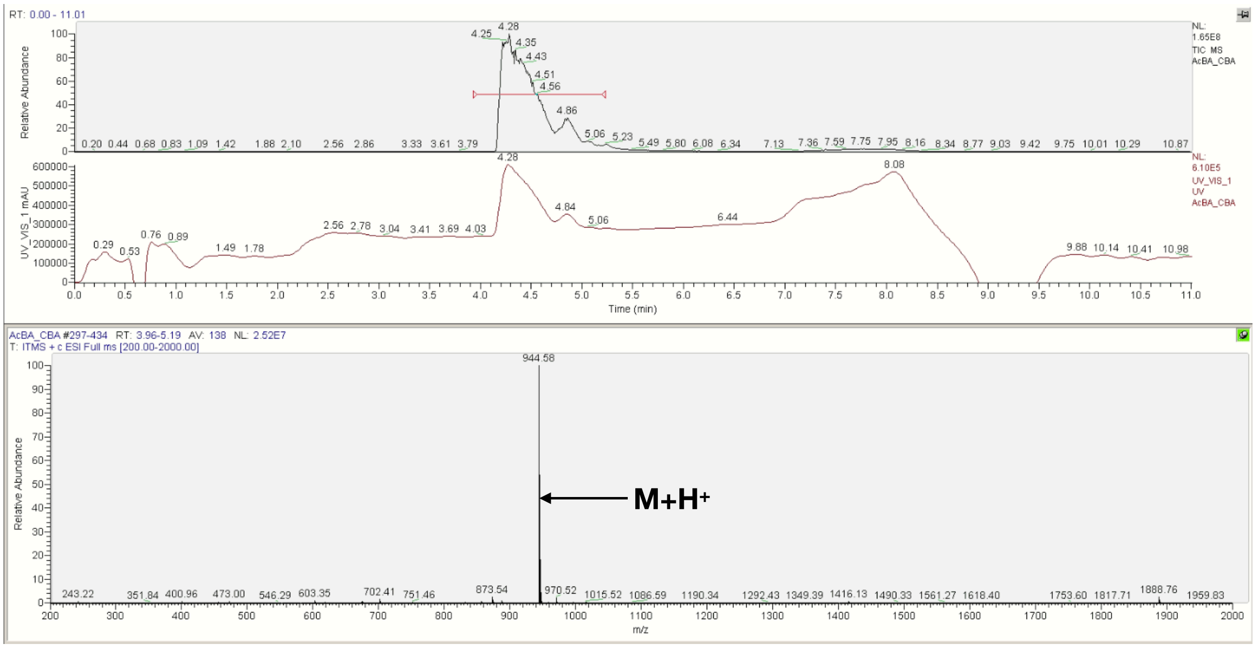


**G Ac-βA-RGEFV-Lys(N$_3$)-Am**

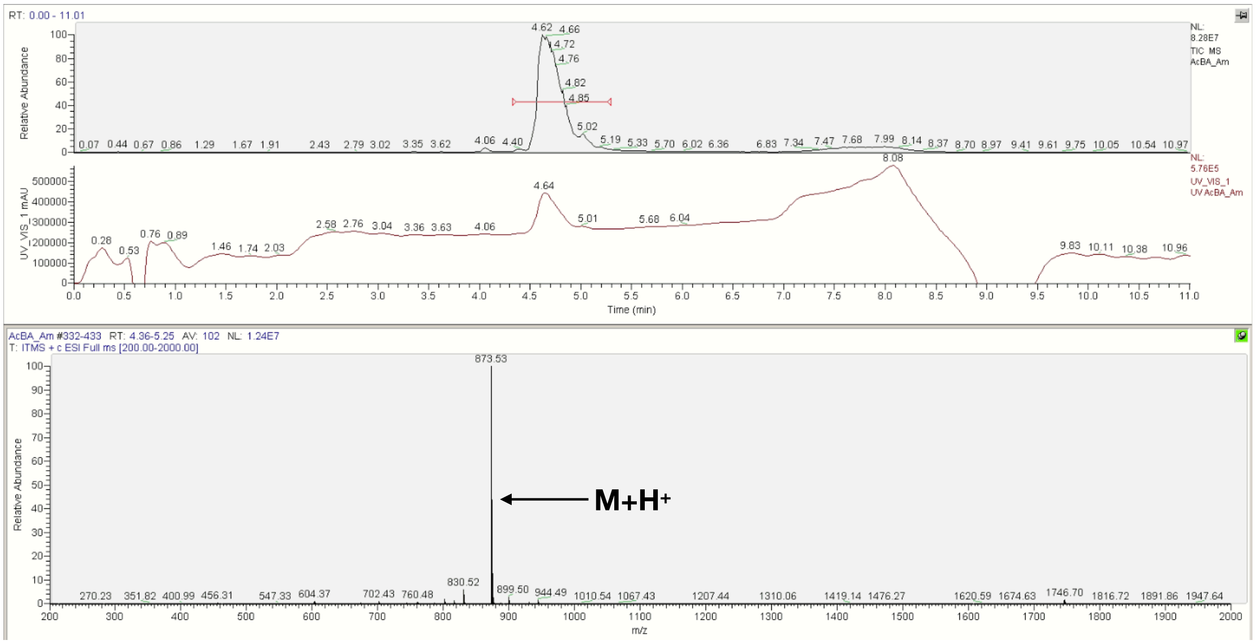

### H Ac-βA-RGEFV-Lys($N_3$)-COOH

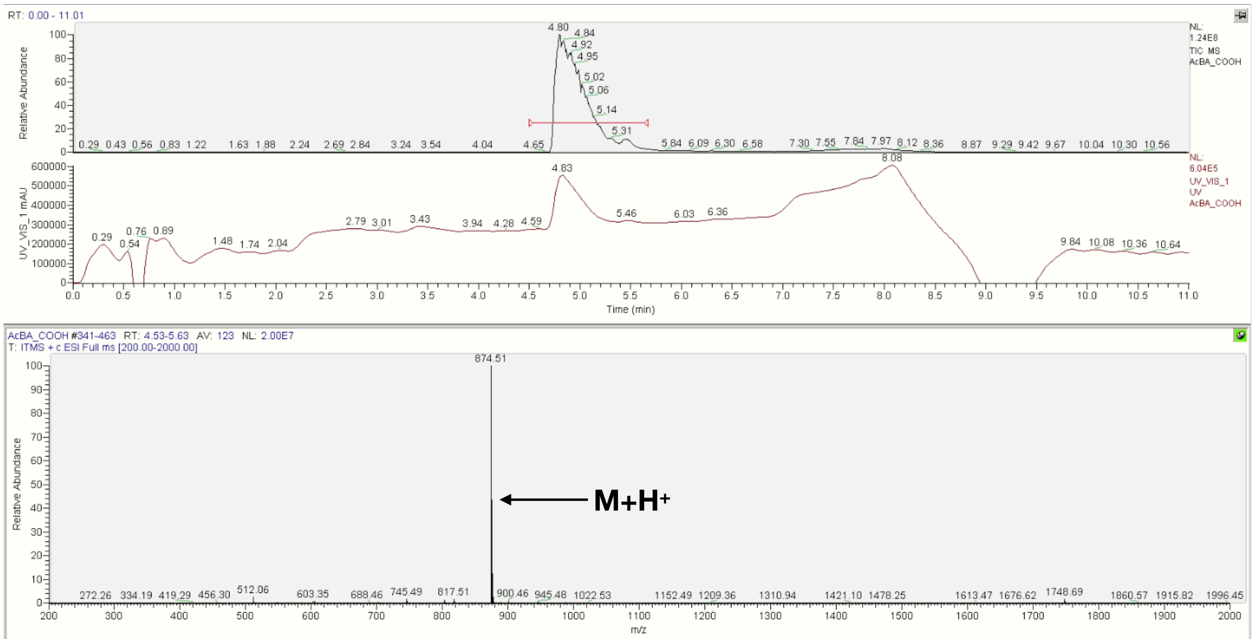


### I Ac-βA-Lys(PEG)-RGEFV-βA-Am

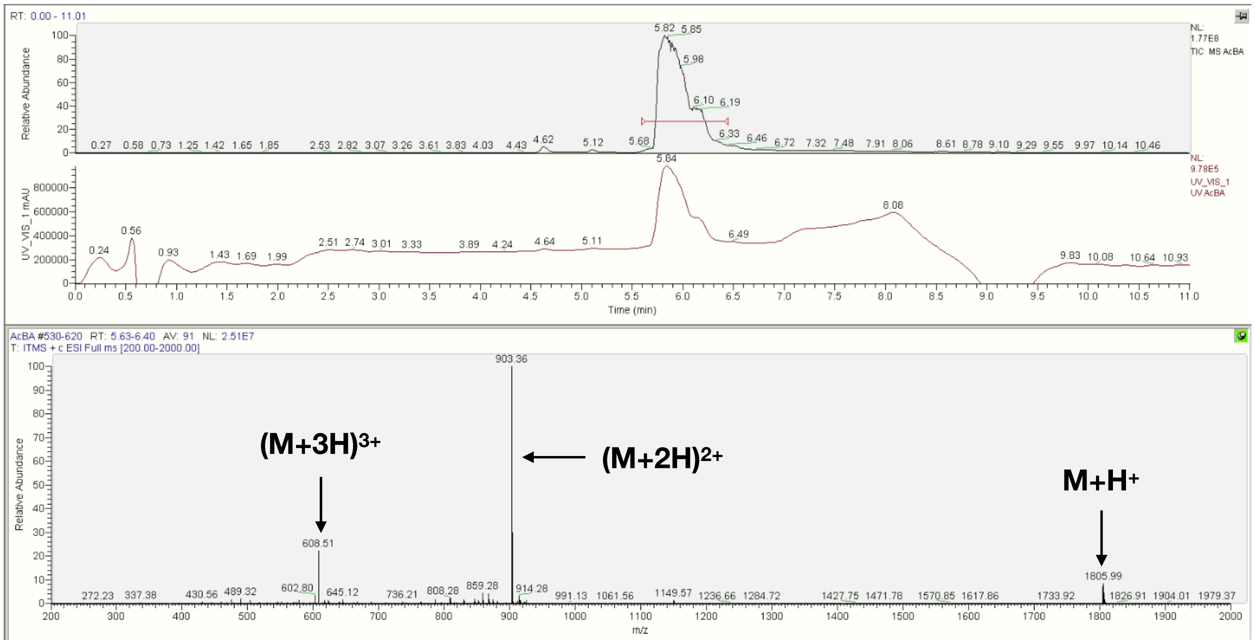

**J Ac-Lys(PEG)-RGEFV-βA-Am**

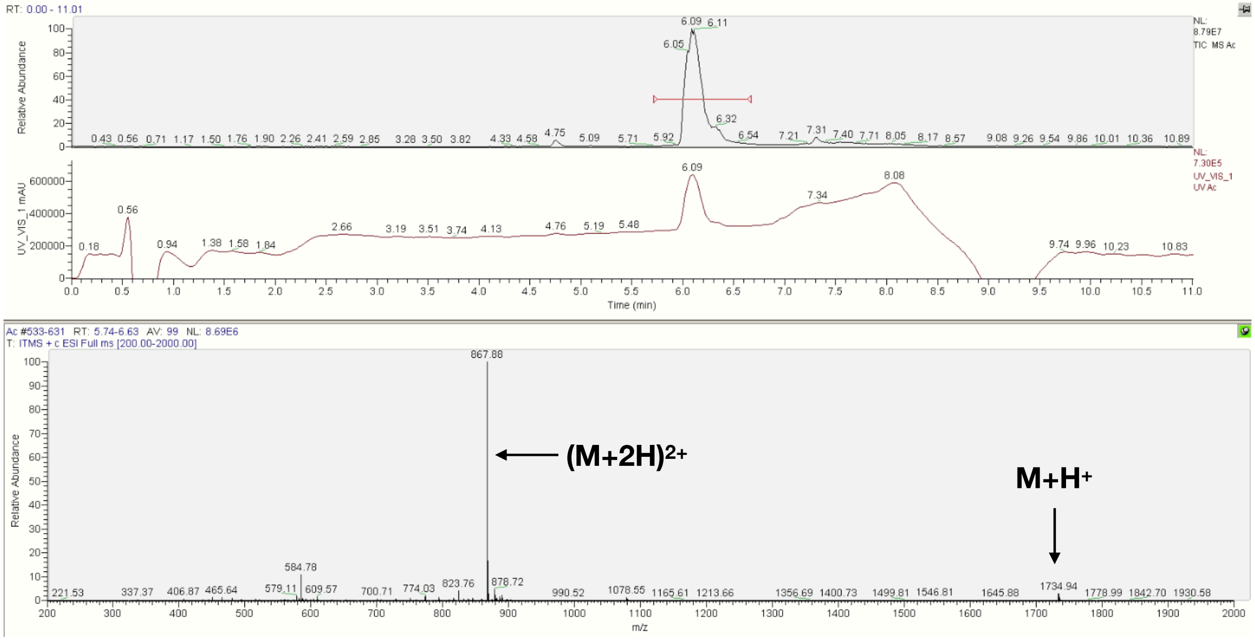


**K $NH_2$-βA-Lys(PEG)-RGEFV-βA-Am**

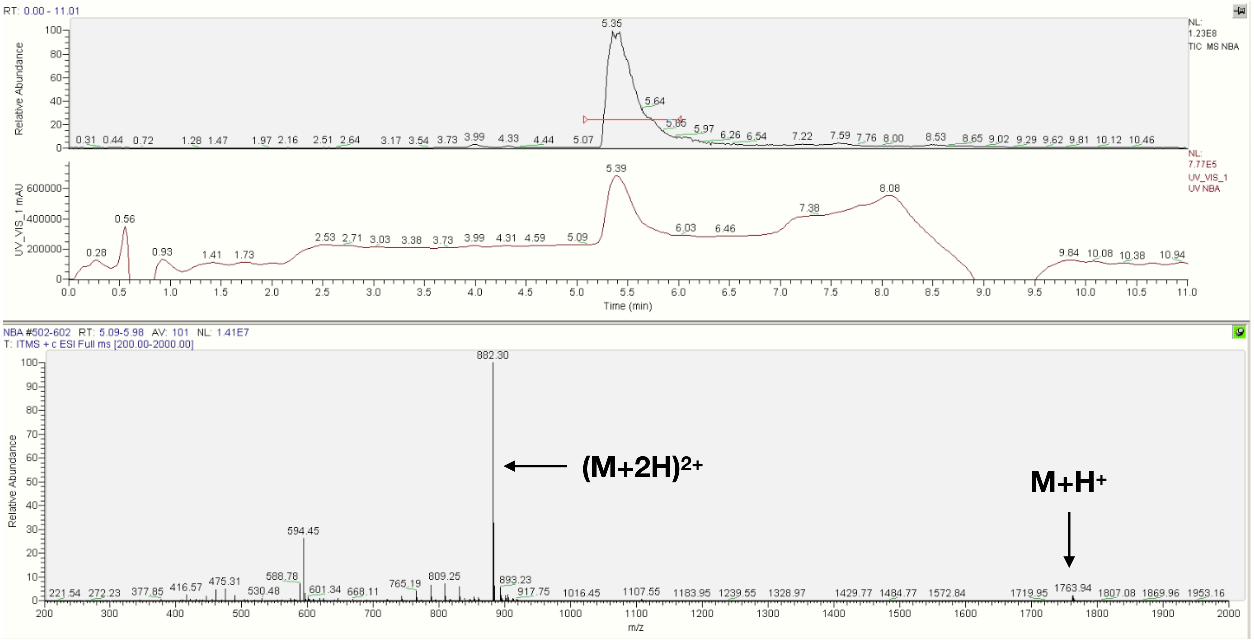

**L NH$_2$-Lys(PEG)-RGEFV-βA-Am**

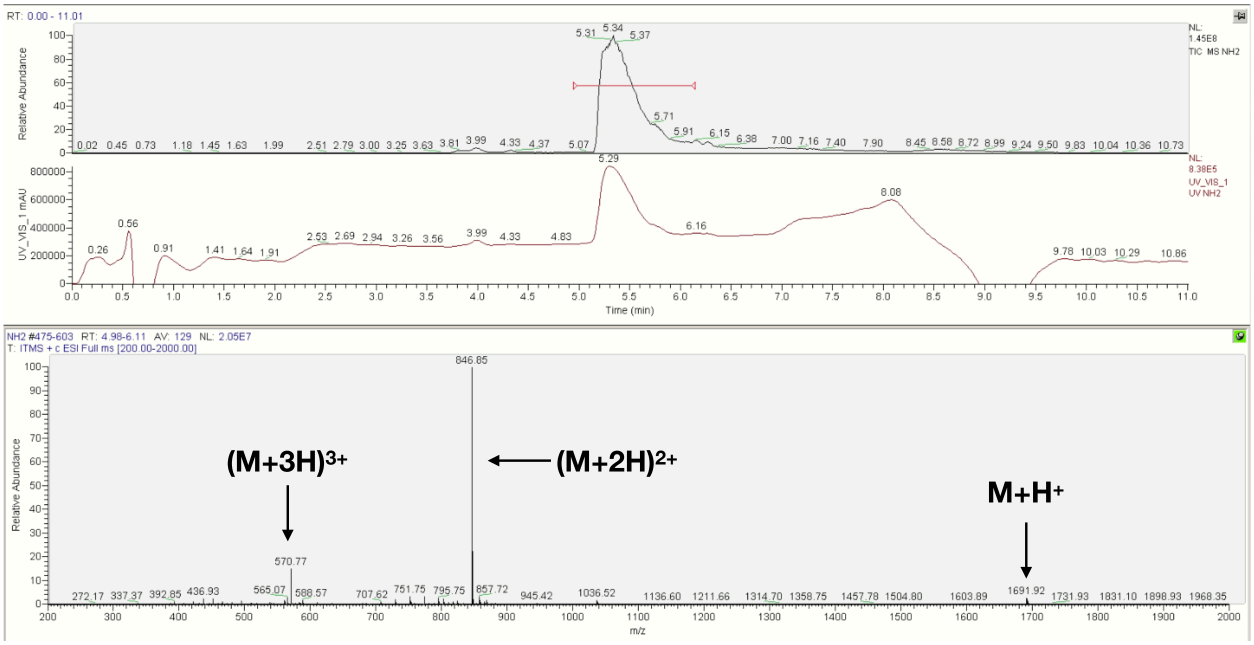


**M Ac-βA-RGEFV-Lys(PEG)-βA-Am**

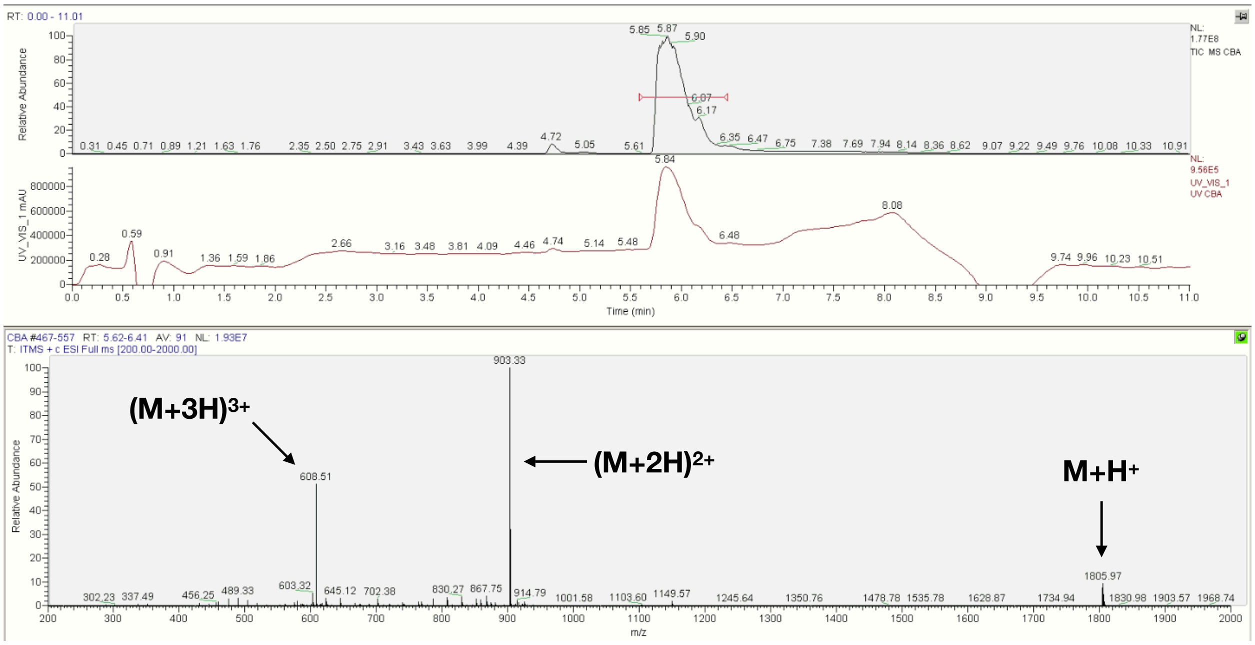

**N   Ac-βA-RGEFV-Lys(PEG)-Am**

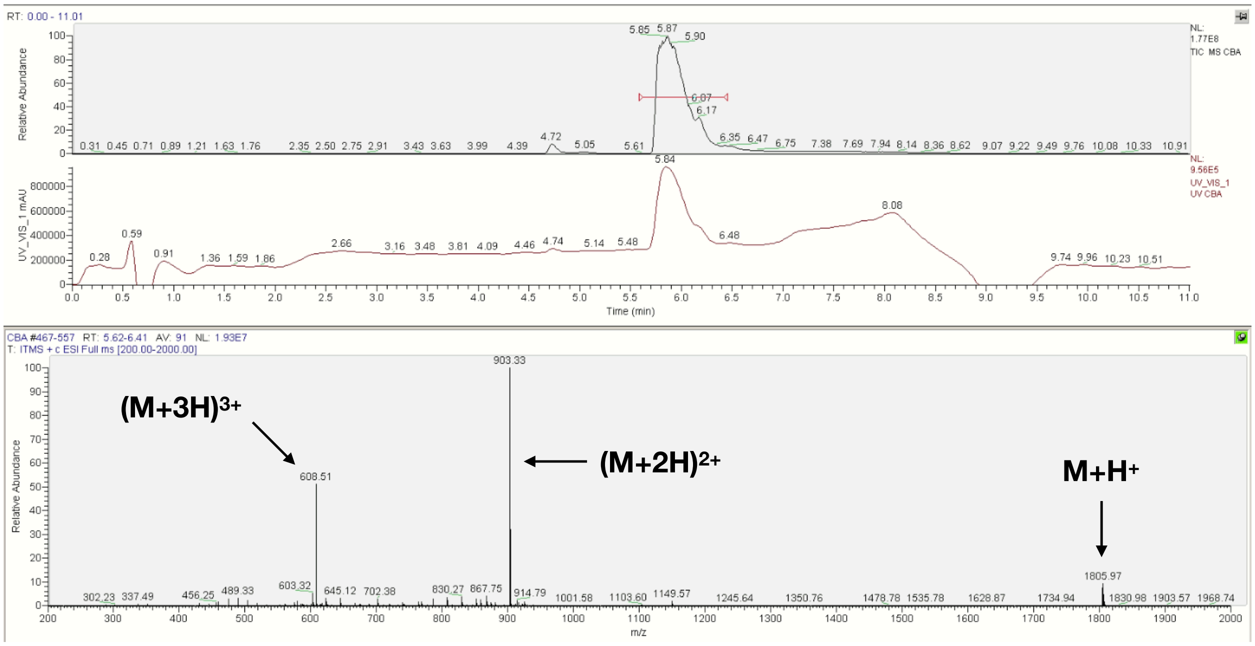


**O   Ac-βA-RGEFV-Lys(PEG)-COOH**

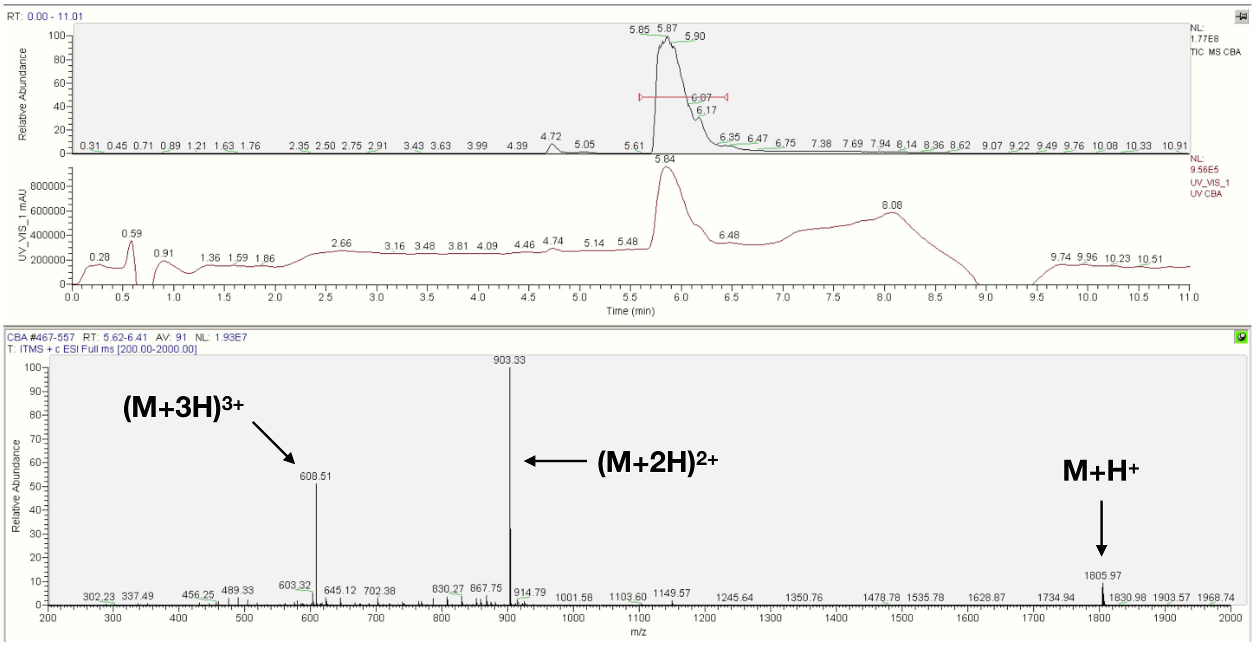


**Figure S22**. LCMS spectra of the Azide and PEG modified peptides in which a glycine was placed on the N-terminus for N-terminal libraries, and the C-terminus for C-terminal libraries. a) The structure of azide and PEG peptides. The azide peptides contained the following functionalizations: b) Ac-βA, c) Ac, d) N-βA, e) $NH_2$, f) C-βA, g) Am, h) COOH. The PEG-modified peptides contained the following functionalizations: i) Ac-βA, j) Ac, k) N-βA, l) $NH_2$, m) C-βA, n) Am, o) COOH.

**A** $N_3$-KGPQGIWGQLys($N_3$)-$NH_2$ (PanMMP)

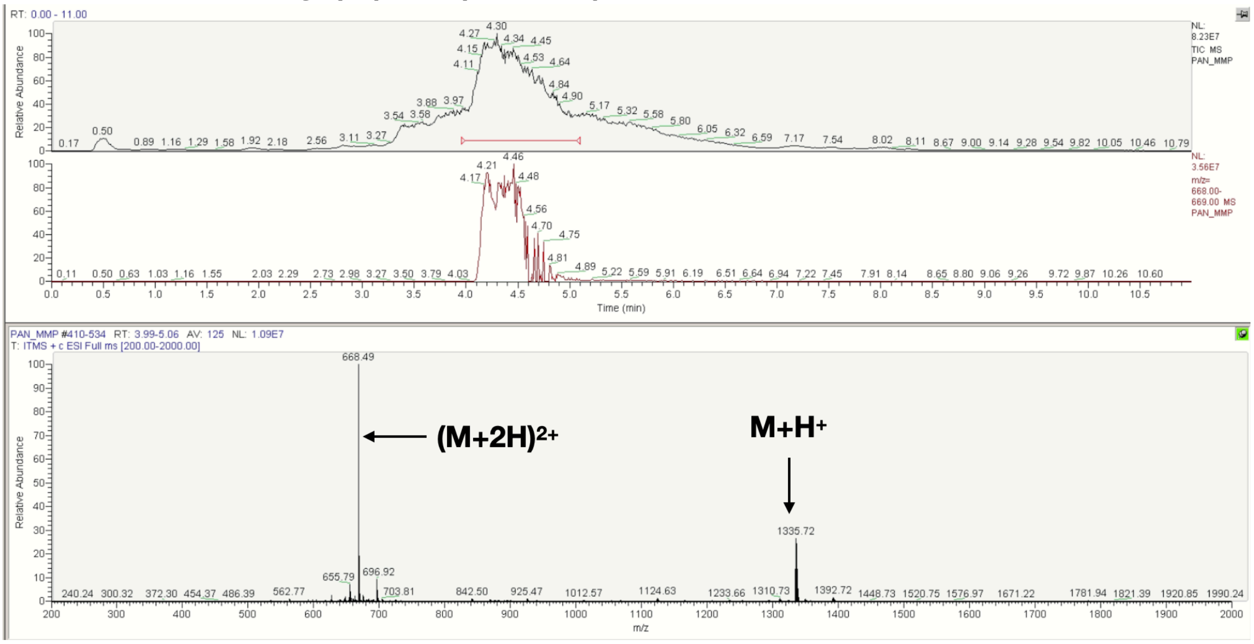


**B** $NH_2$-GRGDS-Lys($N_3$)-$NH_2$

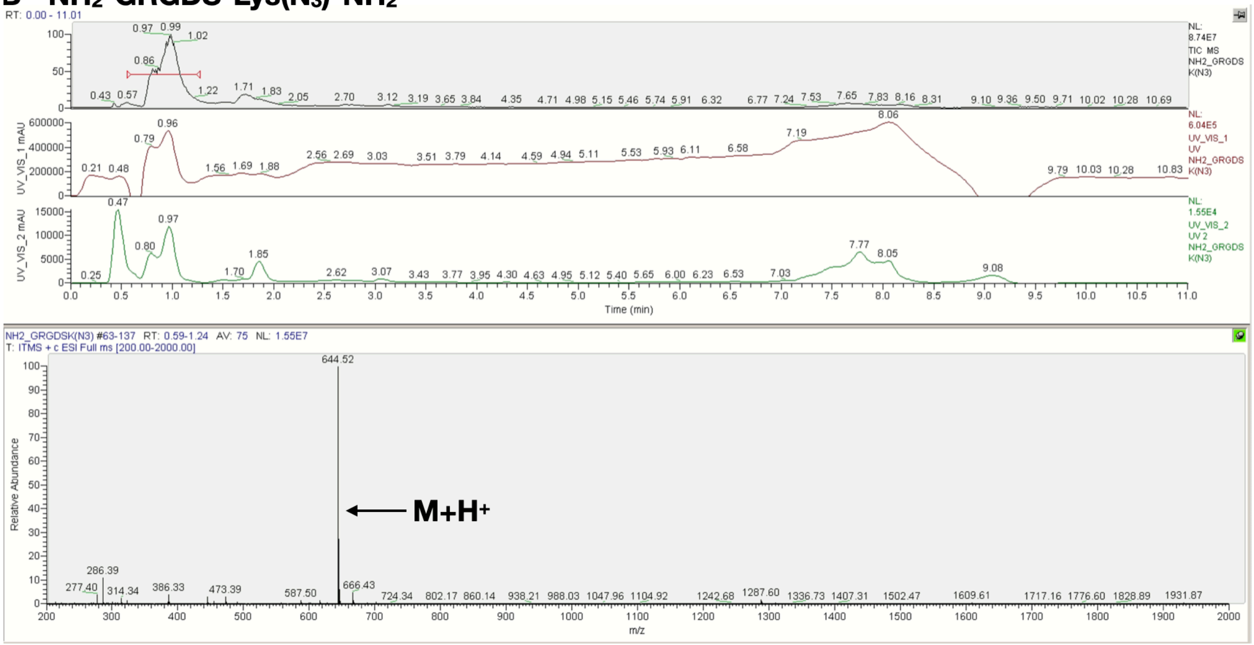

### C Ac-βA-GRGDS-Lys($N_3$)-$NH_2$

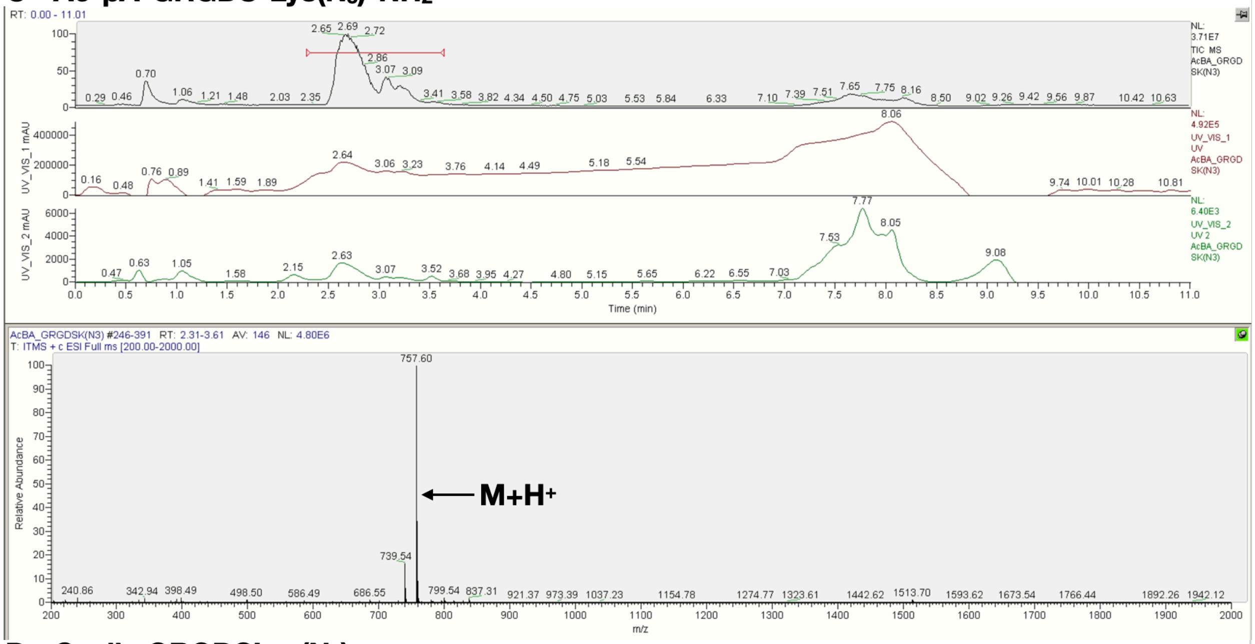


### D Cyclic GRGDSLys($N_3$)

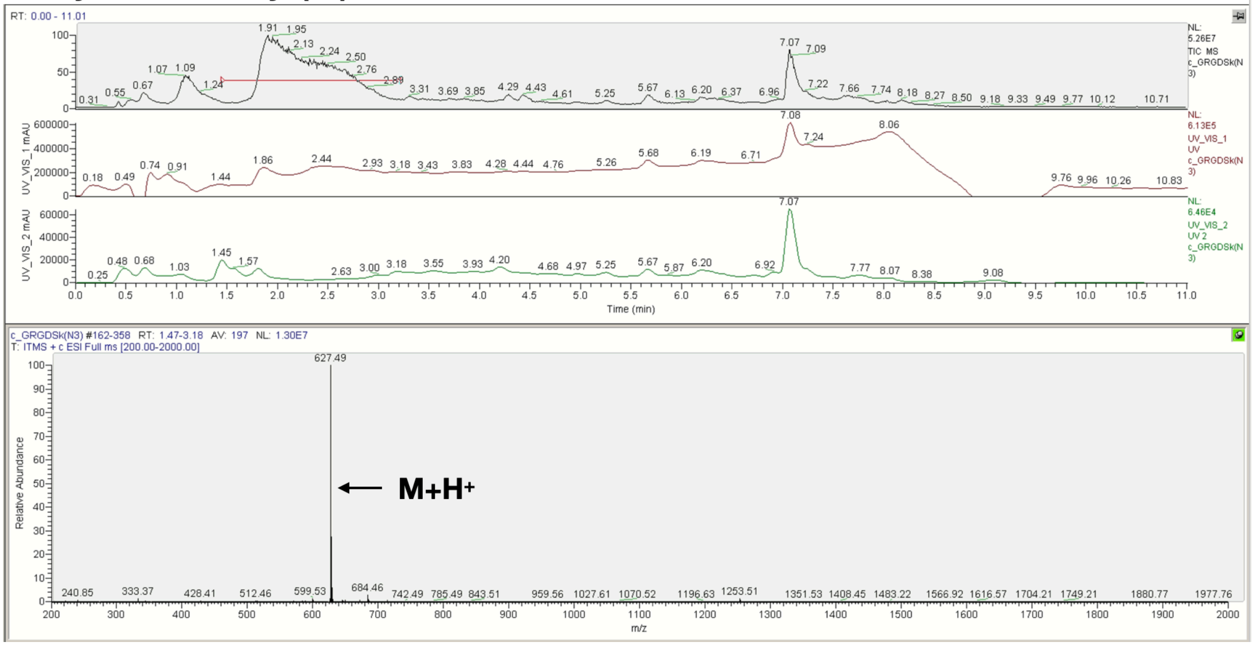

**E NH$_2$-βFβAβAβAβAβA-NH$_2$**

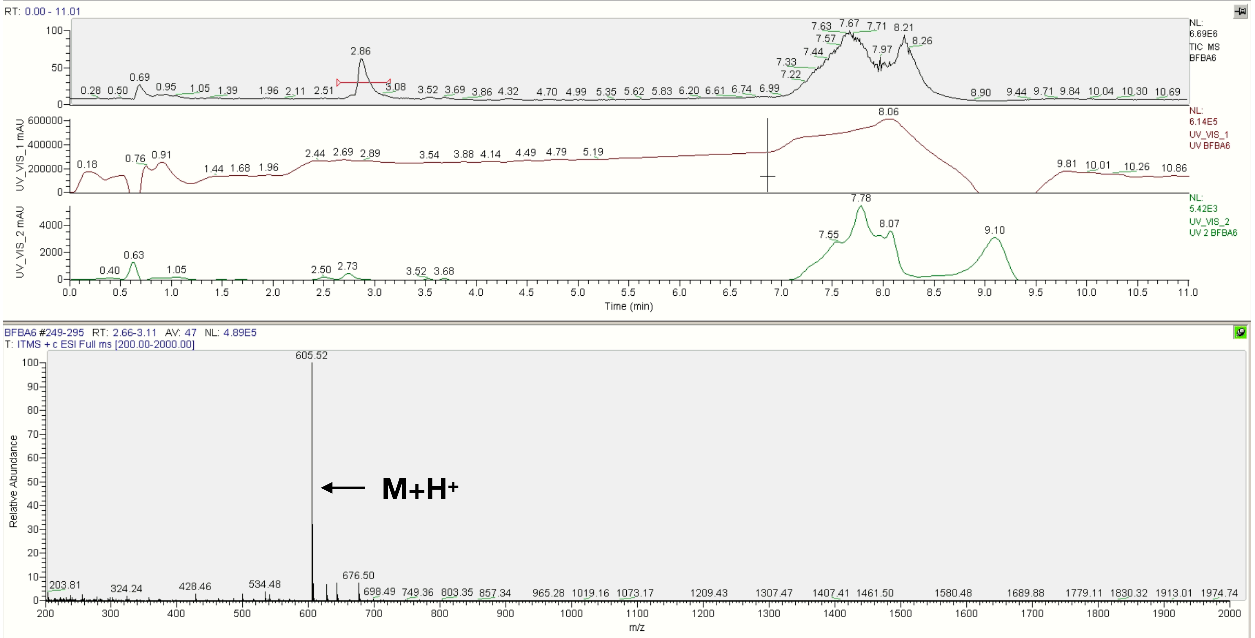


**Figure S23**. LCMS spectra of the peptides used for cell culture: a) N3-KGPQGIWGQK-Lys(N$_3$)-NH$_2$. Note that the N-terminus of the peptide contained an azide-acetic acid moiety. b) NH$_2$-GRGDS-Lys(N$_3$)-NH$_2$, c) Ac-βA-GRGDS-Lys(N$_3$)-NH$_2$, d) cyclic GRGDS-Lys(N$_3$), e) non-proteolytically degradable NH$_2$-βFβAβAβAβAβA-NH$_2$ internal standard used in all peptide degradation studies.

**Statistical Analysis**

**A** **Data for soluble peptides incubated with cells on tissue culture plastic.**

| | Df | Sum Sq | Mean Sq | F value | Pr(>F) |
|---|---|---|---|---|---|
| celltype | 2 | 279.3 | 139.65 | 3125.07 | <2e-16 *** |
| donor | 7 | 12.4 | 1.77 | 39.57 | <2e-16 *** |
| experiment | 7 | 22.7 | 3.24 | 72.56 | <2e-16 *** |
| endgroup | 6 | 376.8 | 62.79 | 1405.17 | <2e-16 *** |
| amino_acid | 17 | 36.4 | 2.14 | 47.91 | <2e-16 *** |
| timepoint | 5 | 607.1 | 121.43 | 2717.35 | <2e-16 *** |
| residuals | 22621 | 1010.9 | | | |

Fit: aov(formula = ave ~ cell + donor + experiment + endgroup + amino_acid + timepoint)

**Cell Type**

| | diff | lwr | upr | p adj |
|---|---|---|---|---|
| hUVEC-hMSC | 0.18174843 | 0.17324958 | 0.19024728 | 0 |
| Macrophage-hMSC | 0.26609418 | 0.25814551 | 0.27404286 | 0 |
| Macrophage-hUVEC | 0.08434575 | 0.07639741 | 0.09229409 | 0 |

**Endgroup - Averaged over all time points**

| | diff | lwr | upr | p adj |
|---|---|---|---|---|
| Am-CBA | -0.035178958 | -0.05066638 | -0.01969154 | 0.0000000 |
| COOH-CBA | -0.147212545 | -0.16269996 | -0.13172512 | 0.0000000 |
| AcBA-CBA | -0.004391847 | -0.01988046 | 0.01109677 | 0.9812038 |
| Ac-CBA | -0.036389911 | -0.05188451 | -0.02089531 | 0.0000000 |
| NBA-CBA | -0.093876888 | -0.10936431 | -0.07838947 | 0.0000000 |
| NH2-CBA | -0.389396425 | -0.40489103 | -0.37390182 | 0.0000000 |
| COOH-Am | -0.112033586 | -0.12751981 | -0.09654736 | 0.0000000 |
| AcBA-Am | 0.030787111 | 0.01529969 | 0.04627453 | 0.0000000 |
| Ac-Am | -0.001210953 | -0.01670436 | 0.01428245 | 0.9999876 |
| NBA-Am | -0.058697929 | -0.07418415 | -0.04321170 | 0.0000000 |
| NH2-Am | -0.354217467 | -0.36971087 | -0.33872406 | 0.0000000 |
| AcBA-COOH | 0.142820698 | 0.12733328 | 0.15830812 | 0.0000000 |
| Ac-COOH | 0.110822633 | 0.09532923 | 0.12631604 | 0.0000000 |
| NBA-COOH | 0.053335657 | 0.03784943 | 0.06882188 | 0.0000000 |
| NH2-COOH | -0.242183880 | -0.25767729 | -0.22669047 | 0.0000000 |
| Ac-AcBA | -0.031998064 | -0.04749267 | -0.01650346 | 0.0000000 |
| NBA-AcBA | -0.089485041 | -0.10497246 | -0.07399762 | 0.0000000 |
| NH2-AcBA | -0.385004578 | -0.40049918 | -0.36950998 | 0.0000000 |
| NBA-Ac | -0.057486976 | -0.07298038 | -0.04199357 | 0.0000000 |
| NH2-Ac | -0.353006513 | -0.36850710 | -0.33750593 | 0.0000000 |
| NH2-NBA | -0.295519537 | -0.31101294 | -0.28002613 | 0.0000000 |

**Amino Acid**

| | diff | lwr | upr | p adj |
|---|---|---|---|---|
| D-A | 0.0911418367 | 0.0617564292 | 0.1205272442 | 0.0000000 |
| E-A | 0.0674525839 | 0.0380671764 | 0.0968379914 | 0.0000000 |
| F-A | -0.0285623966 | -0.0579478041 | 0.0008230109 | 0.0680295 |
| G-A | 0.0074636879 | -0.0219217196 | 0.0368490954 | 0.9999903 |
| H-A | -0.0193680806 | -0.0487534881 | 0.0100173269 | 0.6837704 |
| I_L-A | 0.0150238223 | -0.0143615852 | 0.0444092299 | 0.9485997 |
| K-A | -0.0251598995 | -0.0545453070 | 0.0042255080 | 0.2061101 |
| M-A | -0.0071542074 | -0.0365454494 | 0.0222370346 | 0.9999948 |
| N-A | 0.0307876787 | 0.0014022712 | 0.0601730862 | 0.0286257 |
| P-A | 0.0493901460 | 0.0200047385 | 0.0787755535 | 0.0000005 |
| Q-A | -0.0006024072 | -0.0299878147 | 0.0287830003 | 1.0000000 |
| R-A | -0.0697643361 | -0.0991497436 | -0.0403789286 | 0.0000000 |
| S-A | -0.0104474036 | -0.0398328111 | 0.0189380039 | 0.9990181 |
| T-A | 0.0102079392 | -0.0191774683 | 0.0395933467 | 0.9992686 |
| V-A | 0.0375140807 | 0.0080876607 | 0.0669405006 | 0.0012033 |
| W-A | -0.0644909884 | -0.0939115249 | -0.0350704518 | 0.0000000 |
| Y-A | -0.0120895025 | -0.0414749100 | 0.0172959050 | 0.9943445 |
| E-D | -0.0236892528 | -0.0530746603 | 0.0056961547 | 0.3045992 |
| F-D | -0.1197042334 | -0.1490896409 | -0.0903188259 | 0.0000000 |
| G-D | -0.0836781489 | -0.1130635564 | -0.0542927414 | 0.0000000 |
| H-D | -0.1105099174 | -0.1398953249 | -0.0811245099 | 0.0000000 |
| I_L-D | -0.0761180144 | -0.1055034219 | -0.0467326069 | 0.0000000 |
| K-D | -0.1163017363 | -0.1456871438 | -0.0869163288 | 0.0000000 |
| M-D | -0.0982960442 | -0.1276872861 | -0.0689048022 | 0.0000000 |
| N-D | -0.0603541581 | -0.0897395656 | -0.0309687506 | 0.0000000 |
| P-D | -0.0417516908 | -0.0711370983 | -0.0123662833 | 0.0001051 |
| Q-D | -0.0917442439 | -0.1211296514 | -0.0623588364 | 0.0000000 |
| R-D | -0.1609061728 | -0.1902915803 | -0.1315207653 | 0.0000000 |
| S-D | -0.1015892404 | -0.1309746479 | -0.0722038329 | 0.0000000 |
| T-D | -0.0809338976 | -0.1103193051 | -0.0515484901 | 0.0000000 |
| V-D | -0.0536277561 | -0.0830541760 | -0.0242013361 | 0.0000000 |
| W-D | -0.1556328251 | -0.1850533616 | -0.1262122886 | 0.0000000 |
| Y-D | -0.1032313392 | -0.1326167467 | -0.0738459317 | 0.0000000 |
| F-E | -0.0960149805 | -0.1254003880 | -0.0666295730 | 0.0000000 |
| G-E | -0.0599888960 | -0.0893743035 | -0.0306034885 | 0.0000000 |
| H-E | -0.0868206645 | -0.1162060720 | -0.0574352570 | 0.0000000 |
| I_L-E | -0.0524287616 | -0.0818141691 | -0.0230433541 | 0.0000000 |
| K-E | -0.0926124834 | -0.1219978909 | -0.0632270759 | 0.0000000 |
| M-E | -0.0746067913 | -0.1039980333 | -0.0452155493 | 0.0000000 |
| N-E | -0.0366649053 | -0.0660503128 | -0.0072794978 | 0.0018310 |
| P-E | -0.0180624379 | -0.0474478454 | 0.0113229696 | 0.7898131 |
| Q-E | -0.0680549911 | -0.0974403986 | -0.0386695836 | 0.0000000 |
| R-E | -0.1372169200 | -0.1666023275 | -0.1078315125 | 0.0000000 |
| S-E | -0.0778999875 | -0.1072853950 | -0.0485145800 | 0.0000000 |
| T-E | -0.0572446447 | -0.0866300522 | -0.0278592372 | 0.0000000 |
| V-E | -0.0299385032 | -0.0593649232 | -0.0005120833 | 0.0409923 |
| W-E | -0.1319435723 | -0.1613641088 | -0.1025230357 | 0.0000000 |
| Y-E | -0.0795420864 | -0.1089274939 | -0.0501566789 | 0.0000000 |
| G-F | 0.0360260845 | 0.0066406770 | 0.0654114920 | 0.0025440 |
| H-F | 0.0091943160 | -0.0201910915 | 0.0385797235 | 0.9998155 |
| I_L-F | 0.0435862190 | 0.0142008115 | 0.0729716265 | 0.0000338 |
| K-F | 0.0034024971 | -0.0259829104 | 0.0327879046 | 1.0000000 |

| | | | | |
|---|---|---|---|---|
| M-F | 0.0214081892 | -0.0079830528 | 0.0507994312 | 0.4981737 |
| N-F | 0.0593500753 | 0.0299646678 | 0.0887354828 | 0.0000000 |
| P-F | 0.0779525426 | 0.0485671351 | 0.1073379501 | 0.0000000 |
| Q-F | 0.0279599894 | -0.0014254181 | 0.0573453970 | 0.0844257 |
| R-F | -0.0412019394 | -0.0705873469 | -0.0118165319 | 0.0001460 |
| S-F | 0.0181149930 | -0.0112704145 | 0.0475004005 | 0.7859067 |
| T-F | 0.0387703358 | 0.0093849283 | 0.0681557433 | 0.0005903 |
| V-F | 0.0660764773 | 0.0366500574 | 0.0955028973 | 0.0000000 |
| W-F | -0.0359285917 | -0.0653491282 | -0.0065080552 | 0.0027321 |
| Y-F | 0.0164728942 | -0.0129125133 | 0.0458583017 | 0.8898527 |
| H-G | -0.0268317685 | -0.0562171760 | 0.0025536390 | 0.1237204 |
| I_L-G | 0.0075601345 | -0.0218252730 | 0.0369455420 | 0.9999883 |
| K-G | -0.0326235874 | -0.0620089949 | -0.0032381799 | 0.0129911 |
| M-G | -0.0146178953 | -0.0440091373 | 0.0147733467 | 0.9599599 |
| N-G | 0.0233239908 | -0.0060614167 | 0.0527093983 | 0.3326789 |
| P-G | 0.0419264581 | 0.0125410506 | 0.0713118656 | 0.0000946 |
| Q-G | -0.0080660950 | -0.0374515025 | 0.0213193125 | 0.9999702 |
| R-G | -0.0772280239 | -0.1066134314 | -0.0478426164 | 0.0000000 |
| S-G | -0.0179110915 | -0.0472964990 | 0.0114743160 | 0.8008656 |
| T-G | 0.0027442513 | -0.0266411562 | 0.0321296588 | 1.0000000 |
| V-G | 0.0300503928 | 0.0006239729 | 0.0594768128 | 0.0392224 |
| W-G | -0.0719546762 | -0.1013752127 | -0.0425341397 | 0.0000000 |
| Y-G | -0.0195531903 | -0.0489385978 | 0.0098322172 | 0.6674980 |
| I_L-H | 0.0343919030 | 0.0050064955 | 0.0637773105 | 0.0057134 |
| K-H | -0.0057918189 | -0.0351772264 | 0.0235935886 | 0.9999998 |
| M-H | 0.0122138732 | -0.0171773688 | 0.0416051152 | 0.9936667 |
| N-H | 0.0501557593 | 0.0207703518 | 0.0795411668 | 0.0000002 |
| P-H | 0.0687582266 | 0.0393728191 | 0.0981436341 | 0.0000000 |
| Q-H | 0.0187656735 | -0.0106197340 | 0.0481510810 | 0.7348440 |
| R-H | -0.0503962554 | -0.0797816629 | -0.0210108479 | 0.0000001 |
| S-H | 0.0089206770 | -0.0204647305 | 0.0383060845 | 0.9998777 |
| T-H | 0.0295760198 | 0.0001906123 | 0.0589614273 | 0.0464617 |
| V-H | 0.0568821613 | 0.0274557414 | 0.0863085813 | 0.0000000 |
| W-H | -0.0451229077 | -0.0745434442 | -0.0157023712 | 0.0000130 |
| Y-H | 0.0072785782 | -0.0221068293 | 0.0366639857 | 0.9999933 |
| K-I_L | -0.0401837219 | -0.0695691294 | -0.0107983144 | 0.0002651 |
| M-I_L | -0.0221780298 | -0.0515692717 | 0.0072132122 | 0.4289220 |
| N-I_L | 0.0157638563 | -0.0136215512 | 0.0451492638 | 0.9223924 |
| P-I_L | 0.0343663236 | 0.0049809161 | 0.0637517311 | 0.0057841 |
| Q-I_L | -0.0156262295 | -0.0450116370 | 0.0137591780 | 0.9278480 |
| R-I_L | -0.0847881584 | -0.1141735659 | -0.0554027509 | 0.0000000 |
| S-I_L | -0.0254712260 | -0.0548566335 | 0.0039141815 | 0.1884060 |
| T-I_L | -0.0048158832 | -0.0342012907 | 0.0245695243 | 1.0000000 |
| V-I_L | 0.0224902583 | -0.0069361616 | 0.0519166783 | 0.4040892 |
| W-I_L | -0.0795148107 | -0.1089353472 | -0.0500942742 | 0.0000000 |
| Y-I_L | -0.0271133248 | -0.0564987323 | 0.0022720827 | 0.1127786 |
| M-K | 0.0180056921 | -0.0113855499 | 0.0473969341 | 0.7942535 |
| N-K | 0.0559475782 | 0.0265621707 | 0.0853329857 | 0.0000000 |
| P-K | 0.0745500455 | 0.0451646380 | 0.1039354530 | 0.0000000 |
| Q-K | 0.0245574924 | -0.0048279151 | 0.0539428999 | 0.2435191 |
| R-K | -0.0446044365 | -0.0739898440 | -0.0152190290 | 0.0000176 |
| S-K | 0.0147124959 | -0.0146729116 | 0.0440979034 | 0.9574387 |
| T-K | 0.0353678387 | 0.0059824312 | 0.0647532462 | 0.0035440 |
| V-K | 0.0626739802 | 0.0332475603 | 0.0921004002 | 0.0000000 |

| | | | | |
|---|---|---|---|---|
| W-K | -0.0393310888 | -0.0687516253 | -0.0099105523 | 0.0004429 |
| Y-K | 0.0130703971 | -0.0163150104 | 0.0424558046 | 0.9867309 |
| N-M | 0.0379418861 | 0.0085506441 | 0.0673331280 | 0.0009334 |
| P-M | 0.0565443534 | 0.0271531114 | 0.0859355954 | 0.0000000 |
| Q-M | 0.0065518002 | -0.0228394418 | 0.0359430422 | 0.9999986 |
| R-M | -0.0626101287 | -0.0920013707 | -0.0332188867 | 0.0000000 |
| S-M | -0.0032931962 | -0.0326844382 | 0.0260980458 | 1.0000000 |
| T-M | 0.0173621466 | -0.0120290954 | 0.0467533886 | 0.8385896 |
| V-M | 0.0446682881 | 0.0152360418 | 0.0741005344 | 0.0000177 |
| W-M | -0.0573367809 | -0.0867631450 | -0.0279104169 | 0.0000000 |
| Y-M | -0.0049352951 | -0.0343265371 | 0.0244559469 | 1.0000000 |
| P-N | 0.0186024673 | -0.0107829402 | 0.0479878748 | 0.7480944 |
| Q-N | -0.0313900858 | -0.0607754933 | -0.0020046783 | 0.0222507 |
| R-N | -0.1005520147 | -0.1299374222 | -0.0711666072 | 0.0000000 |
| S-N | -0.0412350823 | -0.0706204898 | -0.0118496748 | 0.0001432 |
| T-N | -0.0205797395 | -0.0499651470 | 0.0088056680 | 0.5741915 |
| V-N | 0.0067264020 | -0.0227000179 | 0.0361528220 | 0.9999980 |
| W-N | -0.0952786670 | -0.1246992035 | -0.0658581305 | 0.0000000 |
| Y-N | -0.0428771811 | -0.0722625886 | -0.0134917736 | 0.0000528 |
| Q-P | -0.0499925532 | -0.0793779607 | -0.0206071457 | 0.0000002 |
| R-P | -0.1191544821 | -0.1485398896 | -0.0897690746 | 0.0000000 |
| S-P | -0.0598375496 | -0.0892229571 | -0.0304521421 | 0.0000000 |
| T-P | -0.0391822068 | -0.0685676143 | -0.0097967993 | 0.0004690 |
| V-P | -0.0118760653 | -0.0413024852 | 0.0175503547 | 0.9954584 |
| W-P | -0.1138811343 | -0.1433016708 | -0.0844605978 | 0.0000000 |
| Y-P | -0.0614796485 | -0.0908650560 | -0.0320942410 | 0.0000000 |
| R-Q | -0.0691619289 | -0.0985473364 | -0.0397765214 | 0.0000000 |
| S-Q | -0.0098449964 | -0.0392304039 | 0.0195404111 | 0.9995420 |
| T-Q | 0.0108103464 | -0.0185750611 | 0.0401957539 | 0.9984965 |
| V-Q | 0.0381164879 | 0.0086900679 | 0.0675429078 | 0.0008703 |
| W-Q | -0.0638885812 | -0.0933091177 | -0.0344680447 | 0.0000000 |
| Y-Q | -0.0114870953 | -0.0408725028 | 0.0178983122 | 0.9968695 |
| S-R | 0.0593169325 | 0.0299315250 | 0.0887023400 | 0.0000000 |
| T-R | 0.0799722752 | 0.0505868677 | 0.1093576827 | 0.0000000 |
| V-R | 0.1072784168 | 0.0778519968 | 0.1367048367 | 0.0000000 |
| W-R | 0.0052733477 | -0.0241471888 | 0.0346938842 | 1.0000000 |
| Y-R | 0.0576748336 | 0.0282894261 | 0.0870602411 | 0.0000000 |
| T-S | 0.0206553428 | -0.0087300647 | 0.0500407503 | 0.5672063 |
| V-S | 0.0479614843 | 0.0185350644 | 0.0773879043 | 0.0000017 |
| W-S | -0.0540435847 | -0.0834641212 | -0.0246230482 | 0.0000000 |
| Y-S | -0.0016420988 | -0.0310275063 | 0.0277433087 | 1.0000000 |
| V-T | 0.0273061415 | -0.0021202784 | 0.0567325615 | 0.1070972 |
| W-T | -0.0746989275 | -0.1041194640 | -0.0452783910 | 0.0000000 |
| Y-T | -0.0222974416 | -0.0516828491 | 0.0070879659 | 0.4180775 |
| W-V | -0.1020050690 | -0.1314665691 | -0.0725435690 | 0.0000000 |
| Y-V | -0.0496035832 | -0.0790300031 | -0.0201771632 | 0.0000004 |
| Y-W | 0.0524014859 | 0.0229809494 | 0.0818220224 | 0.0000000 |

**Donor**

| | diff | lwr | upr | p adj |
|---|---|---|---|---|
| 3088202-3087423 | -0.009686784 | -0.0295486217 | 1.017505e-02 | 0.8747639 |
| 3091412-3087423. | -0.040182618 | -0.0600444554 | -2.032078e-02 | 0.0000000 |
| 310264-3087423 | -0.027216718 | -0.0470917219 | -7.341713e-03 | 0.0006227 |
| 310268-3087423 | -0.019910449 | -0.0397744766 | -4.642087e-05 | 0.0488963 |
| 310280-3087423 | -0.055138247 | -0.0750000843 | -3.527641e-02 | 0.0000000 |
| 4608-3087423 | -0.045885753 | -0.0657607575 | -2.601075e-02 | 0.0000000 |
| 8119-3087423 | -0.025544000 | -0.0454058378 | -5.682163e-03 | 0.0019297 |
| 8478-3087423 | -0.030891304 | -0.0507531412 | -1.102947e-02 | 0.0000378 |
| THP1-3087423 | -0.086540213 | -0.1064042410 | -6.667619e-02 | 0.0000000 |
| 3091412-3088202. | -0.030495834 | -0.0503576714 | -1.063400e-02 | 0.0000520 |
| 310264-3088202 | -0.017529934 | -0.0374049379 | 2.345071e-03 | 0.1394945 |
| 310268-3088202 | -0.010223665 | -0.0300876925 | 9.640363e-03 | 0.8344493 |
| 310280-3088202. | -0.045451463 | -0.0653133002 | -2.558962e-02 | 0.0000000 |
| 4608-3088202 | -0.036198969 | -0.0560739735 | -1.632396e-02 | 0.0000001 |
| 8119-3088202 | -0.015857216 | -0.0357190537 | 4.004622e-03 | 0.2543606 |
| 8478-3088202 | -0.021204519 | -0.0410663571 | -1.342682e-03 | 0.0254446 |
| THP1-3088202 | -0.076853429 | -0.0967174569 | -5.698940e-02 | 0.0000000 |
| 310264-3091412 | 0.012965900 | -0.0069091041 | 3.284090e-02 | 0.5531567 |
| 310268-3091412 | 0.020272169 | 0.0004081412 | 4.013620e-02 | 0.0409855 |
| 310280-3091412 | -0.014955629 | -0.0348174665 | 4.906209e-03 | 0.3363452 |
| 4608-3091412 | -0.005703135 | -0.0255781397 | 1.417187e-02 | 0.9962700 |
| 8119-3091412 | 0.014638618 | -0.0052232200 | 3.450046e-02 | 0.3680951 |
| 8478-3091412 | 0.009291314 | -0.0105705234 | 2.915315e-02 | 0.9003185 |
| | diff | lwr | upr | p adj |
| THP1-3091412 | -0.046357595 | -0.0662216232 | -2.649357e-02 | 0.0000000 |
| 310268-310264 | 0.007306269 | -0.0125709241 | 2.718346e-02 | 0.9777526 |
| 310280-310264 | -0.027921529 | -0.0477965333 | -8.046525e-03 | 0.0003758 |
| 4608-310264 | -0.018669036 | -0.0385571978 | 1.219127e-03 | 0.0873723 |
| 8119-310264 | 0.001672718 | -0.0182022868 | 2.154772e-02 | 0.9999999 |
| 8478-310264 | -0.003674586 | -0.0235495901 | 1.620042e-02 | 0.9998916 |
| THP1-310264 | -0.059323495 | -0.0792006885 | -3.944630e-02 | 0.0000000 |
| 310280-310268 | -0.035227798 | -0.0550918258 | -1.536377e-02 | 0.0000007 |
| 4608-310268 | -0.025975305 | -0.0458524975 | -6.098112e-03 | 0.0014675 |
| 8119-310268 | -0.005633551 | -0.0254975793 | 1.423048e-02 | 0.9965883 |
| 8478-310268 | -0.010980855 | -0.0308448827 | 8.883173e-03 | 0.7671006 |
| THP1-310268 | -0.066629764 | -0.0864959822 | -4.676355e-02 | 0.0000000 |
| 4608-310280 | 0.009252493 | -0.0106225109 | 2.912750e-02 | 0.9029945 |
| 8119-310280 | 0.029594246 | 0.0097324088 | 4.945608e-02 | 0.0001057 |
| 8478-310280 | 0.024246943 | 0.0043851055 | 4.410878e-02 | 0.0044310 |
| THP1-310280 | -0.031401966 | -0.0512659943 | -1.153794e-02 | 0.0000250 |
| 8119-4608 | 0.020341753 | 0.0004667488 | 4.021676e-02 | 0.0398183 |
| 8478-4608 | 0.014994450 | -0.0048805545 | 3.486945e-02 | 0.3335221 |
| THP1-4608 | -0.040654460 | -0.0605316529 | -2.077727e-02 | 0.0000000 |
| 8478-8119 | -0.005347303 | -0.0252091410 | 1.451453e-02 | 0.9977135 |
| THP1-8119 | -0.060996213 | -0.0808602408 | -4.113219e-02 | 0.0000000 |
| THP1-8478 | -0.055648910 | -0.0755129374 | -3.578488e-02 | 0.0000000 |

**B Soluble peptides with cells on tissue culture plastic, 48 hour time point only**

| 48 hours | Df | Sum Sq | Mean Sq | F value | Pr(>F) |
|---|---|---|---|---|---|
| celltype | 2 | 279.3 | 139.65 | 1952.84 | <2e-16 *** |
| donor | 7 | 12.4 | 1.77 | 24.73 | <2e-16 *** |
| experiment | 7 | 22.7 | 3.24 | 45.34 | <2e-16 *** |
| endgroup | 6 | 376.8 | 62.79 | 878.08 | <2e-16 *** |
| amino_acid | 17 | 36.4 | 2.14 | 29.93 | <2e-16 *** |
| residuals | 22626 | 1618 | 0.07 | | |

Tukey multiple comparisons of means
95% family-wise confidence level

Fit: aov(formula = ave ~ cell + donor + experiment + endgroup + amino_acid)

| **Cell type** | diff | lwr | upr | p adj |
|---|---|---|---|---|
| hUVEC-hMSC | 0.18174843 | 0.17099723 | 0.19249964 | 0 |
| Mac-hMSC | 0.26609418 | 0.25603896 | 0.27614940 | 0 |
| Mac-hUVEC | 0.08434575 | 0.07429095 | 0.09440055 | 0 |

| **Endgroup** | diff | lwr | upr | p adj |
|---|---|---|---|---|
| Am-CBA | -0.035178958 | -0.05477083 | -0.01558708 | 0.0000023 |
| COOH-CBA | -0.147212545 | -0.16680442 | -0.12762067 | 0.0000000 |
| AcBA-CBA | -0.004391847 | -0.02398523 | 0.01520154 | 0.9946058 |
| Ac-CBA | -0.036389911 | -0.05599087 | -0.01678895 | 0.0000007 |
| NBA-CBA | -0.093876888 | -0.11346876 | -0.07428501 | 0.0000000 |
| NH2-CBA | -0.389396425 | -0.40899738 | -0.36979547 | 0.0000000 |
| COOH-Am | -0.112033586 | -0.13162395 | -0.09244322 | 0.0000000 |
| AcBA-Am | 0.030787111 | 0.01119524 | 0.05037899 | 0.0000737 |
| Ac-Am | -0.001210953 | -0.02081040 | 0.01838849 | 0.9999969 |
| NBA-Am | -0.058697929 | -0.07828829 | -0.03910757 | 0.0000000 |
| NH2-Am | -0.354217467 | -0.37381691 | -0.33461802 | 0.0000000 |
| AcBA-COOH | 0.142820698 | 0.12322882 | 0.16241257 | 0.0000000 |
| Ac-COOH | 0.110822633 | 0.09122319 | 0.13042208 | 0.0000000 |
| NBA-COOH | 0.053335657 | 0.03374529 | 0.07292602 | 0.0000000 |
| NH2-COOH | -0.242183880 | -0.26178333 | -0.22258443 | 0.0000000 |
| Ac-AcBA | -0.031998064 | -0.05159902 | -0.01239711 | 0.0000305 |
| NBA-AcBA | -0.089485041 | -0.10907691 | -0.06989317 | 0.0000000 |
| NH2-AcBA | -0.385004578 | -0.40460554 | -0.36540362 | 0.0000000 |
| NBA-Ac | -0.057486976 | -0.07708642 | -0.03788753 | 0.0000000 |
| NH2-Ac | -0.353006513 | -0.37261504 | -0.33339799 | 0.0000000 |
| NH2-NBA | -0.295519537 | -0.31511898 | -0.27592009 | 0.0000000 |

**Amino acid**

| | diff | lwr | upr | p adj |
|---|---|---|---|---|
| D-A | 0.0911418367 | 0.0539687514 | 0.1283149221 | 0.0000000 |
| E-A | 0.0674525839 | 0.0302794985 | 0.1046256693 | 0.0000000 |
| F-A | -0.0285623966 | -0.0657354820 | 0.0086106888 | 0.3938383 |
| G-A | 0.0074636879 | -0.0297093975 | 0.0446367732 | 0.9999997 |
| H-A | -0.0193680806 | -0.0565411660 | 0.0178050047 | 0.9393452 |
| I_L-A | 0.0150238223 | -0.0221492630 | 0.0521969077 | 0.9953836 |
| K-A | -0.0251598995 | -0.0623329849 | 0.0120131859 | 0.6374478 |
| M-A | -0.0071542074 | -0.0443346735 | 0.0300262587 | 0.9999999 |
| N-A | 0.0307876787 | -0.0063854067 | 0.0679607640 | 0.2581939 |
| P-A | 0.0493901460 | 0.0122170606 | 0.0865632314 | 0.0005071 |
| Q-A | -0.0006024072 | -0.0377754926 | 0.0365706782 | 1.0000000 |
| R-A | -0.0697643361 | -0.1069374215 | -0.0325912507 | 0.0000000 |
| S-A | -0.0104474036 | -0.0476204890 | 0.0267256818 | 0.9999583 |
| T-A | 0.0102079392 | -0.0269651462 | 0.0473810246 | 0.9999700 |
| V-A | 0.0375140807 | 0.0002891138 | 0.0747390476 | 0.0457894 |
| W-A | -0.0644909884 | -0.1017085126 | -0.0272734641 | 0.0000000 |
| Y-A | -0.0120895025 | -0.0492625879 | 0.0250835829 | 0.9996903 |
| E-D | -0.0236892528 | -0.0608623382 | 0.0134838325 | 0.7380570 |
| F-D | -0.1197042334 | -0.1568773188 | -0.0825311480 | 0.0000000 |
| G-D | -0.0836781489 | -0.1208512343 | -0.0465050635 | 0.0000000 |
| H-D | -0.1105099174 | -0.1476830028 | -0.0733368320 | 0.0000000 |
| I_L-D | -0.0761180144 | -0.1132910998 | -0.0389449290 | 0.0000000 |
| K-D | -0.1163017363 | -0.1534748217 | -0.0791286509 | 0.0000000 |
| M-D | -0.0982960442 | -0.1354765103 | -0.0611155780 | 0.0000000 |
| N-D | -0.0603541581 | -0.0975272435 | -0.0231810727 | 0.0000020 |
| P-D | -0.0417516908 | -0.0789247762 | -0.0045786054 | 0.0109361 |
| Q-D | -0.0917442439 | -0.1289173293 | -0.0545711585 | 0.0000000 |
| R-D | -0.1609061728 | -0.1980792582 | -0.1237330874 | 0.0000000 |
| S-D | -0.1015892404 | -0.1387623258 | -0.0644161550 | 0.0000000 |
| T-D | -0.0809338976 | -0.1181069830 | -0.0437608122 | 0.0000000 |
| V-D | -0.0536277561 | -0.0908527230 | -0.0164027892 | 0.0000738 |
| W-D | -0.1556328251 | -0.1928503493 | -0.1184153009 | 0.0000000 |
| Y-D | -0.1032313392 | -0.1404044246 | -0.0660582538 | 0.0000000 |
| F-E | -0.0960149805 | -0.1331880659 | -0.0588418952 | 0.0000000 |
| G-E | -0.0599888960 | -0.0971619814 | -0.0228158107 | 0.0000025 |
| H-E | -0.0868206645 | -0.1239937499 | -0.0496475792 | 0.0000000 |
| I_L-E | -0.0524287616 | -0.0896018469 | -0.0152556762 | 0.0001263 |
| K-E | -0.0926124834 | -0.1297855688 | -0.0554393981 | 0.0000000 |
| M-E | -0.0746067913 | -0.1117872574 | -0.0374263252 | 0.0000000 |
| N-E | -0.0366649053 | -0.0738379906 | 0.0005081801 | 0.0582158 |
| P-E | -0.0180624379 | -0.0552355233 | 0.0191106475 | 0.9678116 |
| Q-E | -0.0680549911 | -0.1052280765 | -0.0308819057 | 0.0000000 |
| R-E | -0.1372169200 | -0.1743900054 | -0.1000438346 | 0.0000000 |
| S-E | -0.0778999875 | -0.1150730729 | -0.0407269021 | 0.0000000 |
| T-E | -0.0572446447 | -0.0944177301 | -0.0200715594 | 0.0000115 |
| V-E | -0.0299385032 | -0.0671634701 | 0.0072864637 | 0.3088003 |
| W-E | -0.1319435723 | -0.1691610965 | -0.0947260480 | 0.0000000 |
| Y-E | -0.0795420864 | -0.1167151718 | -0.0423690010 | 0.0000000 |
| G-F | 0.0360260845 | -0.0011470009 | 0.0731991699 | 0.0701352 |
| H-F | 0.0091943160 | -0.0279787694 | 0.0463674014 | 0.9999934 |
| I_L-F | 0.0435862190 | 0.0064131336 | 0.0807593044 | 0.0055425 |
| K-F | 0.0034024971 | -0.0337705883 | 0.0405755825 | 1.0000000 |

| | | | | |
|---|---|---|---|---|
| M-F | 0.0214081892 | -0.0157722769 | 0.0585886554 | 0.8654509 |
| N-F | 0.0593500753 | 0.0221769899 | 0.0965231607 | 0.0000036 |
| P-F | 0.0779525426 | 0.0407794572 | 0.1151256280 | 0.0000000 |
| Q-F | 0.0279599894 | -0.0092130959 | 0.0651330748 | 0.4352011 |
| R-F | -0.0412019394 | -0.0783750248 | -0.0040288541 | 0.0133051 |
| S-F | 0.0181149930 | -0.0190580924 | 0.0552880784 | 0.9669115 |
| T-F | 0.0387703358 | 0.0015972504 | 0.0759434212 | 0.0303156 |
| V-F | 0.0660764773 | 0.0288515104 | 0.1033014442 | 0.0000000 |
| W-F | -0.0359285917 | -0.0731461160 | 0.0012889325 | 0.0730109 |
| Y-F | 0.0164728942 | -0.0207001912 | 0.0536459796 | 0.9872413 |
| H-G | -0.0268317685 | -0.0640048539 | 0.0103413169 | 0.5159264 |
| I_L-G | 0.0075601345 | -0.0296129509 | 0.0447332199 | 0.9999997 |
| K-G | -0.0326235874 | -0.0697966728 | 0.0045494980 | 0.1714720 |
| M-G | -0.0146178953 | -0.0517983614 | 0.0225625709 | 0.9966506 |
| N-G | 0.0233239908 | -0.0138490946 | 0.0604970762 | 0.7611846 |
| P-G | 0.0419264581 | 0.0047533727 | 0.0790995435 | 0.0102676 |
| Q-G | -0.0080660950 | -0.0452391804 | 0.0291069903 | 0.9999991 |
| R-G | -0.0772280239 | -0.1144011093 | -0.0400549386 | 0.0000000 |
| S-G | -0.0179110915 | -0.0550841769 | 0.0192619939 | 0.9702995 |
| T-G | 0.0027442513 | -0.0344288341 | 0.0399173367 | 1.0000000 |
| V-G | 0.0300503928 | -0.0071745741 | 0.0672753597 | 0.3021633 |
| W-G | -0.0719546762 | -0.1091722005 | -0.0347371520 | 0.0000000 |
| Y-G | -0.0195531903 | -0.0567262757 | 0.0176198951 | 0.9341904 |
| I_L-H | 0.0343919030 | -0.0027811824 | 0.0715649884 | 0.1100495 |
| K-H | -0.0057918189 | -0.0429649043 | 0.0313812665 | 1.0000000 |
| M-H | 0.0122138732 | -0.0249665929 | 0.0493943394 | 0.9996463 |
| N-H | 0.0501557593 | 0.0129826739 | 0.0873288447 | 0.0003604 |
| P-H | 0.0687582266 | 0.0315851412 | 0.1059313120 | 0.0000000 |
| Q-H | 0.0187656735 | -0.0184074119 | 0.0559387588 | 0.9541257 |
| R-H | -0.0503962554 | -0.0875693408 | -0.0132231701 | 0.0003233 |
| S-H | 0.0089206770 | -0.0282524084 | 0.0460937624 | 0.9999958 |
| T-H | 0.0295760198 | -0.0075970656 | 0.0667491052 | 0.3282941 |
| V-H | 0.0568821613 | 0.0196571944 | 0.0941071282 | 0.0000145 |
| W-H | -0.0451229077 | -0.0823404320 | -0.0079053835 | 0.0031128 |
| Y-H | 0.0072785782 | -0.0298945072 | 0.0444516636 | 0.9999998 |
| K-I_L | -0.0401837219 | -0.0773568073 | -0.0030106365 | 0.0189500 |
| M-I_L | -0.0221780298 | -0.0593584959 | 0.0150024364 | 0.8274584 |
| N-I_L | 0.0157638563 | -0.0214092291 | 0.0529369417 | 0.9920654 |
| P-I_L | 0.0343663236 | -0.0028067618 | 0.0715394090 | 0.1107945 |
| Q-I_L | -0.0156262295 | -0.0527993149 | 0.0215468559 | 0.9927993 |
| R-I_L | -0.0847881584 | -0.1219612438 | -0.0476150730 | 0.0000000 |
| S-I_L | -0.0254712260 | -0.0626443114 | 0.0117018594 | 0.6150779 |
| T-I_L | -0.0048158832 | -0.0419889686 | 0.0323572022 | 1.0000000 |
| V-I_L | 0.0224902583 | -0.0147347086 | 0.0597152252 | 0.8120325 |
| W-I_L | -0.0795148107 | -0.1167323349 | -0.0422972865 | 0.0000000 |
| Y-I_L | -0.0271133248 | -0.0642864102 | 0.0100597606 | 0.4955013 |
| M-K | 0.0180056921 | -0.0191747740 | 0.0551861583 | 0.9688215 |
| N-K | 0.0559475782 | 0.0187744928 | 0.0931206636 | 0.0000225 |
| P-K | 0.0745500455 | 0.0373769601 | 0.1117231309 | 0.0000000 |
| Q-K | 0.0245574924 | -0.0126155930 | 0.0617305777 | 0.6798651 |
| R-K | -0.0446044365 | -0.0817775219 | -0.0074313512 | 0.0037393 |
| S-K | 0.0147124959 | -0.0224605895 | 0.0518855813 | 0.9963790 |
| T-K | 0.0353678387 | -0.0018052467 | 0.0725409241 | 0.0844742 |
| V-K | 0.0626739802 | 0.0254490133 | 0.0998989471 | 0.0000004 |

| | | | | |
|---|---|---|---|---|
| W-K | -0.0393310888 | -0.0765486131 | -0.0021135646 | 0.0256295 |
| Y-K | 0.0130703971 | -0.0241026883 | 0.0502434825 | 0.9991467 |
| N-M | 0.0379418861 | 0.0007614199 | 0.0751223522 | 0.0395549 |
| P-M | 0.0565443534 | 0.0193638873 | 0.0937248195 | 0.0000166 |
| Q-M | 0.0065518002 | -0.0306286659 | 0.0437322663 | 1.0000000 |
| R-M | -0.0626101287 | -0.0997905948 | -0.0254296626 | 0.0000004 |
| S-M | -0.0032931962 | -0.0404736624 | 0.0338872699 | 1.0000000 |
| T-M | 0.0173621466 | -0.0198183196 | 0.0545426127 | 0.9781475 |
| V-M | 0.0446682881 | 0.0074359507 | 0.0819006254 | 0.0037499 |
| W-M | -0.0573367809 | -0.0945616771 | -0.0201118848 | 0.0000114 |
| Y-M | -0.0049352951 | -0.0421157612 | 0.0322451711 | 1.0000000 |
| P-N | 0.0186024673 | -0.0185706181 | 0.0557755527 | 0.9576289 |
| Q-N | -0.0313900858 | -0.0685631712 | 0.0057829996 | 0.2270870 |
| R-N | -0.1005520147 | -0.1377251001 | -0.0633789293 | 0.0000000 |
| S-N | -0.0412350823 | -0.0784081677 | -0.0040619969 | 0.0131501 |
| T-N | -0.0205797395 | -0.0577528249 | 0.0165933459 | 0.9000136 |
| V-N | 0.0067264020 | -0.0304985649 | 0.0439513689 | 0.9999999 |
| W-N | -0.0952786670 | -0.1324961912 | -0.0580611428 | 0.0000000 |
| Y-N | -0.0428771811 | -0.0800502665 | -0.0057040957 | 0.0072404 |
| Q-P | -0.0499925532 | -0.0871656386 | -0.0128194678 | 0.0003878 |
| R-P | -0.1191544821 | -0.1563275675 | -0.0819813967 | 0.0000000 |
| S-P | -0.0598375496 | -0.0970106350 | -0.0226644642 | 0.0000027 |
| T-P | -0.0391822068 | -0.0763552922 | -0.0020091214 | 0.0265046 |
| V-P | -0.0118760653 | -0.0491010322 | 0.0253489016 | 0.9997602 |
| W-P | -0.1138811343 | -0.1510986586 | -0.0766636101 | 0.0000000 |
| Y-P | -0.0614796485 | -0.0986527338 | -0.0243065631 | 0.0000010 |
| R-Q | -0.0691619289 | -0.1063350143 | -0.0319888435 | 0.0000000 |
| S-Q | -0.0098449964 | -0.0470180818 | 0.0273280889 | 0.9999822 |
| T-Q | 0.0108103464 | -0.0263627390 | 0.0479834317 | 0.9999325 |
| V-Q | 0.0381164879 | 0.0008915210 | 0.0753414548 | 0.0379863 |
| W-Q | -0.0638885812 | -0.1011061054 | -0.0266710569 | 0.0000001 |
| Y-Q | -0.0114870953 | -0.0486601807 | 0.0256859901 | 0.9998442 |
| S-R | 0.0593169325 | 0.0221438471 | 0.0964900178 | 0.0000037 |
| T-R | 0.0799722752 | 0.0427991899 | 0.1171453606 | 0.0000000 |
| V-R | 0.1072784168 | 0.0700534499 | 0.1445033837 | 0.0000000 |
| W-R | 0.0052733477 | -0.0319441765 | 0.0424908720 | 1.0000000 |
| Y-R | 0.0576748336 | 0.0205017482 | 0.0948479190 | 0.0000091 |
| T-S | 0.0206553428 | -0.0165177426 | 0.0578284282 | 0.8971104 |
| V-S | 0.0479614843 | 0.0107365174 | 0.0851864512 | 0.0009712 |
| W-S | -0.0540435847 | -0.0912611090 | -0.0168260605 | 0.0000600 |
| Y-S | -0.0016420988 | -0.0388151842 | 0.0355309866 | 1.0000000 |
| V-T | 0.0273061415 | -0.0099188254 | 0.0645311084 | 0.4843365 |
| W-T | -0.0746989275 | -0.1119164518 | -0.0374814033 | 0.0000000 |
| Y-T | -0.0222974416 | -0.0594705270 | 0.0148756438 | 0.8208499 |
| W-V | -0.1020050690 | -0.1392744129 | -0.0647357252 | 0.0000000 |
| Y-V | -0.0496035832 | -0.0868285501 | -0.0123786163 | 0.0004757 |
| Y-W | 0.0524014859 | 0.0151839616 | 0.0896190101 | 0.0001318 |

| Donor | diff | lwr | upr | p adj |
|---|---|---|---|---|
| 3088202-3087423 | -0.009686784 | -0.0348123771 | 0.0154388090 | 0.9694090 |
| 3091412-3087423 | -0.040182618 | -0.0653082108 | -0.0150570248 | 0.0000184 |
| 310264-3087423 | -0.027216718 | -0.0523589667 | -0.0020744686 | 0.0217447 |
| 310268-3087423 | -0.019910449 | -0.0450388124 | 0.0052179149 | 0.2644817 |
| 310280-3087423 | -0.055138247 | -0.0802638397 | -0.0300126536 | 0.0000000 |
| 4608-3087423 | -0.045885753 | -0.0710280023 | -0.0207435042 | 0.0000001 |
| 8119-3087423 | -0.025544000 | -0.0506695932 | -0.0004184071 | 0.0425781 |
| 8478-3087423 | -0.030891304 | -0.0560168965 | -0.0057657105 | 0.0039781 |
| THP1-3087423 | -0.086540213 | -0.1116685768 | -0.0614118494 | 0.0000000 |
| 3091412-3088202 | -0.030495834 | -0.0556214267 | -0.0053702407 | 0.0048322 |
| 310264-3088202 | -0.017529934 | -0.0426721826 | 0.0076123154 | 0.4524902 |
| 310268-3088202 | -0.010223665 | -0.0353520283 | 0.0149046990 | 0.9567040 |
| 310280-3088202 | -0.045451463 | -0.0705770556 | -0.0203258696 | 0.0000002 |
| 4608-3088202 | -0.036198969 | -0.0613412182 | -0.0110567202 | 0.0002255 |
| 8119-3088202 | -0.015857216 | -0.0409828091 | 0.0092683769 | 0.6012720 |
| 8478-3088202 | -0.021204519 | -0.0463301125 | 0.0039210736 | 0.1858617 |
| THP1-3088202 | -0.076853429 | -0.1019817927 | -0.0517250654 | 0.0000000 |
| 310264-3091412 | 0.012965900 | -0.0121763489 | 0.0381081492 | 0.8328010 |
| 310268-3091412 | 0.020272169 | -0.0048561946 | 0.0454005327 | 0.2407125 |
| 310280-3091412 | -0.014955629 | -0.0400812219 | 0.0101699642 | 0.6805488 |
| 4608-3091412 | -0.005703135 | -0.0308453845 | 0.0194391136 | 0.9994131 |
| 8119-3091412 | 0.014638618 | -0.0104869754 | 0.0397642106 | 0.7073104 |
| 8478-3091412 | 0.009291314 | -0.0158342788 | 0.0344169073 | 0.9768263 |
| THP1-3091412 | -0.046357595 | -0.0714859590 | -0.0212292317 | 0.0000000 |
| 310268-310264 | 0.007306269 | -0.0178387489 | 0.0324512868 | 0.9958988 |
| 310280-310264 | -0.027921529 | -0.0530637780 | -0.0027792800 | 0.0160568 |
| 4608-310264 | -0.018669036 | -0.0438279296 | 0.0064898584 | 0.3579557 |
| 8119-310264 | 0.001672718 | -0.0234695315 | 0.0268149665 | 1.0000000 |
| 8478-310264 | -0.003674586 | -0.0288168349 | 0.0214676632 | 0.9999854 |
| THP1-310264 | -0.059323495 | -0.0844685133 | -0.0341784776 | 0.0000000 |
| 310280-310268 | -0.035227798 | -0.0603561616 | -0.0100994343 | 0.0003921 |
| 4608-310268 | -0.025975305 | -0.0511203224 | -0.0008302867 | 0.0362244 |
| 8119-310268 | -0.005633551 | -0.0307619151 | 0.0194948122 | 0.9994664 |
| 8478-310268 | -0.010980855 | -0.0361092185 | 0.0141475089 | 0.9327976 |
| THP1-310268 | -0.066629764 | -0.0917608984 | -0.0414986304 | 0.0000000 |
| 4608-310280 | 0.009252493 | -0.0158897556 | 0.0343947424 | 0.9775736 |
| 8119-310280 | 0.029594246 | 0.0044686535 | 0.0547198395 | 0.0074457 |
| 8478-310280 | 0.024246943 | -0.0008786499 | 0.0493725361 | 0.0691966 |
| THP1-310280 | -0.031401966 | -0.0565303301 | -0.0062736028 | 0.0030867 |
| 8119-4608 | 0.020341753 | -0.0048004959 | 0.0454840021 | 0.2370098 |
| 8478-4608 | 0.014994450 | -0.0101477993 | 0.0401366988 | 0.6780747 |
| THP1-4608 | -0.040654460 | -0.0657994777 | -0.0155094420 | 0.0000137 |
| 8478-8119 | -0.005347303 | -0.0304728964 | 0.0197782897 | 0.9996511 |
| THP1-8119 | -0.060996213 | -0.0861245766 | -0.0358678493 | 0.0000000 |
| THP1-8478 | -0.055648910 | -0.0807772733 | -0.0305205459 | 0.0000000 |

### C Concentration study peptides at 48 hours

| 48 hours | Df | Sum Sq | Mean Sq | F value | Pr(>F) |
|---|---|---|---|---|---|
| celltype | 2 | 7.184 | 3.592 | 97.36 | <2e-16 *** |
| endgroup | 6 | 21.549 | 3.591 | 97.34 | <2e-16 *** |
| concentration | 4 | 3.989 | 0.997 | 27.03 | <2e-16 *** |
| residuals | 302 | 11.142 | 0.037 | | |

Tukey multiple comparisons of means
95% family-wise confidence level
Fit: aov(formula = ave ~ Cell + endgroup + concentration, data = concentration_48_stats)

**Cell type**

```
                    diff        lwr        upr     p adj
hUVEC-hMSC       0.02674739 -0.0356911 0.08918588 0.5717588
macrophage-hMSC  0.33290148  0.2704630 0.39533997 0.0000000
macrophage-hUVEC 0.30615409  0.2437156 0.36859258 0.0000000
```

**Endgroup**

```
                diff         lwr         upr     p adj
Ac_BA-Ac   -0.02869210 -0.14889144  0.09150724 0.9920278
Am-Ac      -0.30279864 -0.42299798 -0.18259929 0.0000000
CBA-Ac     -0.05351273 -0.17371207  0.06668661 0.8415104
COOH-Ac    -0.49469442 -0.61489376 -0.37449508 0.0000000
NBA-Ac     -0.36386417 -0.48406351 -0.24366483 0.0000000
NH2-Ac     -0.76234755 -0.88254689 -0.64214820 0.0000000
Am-Ac_BA   -0.27410654 -0.39430588 -0.15390719 0.0000000
CBA-Ac_BA  -0.02482063 -0.14501997  0.09537871 0.9963800
COOH-Ac_BA -0.46600232 -0.58620166 -0.34580298 0.0000000
NBA-Ac_BA  -0.33517207 -0.45537141 -0.21497273 0.0000000
NH2-Ac_BA  -0.73365544 -0.85385479 -0.61345610 0.0000000
CBA-Am      0.24928590  0.12908656  0.36948525 0.0000000
COOH-Am    -0.19189578 -0.31209513 -0.07169644 0.0000675
NBA-Am     -0.06106553 -0.18126487  0.05913381 0.7399153
NH2-Am     -0.45954891 -0.57974825 -0.33934957 0.0000000
COOH-CBA   -0.44118169 -0.56138103 -0.32098234 0.0000000
NBA-CBA    -0.31035144 -0.43055078 -0.19015209 0.0000000
NH2-CBA    -0.70883481 -0.82903416 -0.58863547 0.0000000
NBA-COOH    0.13083025  0.01063091  0.25102959 0.0229651
NH2-COOH   -0.26765313 -0.38785247 -0.14745378 0.0000000
NH2-NBA    -0.39848338 -0.51868272 -0.27828403 0.0000000
```

**Concentration**

```
                diff         lwr         upr     p adj
19.5-1250 -0.120196418 -0.21411722 -0.02627562 0.0046316
312-1250  -0.067417531 -0.16133833  0.02650327 0.2832901
5000-1250  0.186144359  0.09222356  0.28006516 0.0000011
78-1250   -0.110330119 -0.20425092 -0.01640932 0.0121507
312-19.5   0.052778887 -0.04114191  0.14669968 0.5358579
```

```
5000-19.5  0.306340777  0.21241998  0.40026157 0.0000000
78-19.5    0.009866299 -0.08405450  0.10378710 0.9984852
5000-312   0.253561889  0.15964109  0.34748269 0.0000000
78-312    -0.042912589 -0.13683339  0.05100821 0.7195936
78-5000   -0.296474478 -0.39039527 -0.20255368 0.0000000
```

### D <u>LIAANK peptides at 48 hours</u>

| 48 hours | Df | Sum Sq | Mean Sq | F value | Pr(>F) |
|---|---|---|---|---|---|
| celltype | 2 | 0.082 | 0.0409 | 2.13 | 0.129 |
| endgroup | 6 | 5.773 | 0.9621 | 50.11 | <2e-16 *** |
| residuals | 54 | 1.037 | 0.0192 | | |

Tukey multiple comparisons of means
95% family-wise confidence level
Fit: aov(formula = ave ~ celltype + endgroup)

| **<u>Cell type</u>** | diff | lwr | upr | p adj |
|---|---|---|---|---|
| Macrophage-hUVEC | -0.072990023 | -0.1760419 | 0.03006182 | 0.2119027 |
| hMSC-hUVEC. | -0.079463872 | -0.1825157 | 0.02358797 | 0.1607616 |
| hMSC-Macrophage | -0.006473849 | -0.1095257 | 0.09657800 | 0.9874458 |

| **<u>Endgroup</u>** | diff | lwr | upr | p adj |
|---|---|---|---|---|
| Ac-AcBA | 0.07693082 | -0.123081655 | 0.27694329 | 0.8994233 |
| NBA-AcBA | -0.35614312 | -0.556155600 | -0.15613065 | 0.0000252 |
| Am-AcBA | 0.13334378 | -0.066668695 | 0.33335625 | 0.4015930 |
| CBA-AcBAG | 0.19615704 | -0.003855434 | 0.39616952 | 0.0579509 |
| COOH-AcBA | -0.45166561 | -0.651678083 | -0.25165313 | 0.0000001 |
| NH2-AcBA | -0.63354877 | -0.833561246 | -0.43353630 | 0.0000000 |
| NBA-Ac | -0.43307394 | -0.633086420 | -0.23306147 | 0.0000003 |
| Am-Ac | 0.05641296 | -0.143599515 | 0.25642543 | 0.9763913 |
| CBA-Ac | 0.11922622 | -0.080786254 | 0.31923870 | 0.5373659 |
| COOH-Ac | -0.52859643 | -0.728608903 | -0.32858395 | 0.0000000 |
| NH2-Ac | -0.71047959 | -0.910492066 | -0.51046712 | 0.0000000 |
| Am-NBA | 0.48948691 | 0.289474430 | 0.68949938 | 0.0000000 |
| CBA-NBA | 0.55230017 | 0.352287691 | 0.75231264 | 0.0000000 |
| COOH-NBA | -0.09552248 | -0.295534958 | 0.10448999 | 0.7652009 |
| NH2-NBA | -0.27740565 | -0.477418121 | -0.07739317 | 0.0015813 |
| CBA-Am | 0.06281326 | -0.137199214 | 0.26282574 | 0.9599996 |
| COOH-Am | -0.58500939 | -0.785021863 | -0.38499691 | 0.0000000 |
| NH2G-Am | -0.76689255 | -0.966905026 | -0.56688008 | 0.0000000 |
| COOH-CBA | -0.64782265 | -0.847835124 | -0.44781017 | 0.0000000 |
| NH2-CBA | -0.82970581 | -1.029718287 | -0.62969334 | 0.0000000 |
| NH2-COOH | -0.18188316 | -0.381895638 | 0.01812931 | 0.0975390 |

### E IVKVA peptides at 48 hours

| 48 hours | Df | Sum Sq | Mean Sq | F value | Pr(>F) |
|---|---|---|---|---|---|
| celltype | 2 | 0.263 | 0.1314 | 2.027 | 0.141577 |
| endgroup | 6 | 1.853 | 0.3089 | 4.767 | 0.000578 |
| residuals | 54 | 3.499 | 0.0648 | | |

Tukey multiple comparisons of means
95% family-wise confidence level

Fit: aov(formula = ave ~ celltype + endgroup)

| **Celltype** | diff | lwr | upr | p adj |
|---|---|---|---|---|
| Macrophage-hUVEC | -0.01503481 | -0.20436175 | 0.1742921 | 0.9800181 |
| hMSC-hUVEC | 0.12886108 | -0.06046586 | 0.3181880 | 0.2377607 |
| hMSC-Macrophage | 0.14389589 | -0.04543105 | 0.3332228 | 0.1690231 |

| **Endgroup** | diff | lwr | upr | p adj |
|---|---|---|---|---|
| Ac-AcBA | 0.30851472 | -0.05894836 | 0.6759778073 | 0.1555552 |
| NBA-AcBA | -0.25471927 | -0.62218235 | 0.1127438195 | 0.3546125 |
| Am-AcBA | -0.02456120 | -0.39202429 | 0.3429018828 | 0.9999932 |
| CBA-AcBA | 0.09053903 | -0.27692406 | 0.4580021099 | 0.9881916 |
| COOH-AcBA | -0.18941059 | -0.55687368 | 0.1780524904 | 0.6962825 |
| NH2-AcBA | -0.05986569 | -0.42732878 | 0.3075973939 | 0.9987708 |
| NBA-Ac | -0.56323399 | -0.93069707 | -0.1957709032 | 0.0003595 |
| Am-Ac | -0.33307592 | -0.70053901 | 0.0343871602 | 0.0995625 |
| CBA-Ac | -0.21797570 | -0.58543878 | 0.1494873873 | 0.5431482 |
| COOH-Ac | -0.49792532 | -0.86538840 | -0.1304622322 | 0.0021639 |
| NH2-Ac | -0.36838041 | -0.73584350 | -0.0009173288 | 0.0490434 |
| Am-NBA | 0.23015806 | -0.13730502 | 0.5976211480 | 0.4777630 |
| CBA-NBA | 0.34525829 | -0.02220479 | 0.7127213751 | 0.0786424 |
| COOH-NBA | 0.06530867 | -0.30215441 | 0.4327717556 | 0.9979950 |
| NH2-NBA | 0.19485357 | -0.17260951 | 0.5623166590 | 0.6678880 |
| CBA-Am | 0.11510023 | -0.25236286 | 0.4825633118 | 0.9604966 |
| COOH-Am | -0.16484939 | -0.53231248 | 0.2026136923 | 0.8130976 |
| NH2-Am | -0.03530449 | -0.40276757 | 0.3321585957 | 0.9999420 |
| COOH-CBA | -0.27994962 | -0.64741270 | 0.0875134652 | 0.2477438 |
| NH2-CBA | -0.15040472 | -0.51786780 | 0.2170583686 | 0.8695873 |
| NH2-COOH | 0.12954490 | -0.23791818 | 0.4970079881 | 0.9314373 |

**F Soluble peptides with cells encapsulated in PEG hydrogels at 48 hours**

| 48 hours | Df | Sum Sq | Mean Sq | F value | Pr(>F) |
|---|---|---|---|---|---|
| celltype | 2 | 8.19 | 4.095 | 125.42 | <2e-16 *** |
| endgroup | 6 | 74.10 | 12.350 | 378.28 | <2e-16 *** |
| amino_acid | 1081 | 5.87 | 0.345 | 10.58 | <2e-16 *** |
| residuals | 22626 | 35.29 | 0.033 | | |

Tukey multiple comparisons of means
95% family-wise confidence level
Fit: aov(formula = ave ~ cell + endgroup + amino_acid)

| **Cell type** | diff | lwr | upr | p adj |
|---|---|---|---|---|
| hUVEC-hMSC. | 0.05360174 | 0.02261208 | 0.08459139 | 0.0001559 |
| Macrophage-hMSC | 0.20456374 | 0.17320883 | 0.23591865 | 0.0000000 |
| Macrophage-hUVEC | 0.15096200 | 0.11962755 | 0.18229646 | 0.0000000 |

| **Endgroup** | diff | lwr | upr | p adj |
|---|---|---|---|---|
| AcBA-Ac | 0.03708977 | -0.02221344 | 0.09639299 | 0.5160796 |
| Am-Ac | -0.04452478 | -0.10392001 | 0.01487045 | 0.2885047 |
| CBA-Ac | 0.07091934 | 0.01161613 | 0.13022256 | 0.0078020 |
| COOH-Ac | -0.38434272 | -0.44641560 | -0.32226984 | 0.0000000 |
| NBA-Ac | -0.08609557 | -0.14539878 | -0.02679236 | 0.0003919 |
| NH2-Ac | -0.68822281 | -0.74752602 | -0.62891960 | 0.0000000 |
| Am-AcBA | -0.08161455 | -0.14100978 | -0.02221933 | 0.0010355 |
| CBA-AcBA | 0.03382957 | -0.02547364 | 0.09313278 | 0.6262622 |
| COOH-AcBA | -0.42143249 | -0.48350537 | -0.35935961 | 0.0000000 |
| NBA-AcBA | -0.12318534 | -0.18248856 | -0.06388213 | 0.0000000 |
| NH2-AcBA | -0.72531258 | -0.78461579 | -0.66600937 | 0.0000000 |
| CBA-Am | 0.11544412 | 0.05604890 | 0.17483935 | 0.0000003 |
| COOH-Am | -0.33981794 | -0.40197873 | -0.27765714 | 0.0000000 |
| NBA-Am | -0.04157079 | -0.10096602 | 0.01782444 | 0.3729682 |
| NH2-Am | -0.64369803 | -0.70309326 | -0.58430280 | 0.0000000 |
| COOH-CBA | -0.45526206 | -0.51733494 | -0.39318918 | 0.0000000 |
| NBA-CBA | -0.15701491 | -0.21631813 | -0.09771170 | 0.0000000 |
| NH2-CBA | -0.75914215 | -0.81844536 | -0.69983894 | 0.0000000 |
| NBA-COOH | 0.29824715 | 0.23617427 | 0.36032003 | 0.0000000 |
| NH2-COOH | -0.30388009 | -0.36595297 | -0.24180721 | 0.0000000 |
| NH2-NBA | -0.60212724 | -0.66143045 | -0.54282403 | 0.0000000 |

**Amino acid**

| | diff | lwr | upr | p adj |
|---|---|---|---|---|
| D-A | 0.1757707421 | 0.062276920 | 0.289264564 | 0.0000112 |
| E-A | 0.1257644436 | 0.012270622 | 0.239258265 | 0.0135034 |
| F-A | -0.0196719476 | -0.133165769 | 0.093821874 | 1.0000000 |
| G-A | 0.0399516470 | -0.073542175 | 0.153445469 | 0.9990828 |
| H-A | -0.0762467186 | -0.189740541 | 0.037247103 | 0.6465183 |
| I/L-A | 0.0768747060 | -0.036619116 | 0.190368528 | 0.6318774 |
| K-A | -0.0205584614 | -0.134052283 | 0.092935360 | 0.9999999 |
| M-A | 0.0501176495 | -0.063376172 | 0.163611471 | 0.9871761 |

| | | | | |
|---|---|---|---|---|
| N-A | 0.0738796244 | -0.039614197 | 0.187373446 | 0.7003139 |
| P-A | 0.1097088102 | -0.003785012 | 0.223202632 | 0.0718473 |
| Q-A | 0.0332618800 | -0.080231942 | 0.146755702 | 0.9999196 |
| R-A | -0.0392564012 | -0.156878612 | 0.078365810 | 0.9995371 |
| S-A | 0.0100754237 | -0.103418398 | 0.123569246 | 1.0000000 |
| T-A | 0.1035305588 | -0.010427452 | 0.217488570 | 0.1285330 |
| V-A | 0.0913764269 | -0.022117395 | 0.204870249 | 0.3043976 |
| W-A | -0.0839188068 | -0.197412629 | 0.029575015 | 0.4650114 |
| Y-A | -0.0678269774 | -0.181320799 | 0.045666845 | 0.8217720 |
| E-D | -0.0500062985 | -0.163500120 | 0.063487523 | 0.9874732 |
| F-D | -0.1954426897 | -0.308936512 | -0.081948868 | 0.0000004 |
| G-D | -0.1358190951 | -0.249312917 | -0.022325273 | 0.0040162 |
| H-D | -0.2520174607 | -0.365511283 | -0.138523639 | 0.0000000 |
| I/L-D | -0.0988960361 | -0.212389858 | 0.014597786 | 0.1798358 |
| K-D | -0.1963292035 | -0.309823025 | -0.082835382 | 0.0000003 |
| M-D | -0.1256530926 | -0.239146914 | -0.012159271 | 0.0136767 |
| N-D | -0.1018911177 | -0.215384940 | 0.011602704 | 0.1420506 |
| P-D | -0.0660619319 | -0.179555754 | 0.047431890 | 0.8513349 |
| Q-D | -0.1425088621 | -0.256002684 | -0.029015040 | 0.0016811 |
| R-D | -0.2150271433 | -0.332649354 | -0.097404933 | 0.0000000 |
| S-D | -0.1656953184 | -0.279189140 | -0.052201496 | 0.0000573 |
| T-D | -0.0722401832 | -0.186198194 | 0.041717828 | 0.7420538 |
| V-D | -0.0843943152 | -0.197888137 | 0.029099507 | 0.4539850 |
| W-D | -0.2596895489 | -0.373183371 | -0.146195727 | 0.0000000 |
| Y-D | -0.2435977194 | -0.357091541 | -0.130103898 | 0.0000000 |
| F-E | -0.1454363912 | -0.258930213 | -0.031942569 | 0.0011309 |
| G-E | -0.0858127966 | -0.199306618 | 0.027681025 | 0.4215805 |
| H-E | -0.2020111622 | -0.315504984 | -0.088517340 | 0.0000001 |
| I/L-E | -0.0488897376 | -0.162383559 | 0.064604084 | 0.9901557 |
| K-E | -0.1463229050 | -0.259816727 | -0.032829083 | 0.0010012 |
| M-E | -0.0756467940 | -0.189140616 | 0.037847028 | 0.6603793 |
| N-E | -0.0518848192 | -0.165378641 | 0.061609003 | 0.9816517 |
| P-E | -0.0160556334 | -0.129549455 | 0.097438188 | 1.0000000 |
| Q-E | -0.0925025635 | -0.205996385 | 0.020991258 | 0.2830723 |
| R-E | -0.1650208448 | -0.282643056 | -0.047398634 | 0.0001551 |
| S-E | -0.1156890198 | -0.229182842 | -0.002195198 | 0.0401467 |
| T-E | -0.0222338847 | -0.136191896 | 0.091724126 | 0.9999998 |
| V-E | -0.0343880166 | -0.147881839 | 0.079105805 | 0.9998728 |
| W-E | -0.2096832504 | -0.323177072 | -0.096189429 | 0.0000000 |
| Y-E | -0.1935914209 | -0.307085243 | -0.080097599 | 0.0000005 |
| G-F | 0.0596235946 | -0.053870227 | 0.173117416 | 0.9329647 |
| H-F | -0.0565747710 | -0.170068593 | 0.056919051 | 0.9577398 |
| I/L-F | 0.0965466536 | -0.016947168 | 0.210040475 | 0.2141716 |
| K-F | -0.0008865138 | -0.114380336 | 0.112607308 | 1.0000000 |
| M-F | 0.0697895971 | -0.043704225 | 0.183283419 | 0.7855322 |
| N-F | 0.0935515720 | -0.019942250 | 0.207045394 | 0.2640272 |
| P-F | 0.1293807578 | 0.015886936 | 0.242874580 | 0.0088518 |
| Q-F | 0.0529338276 | -0.060559994 | 0.166427650 | 0.9775717 |
| R-F | -0.0195844536 | -0.137206664 | 0.098037757 | 1.0000000 |
| S-F | 0.0297473713 | -0.083746451 | 0.143241193 | 0.9999835 |
| T-F | 0.1232025064 | 0.009244495 | 0.237160518 | 0.0190700 |
| V-F | 0.1110483745 | -0.002445447 | 0.224542196 | 0.0633443 |
| W-F | -0.0642468592 | -0.177740681 | 0.049246963 | 0.8785525 |
| Y-F | -0.0481550298 | -0.161648852 | 0.065338792 | 0.9916496 |

| | | | | |
|---|---|---|---|---|
| H-G | -0.1161983656 | -0.229692187 | -0.002704544 | 0.0381172 |
| I/L-G | 0.0369230590 | -0.076570763 | 0.150416881 | 0.9996690 |
| K-G | -0.0605101084 | -0.174003930 | 0.052983713 | 0.9241648 |
| M-G | 0.0101660025 | -0.103327819 | 0.123659824 | 1.0000000 |
| N-G | 0.0339279774 | -0.079565844 | 0.147421799 | 0.9998943 |
| P-G | 0.0697571632 | -0.043736659 | 0.183250985 | 0.7861582 |
| Q-G | -0.0066897669 | -0.120183589 | 0.106804055 | 1.0000000 |
| R-G | -0.0792080482 | -0.196830259 | 0.038414163 | 0.6423060 |
| S-G | -0.0298762233 | -0.143370045 | 0.083617599 | 0.9999824 |
| T-G | 0.0635789119 | -0.050379099 | 0.177536923 | 0.8911749 |
| V-G | 0.0514247799 | -0.062069042 | 0.164918602 | 0.9832443 |
| W-G | -0.1238704538 | -0.237364276 | -0.010376632 | 0.0167386 |
| Y-G | -0.1077786243 | -0.221272446 | 0.005715198 | 0.0857491 |
| I/L-H | 0.1531214246 | 0.039627603 | 0.266615246 | 0.0003828 |
| K-H | 0.0556882572 | -0.057805565 | 0.169182079 | 0.9634656 |
| M-H | 0.1263643682 | 0.012870546 | 0.239858190 | 0.0126036 |
| N-H | 0.1501263430 | 0.036632521 | 0.263620165 | 0.0005881 |
| P-H | 0.1859555288 | 0.072461707 | 0.299449351 | 0.0000020 |
| Q-H | 0.1095085987 | -0.003985223 | 0.223002421 | 0.0731963 |
| R-H | 0.0369903174 | -0.080631893 | 0.154612528 | 0.9997893 |
| S-H | 0.0863221424 | -0.027171680 | 0.199815964 | 0.4101494 |
| T-H | 0.1797772775 | 0.065819266 | 0.293735289 | 0.0000065 |
| V-H | 0.1676231456 | 0.054129324 | 0.281116967 | 0.0000422 |
| W-H | -0.0076720882 | -0.121165910 | 0.105821734 | 1.0000000 |
| Y-H | 0.0084197413 | -0.105074081 | 0.121913563 | 1.0000000 |
| K-I/L | -0.0974331674 | -0.210926989 | 0.016060654 | 0.2007222 |
| M-I/L | -0.0267570565 | -0.140250878 | 0.086736765 | 0.9999965 |
| N-I/L | -0.0029950816 | -0.116488903 | 0.110498740 | 1.0000000 |
| P-I/L | 0.0328341042 | -0.080659718 | 0.146327926 | 0.9999328 |
| Q-I/L | -0.0436128260 | -0.157106648 | 0.069880996 | 0.9973099 |
| R-I/L | -0.1161311072 | -0.233753318 | 0.001491104 | 0.0575256 |
| S-I/L | -0.0667992823 | -0.180293104 | 0.046694540 | 0.8393481 |
| T-I/L | 0.0266558529 | -0.087302158 | 0.140613864 | 0.9999969 |
| V-I/L | 0.0145017209 | -0.098992101 | 0.127995543 | 1.0000000 |
| W-I/L | -0.1607935128 | -0.274287335 | -0.047299691 | 0.0001223 |
| Y-I/L | -0.1447016834 | -0.258195505 | -0.031207861 | 0.0012503 |
| M-K | 0.0706761109 | -0.042817711 | 0.184169933 | 0.7680902 |
| N-K | 0.0944380858 | -0.019055736 | 0.207931908 | 0.2485655 |
| P-K | 0.1302672716 | 0.016773450 | 0.243761093 | 0.0079620 |
| Q-K | 0.0538203414 | -0.059673480 | 0.167314163 | 0.9735982 |
| R-K | -0.0186979398 | -0.136320151 | 0.098924271 | 1.0000000 |
| S-K | 0.0306338851 | -0.082859937 | 0.144127707 | 0.9999748 |
| T-K | 0.1240890203 | 0.010131009 | 0.238047031 | 0.0172853 |
| V-K | 0.1119348883 | -0.001558934 | 0.225428710 | 0.0581955 |
| W-K | -0.0633603454 | -0.176854167 | 0.050133476 | 0.8906427 |
| Y-K | -0.0472685160 | -0.160762338 | 0.066225306 | 0.9931985 |
| N-M | 0.0237619749 | -0.089731847 | 0.137255797 | 0.9999994 |
| P-M | 0.0595911607 | -0.053902661 | 0.173084983 | 0.9332725 |
| Q-M | -0.0168557695 | -0.130349591 | 0.096638052 | 1.0000000 |
| R-M | -0.0893740507 | -0.206996261 | 0.028248160 | 0.4120489 |
| S-M | -0.0400422258 | -0.153536048 | 0.073451596 | 0.9990562 |
| T-M | 0.0534129093 | -0.060545102 | 0.167370921 | 0.9764516 |
| V-M | 0.0412587774 | -0.072235044 | 0.154752599 | 0.9986283 |
| W-M | -0.1340364563 | -0.247530278 | -0.020542634 | 0.0050226 |

| | | | | |
|---|---|---|---|---|
| Y-M | -0.1179446269 | -0.231438449 | -0.004450805 | 0.0318220 |
| P-N | 0.0358291858 | -0.077664636 | 0.149323008 | 0.9997782 |
| Q-N | -0.0406177444 | -0.154111566 | 0.072876078 | 0.9988710 |
| R-N | -0.1131360256 | -0.230758236 | 0.004486185 | 0.0755597 |
| S-N | -0.0638042007 | -0.177298023 | 0.049689621 | 0.8846887 |
| T-N | 0.0296509344 | -0.084307077 | 0.143608946 | 0.9999852 |
| V-N | 0.0174968025 | -0.095997019 | 0.130990624 | 1.0000000 |
| W-N | -0.1577984312 | -0.271292253 | -0.044304609 | 0.0001923 |
| Y-N | -0.1417066018 | -0.255200424 | -0.028212780 | 0.0018710 |
| Q-P | -0.0764469302 | -0.189940752 | 0.037046892 | 0.6418642 |
| R-P | -0.1489652114 | -0.266587422 | -0.031343001 | 0.0014252 |
| S-P | -0.0996333865 | -0.213127208 | 0.013860435 | 0.1699198 |
| T-P | -0.0061782513 | -0.120136263 | 0.107779760 | 1.0000000 |
| V-P | -0.0183323833 | -0.131826205 | 0.095161439 | 1.0000000 |
| W-P | -0.1936276170 | -0.307121439 | -0.080133795 | 0.0000005 |
| Y-P | -0.1775357876 | -0.291029609 | -0.064041966 | 0.0000083 |
| R-Q | -0.0725182812 | -0.190140492 | 0.045103930 | 0.7819764 |
| S-Q | -0.0231864563 | -0.136680278 | 0.090307366 | 0.9999996 |
| T-Q | 0.0702686788 | -0.043689332 | 0.184226690 | 0.7817920 |
| V-Q | 0.0581145469 | -0.055379275 | 0.171608369 | 0.9462545 |
| W-Q | -0.1171806869 | -0.230674509 | -0.003686865 | 0.0344542 |
| Y-Q | -0.1010888574 | -0.214582679 | 0.012404964 | 0.1515292 |
| S-R | 0.0493318249 | -0.068290386 | 0.166954036 | 0.9926493 |
| T-R | 0.1427869601 | 0.024716789 | 0.260857131 | 0.0033464 |
| V-R | 0.1306328281 | 0.013010617 | 0.248255039 | 0.0130714 |
| W-R | -0.0446624056 | -0.162284616 | 0.072959805 | 0.9976678 |
| Y-R | -0.0285705762 | -0.146192787 | 0.089051635 | 0.9999946 |
| T-S | 0.0934551351 | -0.020502876 | 0.207413146 | 0.2725897 |
| V-S | 0.0813010032 | -0.032192819 | 0.194794825 | 0.5267417 |
| W-S | -0.0939942305 | -0.207488052 | 0.019499591 | 0.2562335 |
| Y-S | -0.0779024011 | -0.191396223 | 0.035591421 | 0.6076844 |
| V-T | -0.0121541319 | -0.126112143 | 0.101803879 | 1.0000000 |
| W-T | -0.1874493657 | -0.301407377 | -0.073491354 | 0.0000017 |
| Y-T | -0.1713575362 | -0.285315547 | -0.057399525 | 0.0000260 |
| W-V | -0.1752952337 | -0.288789056 | -0.061801412 | 0.0000121 |
| Y-V | -0.1592034043 | -0.272697226 | -0.045709582 | 0.0001557 |
| Y-W | 0.0160918295 | -0.097401992 | 0.129585651 | 1.0000000 |

**G Peptides with either an azide or PEG functionalization at 48 hours**

| 48 hours | Df | Sum Sq | Mean Sq | F value | Pr(>F) |
|---|---|---|---|---|---|
| celltype | 2 | 0.610 | 0.3050 | 8.218 | 0.000459 *** |
| endgroup | 6 | 4.996 | 0.8326 | 22.435 | <2e-16 *** |
| functionalization | 1 | 2.784 | 2.7837 | 75.004 | 3.19e-14 *** |
| residuals | 116 | 4.305 | 0.0371 | | |

Tukey multiple comparisons of means
95% family-wise confidence level
Fit: aov(formula = ave ~ cell + endgroup + functionalization)

| **Cell type** | diff | lwr | upr | p adj |
|---|---|---|---|---|
| hUVEC-hMSC. | 0.09160999 | -0.008199751 | 0.1914197 | 0.0790515 |
| mac-hMSC | 0.17027349 | 0.070463754 | 0.2700832 | 0.0002719 |
| mac-hUVEC. | 0.07866350 | -0.021146233 | 0.1784732 | 0.1516898 |

| **Endgroup** | diff | lwr | upr | p adj |
|---|---|---|---|---|
| AcBA-Ac | 0.003644893 | -0.18905250 | 0.196342286 | 1.0000000 |
| Am-Ac | -0.147102880 | -0.33980027 | 0.045594513 | 0.2574162 |
| CBA-Ac | 0.069312949 | -0.12338444 | 0.262010342 | 0.9329731 |
| COOH-Ac | -0.208820203 | -0.40151760 | -0.016122810 | 0.0245513 |
| NBA-Ac | -0.193966891 | -0.38666428 | -0.001269498 | 0.0473857 |
| NH2-Ac | -0.571832555 | -0.76452995 | -0.379135162 | 0.0000000 |
| Am-AcBA | -0.150747773 | -0.34344517 | 0.041949620 | 0.2308972 |
| CBA-AcBA | 0.065668056 | -0.12702934 | 0.258365449 | 0.9478292 |
| COOH-AcBA | -0.212465097 | -0.40516249 | -0.019767704 | 0.0207275 |
| NBA-AcBA. | -0.197611784 | -0.39030918 | -0.004914391 | 0.0405243 |
| NH2-AcBA | -0.575477449 | -0.76817484 | -0.382780056 | 0.0000000 |
| CBA-Am | 0.216415829 | 0.02371844 | 0.409113222 | 0.0171938 |
| COOH-Am | -0.061717323 | -0.25441472 | 0.130980070 | 0.9612073 |
| NBA-Am | -0.046864011 | -0.23956140 | 0.145833382 | 0.9904628 |
| NH2-Am | -0.424729675 | -0.61742707 | -0.232032282 | 0.0000000 |
| COOH-CBA | -0.278133152 | -0.47083055 | -0.085435759 | 0.0006162 |
| NBA-CBA | -0.263279840 | -0.45597723 | -0.070582447 | 0.0014642 |
| NH2-CBA | -0.641145504 | -0.83384290 | -0.448448111 | 0.0000000 |
| NBA-COOH | 0.014853312 | -0.17784408 | 0.207550705 | 0.9999867 |
| NH2-COOH | -0.363012352 | -0.55570975 | -0.170314959 | 0.0000024 |
| NH2-NBA | -0.377865664 | -0.57056306 | -0.185168271 | 0.0000008 |

| **Functionalization** | diff | lwr | upr. | p adj |
|---|---|---|---|---|
| PEG-azide | 0.2972747 | 0.2292891 | 0.3652603 | 0 |

**Figure S24**. Statistical analyses using multi-way ANOVAs with a Turkey post-hoc test. Three technical replicates were averaged to a single values, and statistics were done on experimental/biological replicates. Statistical analysis on a) soluble peptides cultured with cells on tissue culture plastic, all time points b) soluble peptides cultured with cells on tissue culture plastic, 48 hours, c) concentration study peptides at 48 hours, d) LIAANK peptides at 48 hours, e) IVKVA peptides at 48 hours, f) soluble peptides cultured with cells in PEG hydrogels at 48 hours, g) azide/PEG functionalized peptides cultured with cells at 48 hours.
